# Human Glycolysis Isomerases are Inhibited by Weak Metabolite Modulators

**DOI:** 10.1101/2024.09.26.610356

**Authors:** Yiming Yang Jónatansdóttir, Óttar Rolfsson, Jens G. Hjörleifsson

## Abstract

Modulation of enzyme activity by metabolites represents the most efficient and rapid way of controlling metabolism. Investigating enzyme-metabolite interactions can deepen our understanding of metabolic control and aid in identifying enzyme modulators with potential therapeutic applications. These interactions vary in strength, with dissociation constants (K_d_) ranging from strong (nM) to weak (µM-mM). However, weak interactions are often overlooked due to the challenges in studying them. Despite this, weak modulators can reveal novel binding modes and serve as starting points for compound optimization. In this study, we aim to identify metabolites that weakly modulate the activity of human glucose-6-phosphate isomerase (GPI) and triosephosphate isomerase (TPI), which are potential therapeutic targets in tumor glycolysis. Through a combination of activity and binding assays, the screening revealed multiple weak inhibitors for the two targets, causing partial attenuation of their activity, with K*_d_* and K*_i_* in the low mM range. X-ray crystallography revealed six orthosteric ligands binding to the active sites – four inhibitors of GPI and two of TPI. Our findings underscore the role of weak interactions in enzyme regulation and may provide structural insights that could guide the design of inhibitors targeting human GPI and TPI in cancer intervention.

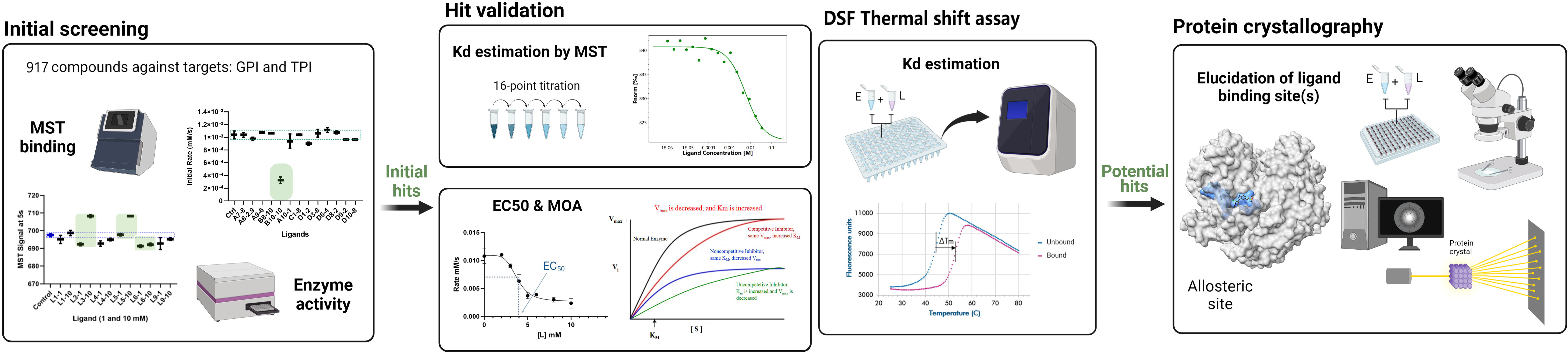

## Introduction

Enzymes are embedded in a sea of metabolites, which may potentially interact with them and modulate their activities. Finding new modulatory interactions has profound importance in the study of biological processes, and it has become an indispensable tactic for the discovery of drugs with novel properties. It can be presumed that all enzymes have one-to-few regulatory sites that are hidden under steady state or in the absence of effector molecules, and they can be occupied when certain metabolites are abnormally enhanced, or when appropriate exogenous binders are present. In general, intracellular metabolites interact with their enzyme partners with an affinity (K_d_) close to their physiological concentrations, which are often in the low-micromolar to - millimolar range [1]. Despite being weak, these interactions can be additive or synergistic, and have substantial influence on pathway behavior [2], resulting in striking differences compared with the somewhat less probable case, wherein no modulatory effect is present at all.

The interactions between enzymes and metabolites can range from strong (K_d_ ≤ nM) to weak (K_d_ = µM-mM). Weak interactions are commonplace in nature, but they are often regarded to be non-specific, as weak binders have been envisioned to be lack of selectivity, exhibiting cross reactivity across multiple targets. Besides, low affinity makes it difficult to detect and characterize weak interactions and their functional effects. The concentration of the screened binders must be high enough such that sufficient occupancy can be achieved to observe binding events or biological response. This may increase the incidence of assay interference and pose solubility issues for certain compounds in aqueous buffers, leading to aggregate formation. Consequently, for a long time, high binding affinity has been the prioritized criterion for identifying promising hits and lead candidates, whereas weak, transient interactions have been largely ignored. Even though weak binders may not be as attractive as strong ones, they still have the potential to probe for new binding sites (subsites) and modes of interactions with targets if structural information can be obtained. In addition, the initial binding affinity of weak binders does not necessarily reflect the potency of mature compounds. Any weak binders may be of high value for subsequent compound optimization and to establish structure-activity relationships.

In this study, we set out to identify weak modulators of two human isomerases in the glycolytic pathway: glucose-6-phosphate isomerase (GPI; EC 5.3.1.9) and triosephosphate isomerase (TPI; EC 5.3.1.1). As a universal metabolic pathway found in all domains of life, glycolysis serves in the critical role of generating fast energy and supplying biosynthetic precursors that are essential for anabolic processes. While glycolysis upregulation is often associated with cancer development [3,4], impairment of glycolytic activity in non-cancerous cells can result in pathological conditions such as hemolytic anemia and neurodegenerative diseases [5,6]. The glycolytic pathway is tightly regulated by feedforward and feedback mechanisms, involving both allosteric and orthosteric modulators [7,8]. Each enzyme in the pathway represents a regulatory point that can be potentially exploited as a therapeutic target for the intervention of diseases related to glycolysis dysfunction or dysregulation.

GPI and TPI both work at a metabolic crossroad, ensuring rapid equilibration of the hexose-phosphates and triosephosphates, respectively. The former catalyzes interconversion of glucose-6-phosphate (G6P) and fructose-6-phosphate (F6P) in the 2^nd^ step of glycolysis, while the latter interconverts dihydroxyacetone phosphate (DHAP) to glyceraldehyde-3-phosphate (GAP) in the 5^th^ step of glycolysis. These four metabolic intermediates directly bridge glycolysis to various other pathways of the central metabolism, including pentose phosphate pathway (PPP), hexosamine biosynthesis, and triglyceride synthesis. Upregulation of GPI and TPI enzyme activities has been implicated in the pathogenesis of various cancers that display addiction to glycolysis, generally referred to as the Warburg effect [4,9–13]. Although seemingly rare, GPI and TPI deficiencies are two of the most severe inborn enzymopathies that manifest as chronic hemolytic anemia and neurological dysfunction, frequently leading to death in early childhood [6,14].

Over the past years, extensive research efforts have been made on targeting glycolysis enzymes to combat tumor development and parasitic infections in humans. It has been shown that activity impairment of GPI and TPI could severely reduce glycolytic flux, and inactivation of GPI has suppressing effects on tumor glycolysis and growth [5,15,16]. But owing to their low ligandability, GPI and TPI have received little attention as therapeutic targets compared with other glycolysis enzymes, such as hexokinase (HK), phosphofructokinase (PFK), and pyruvate kinase (PK) [17,18]. Very few studies have been undertaken for the identification of GPI modulators in the past years [19,20]. Although various screening studies of TPI modulators have been conducted [21–27], they were mostly dedicated to TPI of non-human origins. All the previously known modulators of human GPI and TPI exhibit moderately strong inhibition on the enzymes’ activity, with IC_50_ in the lower-to-upper micromolar range. As compound libraries are conventionally screened at sub-micromolar concentrations or lower, any weak binders can be easily washed out, including those that serve modulatory functions. To identify weak activity modulators for human GPI and TPI, we performed compound screening at micromolar to low-millimolar concentrations. The enzyme targets were screened with 917 compounds (Mw < 600 kDa) comprising a wide range of metabolites found in human tissues and biofluids. A combination of affinity-based and enzyme kinetic assays was used for hit identification, followed by structural validation of the putative hits with X-ray crystallography.

## Methods

### Materials

All chemicals for buffer preparation were purchased from Sigma-Aldrich (USA) and Merk (USA). Culture media components include L-Rhamnose and isopropyl β-D-1-thiogalactopyranoside (IPTG) derived from AppliChem GmbH (Germany); Yeast extract and Casein peptone from Neogen (USA); Bacto agar from Becton, Dickinson (USA); Ampicillin and Chloramphenicol from AppliChem (Germany). The metabolite compound library was purchased from MetaSci (Canada) and a small portion of test compounds from Sigma (USA). Rabbit glyceraldehyde-3-phospahte dehydrogenase (GAPDH; EC 1.2.1.12) and protein standards for molecular weight determination were obtained from Sigma-Aldrich (USA). The luminescence NAD(P)H-Glo™ Detection System (assay kit) was purchased from Promega (USA). Protein Thermal Shift™ Dye Kit (no. 4461146) was obtained from ThermoFisher (USA). FrameStar® 96 Well Semi-Skirted PCR Plate (ABI® FastPlate Style) and qPCR Adhesive Seal were obtained from AZENTA (USA). Crystallization screens – Shotgun (SG) and JCSG-plus™ and other crystallization precipitant and buffer solutions were purchased from Molecular Dimensions (UK). Crystallization tools and silicon grease were derived from Hampton Research (USA). Hulec-5a (Human Lung Microvascular Endothelial cells; CRL-3244™) and U2OS (human osteosarcoma cell line; HTB-96™) were derived from ATCC® cultures (USA).

### Expression and purification of recombinant GPI, TPI, and G6PDH from *E. coli*

The target human enzymes, GPI and TPI, and coupling assay enzyme, glucose-6-phosphate dehydrogenase (G6PDH; EC 1.1.1.49) were expressed in Lemo21(DE3) competent *E. coli* cells purchased from New England Biolabs. The Lemo21(DE3) cells were transformed with pET11a plasmids (all genes subcloned via Ndel/BamHI) carrying the proteins of interest with 6xHis-Tag at N-terminus for TPI and C-terminus for GPI and G6PDH) obtained via GenScript (USA) cDNA ORF cloning services (NM_000365.6 and NM_001329909.1 accession no. for TPI and GPI respectively) or gene synthesis (yeast G6PDH Uniprot ID P11412, codon optimized for *E. coli*). The culture and expression procedures are detailed in supplementary data A.

The protein purification was conducted on an ÄKTA Pure system (Cytiva), using 5 mL HisTrap FF column (Cytiva) at room temperature. A constant flow rate of 5 mL/min was used throughout the purification process. Initially, the column was equilibrated with Buffer A (25 mM Tris pH 7.9, 250 mM NaCl, 1 mM 2-mercaptoethanol, 10 mM imidazole). After loading the protein sample, the column was washed with Buffer A to remove the unbound species. For elution of the His-tagged proteins, an increasing gradient of imidazole (10-100% Buffer B) was employed for 15 minutes, with Buffer B (25 mM Tris pH 7.9, 250 mM NaCl, 1 mM 2-mercaptoethanol, 500 mM imidazole). Re-equilibration of the column was done using Buffer A. The purified proteins were buffer-exchanged and concentrated to 10-40 mg/mL using Merck Amicon™ Ultra-15 Centrifugal Filter Units with a 30 kDa cut-off, from Fisher Scientific (USA). The protein concentrations were determined by A280 measurements on the NanoDrop 2000 Spectrophotometer (Thermo Scientific). All purified proteins were stored at –80°C in Tris-HCl buffer (50 mM Tris pH 7.4, 150 mM NaCl).

### Chemical library and compound preparation

The metabolite library comprising 1250 compounds was purchased from MetaSci (Canada) and received in solid or pure liquid forms. Another smaller custom library consisting of 21 metabolites was obtained from Sigma (USA). All compounds are listed in supplementary data B. A total of 941 compounds were manually reconstituted up to their solubility limit in water, without or with minimum amount of DMSO (≤ 4%). Poorly water-soluble compounds, such as fatty lipids and sterols, were not included in the screening. The pH of the reconstituted compounds was adjusted to around 7.4 by adding HCl or NaOH. All compound stocks and diluted solutions were stored at −80 °C. Due to crystallization or precipitations after freeze-thawing, some compounds were redissolved by sonication in water bath for 30-60 minutes, at ambient temperature. Prior to assays, all compounds were pH-checked with pH strips and the final DMSO% in the assay reagent was kept as low as 4% to avoid protein destabilization. Of the solubilized compounds, 917 were subjected to screening, and 24 compounds were either too intensely colored for absorbance assay or highly prone to precipitation, and thus had to be excluded. See solubility guide Appendix A for further details.

### MST binding assays

The microscale thermophoresis (MST) method [28] was employed to study the binding interactions between enzyme targets and metabolites. Binding assays were performed on a NT.LabelFree Monolith instrument (NanoTemper) using Monolith NT.LabelFree capillaries from NanoTemper GmbH (Germany). All screened chemicals were prepared in MST buffer (100 mM Tris pH 7.4, 150 mM NaCl, and 1 mM DTT). After dilution, the compounds were sonicated in water bath for 30-60 min to ensure no visible particulates nor crystalline solids were present. Before screening, the target enzyme was centrifuged at 10000 × g for 5 min and added to the compound solution in the same buffer before being centrifuged at 5000 × g for 2 min, followed by equilibration at ambient temperature for 10 minutes. The screening was conducted at 25°C, using 20% excitation and 40% MST power, with on-time response set to 5 or 10 seconds, depending on the quality of the MST traces. The enzyme concentration was kept constant at dimeric concentration of 0.5 μM for both GPI and TPI in all runs.

Considering the transient and low-affinity nature of metabolite-enzyme interactions, the ligand stock concentrations were set as high as possible (100 µM–50 mM), depending on their solubility and DMSO concentration. For primary screening, the compounds were screened at two different concentrations, in the 10 μM–10 mM range. Compounds resulting in changes in MST signal > 5 units w.r.t the control were regarded as primary hits. To obtain full binding curves (12-point assay) for K_d_ estimation, 2/3 serial dilutions of selected hits were titrated against a constant concentration of enzyme (0.5 μM), with ligand concentration starting from 10-50 mM, depending on ligand solubility in water. The test compounds were assayed in duplicates in the primary screening and at least triplicates in the full 12 point-titration experiments. The MST binding assay for the enzyme targets was validated by using 6-phosphogluconate (6PGc) and phosphoenolpyruvate (PEP) – competitive inhibitors and binders of GPI and TPI, respectively (Fig. 2S-1 & 2S-2). SDS denaturation (SD) tests were performed in the event of variation in initial fluorescence to discriminate between binding-specific and ligand-induced fluorescence increase or quenching. Prior to SD tests, mixtures containing unbound and bound states of the target were centrifuged at 10000 × g for 5 min, and 7 µL of sample was transferred into a 2X solution containing 4% SDS and 40 mM DTT, and then heated at 95° for 5 min.

### Enzyme activity assays

Activity assays were employed for the initial screening and determination of EC_50_ values and mechanism of action (MOA). The activity of GPI was measured in the direction of F6P to G6P, and TPI in the direction of DHAP to GAP, using both colorimetric and luminescent kinetic assays (Fig. 1). The colorimetric kinetic assays involved coupling the GPI- and TPI-catalyzed reactions to the conversion of NADP+ to NADPH, catalyzed by glucose-6-phosphate dehydrogenase (G6PDH), and reduction of NAD+ to NADH by glyceraldehyde-3-phosphate dehydrogenase (GAPDH), respectively. The test compounds were screened at a concentration range between 20 µM to 10 mM, depending on their water solubility. The rate of GPI- and TPI-catalyzed reactions was measured by monitoring the formation of NAD(P)H at 340 nm. Due to absorption around 340 nm, a portion of compounds (149 in total) were incompatible with the colorimetric assay due to absorbance interference. These near UV-active compounds were instead screened with luminescent assays using the NAD(P)H-Glo™ Detection System (Promega), which monitors the formation of NAD(P)H with the aid of two coupling enzymes, reductase and luciferase, in addition to the dehydrogenases mentioned above. Both the colorimetric and luminescent enzyme assays were performed in 96- and 384-well plate formats, with assay volumes of 60 µL and 30 µL, respectively, using an iD5 multi-mode microplate reader (Molecular Devices). Validation of the activity assays was made using known GPI and TPI inhibitors, 6PGc and PEP, respectively (Fig. 2S-3 & 2S-4). DMSO tolerance of the target enzymes was tested by DMSO titration, from 0.1%-10%. Both enzymes retained their activity and tolerated up to ∼5% DMSO, with less than 10% activity reduction (Table S1). For detailed assay conditions and experimental setups see supplementary data A.

**Figure 1:**
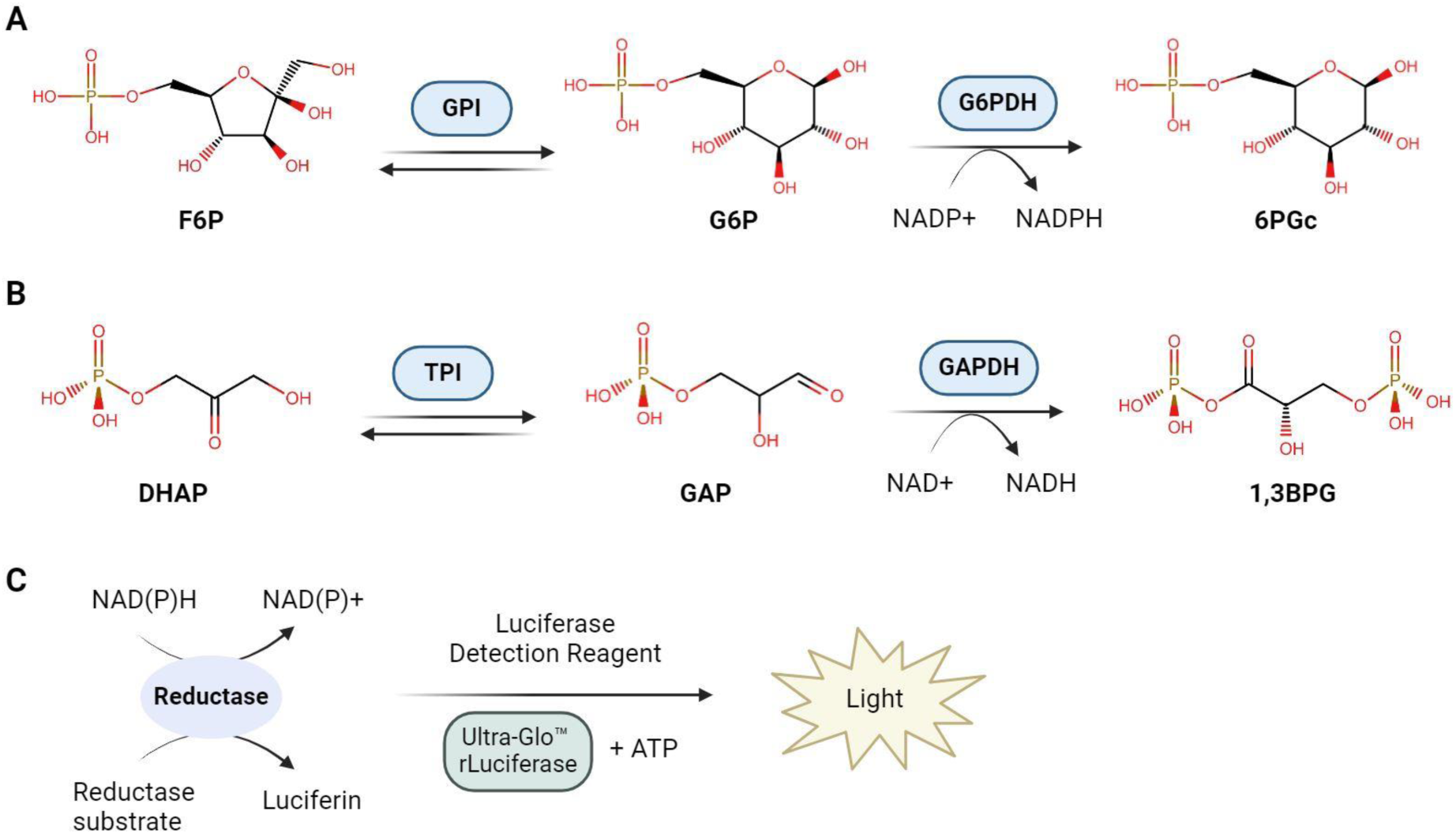
The scheme of enzyme-coupled reactions in colorimetric (A & B) and luminescence-based (C) enzymatic assay. (A) GPI-catalyzed reaction coupled to the reduction of NADP+ to NADPH by G6PDH. (B) TPI-catalyzed reaction coupled to the reduction of NAD+ to NADH by GAPDH. The increase in NAD(P)H absorbance is continuously monitored at 340 nm. (C) NAD(P)H-Glo detection system; utilizes luciferin accumulation followed by its oxidation by luciferase to quantify the formation of NAD(P)H. In the presence of NAD(P)H, a reductase reduces the proluciferin reductase substrate to form luciferin, which is further oxidized by luciferase to produce light signal that is proportional to the amount of NAD(P)H present in the sample.

### Thermal shift assay

Differential scanning fluorometry (DSF) was used as an orthogonal assay for K_d_ estimation of the potential hits. The thermal shift assay was carried out in 96-well qPCR plates, with an assay volume of 20 µL. The fluorescence measurement was performed with QuantStudio™ 3 Real-Time PCR System (ThermoFisher, USA), using a fluorescent reporter, Rox dye (Thermo Scientific), which has an excitation wavelength of 580 ± 10 nm and emission wavelength of 623 ± 14 nm. Assay reagents were comprised of 0.5 µM of enzyme target, 0-40 mM of test compounds, MST buffer supplemented with 0.001% (w/v) Triton X-100, and 1X Rox dye. The ligand binding assays were conducted following the manufacturers’ protocol. Mixtures of enzymes and ligands of varying concentrations (0-45 mM) were incubated at 4°C for 10 minutes, followed by the addition of the dye, and then centrifugation of the assay mixtures at 3000xg and 4°C for 3 minutes. Prior to measurement, the assay mixtures were incubated at 25°C for 2 minutes in the PCR machine, and the change in fluorescence was monitored at a temperature range from 25 to 85°C, with an increment rate of 0.05°C/s. Validation of the assay was made using GTP and PEP – known inhibitors of GPI and TPI, respectively. No background signal was observed when the test compounds were assayed in the absence of the enzymes. Using GraphPad Prism, the first derivatives of the relative fluorescence unit (RFU) as a function of temperature were obtained from the melt curve data. The midpoint of thermal denaturation (T_m_) was approximated by fitting the first derivative curves in a defined range with quadratic polynomials (f(x) = ax^2^ + bx + c) and finding the vertex (x = –b/2a) of the quadratic function.

### Protein X-ray crystallography

Co-crystallization and soaking experiments were performed to obtain enzyme-ligand complexes. GPI and TPI crystals were grown at ambient temperature (25°C) using hanging-drop diffusion. Initial screens, SG and JCSG+, (Molecular Dimensions, UK) were performed for both enzymes to identify suitable crystallization conditions. Prior to crystallization, all protein samples and ligand solutions were passed through 0.22 µm filters. The GPI and TPI samples were stored in 25 mM Tris and 50 mM NaCl, pH 7.4. The best GPI crystals were produced in conditions containing 20-23% PEG3350, 0.16 M CaCl_2_, and 58 mM HEPES, pH 7.0, whereas for TPI, the crystals grew optimally under a condition containing 0.14 M KBr and 24% PEG 2000 MME. In both cases, micro-seeding was necessary to obtain suitably large, single crystals. Crystal drops were prepared by mixing 1.5 µL of reservoir solution with 1.5 µL of 8 mg/mL GPI or 10 mg/mL TPI solution (with or without ligand), and 0.5 µL crystal seeds of corresponding protein.

For ligand co-crystallization, the target enzymes were pre-incubated with 10-25 mM ligand at 4°C for 30 minutes before being mixed with the reservoir and seed solutions. After 4-5 days of incubation, single crystals of medium sizes (100-200 µm) were harvested and flash-frozen after dipping into a cryoprotectant mixture containing mother liquor, 20% glycerol, and 10-30 mM ligand. For ligands that did not produce visible crystals, soaking experiments were performed using the apo crystals, which were dipped into the cryoprotectant mixture mentioned above. The results of co-crystallization trials are detailed in Fig. 8S in supplementary data A. The length of soaking time was between 1-10 min, depending on the tolerance of the protein crystals. To prevent dehydration, the coverslips were sealed during soaking. The crystals were harvested if no cracking was visualized after soaking. The harvested crystals were snap-frozen and sent to MAX-IV synchrotron facility in Lund for X-ray diffraction experiments. The diffraction data were collected up to 1.1-1.8 Å resolution for the protein-ligand complex structures submitted to the PDB. The X-ray data obtained for GPI crystals were processed in space group P_21 21 21_ for GPI, with cell dimensions a = ∼81.0 Å, b = ∼105.0 Å, and c = ∼272.0 Å. The TPI diffraction data were processed in P_61 2 2_, with cell dimensions a = ∼48.5 Å, b = ∼48.5 Å, and c = ∼342.0 Å. Diffraction data were processed in XDS [29] and scaled in Aimless within the CCP4 program suite [30]. Initial estimation of diffraction data quality was carried out in XTRIAGE [31] and phases solved using molecular replacement (MR), in Phaser [32] for GPI and MoRDa [33] for TPI, using PDB accession numbers 1JLH and 2JK2 respectively as reference models. Model refinement was carried out in phenix.refine [34] within the Phenix software suite (v. 1.20.1-4487), and manual model building in COOT [35].

### Metabolic flux and cell viability assays

The effect of PEP on cellular glycolytic flux and cell viability was examined for two human cell lines – Hulec-5a and U2OS. Real-time measurement of extracellular acidification rate (ECAR) was conducted using the Agilent Seahorse XFe96 analyzer. Two days before the assay, the HULEC-5a and U2OS cells were separately seeded at a density of ∼40.000 cells/100 μL in Seahorse cell culture plates and incubated at 37°C, 5% v/v CO_2_. 24 hours prior to measurement, sensor plates were hydrated with the Agilent Seahorse Calibrant and incubated at 37°C, 0% CO_2_. On the day of measurement, 8 μL of PrestoBlue reagent was added into each well of the culture plate, which was then incubated for 1 hour at 37°C, 5% CO_2_, followed by the removal of the media for absorbance measurement and cell viability normalization. After removing the media and rinsing the cells with 1x PBS, 180 μL of Agilent DMEM Running media (5 mM HEPES pH 7.4, 10 mM glucose, 1 mM pyruvate, and 2 mM glutamine) was dispensed into each well, and then the culture plate was incubated at 37°C, 0% CO2 for 1 hour. The sensor cartridge ports were loaded with test compounds and inhibitors to reach a final assay volume of 250 µL. The components added and their final concentrations were as follows: test compounds – 5-25 mM PEP (pH-adjusted to 7.4) and assay inhibitors – 5 µM rotenone/antimycin (Rot/AA) and 50 mM 2-Deoxy-glucose (2-DG); all compounds were dissolved in DMEM media. For positive and negative controls, 5-20 mM 6PGc (GPI inhibitor) and DMEM media were utilized, respectively. After calibration of the sensor cartridge, the assay was run 37°C and five measurements were taken after each injection. Six replicates were taken for each condition. For cell viability normalization, the media with PrestoBlue reagent (Invitrogen™) were measured for absorbance at 570 nm and 600 nm. Proton efflux rate (PER) was calculated from the measured ECAR and used as an approximation of the glycolytic flux rate.

The cell viability assays were conducted using PrestoBlue reagent as a cell viability indicator. Prior to assays, the HULEC-5a and U2OS cells were seeded at a density of ∼5000 cells/100 μL in Nunc™ MicroWell™ 96-Well Microplates (ThermoFisher, USA), to reach 50% confluency to ensure optimal compound and dye uptake. The HULEC-5a and U2OS cells were cultured at 37°C, 5% CO_2_, in DMEM-GlutaMAX^TM^ (Gibco) supplemented with 10% FBS. The viability of the cells was monitored over a period of three days by absorbance measurement at 570 nm before and after 4, 8, 24, 48, and 68 hours from the first day of compound treatment. The cells were divided into three groups: the experimental groups were treated with 10, 20, and 30 mM of PEP, whereas the positive and negative control groups were fed with 30 mM 2-deoxyglucose (2-DG; hexokinase inhibitor) and plain media, respectively. For each condition, six replicates were prepared and measured. Prior to the absorbance measurement, the cells were treated with 5 µL PrestoBlue reagent and incubated for 1 hour at 37°C, 5% CO_2_, followed by transfer of the media into 96-well microplates. After removing the media, the cells were quickly rinsed with 100 uL Gibco™ PBS solution and supplied with 100 µL of fresh media containing their respective treatments. All media, reagent, and buffer solutions were prewarmed at 37°C before feeding to the cells. The absorbance measurement at 570 nm was conducted using the iD5 multi-mode microplate reader (Molecular Devices).

## Results

### Metabolite screening revealed several weak modulators of GPI and TPI

A total of 917 water-soluble metabolites were screened against human GPI and TPI. The results from primary and secondary screening are detailed in supplementary data A and B, and the hit selection strategy is illustrated in Fig 2. A low threshold was set to the initial screens to coarsely identify all possible weak modulators. All compounds that caused changes in activity ≥ 25% and/or ≥ 5 MST F_norm_ (‰) units in the initial screens were regarded as primary hits. The initial screening performed for GPI resulted in 123 activity- and 174 MST binding hits, 30 of which showed up as primary hits in both activity and MST assays. The initial screening against TPI generated 135 activity assay- and 186 MST hits, 45 of which were both activity assay and MST hits. Of the primary activity hits, 78 GPI – and 98 TPI hits were derived from the colorimetric kinetic assay; the remaining 45 GPI – and 37 TPI hits (of 149 screened) were obtained from the luminescence assay. To exclude false positives, the primary hits were screened against the coupling enzymes at two ligand concentrations (0.1-10 mM) using the activity assays. Compounds that altered the coupling enzyme activity < 15% were then re-assessed (at two concentrations) against the target enzymes in secondary screens. Compounds showing a concentration-dependent effect on the targets were defined as secondary hits and further examined.

**Figure 2:**
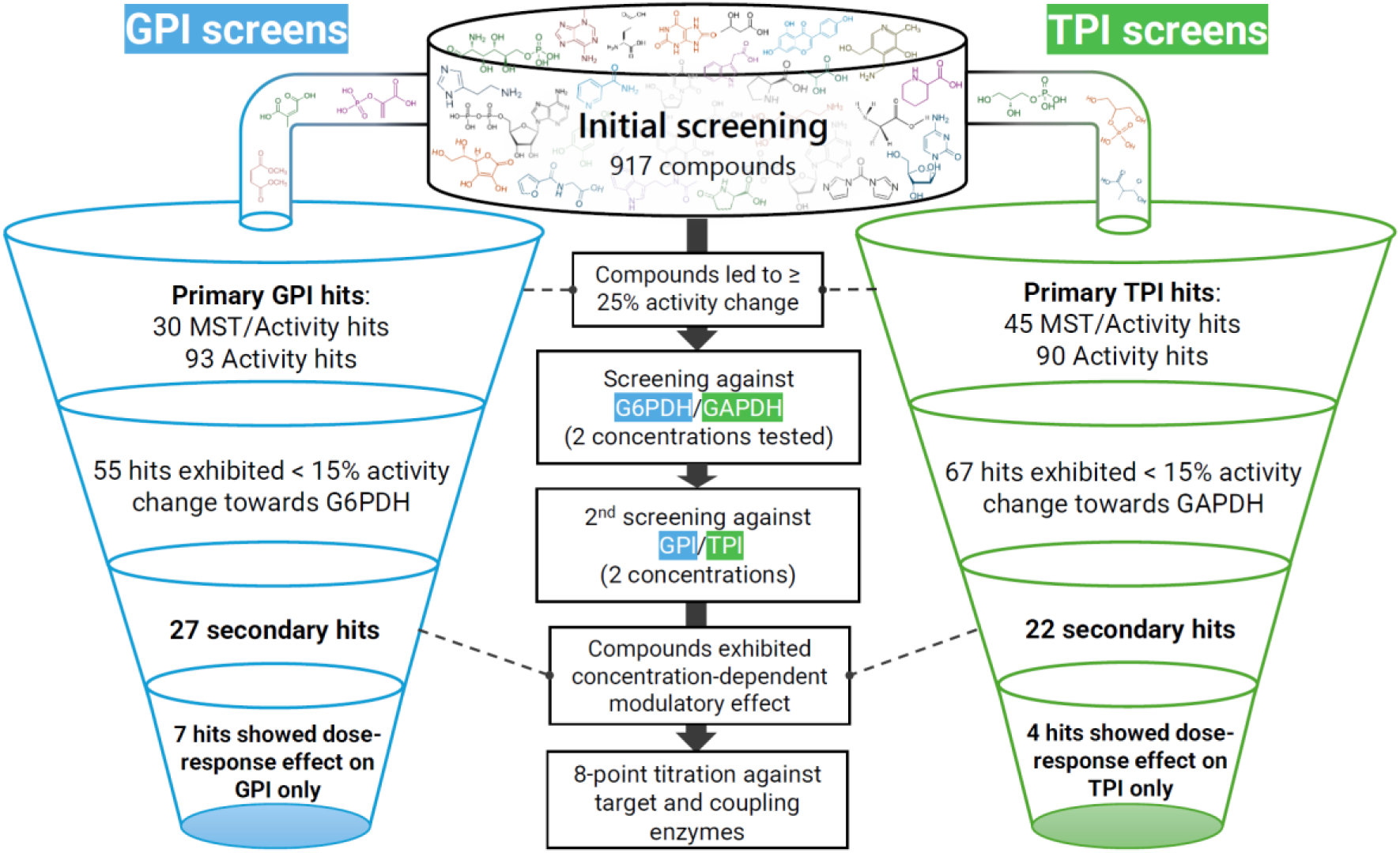
Screening and hit selection strategy for identifying potential GPI and TPI modulators. The GPI- and TPI-catalyzed reactions were coupled to NADP+/NADH+ reduction catalyzed by G6PDH and GAPDH, respectively. Primary activity hits: compounds that caused ≥ 25% and 30% activity change in absorbance and luminescent activity assays, respectively. Primary MST hits: compounds that altered MST signal by at least 5 F_norm_ (‰) units for at least one test concentration. Primary hits were reassessed against the coupling enzymes (G6PDH and GAPDH) and target enzymes (GPI and TPI) at two ligand concentrations (0.1–10 mM range, depending on solubility). Secondary hits: compounds that exhibited < 15% activity change towards GAPDH while showing concentration-dependent effect on target activity. 8-point titration of the secondary hits (2-fold serial dilution from 24-30 mM) was conducted against both target and coupling enzymes. 7 GPI and 4 TPI hits showed dose-response inhibition of the targets but having almost no effect on the coupling enzyme activity.

Altogether, there were 7 compounds showing weak dose response inhibition on GPI activity, and 4 compounds that exhibited weak inhibitory effects towards TPI in dose response manner. The EC_50_, K_i_, and K_d_ values of all identified GPI and TPI hits, and their potential mode of action (MOA) and binding sites (derived by X-ray crystallography) are summarized in Table 1 and Table 2. All identified hits, except for glycerol-3-phosphate (G3P), are considered novel with respect to the human GPI and TPI. G3P was included for further assessment as there were no binding and crystallographic studies conducted for this TPI ligand previously. Apart from the GPI and TPI hits, there were multiple compounds that displayed apparent inhibition towards the coupling enzymes – yeast G6PDH and rabbit GAPDH (Fig. 5S-2 & 5S-4).

**Table 1:** Summary of identified GPI inhibitors. The EC_50_ and K_d_ values are expressed as mean ± standard deviation of two duplicates and triplicates, respectively.

| Compounds | EC<sub>50</sub> (mM) | K<sub>i</sub> (mM) | K<sub>d</sub> (mM) |  | Potential MOA | Binding site | Metabolic pathway |
| --- | --- | --- | --- | --- | --- | --- | --- |
|  |  |  | MST | DSF |  |  |  |
| Dihydroxyacetone phosphate (DHAP) | 8.24 ± 1.94 | 4.47 | 17.48 ± 0.98 | 9.67 ± 0.97 | Mixed | Active site | Glycolysis |
| Phosphoenolpyruvate (PEP) | 11.84 ± 1.33 | 5.61 | 11.14 ± 1.44 | 13.36 ± 1.15 | Mixed | Active site | Glycolysis |
| Citraconate (CTA) | 8.05 ± 2.34 | 2.17 | 20.13 ± 2.03 | 14.73 ± 2.13 | Competitive | Active site | Fatty acid metabolism |
| Maleate (MAE) | 9.48 ± 2.85 | 7.15 | 14.44 ± 2.07 | 17.34 ± 2.38 | Mixed | Active site | Gentisate pathway |
| Histamine | 7.35 ± 1.26 | 9.25 | 13.10 ± 0.64 * | N/A | Mixed | N/A | Histidine metabolism |
| ADP | 8.66 ± 1.37 | 3.97 | 19.27 ± 0.96 * | 15.32 ± 1.44 ** | Competitive | N/A | Energy metabolism |
| Hydroxylamine <sup>P</sup> | 14.63 ± 1.99 | 21.13 | 17.24 ± 3.39 * | N/A | Non-specific | N/A | Nitrogen metabolism |

\* Ligand-induced fluorescence change (up to 20%) / aberrant MST traces
\*\* T<sub>m</sub> decreased with increasing ligand concentration
<sup>P</sup> Slight inhibition (15% at 10 mM) also observed towards TPI

**Table 2:**
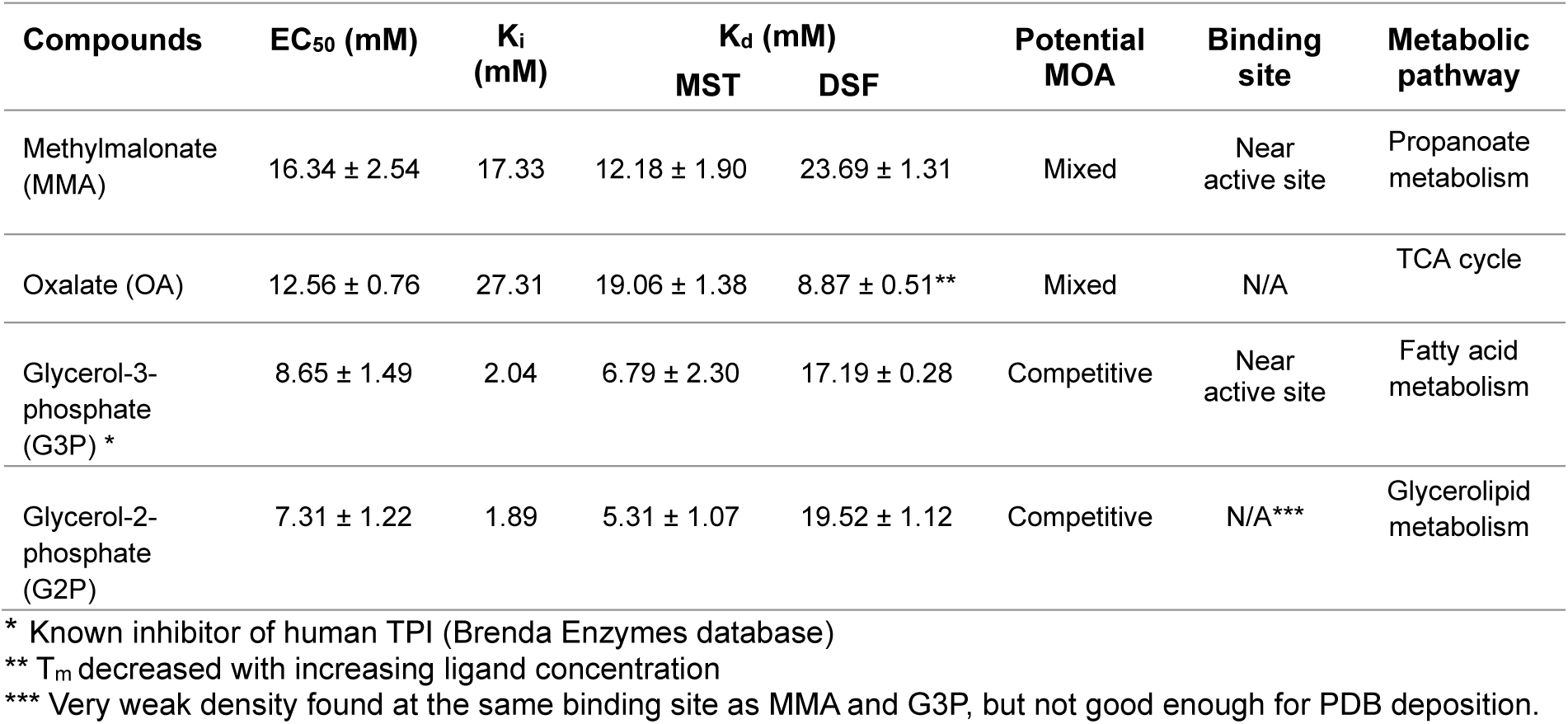
Summary of identified TPI inhibitors. The EC_50_ and K_d_ values are expressed as mean ± standard deviation of two duplicates and triplicates, respectively.

| Compounds | EC<sub>50</sub> (mM) | K<sub>i</sub> (mM) | K<sub>d</sub> (mM) |  | Potential MOA | Binding site | Metabolic pathway |
| --- | --- | --- | --- | --- | --- | --- | --- |
|  |  |  | MST | DSF |  |  |  |
| Methylmalonate (MMA) | 16.34 ± 2.54 | 17.33 | 12.18 ± 1.90 | 23.69 ± 1.31 | Mixed | Near active site | Propanoate metabolism |
| Oxalate (OA) | 12.56 ± 0.76 | 27.31 | 19.06 ± 1.38 | 8.87 ± 0.51** | Mixed | N/A | TCA cycle |
| Glycerol-3-phosphate (G3P) * | 8.65 ± 1.49 | 2.04 | 6.79 ± 2.30 | 17.19 ± 0.28 | Competitive | Near active site | Fatty acid metabolism |
| Glycerol-2-phosphate (G2P) | 7.31 ± 1.22 | 1.89 | 5.31 ± 1.07 | 19.52 ± 1.12 | Competitive | N/A*** | Glycerolipid metabolism |

\* Known inhibitor of human TPI (Brenda Enzymes database)
\*\* T<sub>m</sub> decreased with increasing ligand concentration
\*\*\* Very weak density found at the same binding site as MMA and G3P, but not good enough for PDB deposition.

All identified hits resulted in weak inhibition of the enzyme activity, with EC_50_, K_i_ values and binding affinity (K_d_) in the low millimolar range. Because of low affinities, it was difficult to obtain full saturation curves for these ligands, yet the dose response was clear in both activity and binding assays (Figure 3 & 4). Along with the MST assay, DSF thermal shift experiments were performed to measure the hit binding affinities. The MST binding – and melt curves obtained for the GPI and TPI hits are shown in the supplementary data (Fig. 6S & 7S). Furthermore, none of these hits exhibited apparent inhibitory dose response effect on the coupling enzymes when full titration experiments were conducted (Fig. 5S-1 & 5S-2), suggesting the observed activity decrease was due to inhibition of the target enzymes. Since full saturation of binding/inhibition was not reached at the highest concentrations tested, the EC_50_ values were determined with some degree of uncertainty.

**Figure 3:**
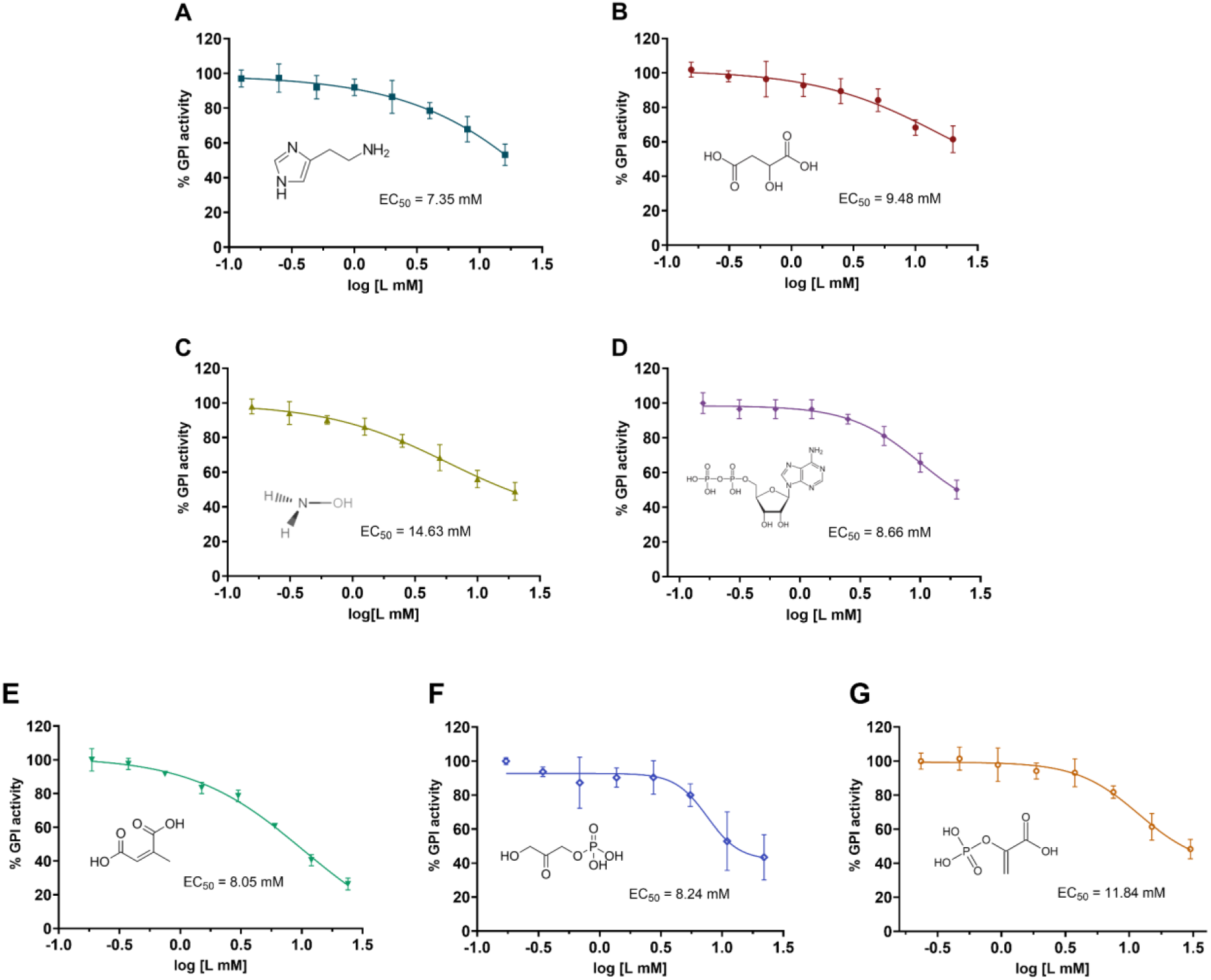
Metabolite inhibitors of human GPI and their estimated EC_50_ values. (A) Histamine. (B) Maleate. (C) Hydroxylamine. (D) ADP. (E) Citraconate. (F) DHAP. (G) PEP. The dose-response curves were obtained by measuring the activity of GPI using enzyme-coupled assay. Each data point represents mean ± SD of two duplicates or triplicates measured on two separate days. Data were fitted using the log(inhibitor) vs. response - Variable slope (four parameters) in Prism GraphPad.

**Figure 4:**
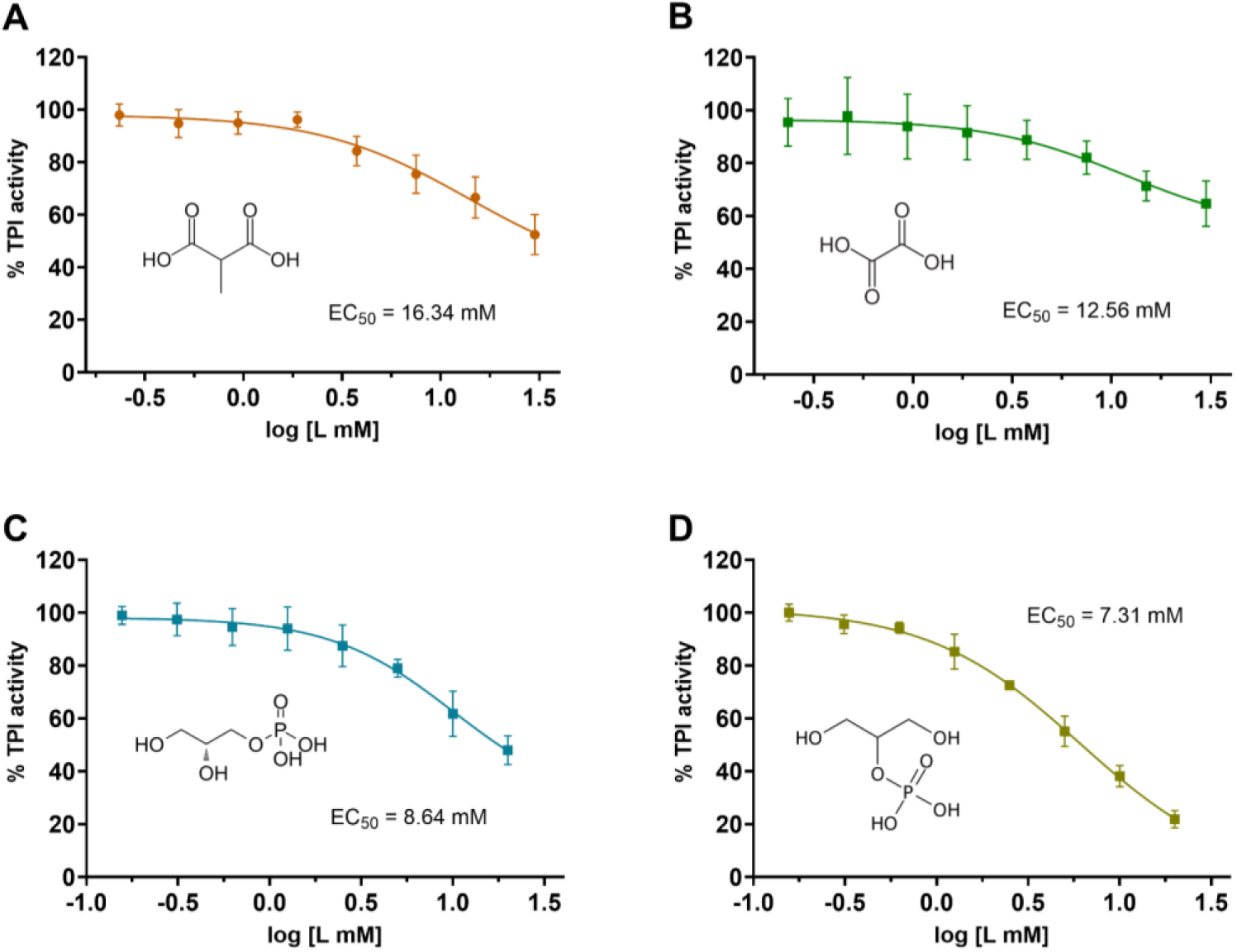
Metabolite inhibitors of human TPI and their estimated EC_50_ values. (A) Methylmalonate. (B) Oxalate. (C) Glycerol-3-phosphate. (D) Glycerol-2-phosphate. The dose-response curves were obtained by monitoring the activity of TPI using enzyme-coupled assay. Each data point represents mean ± SD of two duplicates or triplicates measured on two different days. Data were fitted using the log(inhibitor) vs. response - Variable slope (four parameters) in Prism GraphPad.

### Identified inhibitors show competitive or mixed mode of inhibition

To determine the mechanisms of action (MOA) for the metabolite inhibitors, Michaelis-Menten (MM) steady-state kinetics were measured for GPI and TPI (Figure 5 & 6). Four of the GPI hits, histamine, MAE, DHAP, and PEP, exhibited increased K_m_ and decreased V_max_ (mixed inhibition), while ADP and CTA showed increased K_m_ but almost unaltered V_max_ (nearly competitive), and hydroxylamine had a slight increase in K_m_ and decrease in V_max_ (unclear MOA; data not shown). Of the TPI hits, MMA, G3P, and G2P showed enhanced K_m_ but unchanged V_max_ (pure competitive), and oxalate displayed an increase in K_m_ and a slight decrease in V_max_ (mixed inhibition).

**Figure 5:**
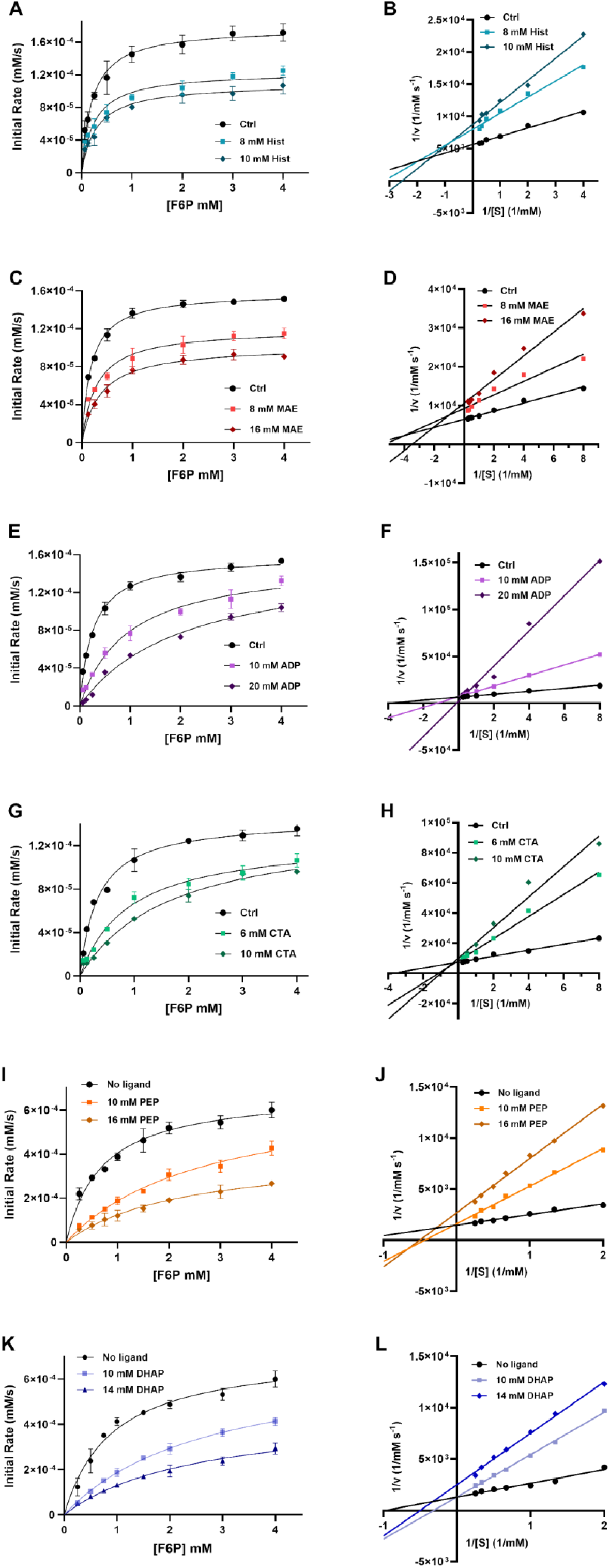
Michaelis-Menten (MM) kinetics and Lineweaver Burk plots of human GPI inhibitors: (A-B) Histamine. (C-D) Maleate. (E-F) ADP. (G-H) Citraconate. (I-J) DHAP. (K-L) PEP. The enzyme activity was measured at 8 different substrate concentrations (0.05-4 mM F6P). Each data point represents 2-3 replicates and error bars denote ± SD of mean.

**Figure 6:**
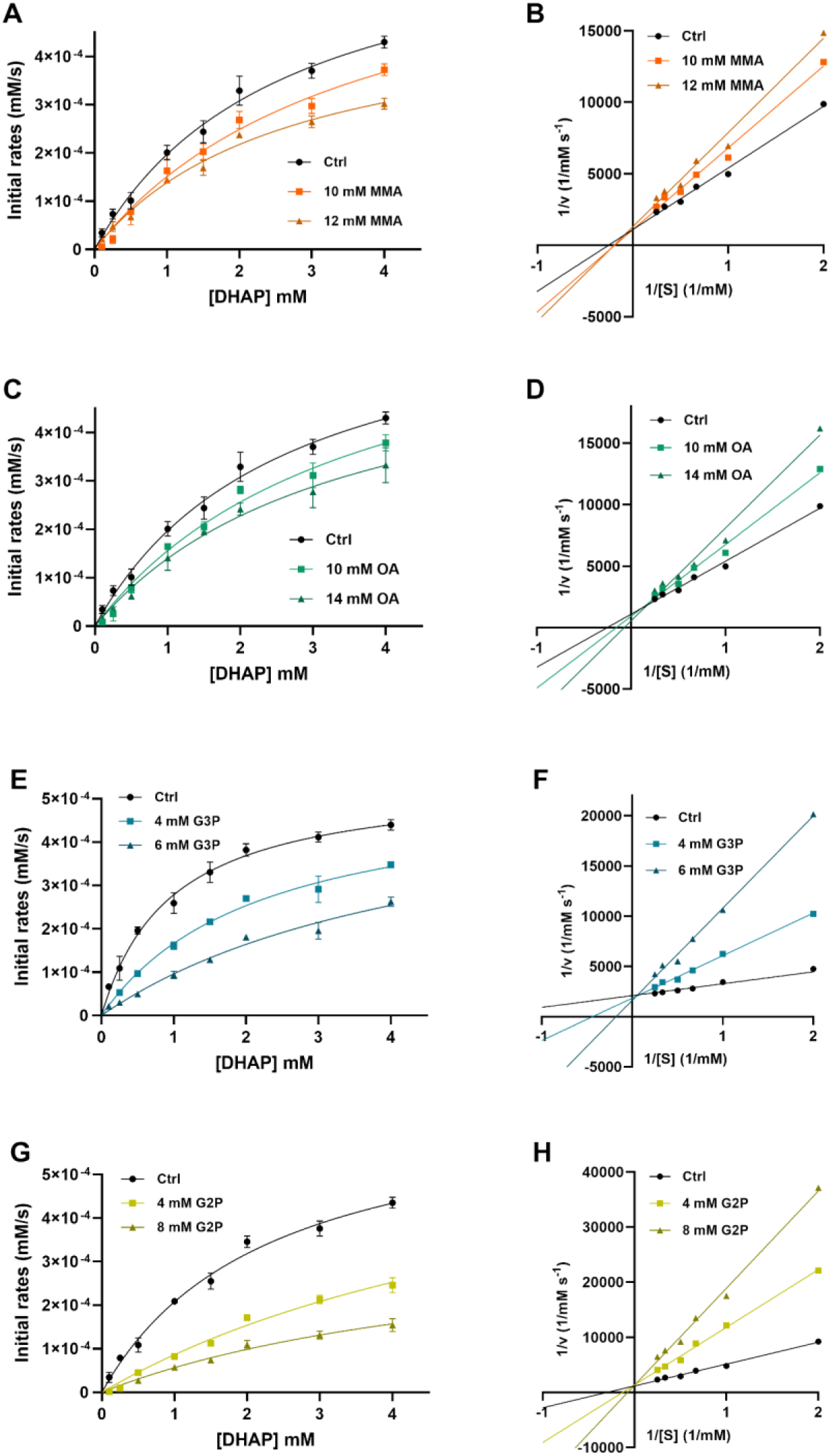
Michaelis-Menten (MM) kinetics and Lineweaver Burk plots of human TPI inhibitors: (A) Methylmalonic acid (MMA). (B) Oxalic acid (OA). (C) glycerol-3-phosphate (G3P). (D) beta-glycerophosphate (G2P). The enzyme activity was measured at 8 different substrate concentrations (0.05-4 mM DHAP). Each data point represents a duplicate and error bars denote ± SD of mean.

### X-ray crystallography revealed orthosteric binding site for identified GPI hits and a new ligand binding site for TPI

To elucidate the binding sites of the GPI and TPI inhibitors, X-ray crystallography was employed. The X-ray diffraction data for human GPI and TPI crystals were collected up to a resolution of 1.17-1.80 Å (Table 3). The crystal structures of GPI had four subunits (two homodimers) in the asymmetric unit (ASU), whereas the TPI structures contained one subunit in the ASU (Fig. 7 & 8). Ligand densities were revealed in the GPI crystal structures co-crystallized with DHAP, PEP, MAE, and CTA; all binding to the active site as shown in Figure 7. DHAP binds in a similar geometry to the previously published structures of GPI complexed with phosphate-containing inhibitors (Fig. 8S), whereas PEP adopts a slightly different position but with the phosphate group similarly oriented in the phosphate-binding region. Both MAE and CTA are bound to the phosphate-binding region in the GPI active site. These four GPI hits interact with various active site residues by means of H-bonding (Fig. 9). The common interacting residues include Ser^210^, Thr^212^, and Thr^215^, all of which have been reported in other GPI-ligand complexes (supplementary data C).

**Table 3:** X-ray diffraction data collection and refinement statistics.

| Structure | GPI-MAE<br>PDB:9FCW | GPI-PEP<br>PDB:9FKF | GPI-DHAP<br>PDB:9FHF | GPI-CTA<br>PDB:9FKC | TPI-MMA<br>PDB:9F69 | TPI-G3P<br>PDB:9FFC |
| --- | --- | --- | --- | --- | --- | --- |
| Wavelength | 0.976 | 0.976 | 0.976 | 0.976 | 0.976 | 0.976 |
| Space group | P 21 21 21 | P 21 21 21 | P 21 21 21 | P 21 21 21 | P 61 2 2 | P 61 2 2 |
| Resolution range | 37.78 - 1.4<br>(1.45 - 1.4) | 46.19 - 1.6<br>(1.657 - 1.6) | 45.05 - 1.8<br>(1.864 - 1.8) | 48.5-1.60<br>(1.63-1.60) | 31.83 - 1.17<br>(1.21-1.17) | 24.43 - 1.25<br>(1.295-1.25) |
| Total reflections | 5879303<br>(393048) | 4184252<br>(417435) | 2968155<br>(298270) | 2820899<br>(138671) | 2864457<br>(163955) | 2510506<br>(249752) |
| Unique reflections | 454749<br>(41170) | 310109<br>(30723) | 220113<br>(21752) | 310649<br>(15137) | 82312<br>(8037) | 68952<br>(6688) |
| Multiplicity | 12.9 (9.5) | 13.5 (13.6) | 13.5 (13.7) | 9.1 (9.2) | 34.8 (20.4) | 36.4 (37.3) |
| Completeness | 98.60<br>(89.35) | 99.76<br>(99.79) | 99.87<br>(99.91) | 99.8<br>(99.1) | 99.81<br>(99.06) | 99.59<br>(99.46) |
| mean(I) / sig(I) | 11.96 (1.47) | 9.89 (1.49) | 9.74 (1.56) | 8.7 (0.9) | 23.55 (1.23) | 28.46 (4.10) |
| Wilson B-factor | 15.21 | 19.17 | 21.88 | 21.65 | 13.39 | 14.55 |
| $R_{merge}$ | 0.1278<br>(2.239) | 0.1929<br>(3.025) | 0.2689<br>(3.017) | 0.18<br>(3.92) | 0.1163<br>(2.320) | 0.06885<br>(0.9524) |
| $CC_{1/2}$ | 0.999<br>(0.478) | 0.999<br>(0.671) | 0.998<br>(0.695) | 1.00<br>(0.44) | 0.999<br>(0.437) | 1.000<br>(0.997) |
| $R_{work}$ | 0.1427 | 0.187 | 0.185 | 0.197 | 0.1465 | 0.1449 |
| $R_{free}$ | 0.1896 | 0.224 | 0.227 | 0.227 | 0.1837 | 0.1811 |
| Ramachandran outliers (%) | 0 | 0 | 0 | 0 | 0 | 0 |
| Sidechain outliers (%) | 0.8 | 1.3 | 1.8 | 1.5 | 0.5 | 0.5 |
| RSRZ outliers (%) | 2.5 | 4.1 | 3.4 | 5.0 | 2.4 | 2.4 |
| Clash score | 2 | 3 | 3 | 3 | 1 | 1 |

**Figure 7:**
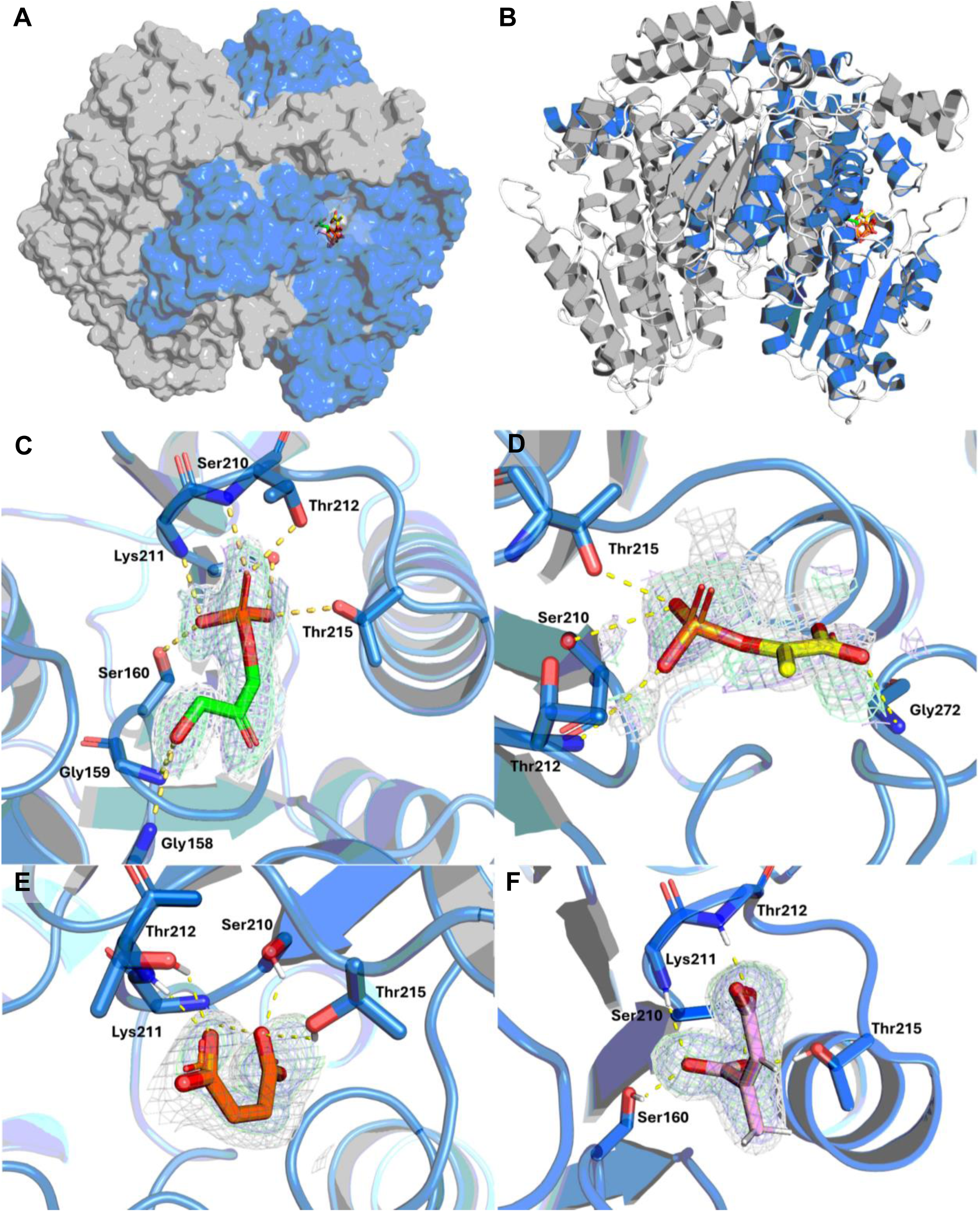
Crystal structures of human GPI dimer in complex with metabolite inhibitors. (A & B) Overall structure of human homodimeric GPI crystallized in 20% PEG3350, 0.16 M CaCl_2_, 0.058M HEPES, pH 7.0. Four bound ligands: DHAP (green), PEP (yellow), MAE (orange), and CTA (pink), shown in one of the subunits. (C) GPI co-crystallized with DHAP (ligand density in chain A); refined to 1.80 Å. (D) GPI co-crystallized with PEP (ligand density in chain C); refined to 1.60 Å. (E) GPI co-crystallized with MAE (ligand density in chain A); refined to 1.40 Å. (F) GPI co-crystallized with CTA (ligand density in chain D); refined to 1.60 Å. 2mFo-DFc electron density maps for the ligands are shown as blue mesh at 1.0 σ, difference omit maps as green mesh at 3.0 σ, and polder maps as gray mesh at 3.0-3.5 σ. The yellow dashed lines represent the polar contacts between the ligands and the active site residues.

**Figure 8:**
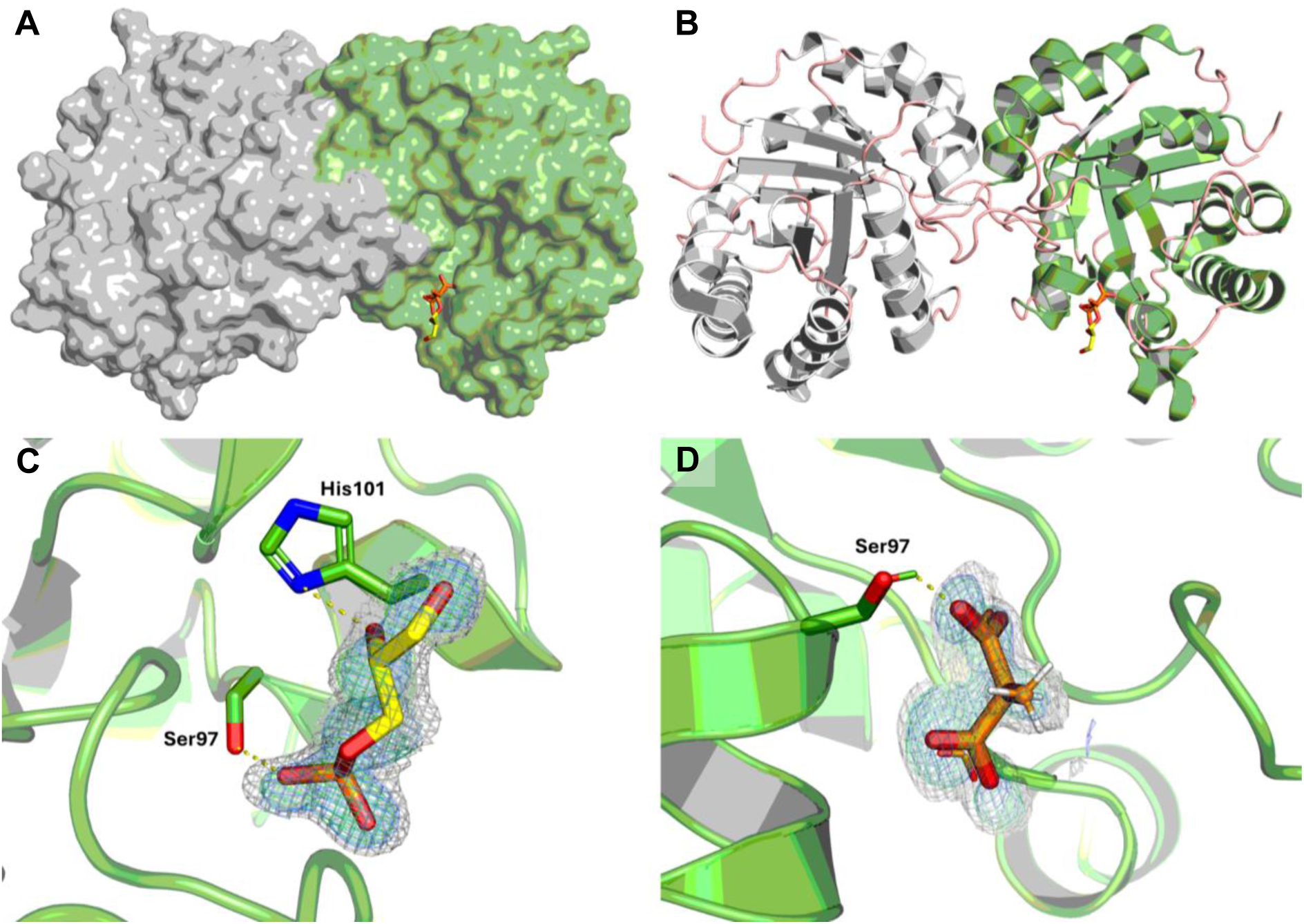
Crystal structures of human TPI dimer in complex with metabolite inhibitors. (A & B) Overall structure of homodimeric GPI crystallized in 0.14 M KBr and 24% PEG 2000 MM. The bound ligands: G3P and MMA are shown in yellow and orange, respectively. (C) TPI crystal soaked with G3P; refined to 1.25 Å. (D) TPI crystal soaked with MMA; refined to 1.17 Å. 2mFo-DFc electron density maps for the ligands are shown as blue mesh at 1.0 σ, difference omit maps as green mesh at 3.0 σ, and polder maps as gray mesh at 3.0-3.5 σ. The yellow dashed lines represent the polar contacts between the ligands and the active site residues.

**Figure 9:**
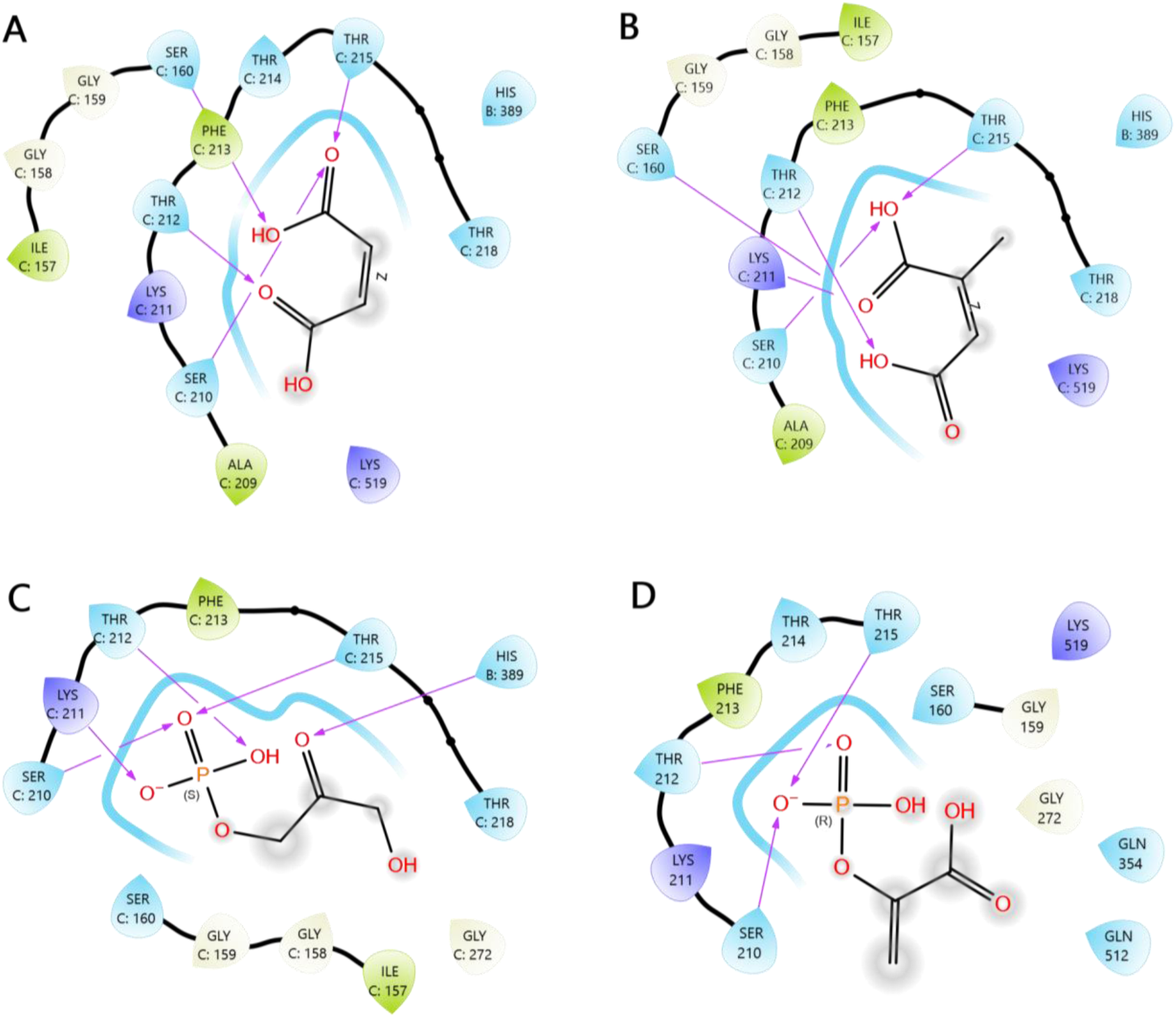
Ligand-interaction diagrams showing the possible interactions between the GPI active site residues and identified GPI hits – (A) maleate (MAE). (B) citraconate (CTA). (C) DHAP. (D) PEP. The interaction maps were generated using Ligand Interaction script in Maestro (Schrödinger Inc), with cutoff set to 4.0 Å. Pink arrows represent the H-bond interactions.

Despite relatively low occupancy, electron densities matching two TPI hits, G3P and MMA, were found in a cleft ∼10 Å from the substrate-binding site (Figure 8), occupying a position different from those observed for the substrate (DHAP) and other TPI inhibitors. The binding position of MMA overlaps with that of the phosphate moiety in G3P (Fig. 8S-2). The interactions of these two ligands with TPI occur through H-bonding with Ser^97^ and His^101^ residues, both of which have not been reported to be associated with ligand binding. However, residue Ser^97^ is largely conserved and has been recognized for controlling water near the active site and positioning of the catalytic base Glu^165^ [36,37]. In the unliganded state, there is an H-bond between Ser^97^ and Glu^165^; this H-bonding interaction is disrupted when a substrate binds [38]. In contrast to the identified hits, many known TPI inhibitors, such as 2-phosphoglycolate (PGA, substrate analog) [39], PEP [7], and phosphoglycolo-hydroxamate (PGH) [40], bind to the substrate-binding site and interact with highly conserved residues in the active site, including Asn^11^, Lys^13^, His^95^, Glu^165^, Gly^171^, and Ser^211^ (supplementary data C), all of which are essential for the catalytic function of TPI [39,41–43].

For most of the GPI X-ray diffraction data, the structures were close to identical to the previously published apo structures (ligand-free active sites) of the enzymes (PDB: 1IAT and 2JK2 for GPI and TPI respectively), with Cα root-mean-square deviations (RMSD) in the range of 0.2-0.3 Å (Table 2S). Whereas for TPI, the RSMDs were significantly higher, or up to around 0.6 Å. In the subunit alignment for GPI, there was local dimer asymmetry present at the “large” domain at α20-turn-α21 motif and is likely a crystal-contact artefact (Fig. 9S A-H). In the GPI structures complexed with DHAP and CTA, the ligands densities were found in all four subunits of the asymmetric unit, but in the PEP- and MAE-bound structures, only two of the subunits were occupied. Furthermore, PEP showed an asymmetric binding, where a local increase in RSMD (dimer asymmetry) was observed in the active site loop (situated between β3 and α11) when the PEP-bound and unbound subunits were aligned.

The alignment of the two TPI structures to the apo form showed that they were highly similar except at surface crystal contact regions (Fig. 9S I-L). Interestingly, when the structures were aligned with a structure containing the substrate analogue (PDB: 6UPF), PGA, a loop transition seen in the PGA-bound TPI, was not observed for the identified TPI inhibitors (Fig. 9S J & L).

### Inhibition of glycolytic flux by phosphoenolpyruvate (PEP)

To assess if, at high concentration, any of the hits can potentially perturb glycolysis, extracellular flux measurement was performed for one of the identified hits, PEP. The flux measurement was undertaken with two cell lines – Hulec-5a (Human Lung Microvascular Endothelium cell line) and U2OS (human osteosarcoma cell line), using the seahorse FeX96 analyzer (Agilent USA). The cellular glycolytic flux (Fig. 10 & 11) was approximated using the proton efflux rate (PER) calculated from the extracellular acidification rate (ECAR) measured in real time. Treatment with PEP (10-30 mM) led to a decrease in the PER of both cell lines in a concentration-dependent manner, with more significant impact on U2OS cell line (Fig. 11). A comparable drop in PER was observed for the GPI competitive inhibitor – 6PGc. In the case of U2OS, the attenuation of PER lasted throughout the measurement (Fig. S10-B). But in that of Hulec-5, the inhibitory effect persisted for a limited timeframe (∼40 min), and the flux partially recovered after longer incubation time (Fig. S10-A). Nevertheless, the observed PER reduction was primarily due to the inhibition of glycolysis. This was indicated by the drastic decline of PER after successive injection of mitochondrial inhibitors (Rot/AA) and glycolytic blocker (2-DG).

**Figure 10:**
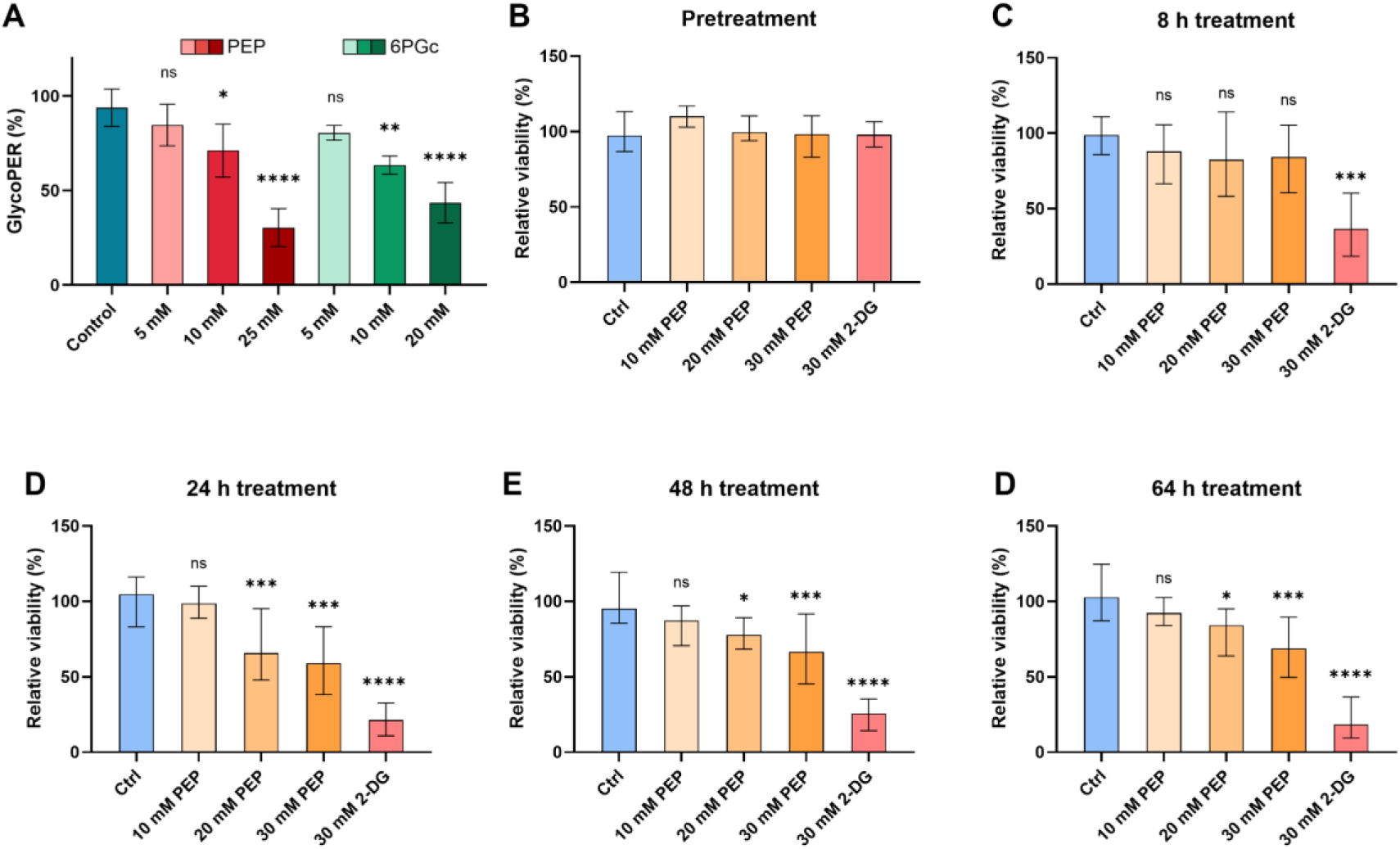
Percentage glycolytic proton efflux (PER; A) and cell viability (B-F) measured for Hulec-5a cells in the absence and presence of phosphoenolpyruvate (PEP) and positive controls: 6-phosphogluconate (6PGc) and 2-deoxyglucose (2-DG). The glycolytic PER was measured after effector compound and Rot/AA injection. Viability assessed before compound addition (B) and after 8 hours (C), 24 hours (D), 48 hours (E), and 68 hours (F) of treatment. Data were normalized with respect to the baseline (basal glycoPER). Each bar represents the mean value ± SD for six replicates (n = 6) per condition. Statistical p-values: ns (p ≥ 0.05), * (p = 0.01-0.05), ** (p = 0.001-0.01), *** (p = 0.0001-0.001), and **** (p < 0.0001), obtained from one-way ANOVA in GraphPad Prism.

**Figure 11:**
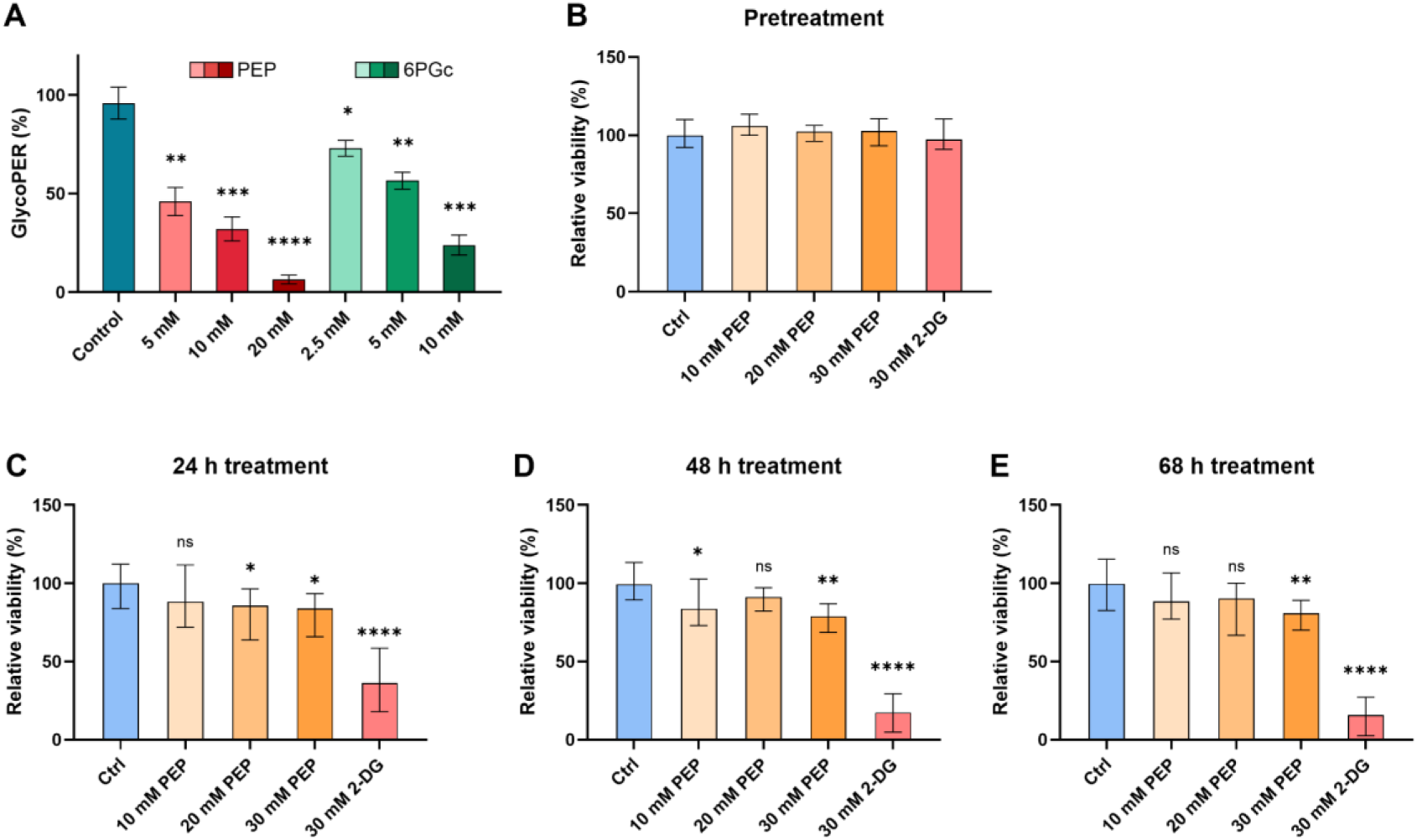
U2OS cell glycolytic proton efflux (PER; A) and viability (B-E) measured the absence and presence of phosphoenolpyruvate (PEP) and positive controls. The glycolytic PER was measured after effector compound and Rot/AA injection. Viability assessed before compound addition (B) and after 24 hours (C), 48 hours (D), 68 hours (E) of treatment. 6-phosphogluconate (6PGc) and 2-deoxyglucose (2-DG) were used as positive controls in the flux measurement and viability assay, respectively. Data were normalized with respect to the baseline (basal glycoPER). Each bar represents the mean value ± SD for six replicates (n = 6) per condition. Statistical p-values: ns (p ≥ 0.05), * (p = 0.01-0.05), ** (p = 0.001-0.01), *** (p = 0.0001-0.001), and **** (p < 0.0001), obtained from one-way ANOVA in GraphPad Prism.

In addition, the viability of Hulec-5a and U2OS cell lines was assessed before and after continuous PEP treatment (5-30 mM) over a period of three days. Compared to the negative control (no inhibitor), there was an apparent decrease in the Hulec-5a cell viability over the course of treatment (Fig. 10), especially at high PEP concentration (30 mM). However, no substantial changes to the viability were observed for the U2OS cell line (Fig.11); even at high concentration (30 mM), PEP did not effectively inhibit the cell growth. The positive control – 2-deoxyglucose (2-DG), by contrast, led to a drastic decrease in the viability of both cell lines as a result of glycolysis inhibition. 2-DG was able to inhibit both the glycolytic flux and cell viability in an effective manner, whereas PEP reduced the PER without having substantial effect on the viability. Being a relatively potent inhibitor, 2-DG is capable of blocking the dual metabolic flux of glycolysis and oxidative PPP, and it is also a stable glucose analogue that cannot be fully metabolized after being phosphorylated by hexokinase and is unlikely to be used as an energy source. Conversely, PEP can be metabolized over time, thus having no pronounced impact on the cell growth.

## Discussion

According to the Brenda Enzymes database and various literatures [7,12,19–21], all previously reported modulators of human GPI and TPI are either competitive or mixed inhibitors that act by occupying the active sites (some of them are listed in supplementary data C). In fact, one of the inhibitor identified here, G3P, has already been reported as a competitive inhibitor of human TPI [44], but up till now, there was no available structural information on its binding site. Three of the identified GPI hits in this study have been reported to inhibit homologous GPI of non-human origin. PEP has been shown to be a competitive inhibitor of *E. coli* GPI [45], whereas DHAP has been proposed to weakly inhibit GPI from *S. oleracea* [46], and maleate was reported a weak inhibitor of GPI in *T. aestivum* [47].

The identified GPI hits – DHAP, PEP, citraconate, and maleate all bind to the active site (Fig. 8S), in a manner similar to known competitive inhibitors (by forming H-bonds with conserved active site residues). In GPI, the binding of phosphate moiety is essential for anchoring the substrate to the active site and the initiation of the isomerization reaction [48]. Both citraconate and maleate act as mimics of the phosphate anchor, occupying the region where the phosphate group of GPI substrates is normally bound. The TPI hits, MMA and G3P, were observed in a cleft ∼10 Å away from the substrate binding site. Another TPI inhibitor, G2P, had a similar interaction site as the two above, but the electron density was too weak for accurate modeling. Unlike other competitive inhibitors, these TPI hits did not cause a conformational change of the TPI catalytic loop upon binding. Presumably, either the binding of these inhibitors to an alternative site has hindered the movement of the loop within the catalytic cycle of the enzyme, or the structures were locked in their open-loop conformation (apo form) upon crystallization.

Despite showing inhibitory effect on GPI activity, the binding of histamine, ADP, and hydroxylamine to GPI, was not fully supported by binding assay results nor confirmed by crystallographic data. However, the kinetic studies showed that ADP acts by a competitive-inhibition mode, suggesting that it binds to the GPI active site. Histamine, on the other hand, appeared to be a mixed-type inhibitor, but it may also be an orthosteric binder. In a recent study, Ahmad et al. [20] performed an N-substitution of GPI transition-state analogs with various amine groups, one of which was histamine. All the N-substituted derivatives showed relatively potent inhibition towards GPI (IC_50_ in low µM), and two of them displayed a new binding mode where the amine substituents occupied a subsite in the active site. Although no crystallographic data was reported for the histamine-containing inhibitor, we suspect that it adopts a similar binding mode as that of its counterparts, with the histamine motif bound to a subsite within the active site. Unlike the other GPI hits, hydroxylamine did not exhibit a clear mode of action, and no useful binding data was obtained from the MST and DSF assays. Weak inhibition of human GPI by hydroxylamine has been previously reported [49], but the mechanism is still unclear. The inhibitory effect might be a result of protein destabilization, as hydroxylamine could act as a nucleophile, inducing cleavage at Asn-Gly (NG) sites in proteins [50,51]. There are three and two NG sites on GPI and TPI, respectively. Two of the GPI NG sites and both of the TPI NG sites are located in a solvent-accessible region of the enzyme dimer interface, making them prone to nucleophilic attack and subsequent hydrolysis of the NG sites, leading to structural destabilization and reduced catalytic activity.

### Can weak modulators potentially affect pathway activity?

Currently, the search for selective, strong inhibitors targeting glycolysis enzymes still remains a central dogma in cancer drug discovery. Most of the known small-molecule inhibitors of human glycolysis enzymes are relatively potent, with IC_50_ in the nM-µM range [12,17,18,52]. Strong inhibitors may have limited clinical use due to the high toxicity they pose to healthy cells, including immune cells (with anti-tumor functions) and other proliferating non-tumor cells [15,53]. The GPI and TPI hits identified here are all weak binders, with K_d_ in the mM range. As most weak binders lack target specificity, they tend to show high cross reactivity towards several proteins, resulting in off-target effects. However, such cross reactivity may not be a disadvantage if multi-target modulation allows for maximum therapeutic efficacy. It has been proposed that weak binders acting on multiple targets in a synergistic manner can be as effective as high-affinity modulators that selectively interact with a single target [54]. Moreover, a binding efficiency analysis of natural products demonstrated that high binding affinity (K_d_ < low µM) is not a necessary prerequisite for therapeutic efficacy [55]; despite having poor binding efficiency, many natural products were able to obtain therapeutic efficacy in animal studies. Furthermore, a study of metabolic network indicated that partial attenuation of a small number of targets could be even more efficient than complete inhibition of a single target [56], further highlighting the therapeutic potential of weak modulators.

To examine the impact of weak modulation on glycolytic pathway activity, we performed extracellular flux measurement using PEP. The PEP-treated cells exhibited a dose-response decrease in the PER. This reduced PER could be attributed to the combined effects of direct inhibition on glycolysis enzymes and indirect inhibition of hexokinase (HK) and lactate dehydrogenase (LDH). Besides showing weak inhibition towards GPI, PEP has been reported to inhibit human TPI [7] and bind more strongly to the latter enzyme (K_d_ = 1-2 mM). High amount of PEP could also induce secondary inhibitory effects such as inhibition of LDH activity through pyruvate buildup [57], resulting in diminished lactate production. Furthermore, the inhibition of GPI could lead to accumulation of G6P, which may in turn inhibit HK activity, exacerbating the inhibitory effect. In addition, PEP might inhibit another human glycolysis enzyme – phosphofructokinase I (PFK-I), as it has been reported to be a non-competitive inhibitor of PFK-I in various species, including mouse [45,58], whose PFK-1 shares a sequence homology of 89% with the human counterpart.

It seems reasonable that PEP acts as a feedback inhibitor of glycolytic enzymes, given that it is an intermediate from glycolysis downstream. Similarly, the accumulation of DHAP and ADP might also inhibit GPI activity, and increased levels of PEP and G3P might subject TPI to feedback regulation, as the enzyme bridges between glycolysis and triglyceride synthesis. However, it is unlikely for every weak modulator to be able to effectively alter the activity of a particular pathway simply with enhanced concentration, considering the possibility of metabolic rerouting and the presence of other competing interactions. Inhibition of glycolytic enzymes can result in accumulation of pathway intermediates, which may be redirected back to glycolysis via other pathways. For instance, a recent metabolic analysis showed that inhibition of GPI activity will result in the buildup of G6P, which has two fates: it may either inhibit HK, lowering the glycolytic flux, or it may be channeled to the PPP and converted into glycolytic intermediates, depending on metabolic demands [59]. While the oxidative phase of PPP can provide a positive feedback loop that further promotes GPI inhibition via 6PGc production, the oxidative and non-oxidative branches of PPP together, may compensate for GPI inhibition by providing F6P and GAP to recover glycolytic flux through the upper and lower parts of glycolysis [59]. Accumulation of metabolites can also bring about detrimental consequences as some compounds can become undesirably toxic at high levels, or worse yet, bear oncogenic potential that promotes tumor development. Even relatively non-reactive, non-toxic metabolites, such as DHAP and MMA, can spontaneously break down into toxic agents when their levels exceed the normal physiological concentrations [60–62].

### Assay interference in low-affinity compound screening

Weak binding or biological interactions are generally little studied, partially because of the difficulties in screening and evaluating them. To probe for subtle interactions, high ligand concentration is required, and this can result in prominent assay interference and leads to high propensity for false positive results. This was clearly evident by the high number of frequent hitters (167 compounds) appeared in the primary screening against both GPI and TPI. Solubility also becomes an issue when screening compounds at high concentrations. A portion of the MetaSci metabolites (309 out of 1250) had to be excluded due to low water solubility. The average initial hit rates of the colorimetric activity – and the MST binding assay were 9.6% and 16.1%, respectively. However, the fraction of non-specific hits (frequent hitter) was much greater in the binding assay. Despite the high initial hit rate of 27.5% in the luminescent assay, it did not reveal any hits specific to the target enzymes, and the screening results obtained for the two targets were similar. This high primary hit rate and large number of common hitters was partly due to compound interference. Colored compounds, particularly intensely colored ones (with absorbance range in 320-400 nm), could result in reduced absorbance and/or total luminescence, and in some cases to a large extent that may be mistaken as strong inhibition. And certain colorless compounds, such as chlorogenic acid, gallic acid, gentisic acid, and 3,5-Diiodo-L-tyrosine, developed color after being incubated with assay reagents, causing readout interference. The color changes might be due to oxidation or side reactions between the LDR substrates and test compounds. A 2008 study by Auld et al. [63] found that 0.9% of the 70,000 compounds tested in the screening against luciferase were assay-interfering false positives, and they exhibited at least 50% inhibition on the enzyme at ligand concentrations below 10 µM. Since our intention was to find weak modulators, screening the compounds at high µM to low mM concentrations would inevitably lead to high assay interference. It can thus be concluded that the LDR assay is not suitable for screening weak modulators.

Aside from compound interference, enzyme-coupled assays also suffered from unwanted modulatory effects on the auxiliary enzymes, particularly in the case of the luminescence assay. The activity screening conducted without GPI or TPI revealed many non-target-specific hits, some of which may interact with the coupling enzymes in a productive way. As the enzyme substrates and products cannot be directly detected by optical means, to monitor their formation without the aid of coupling enzymes, one could utilize chromogenic or fluorogenic probes. An assay that employs such chemical probes is the Seliwanoff’s test (endpoint assay), which can detect the formation of F6P by absorbance measurement (at 410-460 nm) of a colored product formed from a reaction with resorcinol and heat treatment [64]. This method was initially used in the GPI screening to measure the rate of GPI-catalyzed conversion of G6P to F6P in an endpoint manner. But it was disused due to its laborious nature, low consistency, and reactivity of resorcinol with various aldehydes and ketones, resulting in high background signal. An attempt was also made to monitor the activity of TPI using a non-enzyme-coupled, colorimetric kinetic assay that was derived from a cell-based assay using a chemical probe (2-ABAO) showing sensitivity and selectivity towards a group of aldehydes [65]. The TPI-catalyzed reaction can be followed in the direction from GAP to DHAP, which reacts with 2-ABAO to form a final product that has a maximal absorption at around 405 nm. However, this method was only feasible at low pH (4.5-6.0), and it was susceptible to interference by certain aldehyde- and ketone-containing compounds, thus not being further employed.

The label-free MST was chosen as the main binding assay as it does not require labeling nor target immobilization. This method has few drawbacks, such as being prone to interference by compounds who exhibit fluorescence and quenching properties. Apart from that, a high ligand concentration (> 10 mM) was needed for weak binders to produce apparent changes in the MST signal to indicate target binding. This was noticed in the primary screening (ligand concentration ≤ 10 mM), where many of the identified hits were missed by the MST method. All the identified inhibitors showed up as hits in the primary activity screening, but only three of them (PEP, G2P, and G3P) came up as hits in the screening with MST. When reassessing those activity hits with MST assay by 12-point serial dilution (from 40-50 mM downward), all compounds exhibited dose-response pattern and most with K_d_ values above 10 mM. By contrast, when primary MST hits (non-activity assay hits) were re-evaluated using the activity assay, none of them showed no apparent effect (≥ 25% change) on the enzyme activity, suggesting they were either false-positives, or possibly, silent binders. Comparing binding analysis screening methods, surveys have shown that hits tested on the same libraries can vary substantially across different screening methods [66]. Nevertheless, when it comes to false-negative rate, which has been observed up to 50% for known binders, MST should be less of concern than other affinity-based methods such as DSF and surface plasmon resonance (SPR) [67].

Finally, in addition to the label-free MST, DSF thermal shift assay was employed to validate the potential GPI and TPI hits. In most cases, a small T_m_ shift (1-2°C) was observed for the enzyme targets, and it was difficult to obtain log-sigmoidal dose-response curves. Even in the presence of relatively strong GPI and TPI inhibitors (6PGc and PEP), the target enzymes did not display regular dose-response behavior in the DSF assay. Based on the results, the DSF assay may not be an ideal method for screening GPI and TPI binders, however, it could still serve as a hit validation tool to improve hit confidence.

### Conclusion

In summary, we conducted a screening on common human metabolites at high concentrations to identify weak modulators of human GPI and TPI. To maximize the chances of finding potential hits, activity and binding assays were employed in parallel. The screening revealed that GPI and TPI can be weakly modulated by small-molecule inhibitors that bind in proximity to the enzyme active sites. All the identified hits, except for G3P, are considered novel with respect to the human enzyme targets. Although these compounds are relatively weak inhibitors, together with other known inhibitors, they could serve as starting points for the structural optimization and design of novel inhibitors targeting human GPI and TPI. The common binding mode of the identified GPI inhibitors indicated that the phosphate group, which serves as molecular anchor of most GPI inhibitors, was replaceable by anionic moieties (MAE and CTA) other than sulfates. Moreover, the binding of the identified TPI inhibitors to a different position from other orthosteric TPI inhibitors suggested a possibility of designing novel inhibitors with new binding modes. Such inhibitors may possess enhanced selectivity over typical orthosteric binders and may also be less prone to drug resistance – a problem that could be encountered by modulators acting on the less conserved allosteric sites.

## Supporting information

Appendix A - Ligand solubility guide

PDB validation report for PDB 9F69

PDB validation report for PDB 9FCW

PDB validation report for PDB 9FFC

PDB validation report for PDB 9FHF

PDB validation report for PDB 9FKC

PDB validation report for PDB 9FKF

## Abbreviations

GPI: glucose-6-phosphate isomerase;
TPI: triosephosphate isomerase;
G6PDH: glucose-6-phosphate dehydrogenase;
GAPDH: glyceraldehyde-3-phosphate dehydrogenase;
HK: hexokinase;
PFK: phosphofructokinase;
PK: pyruvate kinase;
LDH: lactate dehydrogenase;
G6P: glucose-6-phosphate;
F6P: Fructose-6-phosphate;
DHAP: dihydroxyacetone phosphate;
GAP: glyceraldehyde-3-phosphate;
6PGc: 6-Phosphogluconate;
PEP: phosphoenolpyruvate;
CTA: citraconate;
MEA: maleate;
ADP: adenosine diphosphate;
MMA: methylmalonate;
G2P: glycerol-2-phosphate;
G3P: glycerol-3-phosphate;
PGA: 2-Phosphoglycolate;
PGH: phosphoglycolo-hydroxamate;
2-ABAO: aminobenzamide oxime;
EC_50_: half maximal effective concentration;
IC_50_: Half maximal inhibitory concentration;
K_i_: inhibition constant;
MOA: mode/mechanism of action;
MST: microscale thermophoresis;
F_norm_: normalized fluorescence change (F_hot_/F_cold_);
DSF: differential scanning fluorometry;
ASU: asymmetric unit;
RMSD: root-mean-square deviations;
ECAR: extracellular acidification rate;
PER: proton efflux rate;
GlycoPER: glycolytic proton efflux rate;
2-DG: 2-Deoxy-glucose;
Rot/AA: rotenone/antimycin;
SPR: surface plasmon resonance.

## ASSOCIATED CONTENT

### Supporting information

Manuscript supplementary data A

List of metabolites supplementary data B

Known inhibitors interactions supplementary data C

Appendix A: Solubility guide for MetaSci Complete metabolite library.

### Author Contributions

JGH and YYJ planned experiments; YYJ performed experiments; JGH and YYJ analyzed data; JGH and ÓR contributed reagents and access to facilities; JGH and YYJ wrote the paper; All authors took part in revision of the manuscript draft.

### Funding Sources

Financial support from the Icelandic Research Fund (project 239695-051), the Icelandic cancer society, and the Blái Naglinn initiative is gratefully acknowledged.

## Acknowledgments

We acknowledge MAX IV Laboratory for time on Beamline BioMAX under Proposal 20230497. Research conducted at MAX IV, a Swedish national user facility, is supported by the Swedish Research council under contract 2018-07152, the Swedish Governmental Agency for Innovation Systems under contract 2018-04969, and Formas under contract 2019-02496. Heidi Erlandssen and Ronny Helland at the Department of Chemistry, The Arctic University of Tromsø are thanked for their guidance in crystallization screens. Arnar Ingi Vilhjálmsson at the Department of medicine at the University of Iceland is gratefully thanked for assistance and guidance in glycolytic flux and cell viability studies.

