## Supplementary data A for "Human Glycolysis Isomerases are Inhibited by Weak Metabolite Modulators"

#### Experimental procedures

##### Expression and purification of recombinant GPI, TPI, and G6PDH from *E.coli*

Transformed cells were cultivated in 1 L LB broth in 5 L culture flask supplemented with L-rhamnose (200 µM), ampicillin (100 µg/mL), and chloramphenicol (30 µg/mL). Prior to protein expression, the cell culture was incubated at 37°C for 2-3 hours with 180 RPM shaking (5 cm throw diameter) until reaching an OD600 of 0.4-0.8. The expression of the target enzymes was induced by the addition of IPTG (400 µM) to the cell culture. Following induction, the cells were incubated for 5-7 hours at 37°C (for GPI and TPI expression) or overnight at 25°C (for G6PDH expression), and then harvested by centrifugation at 4000 x g at 4°C for 10 minutes. The harvested cells were resuspended in a lysis buffer (0.01% Triton X-100, 0.1 mg/mL lysozyme, 1 mM 2-mercaptoethanol, 25 mM Tris pH 7.9) and disrupted (under an ice bath) by ultrasonication for 5 minutes, with 5-second pulses and 15-second pauses using the UP200St Ultrasonic Lab Homogenizer (Hielscher Ultrasonics, Germany). The cell suspension was centrifuged at 40000 x g at 4°C for 30 minutes to collect the supernatant, which was filtered through a 0.22 µm pore filter before purification.

##### Enzyme activity assays

*Enzyme activity hit screening and validation*: The screening against the targets was first performed at a single point (in 20 µM – 10 mM range) to identify primary hits – compounds that showed changes in initial rate ≥ 25%, w.r.t the control (no ligand). The threshold for the luminescent assay was set to ≥ 30%, as more enzymes and substrates were involved, the false-positive hit rate was much greater. As a control measure, primary hits were then screened against the auxiliary enzymes to remove false positives. Primary hits that showed ≤ 15% alteration to the dehydrogenase activity were re-screened against the targets at two concentrations (≤ 10 mM) to check for concentration-dependent effect. Following that, an 8-point titration was conducted for the secondary hits against the targets for $EC_{50}$ estimation, and also against coupling enzymes in separate assays to ensure the observed modulation was target-specific. Finally, hits showing dose-response modulatory effect towards the target activity were assessed for their mechanism of action (MOA) via Michaelis-Menten (MM) steady-state kinetics and Lineweaver-Burk plots. Compounds showing spurious dose-response patterns, such as bell-shaped steady-state curves, were not further assessed for MOA. Each sample was measured in duplicates in the initial screen and at least two duplicates for $EC_{50}$ and MOA assessments. The $EC_{50}$ and inhibition constant ($K_i$) values for all hits were determined in GraphPad Prism using log(inhibitor) vs. response - Variable slope (four parameters) and competitive or mixed-mode inhibition fit, respectively.

*Activity assay conditions:* For GPI screening and dose-response tests, the final assay concentrations of enzyme and reagents were as follow: 1.26 ng/mL of dimeric GPI, 51.2 µg/mL G6PDH, 2 mM F6P, 10 mM NADP+, and 0-20 mM test compounds (metabolites), and 50% v/v of luciferase detection reagent (LDR) for luminescent measurement. The Michaelis-Menten steady-state kinetics of GPI were obtained with same assay components described above, except that the concentrations of F6P was ranged from 0.05 to 4 mM. The activity of G6PDH (control test) was measured with an assay mixture containing 25.6 µg/mL G6PDH, 0.05-4 mM G6P, 8 mM NADP+, and 0-24 mM compounds (and 50% v/v LDR). Besides LDR, all reagents and test compounds used in GPI and G6PDH screening were prepared in buffer containing 100 mM Tris, pH 7.4 and 150 mM NaCl. In TPI screening, the assay mixture contained 80 ng/mL TPI, 0.1 mg/mL GAPDH, 2 mM DHAP, 8 mM NAD+, 6 mM Arsenate, 5 mM EDTA, and 0-20 mM compounds, and 50% v/v of LDR for luminescent assay. The Michaelis-Menten kinetics for TPI were obtained using same assay components, except that the DHAP concentrations was varied from 0.1 to 4 mM. The reaction mixture for GAPDH activity test (control assay) comprised of 30 µg/mL GAPDH, 0.2-6 mM DL-GAP, 8 mM NAD+, 6 mM Arsenate, 5 mM EDTA, and 0-24 mM compounds (and 50% v/v LDR). Except for the LDR and test compounds (metabolites), other reagents used for TPI and GAPDH screens were prepared in 100 mM triethanolamine (TEA), pH 7.4. The test compounds were assayed in a concentration range of 10 µM–24 mM, depending on their solubility and DMSO concentration. The negative controls, wherein the screened enzyme was 100% active (no inhibition), contained all relevant assay components except for test compounds. For metabolites dissolved in DSMO, negative controls containing the same concentration (%v/v) of DMSO were used instead. To confirm that the target enzymes were rate-limiting, their activities were assessed and showed no apparent changes with increasing concentrations (5-100x fold) of the coupling enzymes.

*Spectrophotometry:* The colorimetric assay was executed in the following manner: dispense of the pre-equilibrated enzyme mixture in wells containing test compounds, incubation of the mixture at 26°C for 5-10 minutes, addition of substrate mixture to the assay plate, and absorbance measurement at 340 nm; monitored over a period of 3-4 min at 26°C (26°C was used for the assays due to temperature fluctuation at 25°C in the microplate reader chamber). The luminescent assay was conducted according to the manufacturer's protocol, and the luminescence measured over a duration of 10-12 minutes at 26°C, with integration time set to 1000 ms.

#### Figures 1S – Protein quality control

For quality control, the sample purity was monitored during and after the purification and ultrafiltration with SDS-PAGE using the SurePAGE™ 4-12% gradient gels from GenScript (USA). For structural and stability assessment, Circular Dichroism (CD) analysis of protein secondary structure and melting temperature assays were performed on a JASCO 1100 CD spectropolarimeter. In addition, the dimeric molecular weights (Mw) of the purified proteins were confirmed by the size exclusion chromatography (SEC) method, using a Superdex® 200 10/300 GL column from Cytiva. The SEC analysis was performed using the MST buffer (100 mM Tris, 150 mM NaCl, pH 7.4), at a flow rate of 0.5 mL/min. For Mw calibration, protein standards of various sizes were used: myoglobin (17.7 kDa), ovalbumin (OA, 45 kDa), Vibrio alkaline phosphatase (VAP, 120 kDa purified inhouse) [1], Lactate dehydrogenase (LDH, 140 kDa), and lactate oxidase (LO, 160 kDa) from Sigma (USA).

1) Purification of recombinant human GPI and TPI from *E. coli.*

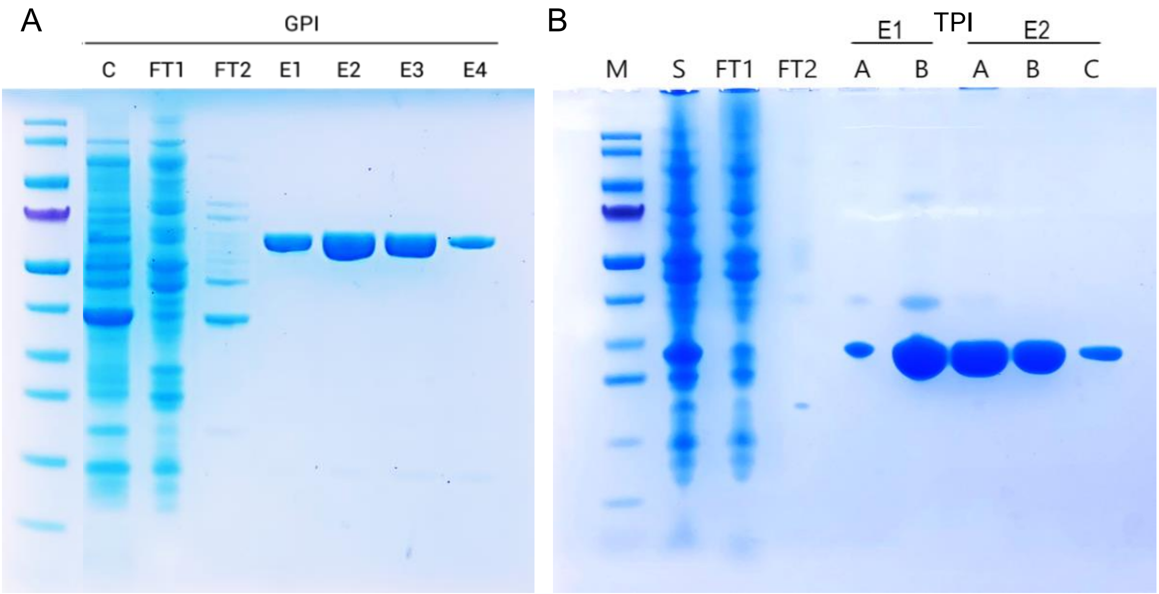


Figure 1S-1: SDS-PAGE analysis of GPI purification (A) and purified TPI (B) by His-Trap IMAC column. (A) C: clarified lysate; FT1-2: flow-through; E1-3: elution peaks from His-Trap column; E4: buffer-exchanged and diluted elution fraction. (B) M: protein marker; S: cell lysate; FT1-2: flow-through; E1-E2: elution peak fractions from His-Trap column (A) and after buffer-exchange (B) and dilution (C). Both GPI and TPI purified to > 95% purity, with yield close to 20 mg/L culture.

2) Purification of recombinant yeast G6PDH from *E. coli*.

G6PDH (57.5 kDa) – 25°C overnight culture; yield = 7.4 mg/L

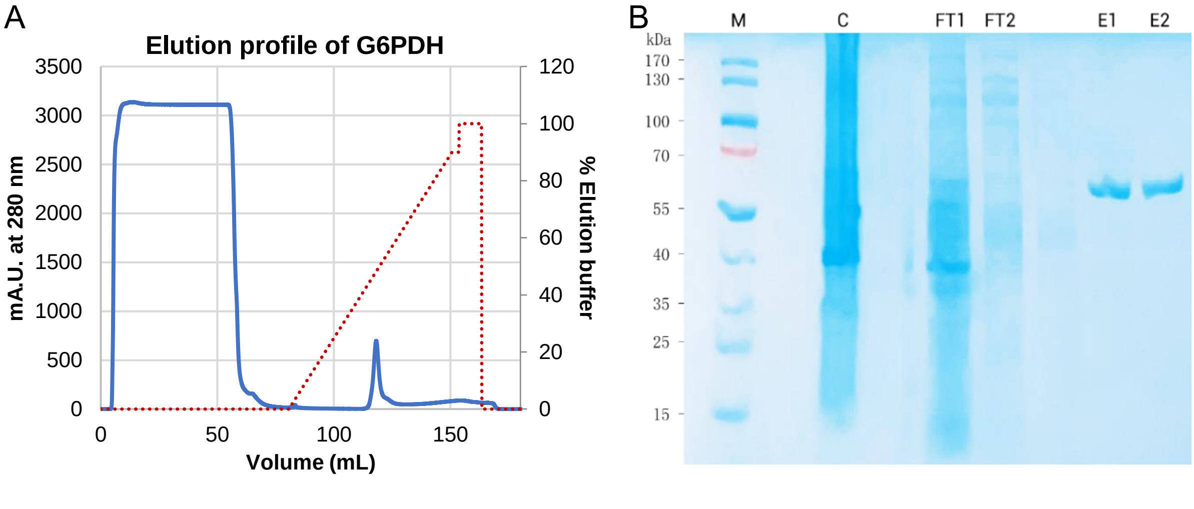


Figure 1S-1: His-Trap affinity chromatography purification of G6PDH from Lemo21(DE3) *E. coli* cells. (A) Elution profile of G6PDH monitored by absorbance at 280 nm. (B) Gradient (4-12%) SDS-PAGE of crude extract (C), flow-through (FT1 and FT2), and elution fractions (E1 and E2) from His-trap column.

#### 3) Circular dichroism

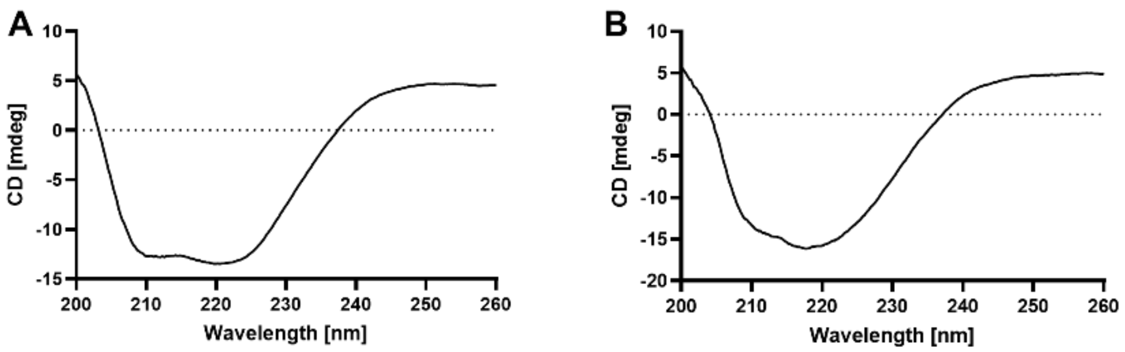


Figure 1S-3: Far-UV circular dichroism CD spectra of purified recombinant proteins (0.15 mg/mL) – GPI (A) and TPI (B) in 10 mM HEPES (pH 7.4) and 2 mM $MgCl_2$. Measured using JASCO 1100 CD spectropolarimeter, with 1 mm cell. CD spectra collected between 200-250 nm, at 25°C, with 1 nm bandwidth. Spectral features of an alpha-beta secondary structure were observed for both enzymes, with characteristic minima near 208 and 222 nm.

### Figures 2S – Assay validation

#### MST-binding assay – positive controls

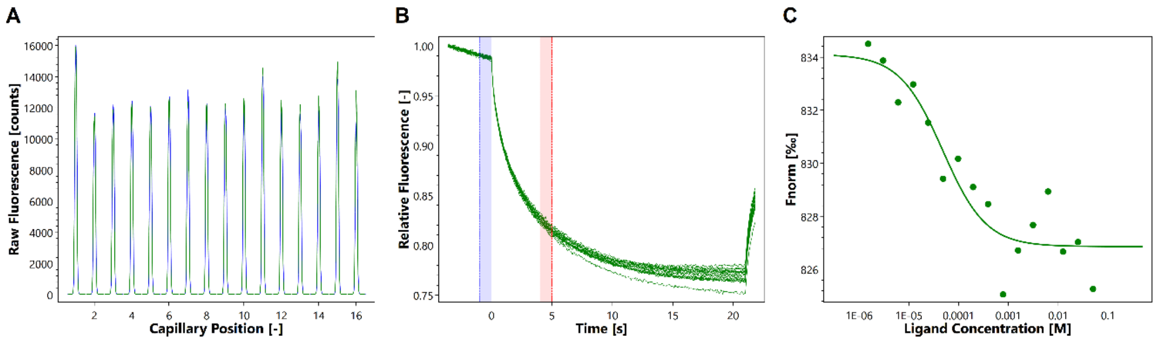


Figure 2S-1: Binding of 6-phosphogluconic acid (6PGc) to glucose-6-phosphate isomerase (GPI) in MST buffer (pH 7.4), monitored using label-free MST instrument, with medium MST power, 40% excitation power, and on-time interval of 30 seconds. (A) Capillary scans of initial fluorescence across different ligand concentrations. (B) MST traces before, during, and after the IR-laser was turned on. (C) Dose response curve of 6PGc to GPI, with an estimated $K_d$ of 578 µM, obtained using an on-time of 5 sec. Unbound (left) and bound states (right). The $K_d$ value was determined using the $K_d$ fit model in the MO.Affinity Analysis software.

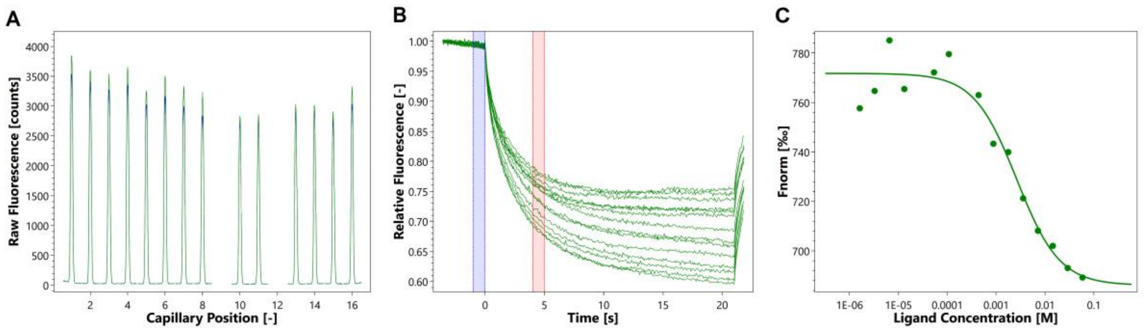


Figure 2S-2: Binding of phosphoenolpyruvate (PEP) to triosephosphate isomerase (TPI) in MST buffer (pH 7.4), monitored using label-free MST instrument, with medium MST power, 40% excitation power, and on-time interval of 30 seconds. (A) Capillary scans of initial fluorescence across different ligand concentrations, with two outliers discarded. (B) MST traces before, during, and after the IR-laser was turned on. (C) Dose response curve of PEP to TPI, with an estimated $K_d$ of 2.28 mM, using the $K_d$ fit model in the MO.Affinity Analysis software. A on-time of 5 sec was used for analysis. Unbound (left) and bound states (right).

#### Validation of enzyme-coupled activity assays using competitive inhibitors:

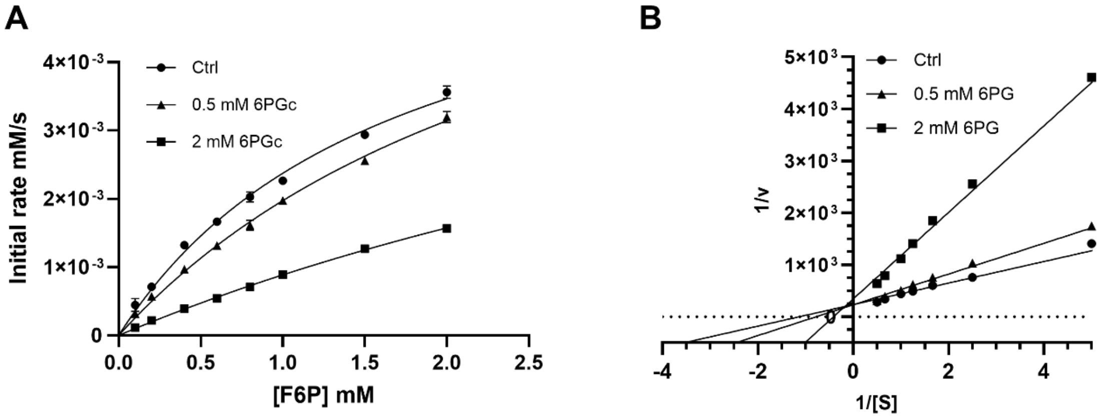


Figure 2S-3: Michaelis-Menten Kinetics (A) and Lineweaver-Burk plots (B) for GPI-catalyzed reaction in the absence and presence of 0.5- and 2-mM 6PGc. The GPI activity was measured by monitoring the absorbance of NADPH at 340 nm. The inhibition mode of 6PG was revealed by the Lineweaver-Burk plots (constructed using GraphPad Prism).

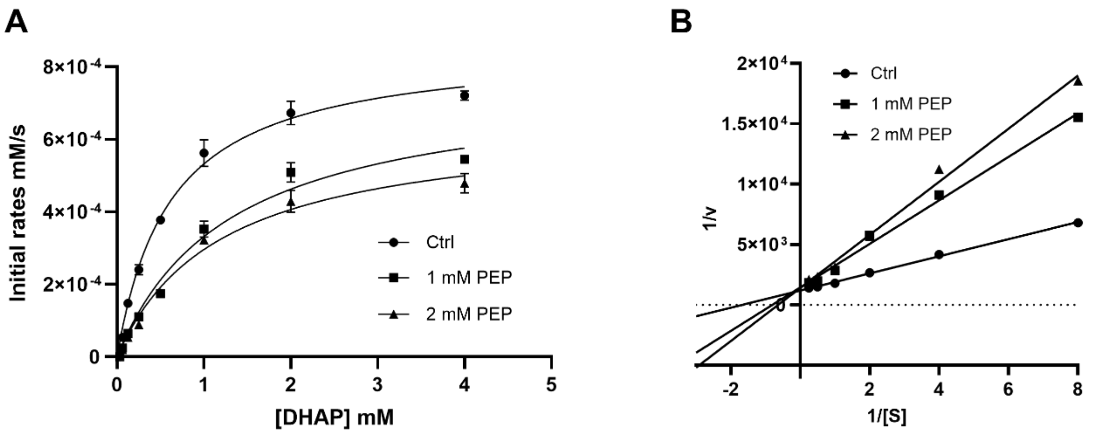


Figure 2S-4: Michaelis-Menten Kinetics (A) and Lineweaver-Burk plots (B) for TPI-catalyzed reaction in the absence and presence of 1- and 2-mM PEP. The TPI activity was determined by monitoring the absorbance of NADPH at 340 nm. PEP inhibited TPI via competitive mechanism as indicated by the Lineweaver-Burk plots constructed using GraphPad Prism.

#### Table S1 – DMSO tolerance test on GPI and TPI

| % DMSO | % GPI activity | |
|---|---|---|
| **0.08** | 100 | 100 |
| **0.16** | 89 | 96 |
| **0.31** | 94 | 99 |
| **0.63** | 97 | 92 |
| **1.25** | 99 | 96 |
| **2.50** | 89 | 95 |
| **5.00** | 98 | 93 |
| **10.00** | 95 | 84 |

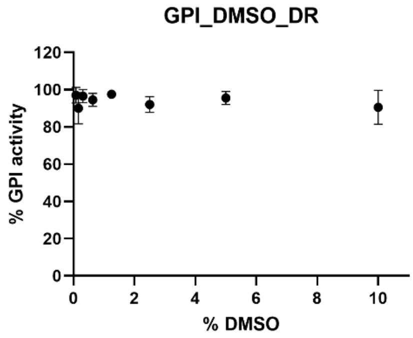


| % DMSO | % TPI activity | |
|---|---|---|
| **0.08** | 100 | 100 |
| **0.16** | 94 | 91 |
| **0.31** | 95 | 93 |
| **0.63** | 94 | 90 |
| **1.25** | 97 | 95 |
| **2.50** | 98 | 96 |
| **5.00** | 90 | 88 |
| **10.00** | 80 | 78 |

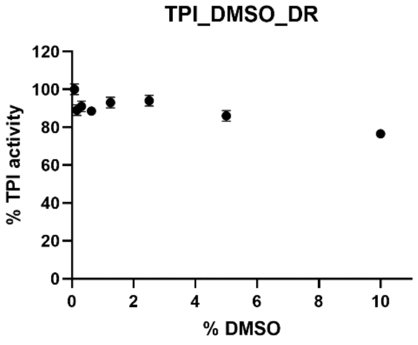


Prior to measurement, the enzymes were incubated with 0.08-10% DSMO for 10 minutes, at 25°C.

#### Figure 3S – GPI screening summary

##### Initial binding and activity screens – GPI

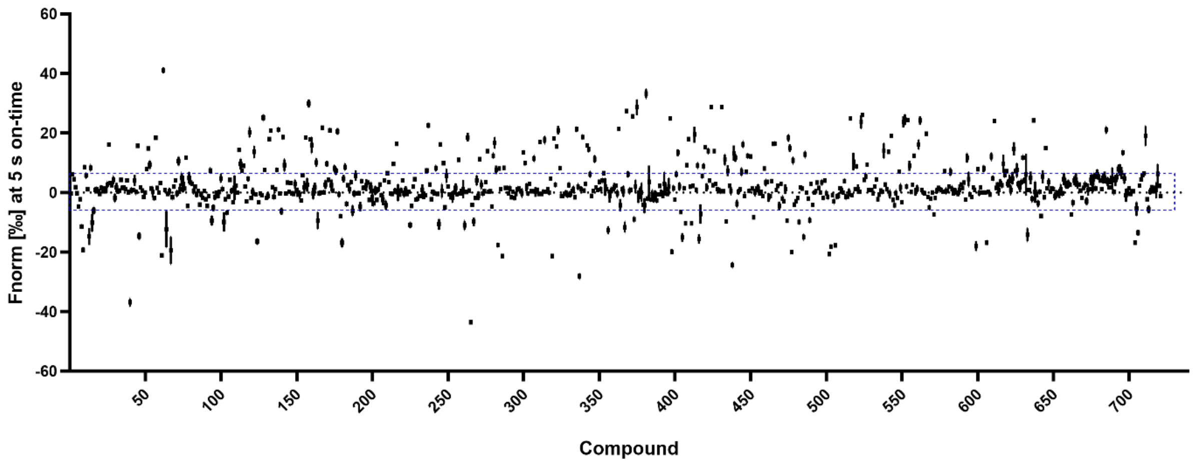


Figure 3S-1: Initial binding screens – GPI. Screening of the MetaSci metabolites (721 in total) against human GPI using label-free Microscale Thermophoresis (MST). Compounds that fall outside the dashed lines (MST signals changed by at least 5 units) are considered initial hits. The compounds were screened at two different concentrations (in 10 µM – 10 mM range, depending on their solubility); only the highest concentrations (20 µM – 10 mM ) are shown here. The screening was conducted at 25°C, with on-time response set to 5 seconds. At least two duplicates were performed for each compound. The dash box represents compounds that fall within the threshold.

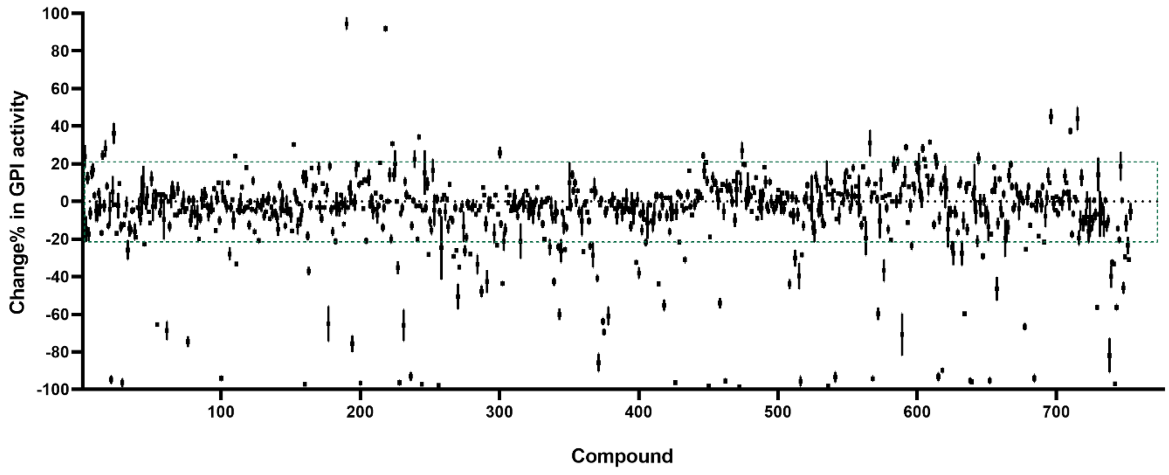


Figure 3S-2: Initial activity screens – GPI. Enzymatic screening of the MetaSci metabolites (753 in total) against human GPI using colorimetric G6PDH-coupled assay. The initial rates were determined by monitoring the formation of NADPH at 340 nm. Compounds that fall outside the dashed lines (showing activity changes > 20%) are considered initial hits. The compounds were screened at one concentration (in 20 µM – 10 mM range). The screening was conducted in a 96-well format, at 26°C, using a microplate reader. At least two replicates were performed for each compound. The dash box represents compounds that fall within the threshold.

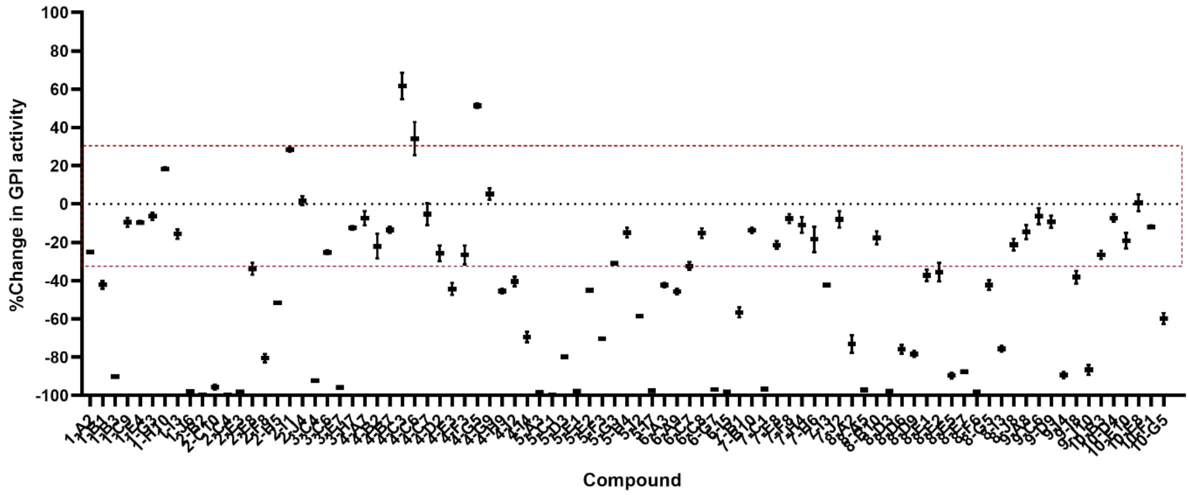


Figure 3S-3: Luminescent initial activity screens – GPI. Enzymatic screening of MetaSci metabolites at a single concentration (in 20 µM – 10 mM range), using the NAD(P)-Glo assay. 149 compounds were screened; 52 compounds were excluded (not shown) due to assay readout interference. The dash box represents compounds that fall within the threshold.

#### Figure 4S – TPI screening summary

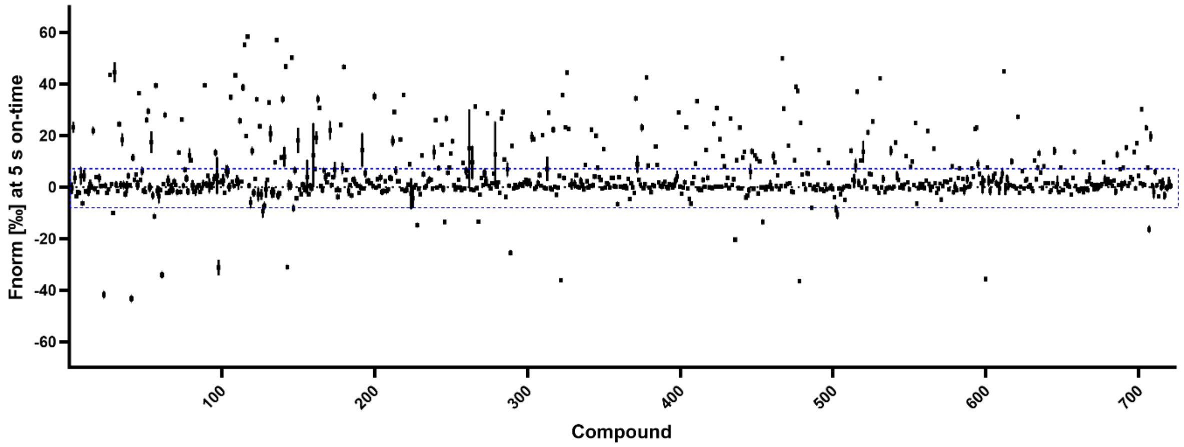


Figure 4S-1: Initial binding screens – TPI. Microscale Thermophoresis (MST) screening of the MetaSci metabolites (721 in total) against human TPI. Compounds that fall outside the dashed lines (MST signals changed by at least 5 units) are considered initial hits. The compounds were screened at two different concentrations (in 10 µM – 10 mM range, depending on their solubility); only the highest concentrations (20 µM – 10 mM) are shown here. The screening was conducted at 25°C, with on-time response set to 5 seconds. At least two duplicates were performed for each compound. The dash box represents compounds that fall within the threshold.

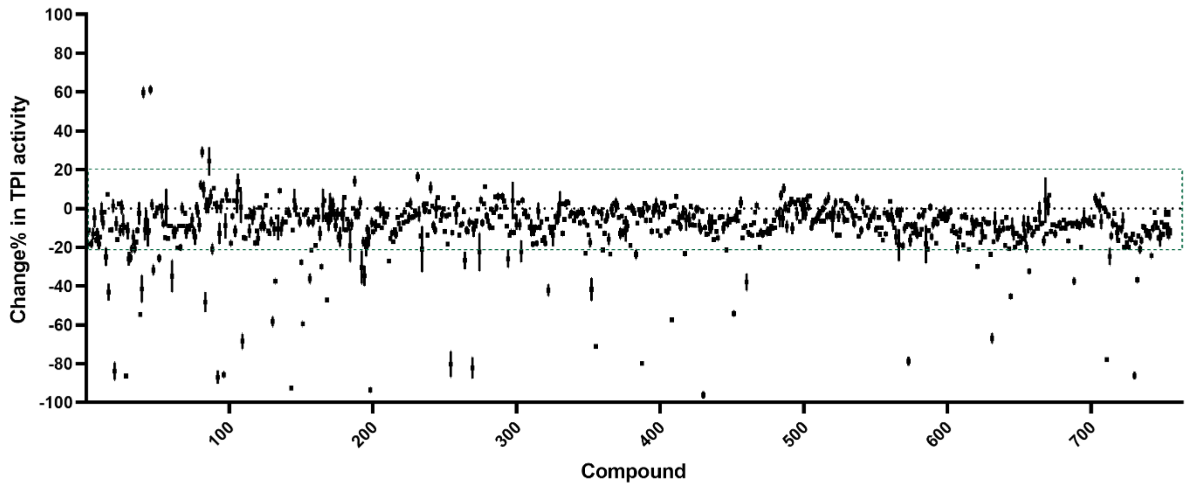


Figure 4S-2: Initial activity screens – TPI. Enzymatic screening of the MetaSci metabolites (754 in total) against human TPI using colorimetric GAPDH-coupled assay. The initial rates were determined by monitoring the formation of NADH at 340 nm. Compounds that fall outside the dashed lines (showing activity changes > 20%) are considered initial hits. The compounds were screened at one concentration (in 10 μM – 10 mM range). The screening was conducted in a 96-well format, at 26°C, using a microplate reader. At least two replicates were performed for each compound.

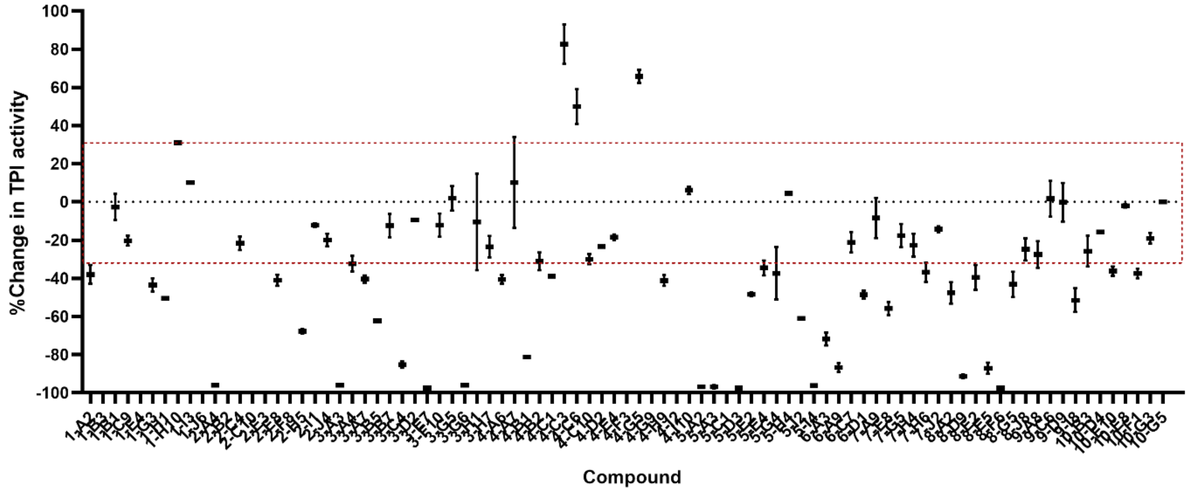


Figure 4S-3: Luminescent initial activity screens – TPI. Enzymatic screening of MetaSci metabolites at a single concentration (in 20 μM – 10 mM range), using the NAD(P)-Glo assay. 149 compounds were screened; 52 compounds were excluded (not shown) due to assay readout interference.

### Figure 5S – Evaluation of dose-response activity against target and coupling enzymes

The coupling enzyme used for GPI assay, G6PDH, was weakly inhibited by glucosamine-6-phosphate ($EC_{50}$ = 5.94 mM and $K_d$ = 5.74 mM) and few other compounds that contain a phosphate or indole-like group. In addition, G6PDH showed reduced activity in the presence of a metal chelator, EGTA, suggesting that the enzyme might be metal ion-dependent. The coupling enzyme used in TPI assay, GAPDH, was found to be inhibited by 3-methyl-2-oxindole ($EC_{50}$ = 1.03 mM) and 2,3-diaminopropionic acid ($EC_{50}$ = 9.77 mM), and several other compounds. G6PDH and GAPDH are widely studied targets for the inhibition of the rate-limiting step in pentose phosphate pathway (PPP) [2] and a potential flux-limiting step in aerobic glycolysis [3], respectively. The G6PDH and GAPDH used in the coupled assays were of yeast and rabbit origins, which share a sequence similarity of 46.84% and 94.89% to their human counterpart, respectively.

#### 8-point titration of potential GPI hit against GPI and G6PDH

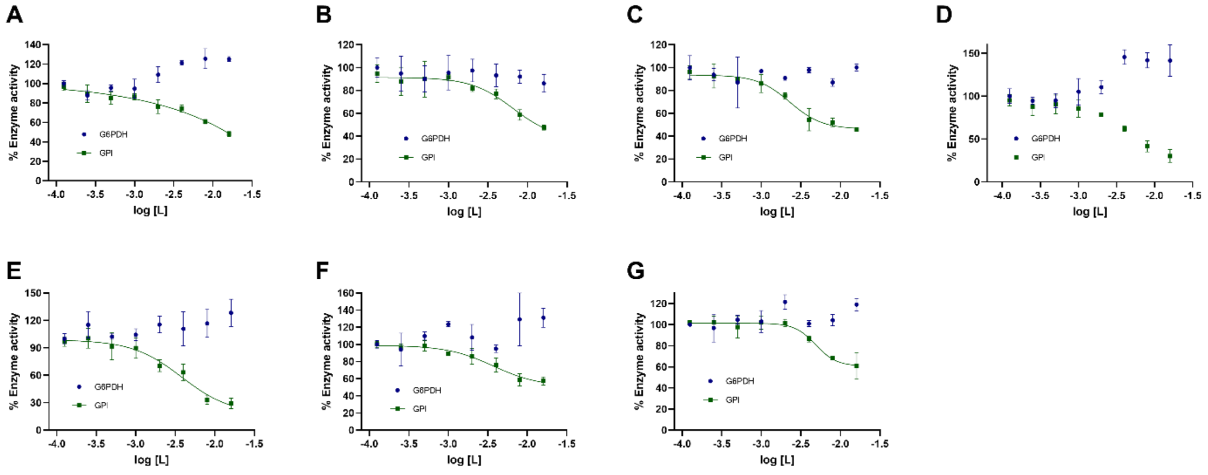


Figure 5S-1: Dose-response curves obtained for GPI (green) and G6PDH (blue) in the presence of histamine (A), maleate (B), hydroxylamine (C), ADP (D), citraconate (E), DHAP (F), and PEP (G). Data were fitted using the Sigmoidal dose-response (variable slope) in Prism GraphPad. Each data point represents mean ± SD of two duplicates.

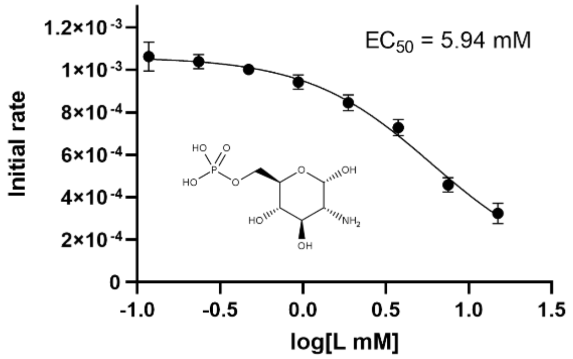


Figure 5S-2: Inhibition of yeast G6PDH by glucosamine-6-phosphate revealed by full-titration of the compound against the coupling enzyme. Each data point represents mean ± SD of two duplicates.

#### 8-point titration of potential TPI hit against TPI and GAPDH

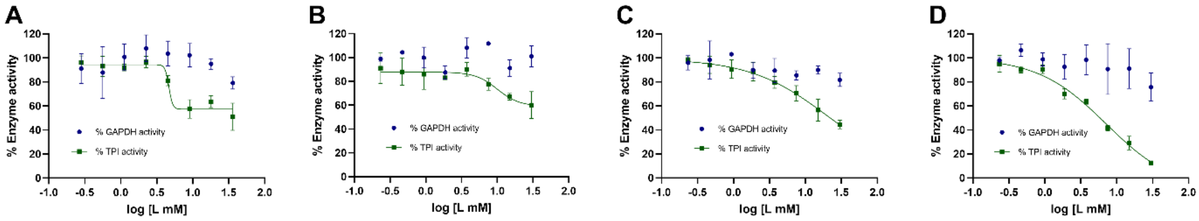


Figure 5S-3: Dose-response curves obtained for methylmalonate (A), oxalate (B), glycerol-3-phosphate (C), and glycerol-2-phosphate (D). Data were fitted using the Sigmoidal dose-response (variable slope) in Prism GraphPad. Each data point represents mean ± SD of two duplicates.

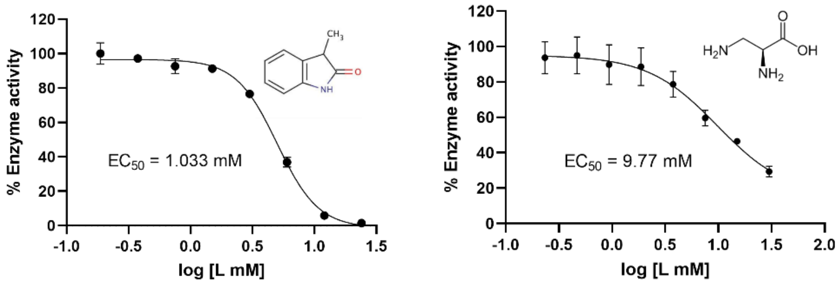


Figure 5S-4: Inhibition of rabbit GAPDH by 3-Methyl-2-oxindole and 2,3-diaminopropionic acid revealed by full titration of compounds against the coupling enzyme. Each data point represents mean ± SD of two duplicates.

### Figure 6S – MST binding curves

#### $K_d$ estimation of GPI hits

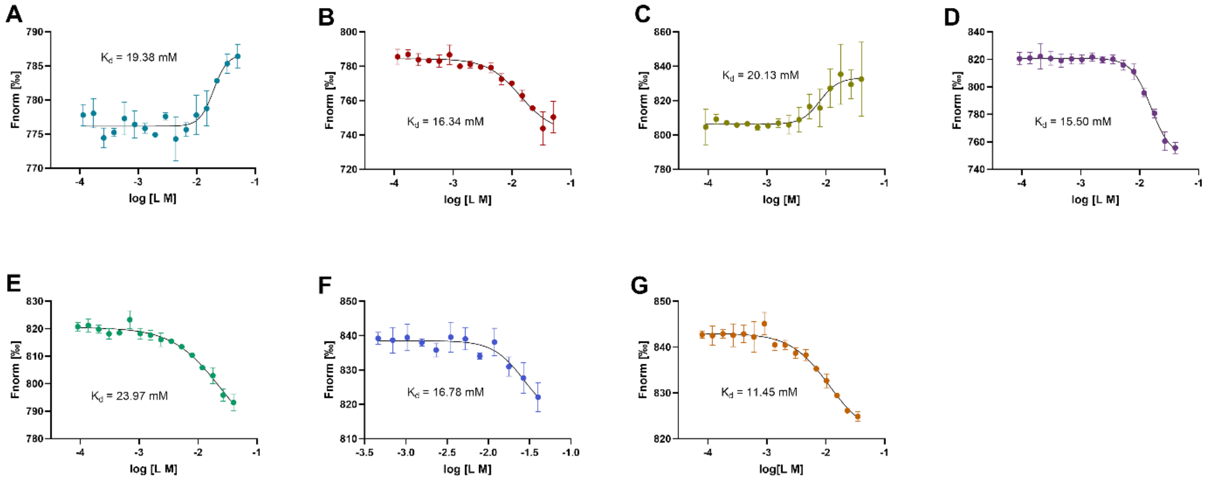


Figure 6S-1: Binding of the identified metabolite inhibitors to human GPI and their estimated $K_d$ values. (A) Histamine. (B) Maleate. (C) Hydroxylamine. (D) ADP. (E) Citraconate. (F) DHAP. (G) PEP. The binding curves were obtained using label-free MST. Compounds were titrated by a 2/3-fold serial dilution starting at 40-50 mM. Data were fitted using the Sigmoidal dose-response (variable slope) in Prism GraphPad. Each data point represents mean ± SD of triplicates or quadruplicates.

#### $K_d$ estimation of TPI hits

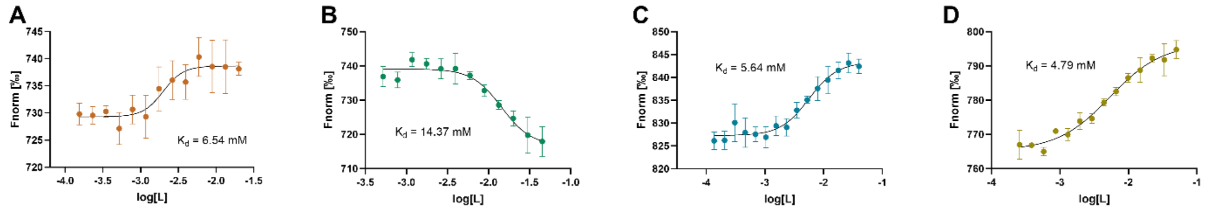


Figure 6S-2: Binding of the identified metabolite inhibitors to human TPI and their estimated $K_d$ values. (A) Methylmalonate. (B) Oxalate. (C) Glycerol-3-phosphate. (D) Glycerol-2-phosphate. The binding curves were obtained using label-free MST. Compounds were titrated by a 2/3-fold serial dilution starting from 40-50 mM. Data were fitted using the Sigmoidal dose-response (variable slope) in Prism GraphPad. Each data point represents mean ± SD of triplicates or quadruplicates.

### Figure 7S – DSF Thermal shift assays

In the thermal shift assay for GPI, all compounds, including the positive control (GTP), had a small-yet-measurable thermal shift, with a 1-2°C change in the $T_m$ over a concentration range of 0.3-40 mM (Fig. 7S). The GPI competitive inhibitor, 6PG, which was used as a positive control for GPI throughout the study, produced an unusual dose response pattern (U-shaped) in the DSF assay. Thus, another competitive inhibitor of GPI – GTP [4] was instead used as the positive control in the DSF assay for GPI. All the GPI hits, except for histamine and hydroxylamine, showed dose-response behavior. In most cases, the $T_m$ increased in the presence of the ligand, indicating stabilization of the protein structure by ligand binding. However, a decrease in $T_m$ was noticed in the presence of ADP, histamine, and oxalate. Such phenomenon is less common but may occur if ligands destabilize the native structure, due either specific or non-specific interaction, or if they preferentially bind to the unfolded state of the protein [5].

#### GPI thermal shift assays

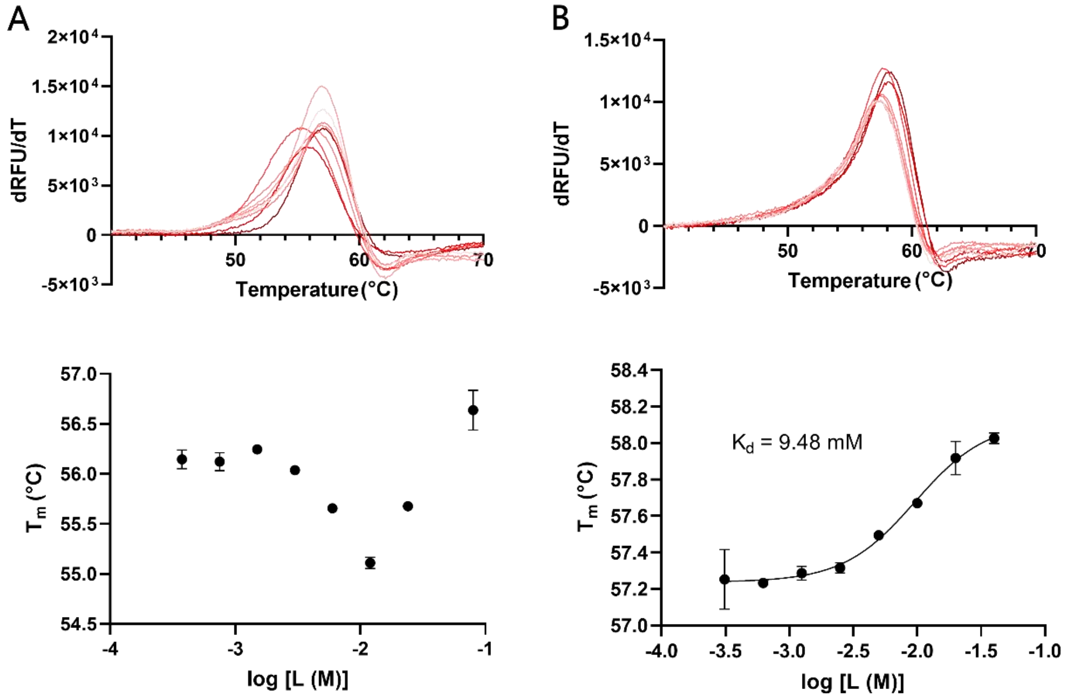


Figure 7S-1: DSF melting curves (dRFU/dT) and changes in the melting temperatures (°C) of GPI in response to competitive GPI inhibitors - 6PG (A) and GTP (B). The compounds were assessed at a concentration range of 0-40 mM. Data were fitted using the Sigmoidal dose-response (variable slope) in Prism GraphPad. Each data point represents mean ± SD of triplicates.

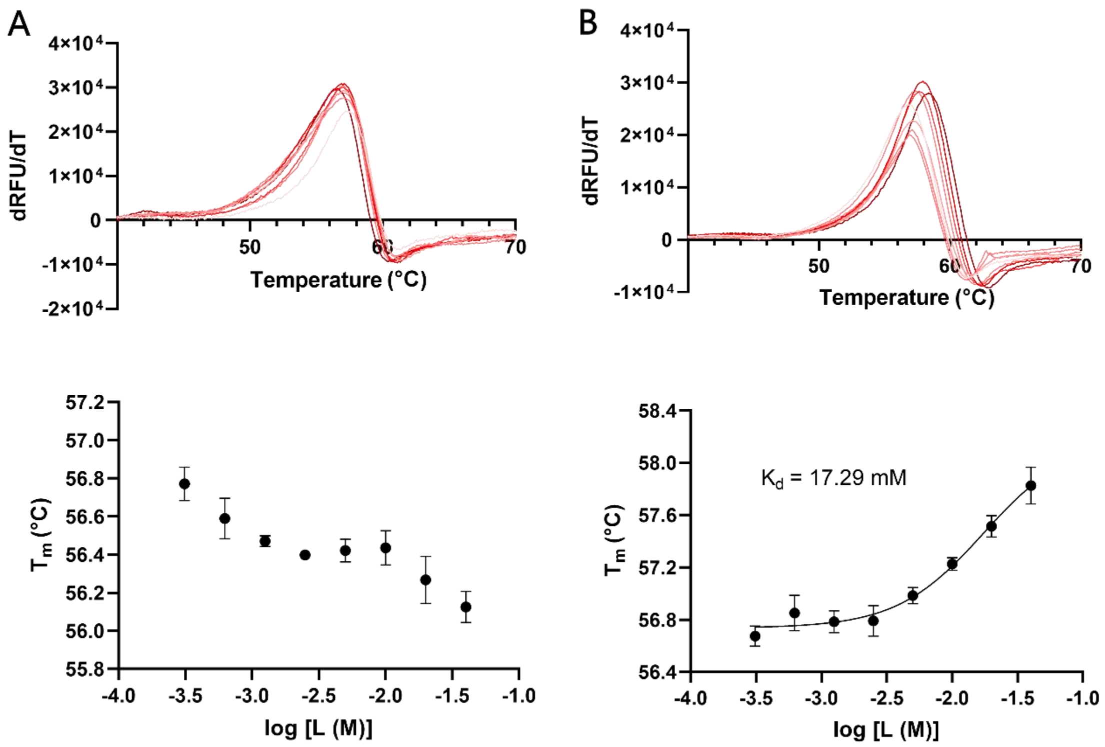


Figure 7S-2: DSF melting curves (dRFU/dT) and changes in the melting temperatures (°C) of GPI in the presence of histamine (A) and GTP (B). The compounds were assessed at a concentration range of 0-40 mM. Data were fitted using the Sigmoidal dose-response (variable slope) in Prism GraphPad. Each data point represents mean ± SD of triplicates.

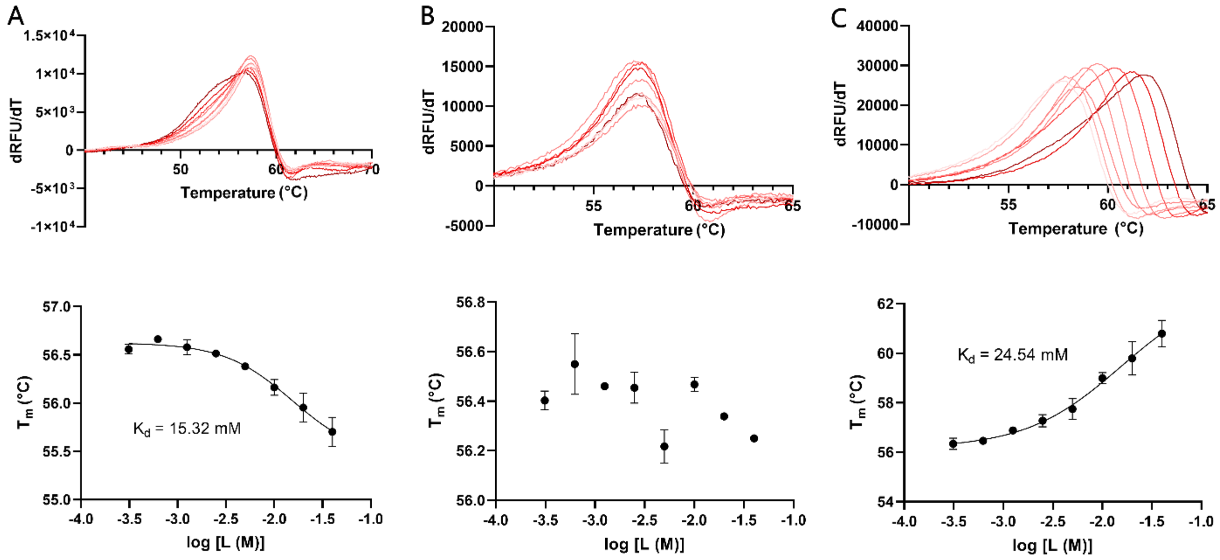


Figure 7S-3: DSF melting curves (dRFU/dT) and changes in the melting temperatures (°C) of TPI in response to varying concentrations (0-40 mM) of ADP (A), hydroxylamine (B), and citraconate (C). Data were fitted using the Sigmoidal dose-response (variable slope) in Prism GraphPad. Each data point represents mean ± SD of triplicates.

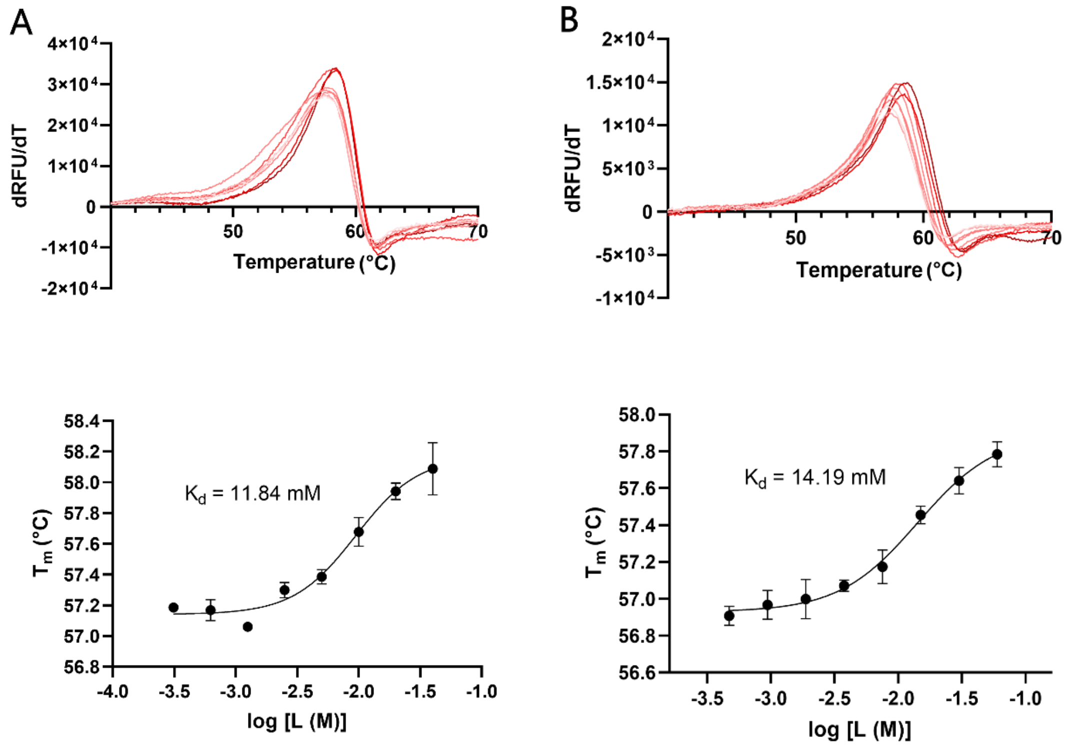


Figure 7S-4: DSF melting curves (dRFU/dT) and changes in the melting temperatures (°C) of GPI in response to competitive GPI inhibitors - DHAP (A) and PEP (B). The compounds were assessed at a concentration range of 0-40 mM. Data were fitted using the Sigmoidal dose-response (variable slope) in Prism GraphPad. Each data point represents mean ± SD of triplicates.

#### TPI thermal shift assays

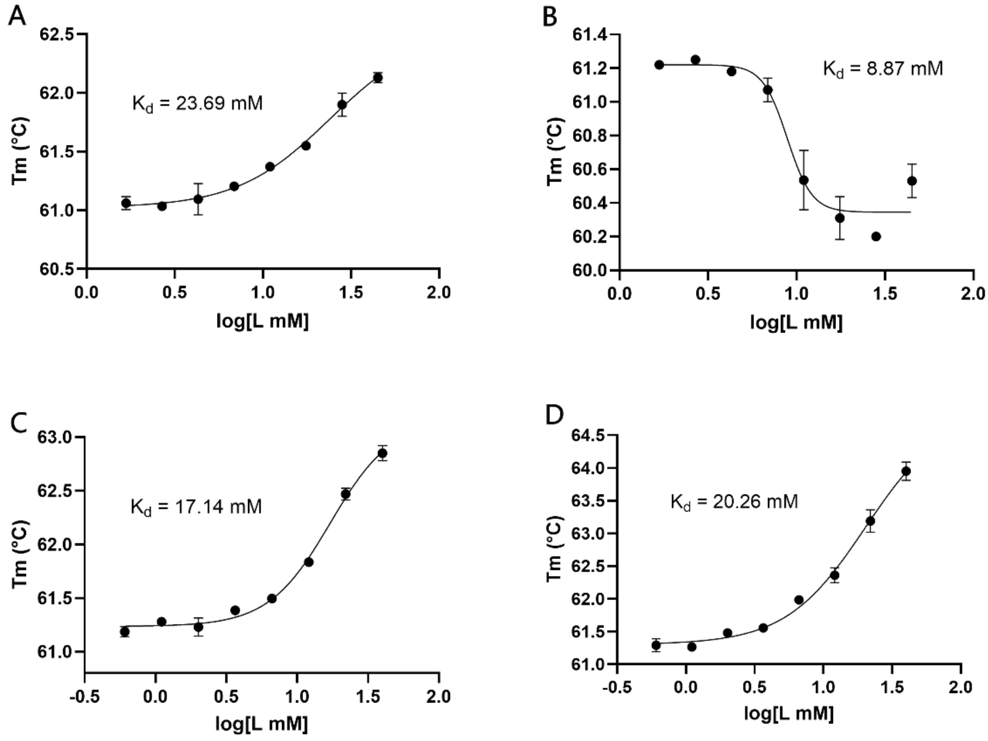


Figure 7S-5: DSF melting curves (dRFU/dT) and changes in the melting temperatures (°C) of TPI in response to varying concentrations (0-40 mM) of methylmalonate (A), oxalate (B), glycerol-3-phosphate (C), and glycerol-2-phospahte (D) . Data were fitted using the Sigmoidal dose-response (variable slope) in Prism GraphPad. Each data point represents mean ± SD of duplicates.

#### Figure 8S – Protein crystallization trials

In the GPI co-crystallization trials, all potential hits except for ADP, resulted in single crystals of medium sizes after 5 days of incubation (Fig. 8S-1). The addition of ADP caused immediate precipitation of the protein drops; therefore, soaking was performed (but no ligand density was observed). Co-crystallization of TPI with oxalate (OA) gave single needle-shaped crystals after 4-5 days of incubation (no ligand density observed in data). The other three hits, methylmalonate (MMA), glycerol-2-phosphate (G2P), and glycerol-3-phosphate (G3P), were soaked with Apo TPI crystals, as they did not produce any single crystals in the co-crystallization trials; only intertwined needles and sea urchins were observed (Fig. 8S-2). It was noticed that TPI tended to form small needle-like crystals in the presence of phosphate-containing compounds (e.g., PEP, G2P, and G3P).

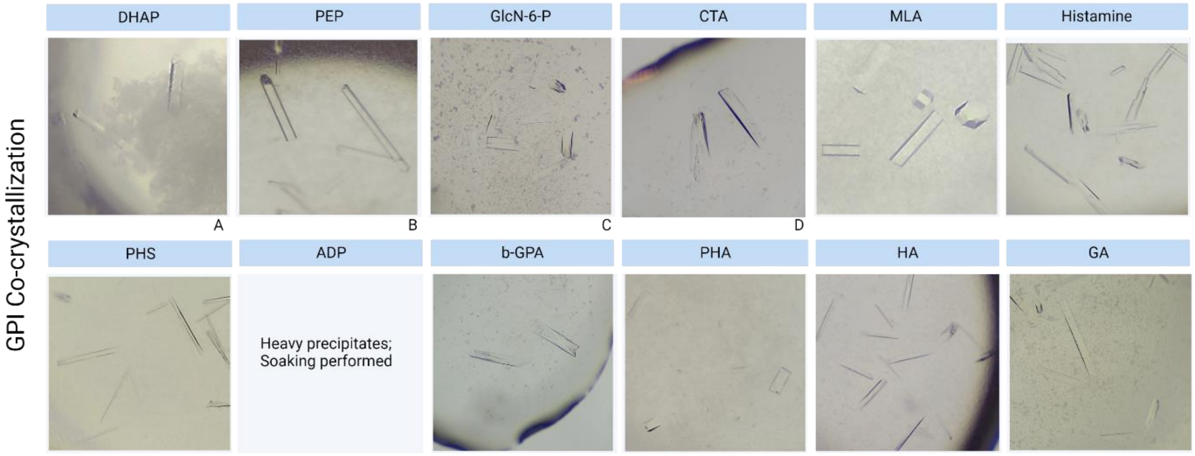


Figure 8S-1: Crystals produced from the co-crystallization trials of GPI with putative hits. The GPI crystals were grown from microseeds, in condition containing 21% PEG3500, 0.16 M $CaCl_2$, and 0.058 M HEPES, pH 7.0, at room temperature, using hanging-drop vapor diffusion method. Most of the crystals diffracted to 1.3-2.5 Å. Protein drops prepared from 1.5 µL of 8 mg/mL GPI solution (pre-incubated with ligands at 15-25 mM), 1.5 µL of reservoir, and 0.5 µL of 4-fold serially diluted seed solution. Crystals were visible after 2-5 days of incubation.

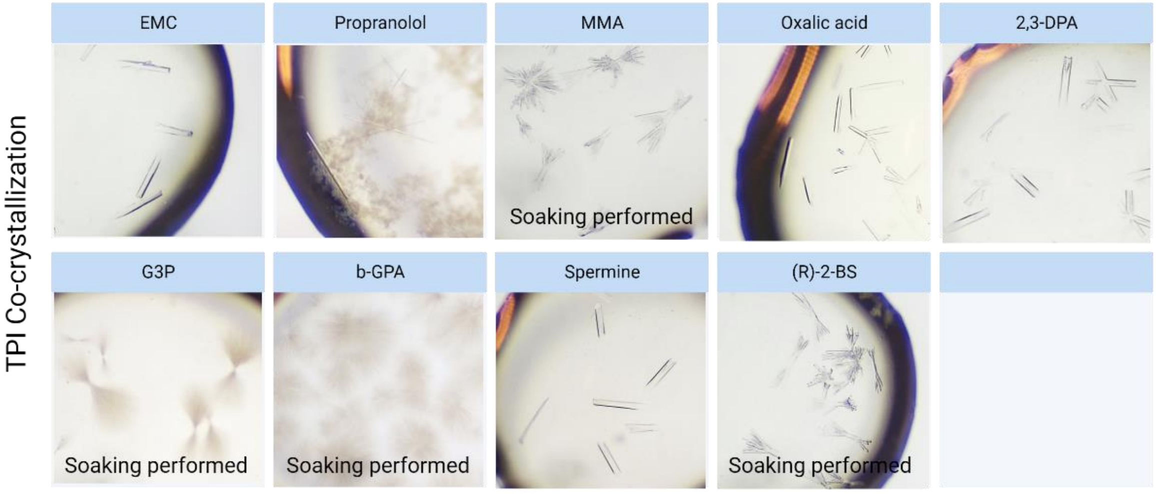


Figure 8S-2: Crystals produced from the co-crystallization trials of TPI with 9 potential inhibitors. The TPI crystals were grown from microseeds, in condition containing 24% PEG2000 MME and 0.14 M KBr, at room temperature, using hanging-drop vapor diffusion method. Protein drops prepared from 1.5 µL of 9 mg/mL TPI solution (pre-incubated with ligands at 20 mM), 1.5 µL of reservoir, and 0.5 µL of 7-fold serially diluted seed solution. Crystals were seen after 2-3 days of incubation.

### Figure 8S – GPI and TPI crystals in complex with inhibitors

#### Ligands bound to human GPI

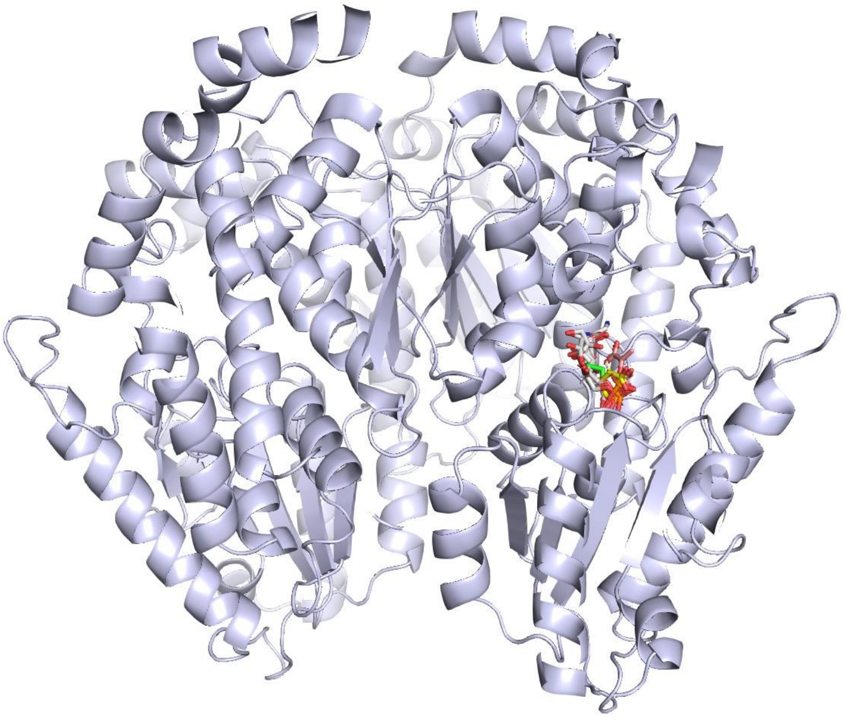

Figure 8S-1: Identified hits bound to human GPI and aligned with known inhibitors (E4P, PA5, 6PG) – all found in the enzyme active site.

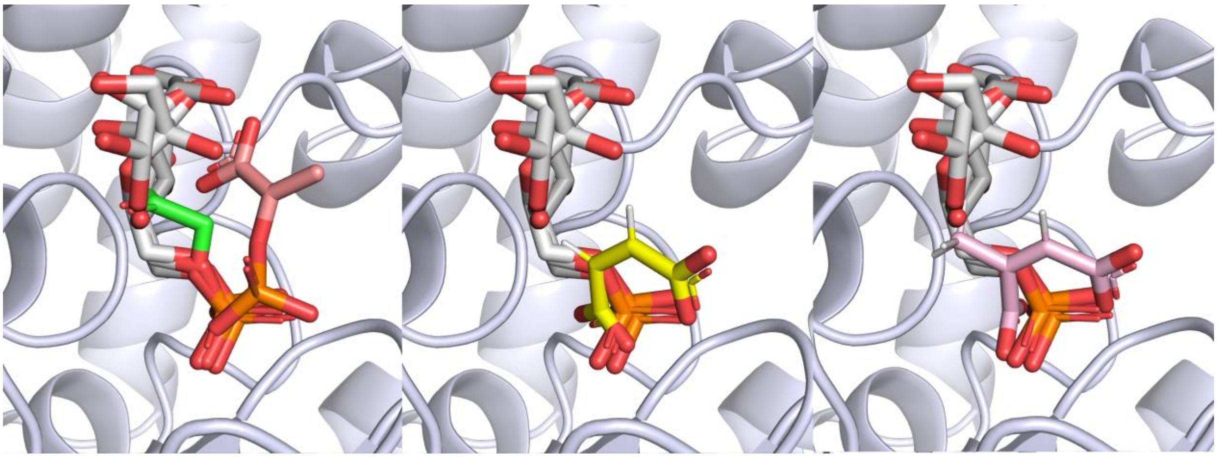

Figure 8S-2: Binding poses of DHAP (green), PEP (red), MAE (yellow), and CTA (pink) complexed with human GPI, and superimposed with known inhibitors (grey; E4P, PA5, 6PG) within the enzyme active site.

#### Ligands bound to human TPI

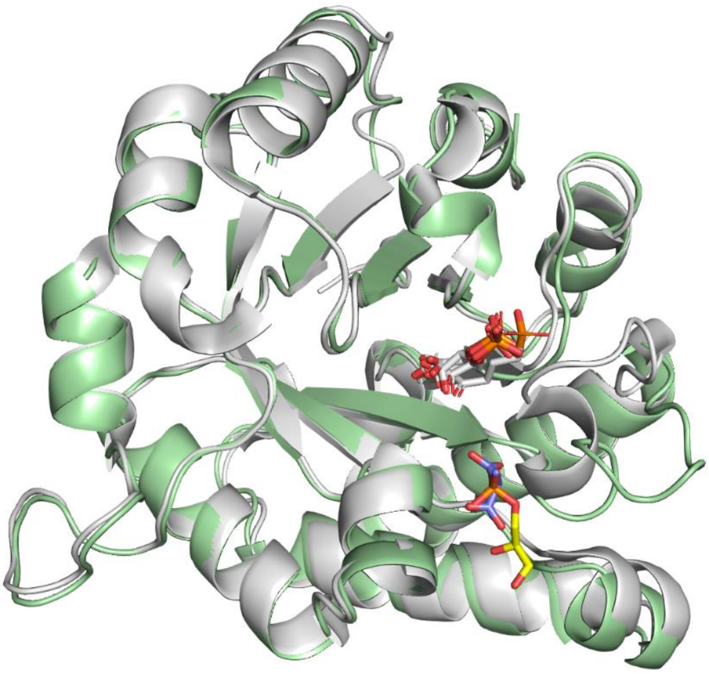

Figure 8S-2: Binding poses of identified TPI hits, G3P (yellow) and MMA (blue), and known inhibitors (grey; E4P, PA5, 6PG) complexed with TPI. The published human TPI structure (1HTI), shown in grey, is used as a representative showing the closed-loop conformation upon orthosteric binding.

#### Table 2S – Structural comparison

Table 2S: Cα Root-mean-square deviation (RMSD) distances after monomer subunit alignments (RMSD asymmetric unit) and alignments to a published apo structure (RMSD apo structure). The PDB IDs for the apo structure references were 1IAT and 2JK2 for GPI and TPI respectively. Alignments and RMSD calculations were performed in UCSF Chimera.

| Crystal structure | RMSD asymmetric unit (Å) | RSMD apo structure |
|---|---|---|
| **GPI_CTA** | 0.225 | 0.256 |
| **GPI_DHAP** | 0.230 | 0.257 |
| **GPI_PEP** | 0.335 | 0.268 |
| **GPI_MAE** | 0.483 | 0.382 |
| **TPI_G3P** | N/A | 0.586 |
| **TPI_MAE** | N/A | 0.587 |

#### Figure 9S – Structural comparison

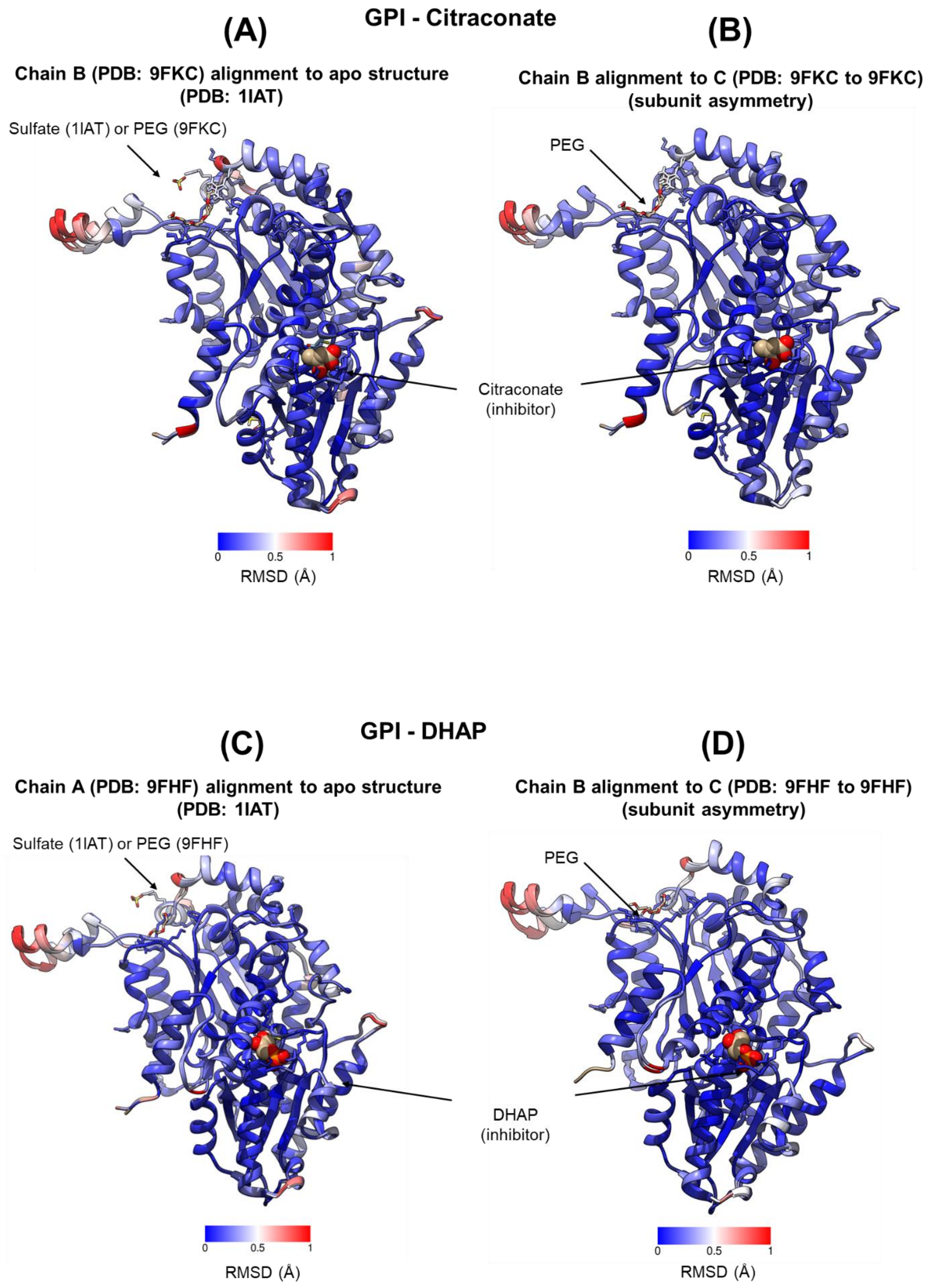

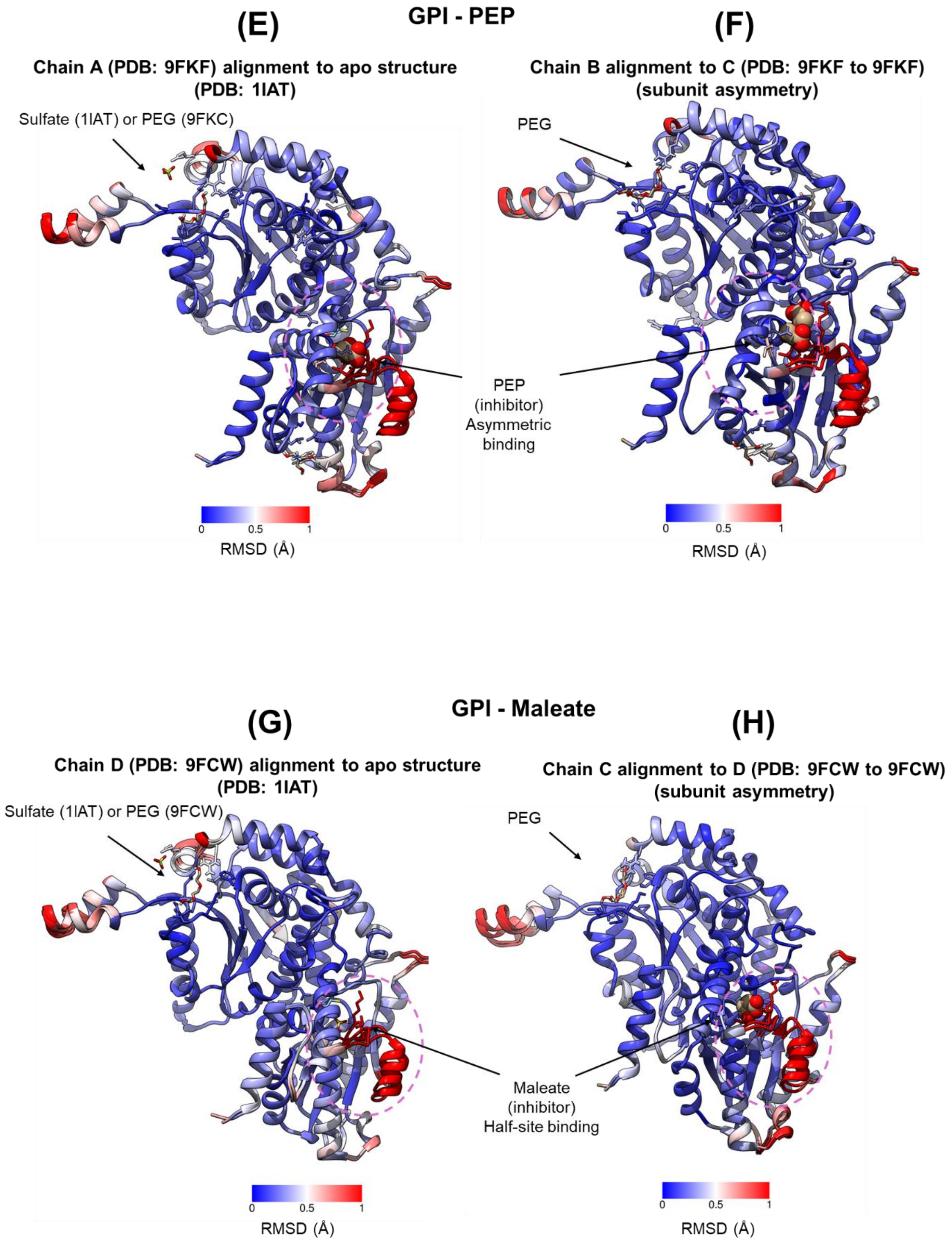
GPI - PEP
(E)
Chain A (PDB: 9FKF) alignment to apo structure (PDB: 1IAT)
Sulfate (1IAT) or PEG (9FKC)
0
0.5
1
RMSD (Å)
(F)
Chain B alignment to C (PDB: 9FKF to 9FKF) (subunit asymmetry)
PEG
PEP (inhibitor) Asymmetric binding
0
0.5
1
RMSD (Å)
GPI - Maleate
(G)
Chain D (PDB: 9FCW) alignment to apo structure (PDB: 1IAT)
Sulfate (1IAT) or PEG (9FCW)
0
0.5
1
RMSD (Å)
(H)
Chain C alignment to D (PDB: 9FCW to 9FCW) (subunit asymmetry)
PEG
Maleate (inhibitor) Half-site binding
0
0.5
1
RMSD (Å)

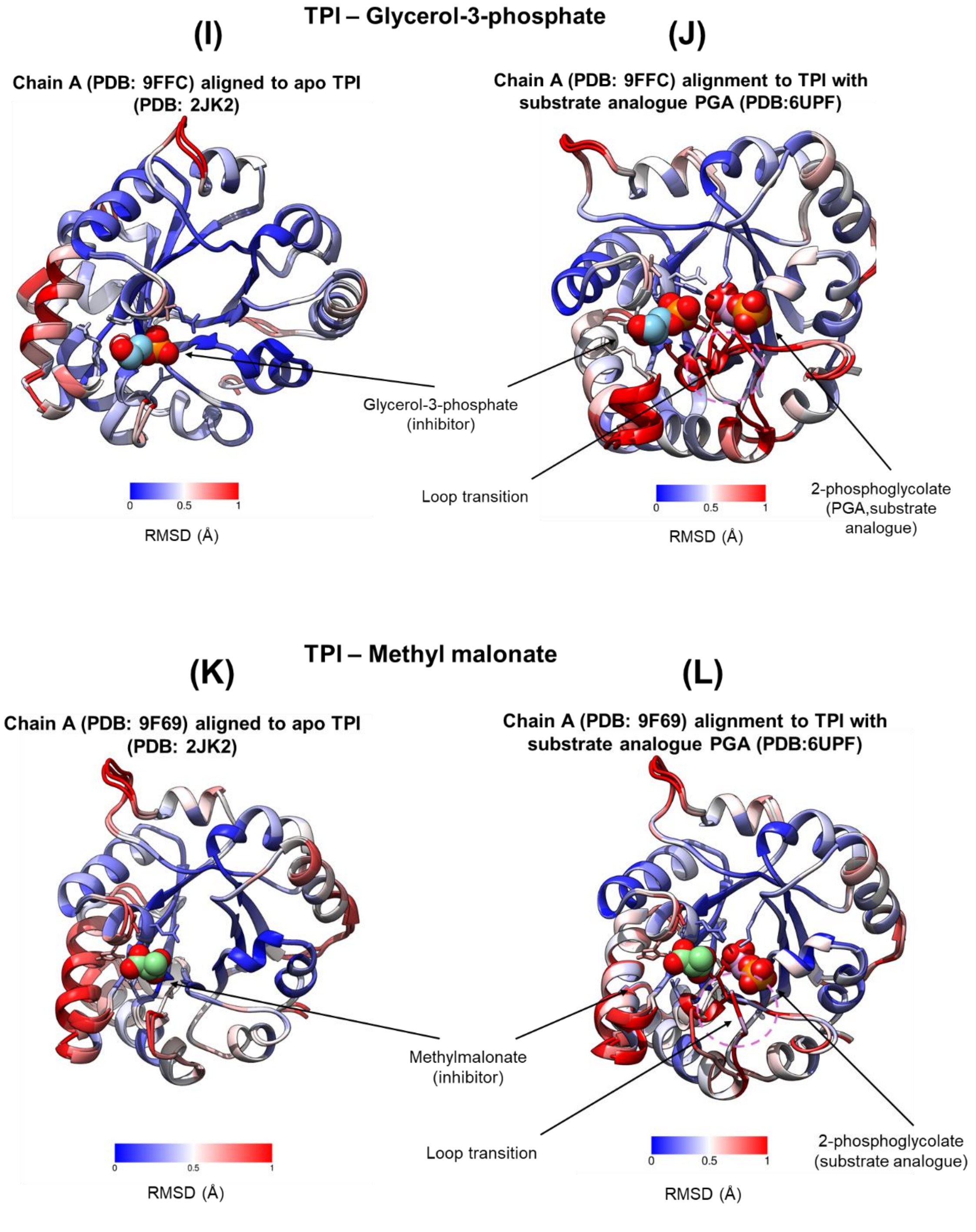


Figure 9S. RMSD color rendering of structures. The left panes show alignment of the structures to published apo structures and the right pane shows subunit asymmetry within the same structure (only applies to GPI, as TPI has a monomer in the asymmetric unit). Panes A-H and I-L represent GPI and TPI structures, respectively, utilizing apo PDB IDs 1IAT and 2JK2 for GPI and TPI, respectively. For TPI, the right panes show alignment to a structure containing a substrate analogue, 2-phosphoglycolate (PGA; PDB ID: 6UBF). A loop transition upon substrate analogue binding (PGA) is highlighted, which is absent in both TPI structures with inhibitor bound. Alignments and RMSD calculations were performed in UCSF Chimera.

#### Figure 10S – Cellular flux assays

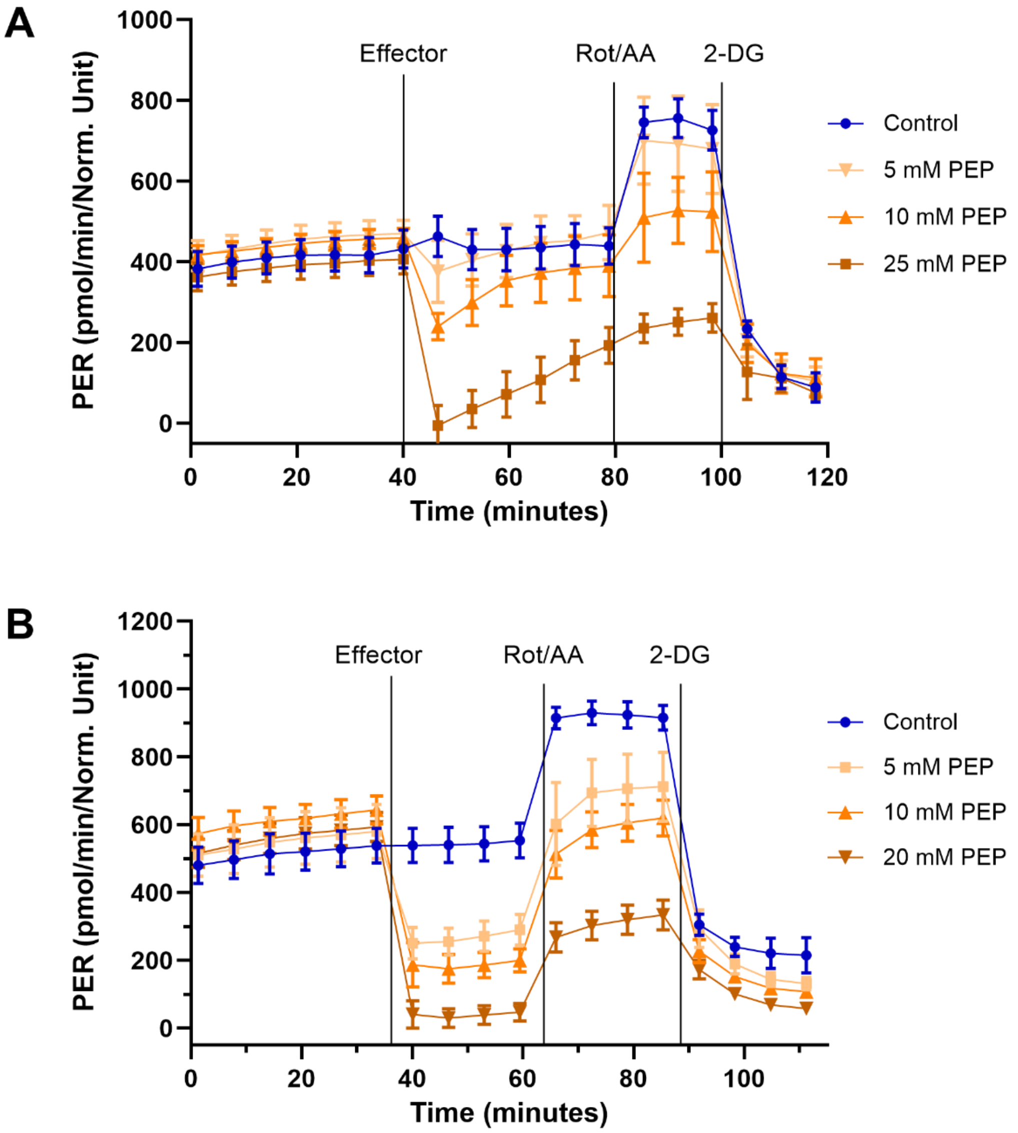


Figure 10S: Real-time analysis of proton efflux rate (PER) for Hulec-5a (A) and U2OS (B) cells, measured at 37°C, using Seahorse XFe96 analyzer, before and after injection of effector (PEP), mitochondrial inhibitors (Rot/AA), and glycolytic inhibitor (2-DG). After the baseline measurement, effectors were added to the cells to a final concentration of 5, 10, and 20-25 mM. The PER data were normalized to relative viable cells in each well, quantified with Prestoblue and absorption at 570 and 600 nm. Each data point represents the mean ± SD of six replicates.
