## Supplementary data B for "Human Glycolysis Isomerases are Inhibited by Weak Metabolite Modulators"

#### List of metabolites screened against GPI and TPI

MetaSci metabolites – Box 1-13 (1250 compounds in total)

Custom library – 21 compounds from Sigma-Aldrich

| Column | Description |
|---|---|
| Assayed | Green = compounds assayed using a combination of activity – and MST binding assays. |
| MST assay | Green = screened using MST assay |
| Activity assay | Green = screened using absorbance assay<br>Yellow = screened using luminescence assay |
| Abs interference | Dark grey = compounds caused interference in the absorbance assay |
| AF or quenching | Dark grey = compounds with auto-fluorescent or fluorescence-quenching properties |
| Initial MST hit | Blue = compounds showing changes in MST > 5 units |
| Initial activity hit | Red = compounds exhibiting activity changes > 20% |
| G6PDH/GADPH hit | Red = compounds showing activity changes > 15% when screening against the coupling enzyme |
| DR pattern | Orange = compounds exhibiting dose-repose pattern in the activity assay against the target<br>Dark grey = dose-response effects, but low reproducibility |
| Frequent hitter | Purple = compounds show up as initial hits in both the GPI and TPI screens |

### GPI screening: MetaSci – Box 1

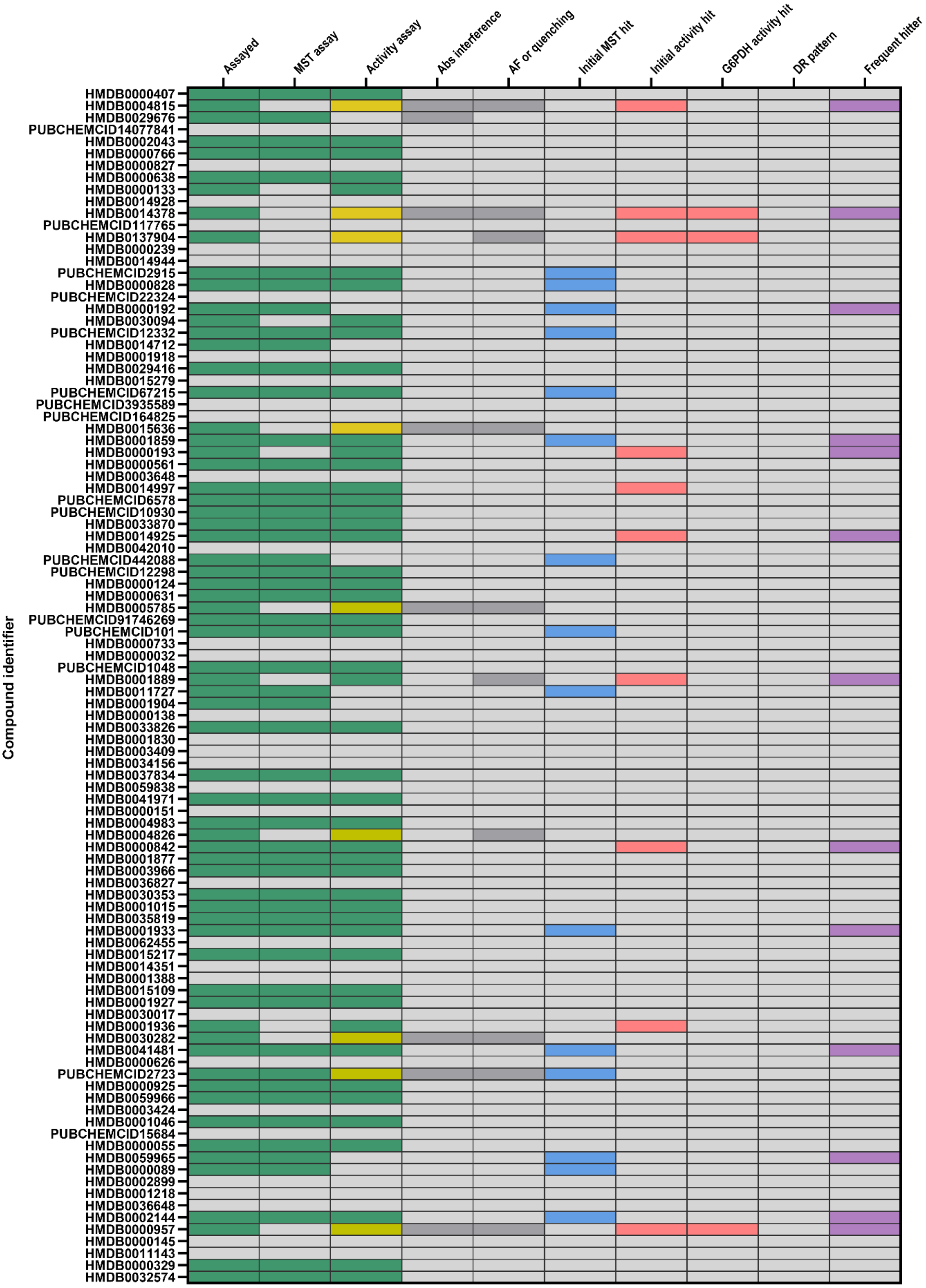

### GPI screening: MetaSci – Box 2

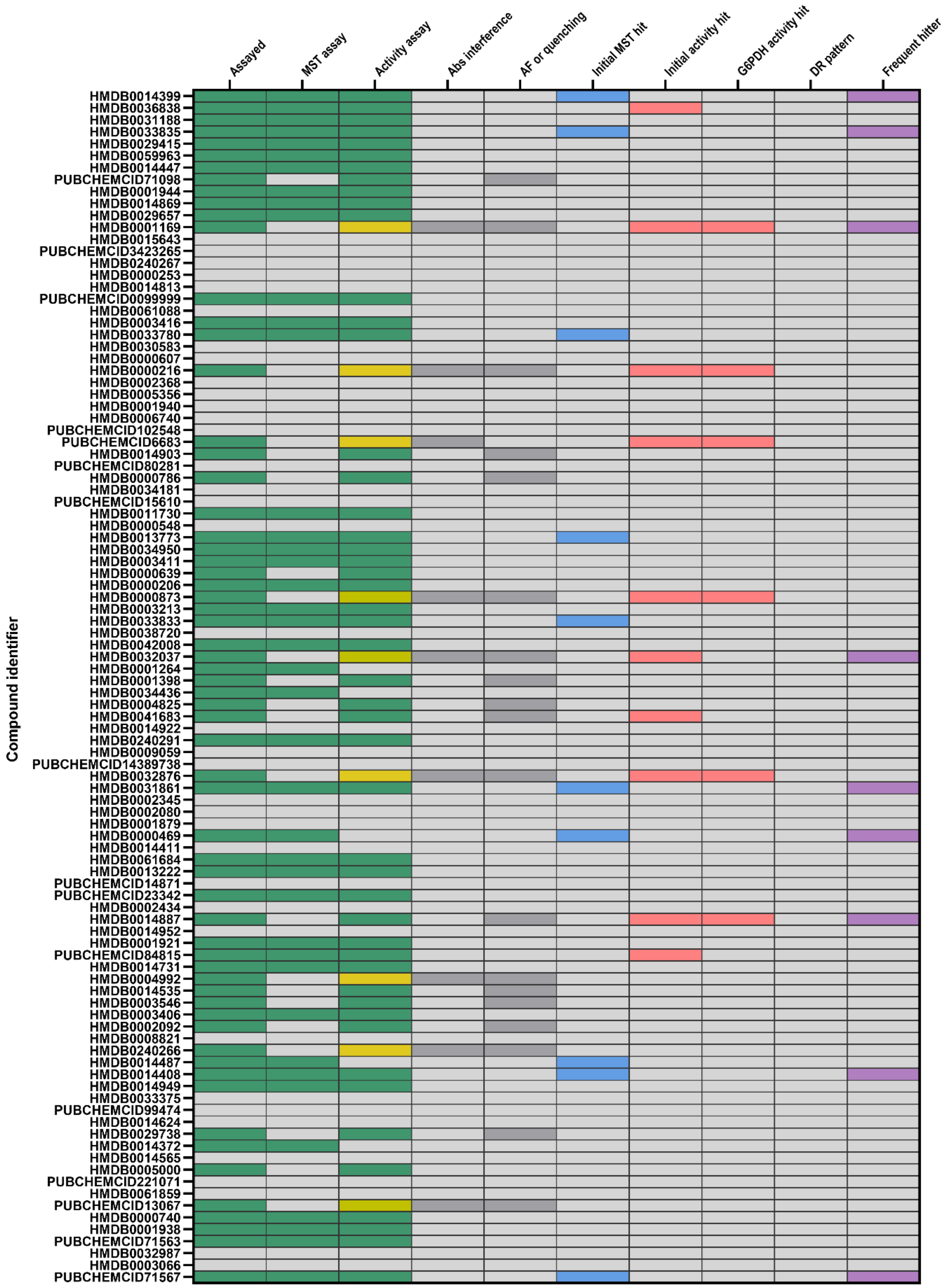

### GPI screening: MetaSci – Box 3

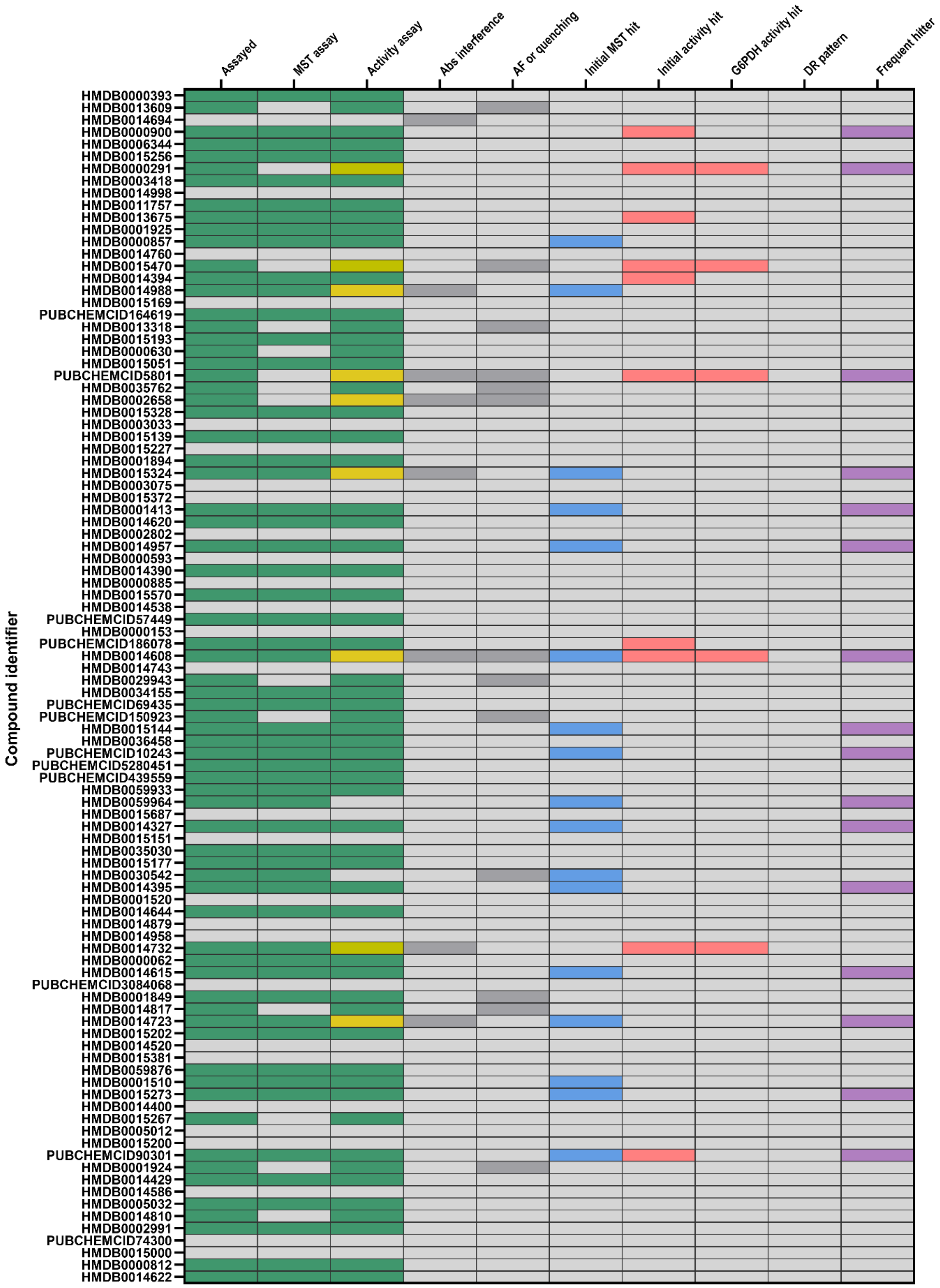

GPI screening: MetaSci – Box 4

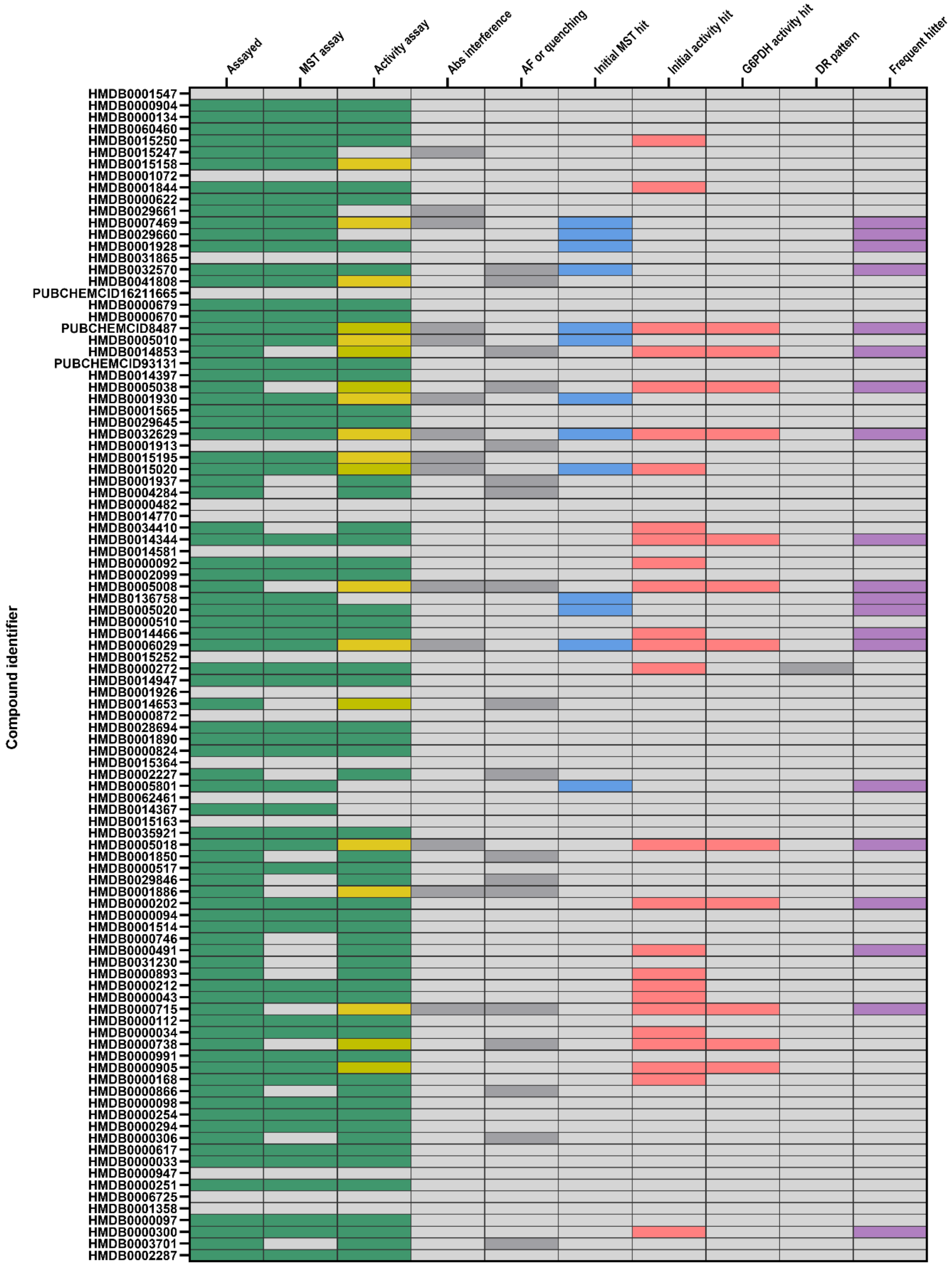

### GPI screening: MetaSci – Box 5

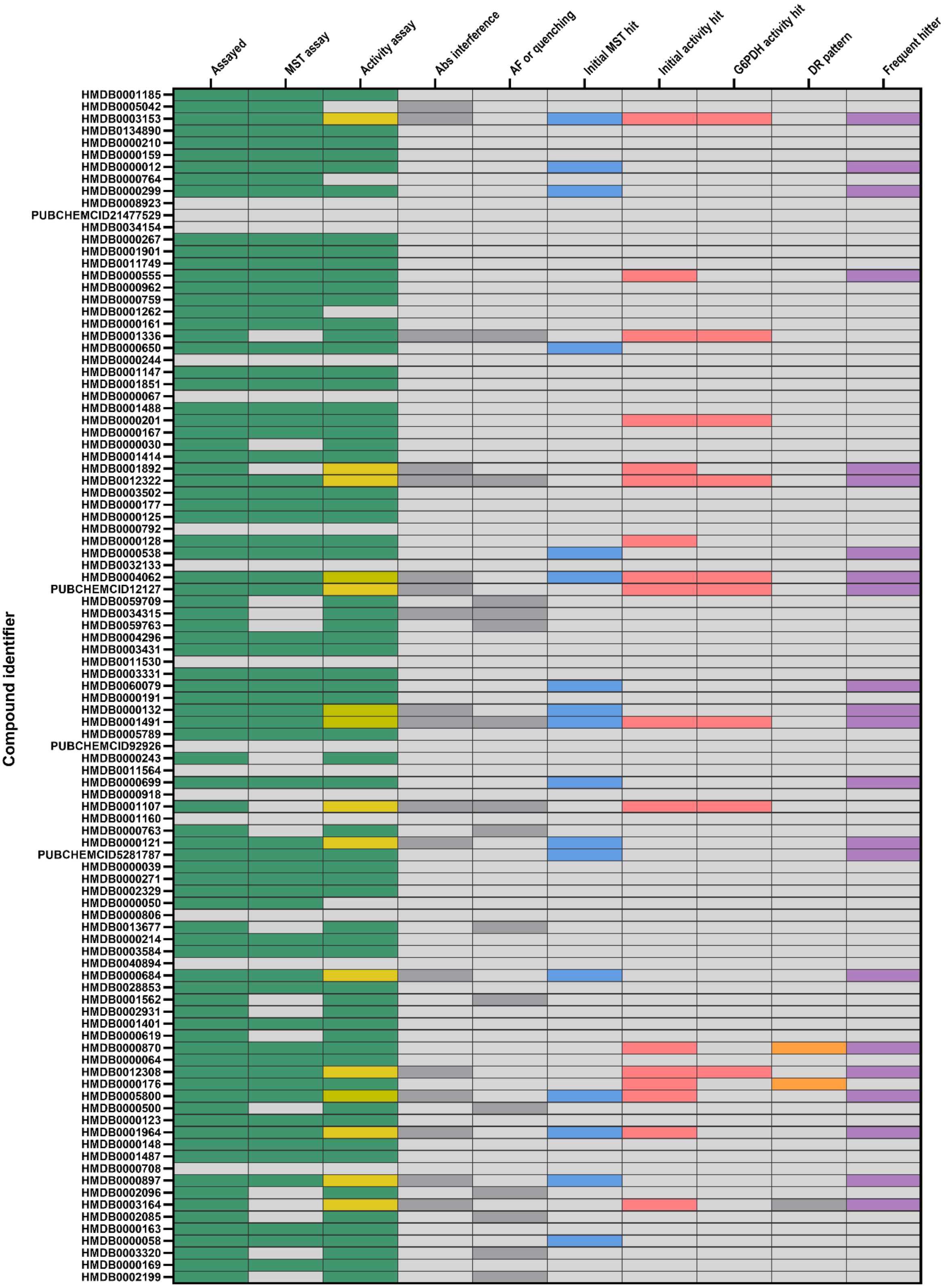

### GPI screening: MetaSci – Box 6

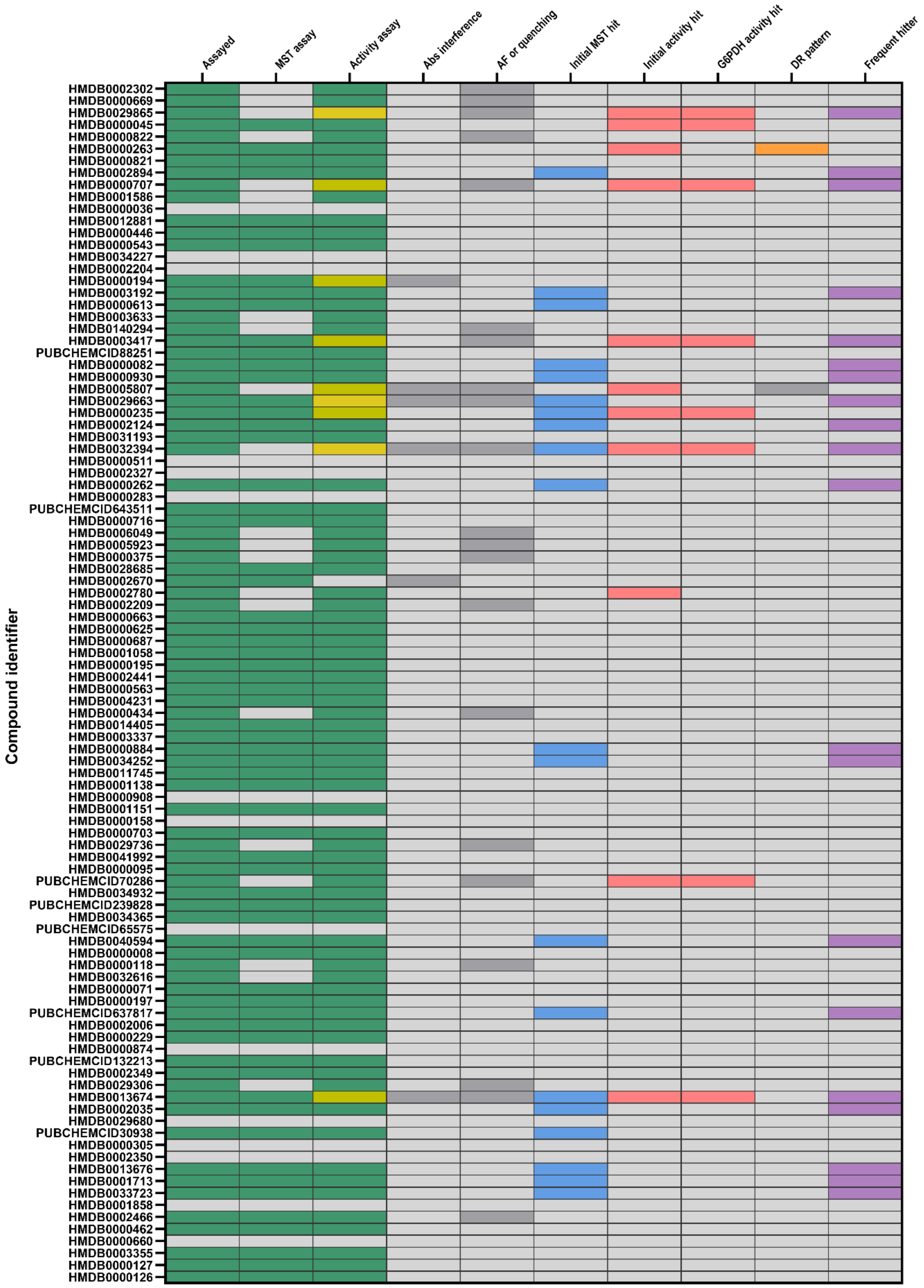

### GPI screening: MetaSci – Box 7

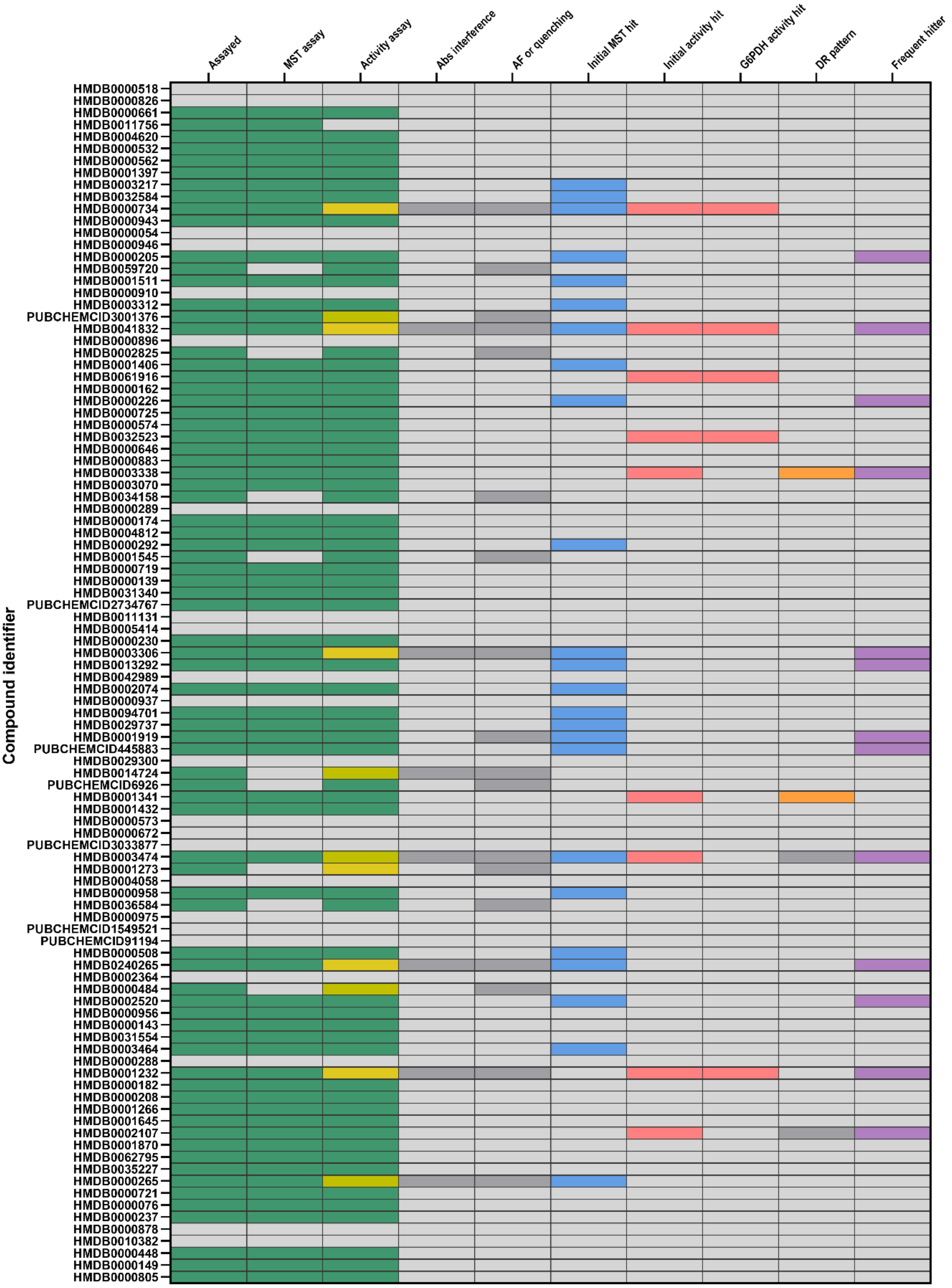

GPI screening: MetaSci – Box 8

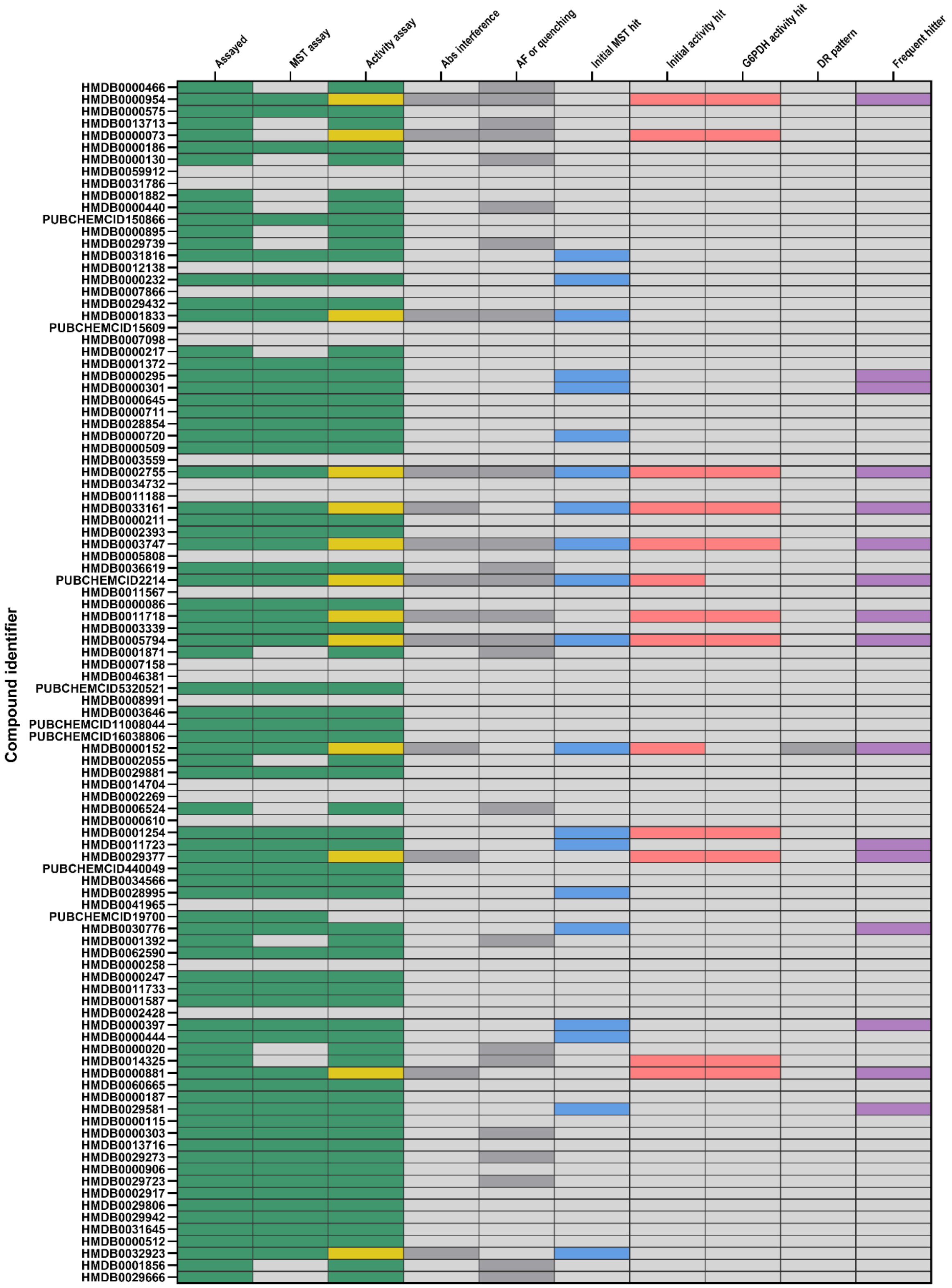

GPI screening: MetaSci – Box 9

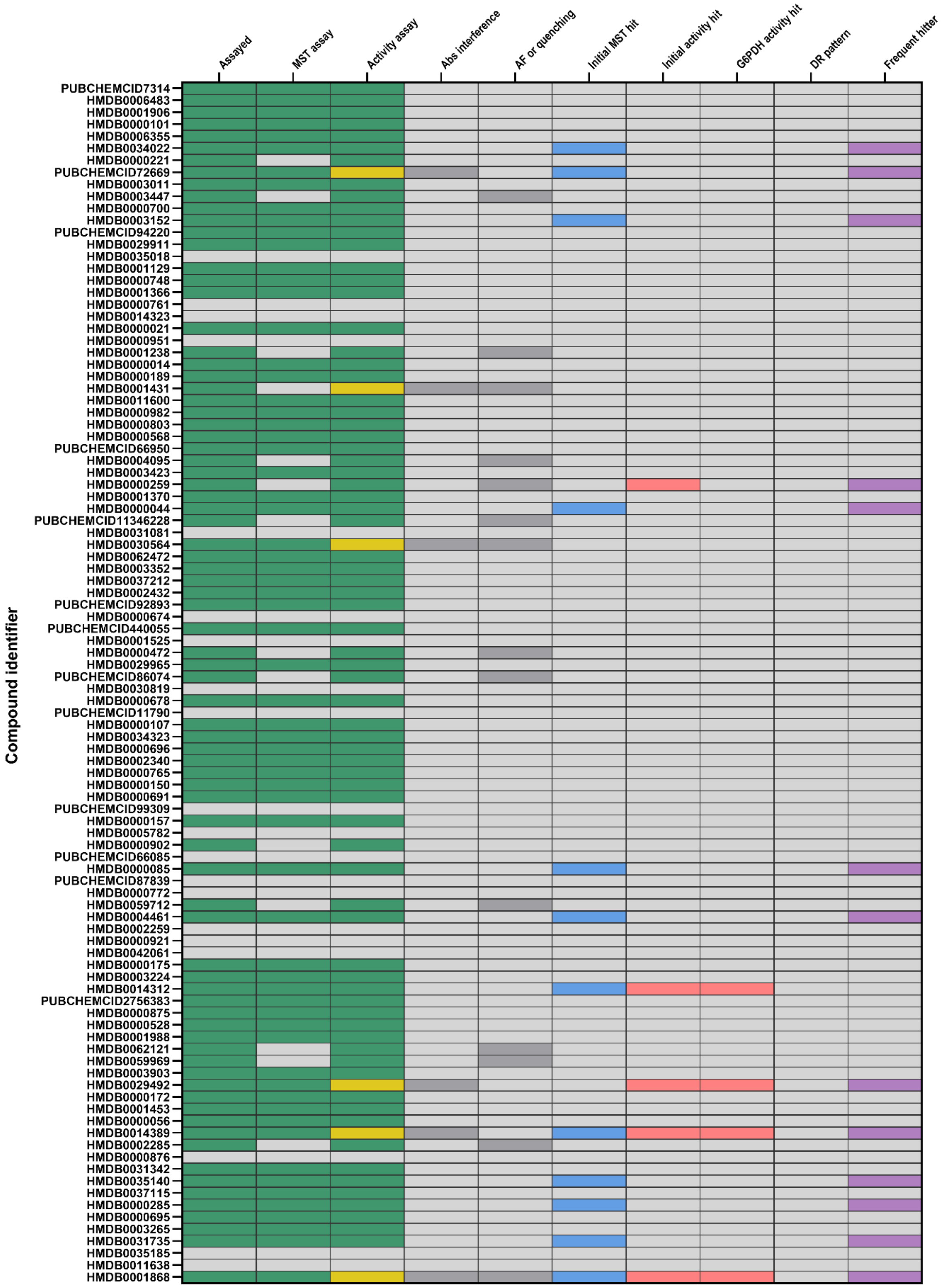

### GPI screening: MetaSci – Box 10

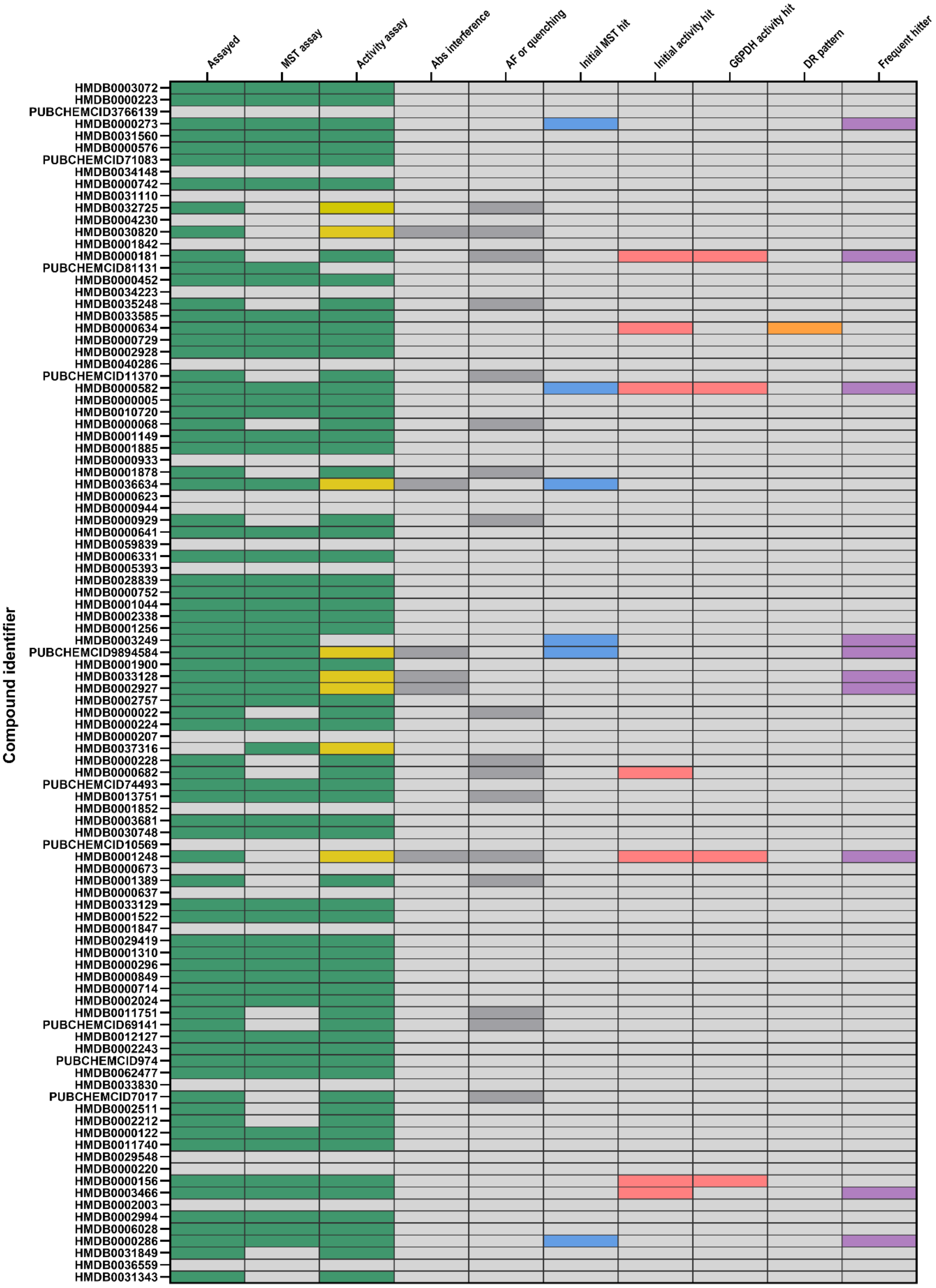

### GPI screening: MetaSci – Box 11

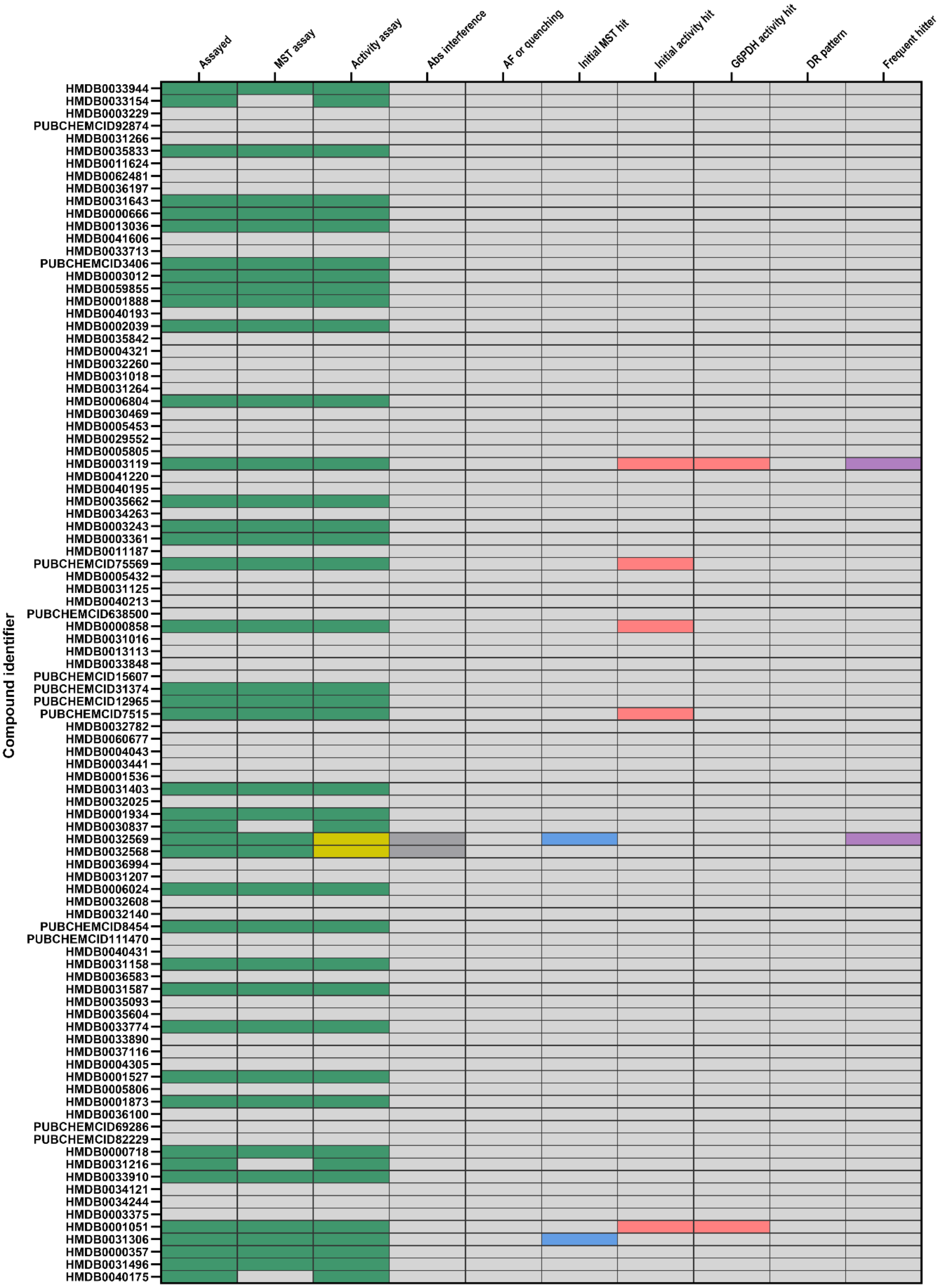

### GPI screening: MetaSci – Box 12

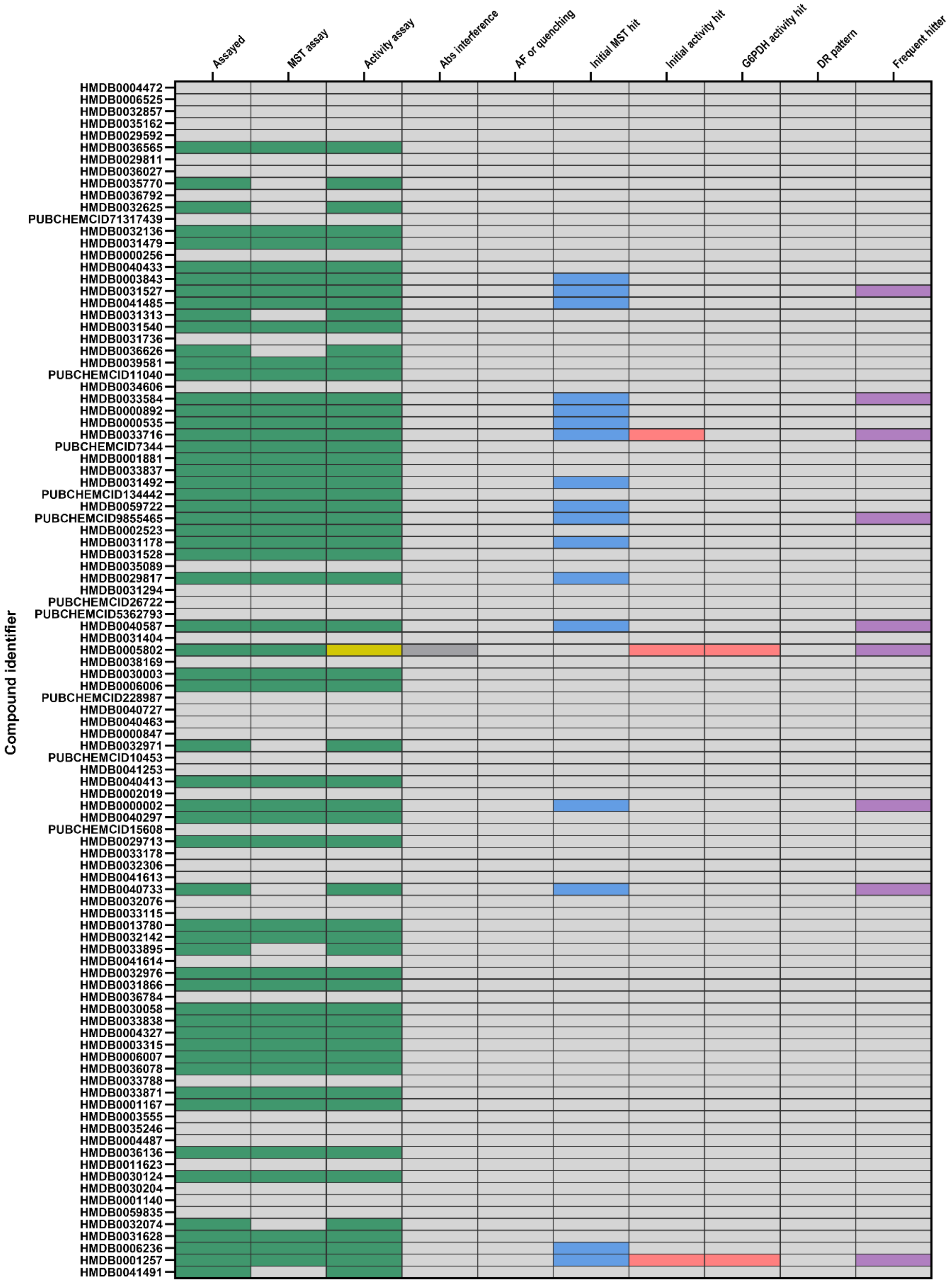

### GPI screening: MetaSci – Box 13 + Custom library

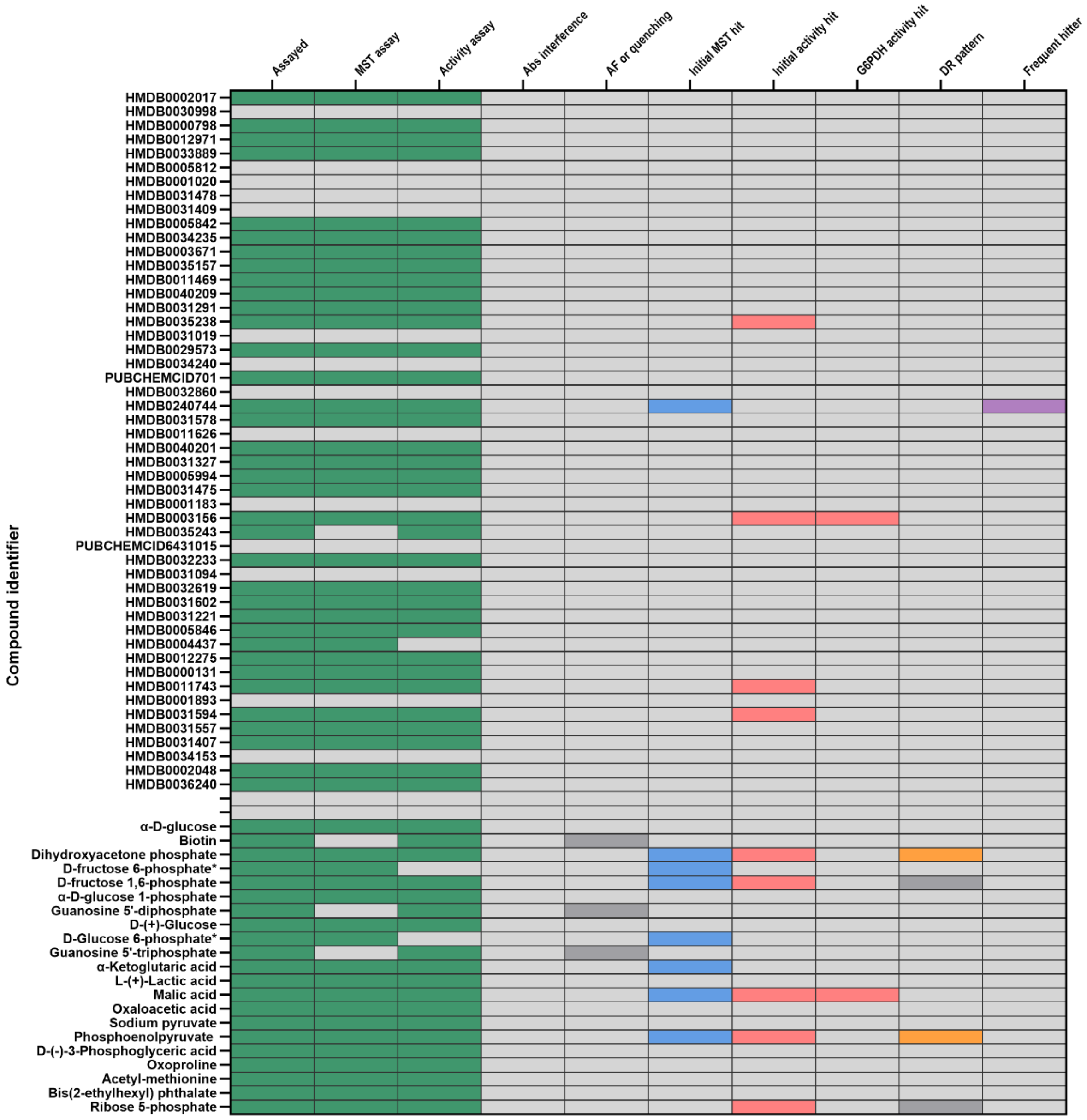


* Enzyme substrate

### TPI screening: MetaSci – Box 1

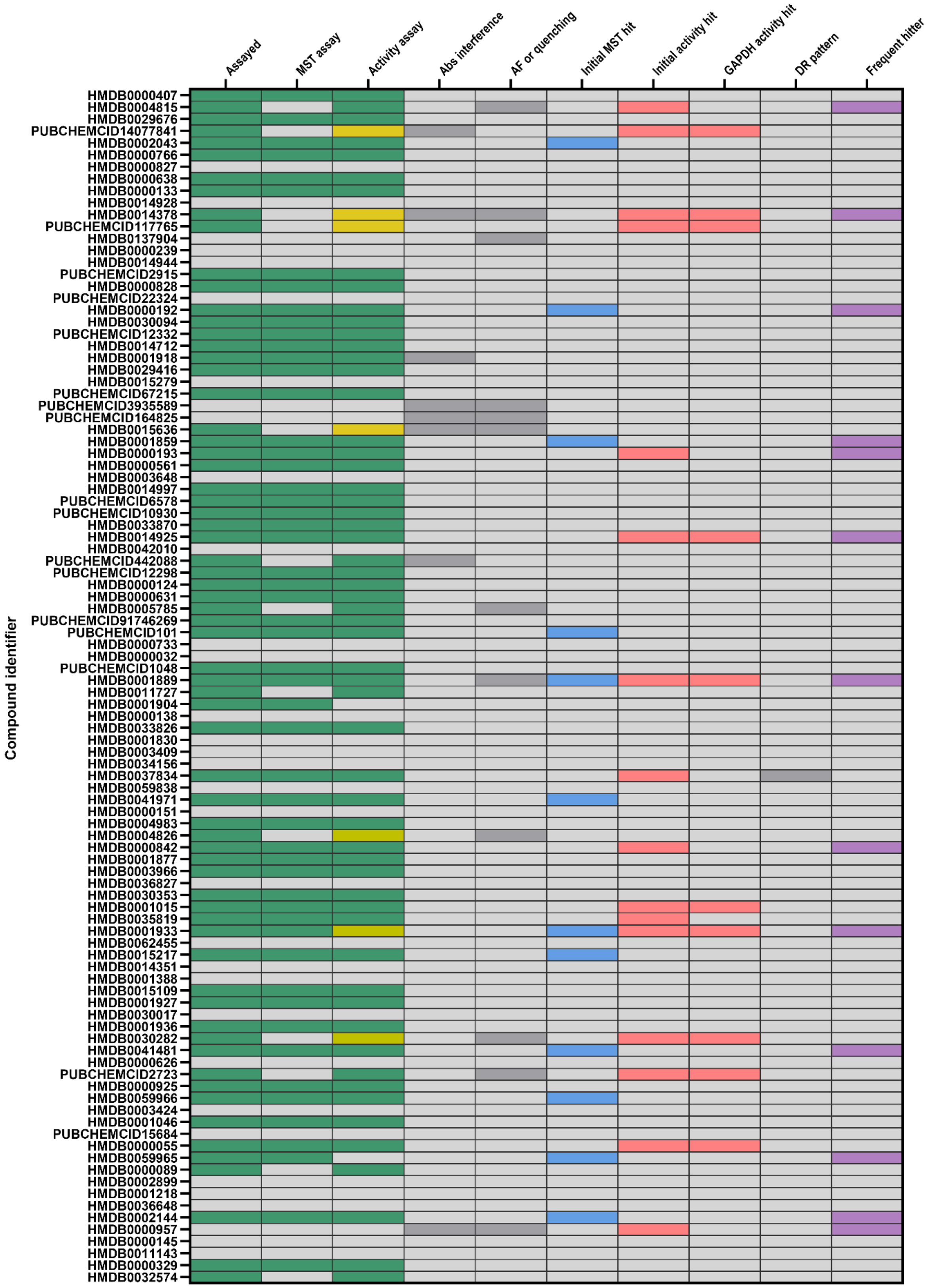

TPI screening: MetaSci – Box 2

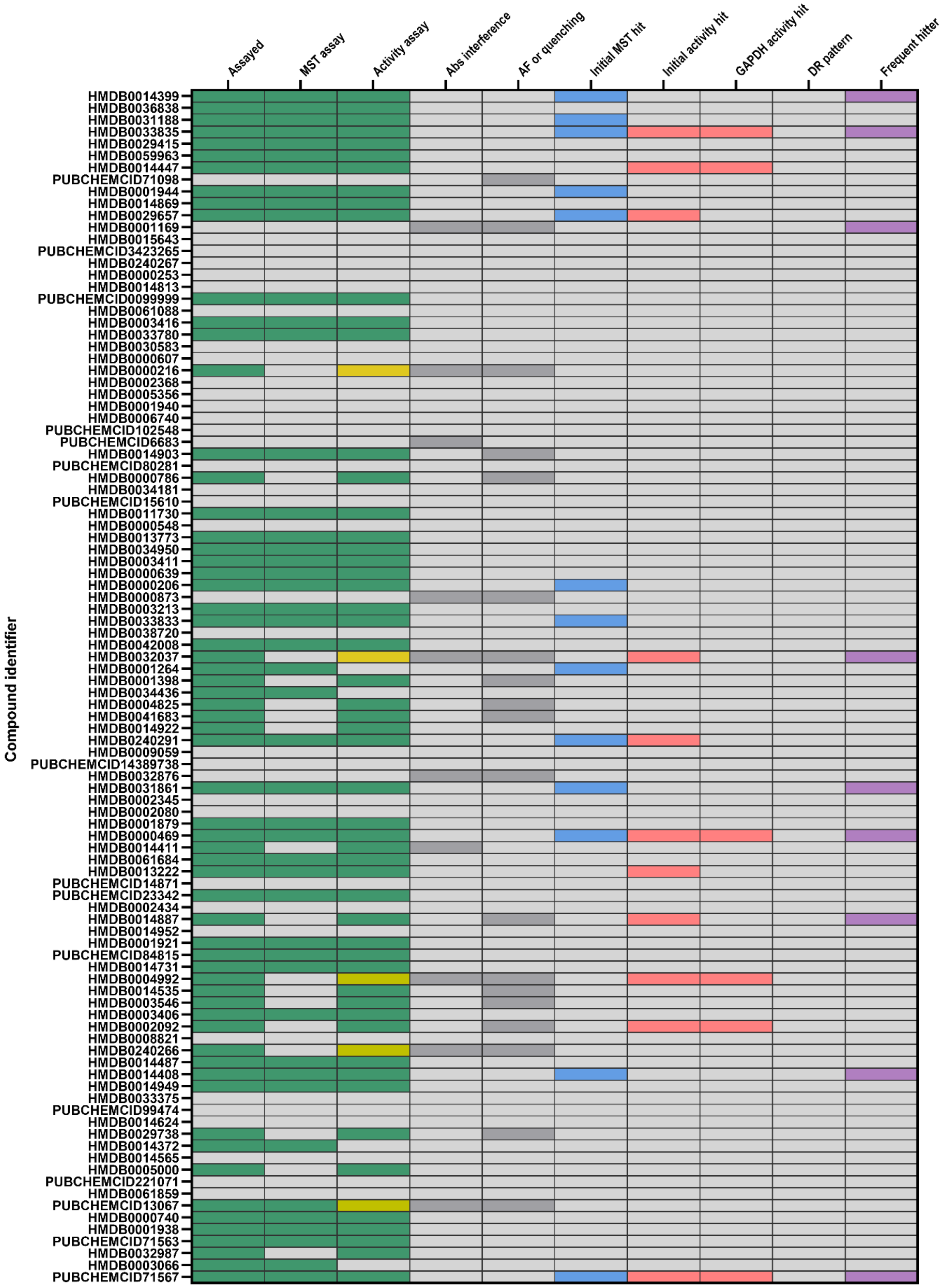

TPI screening: MetaSci – Box 3

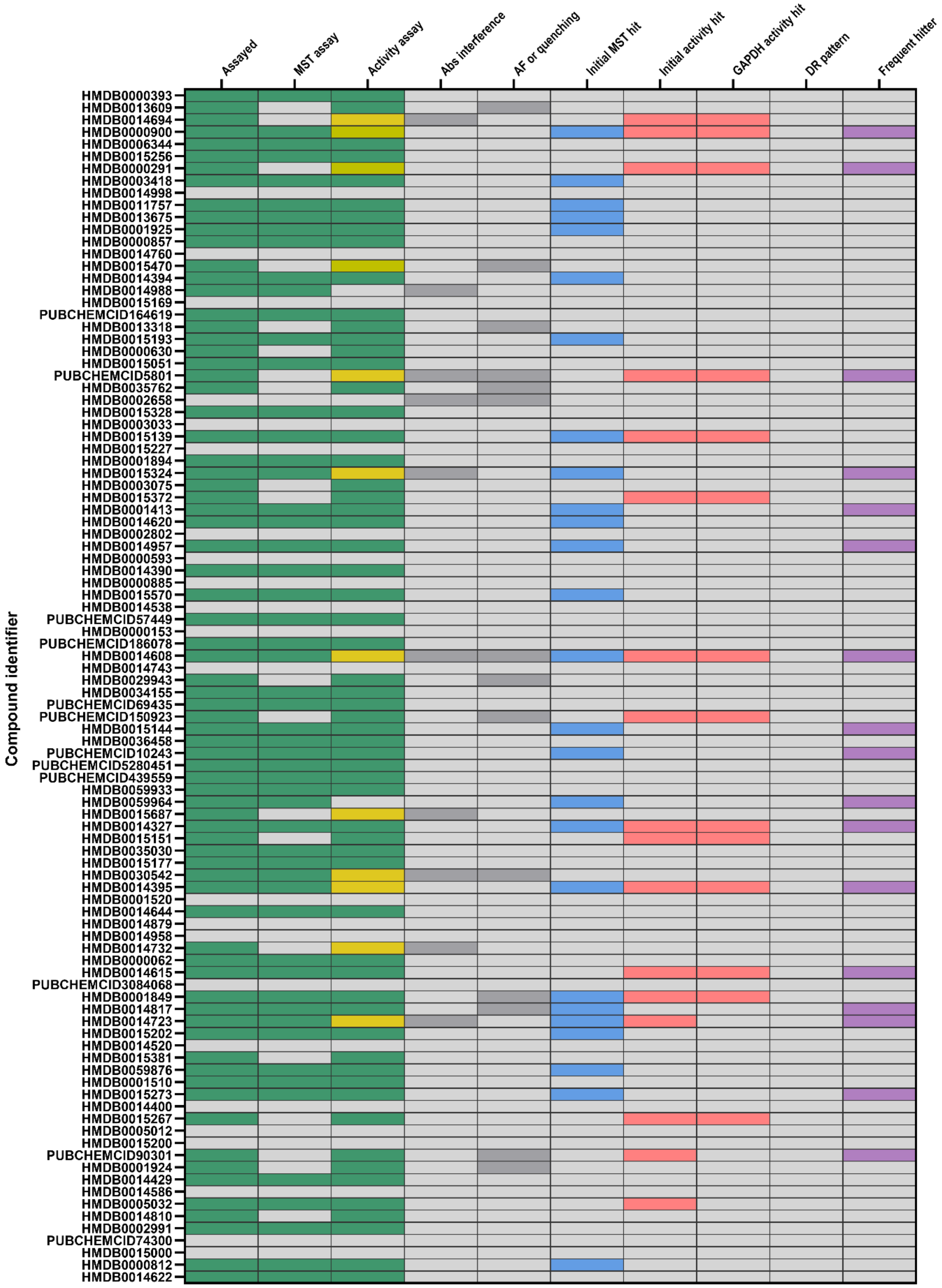

### TPI screening: MetaSci – Box 4

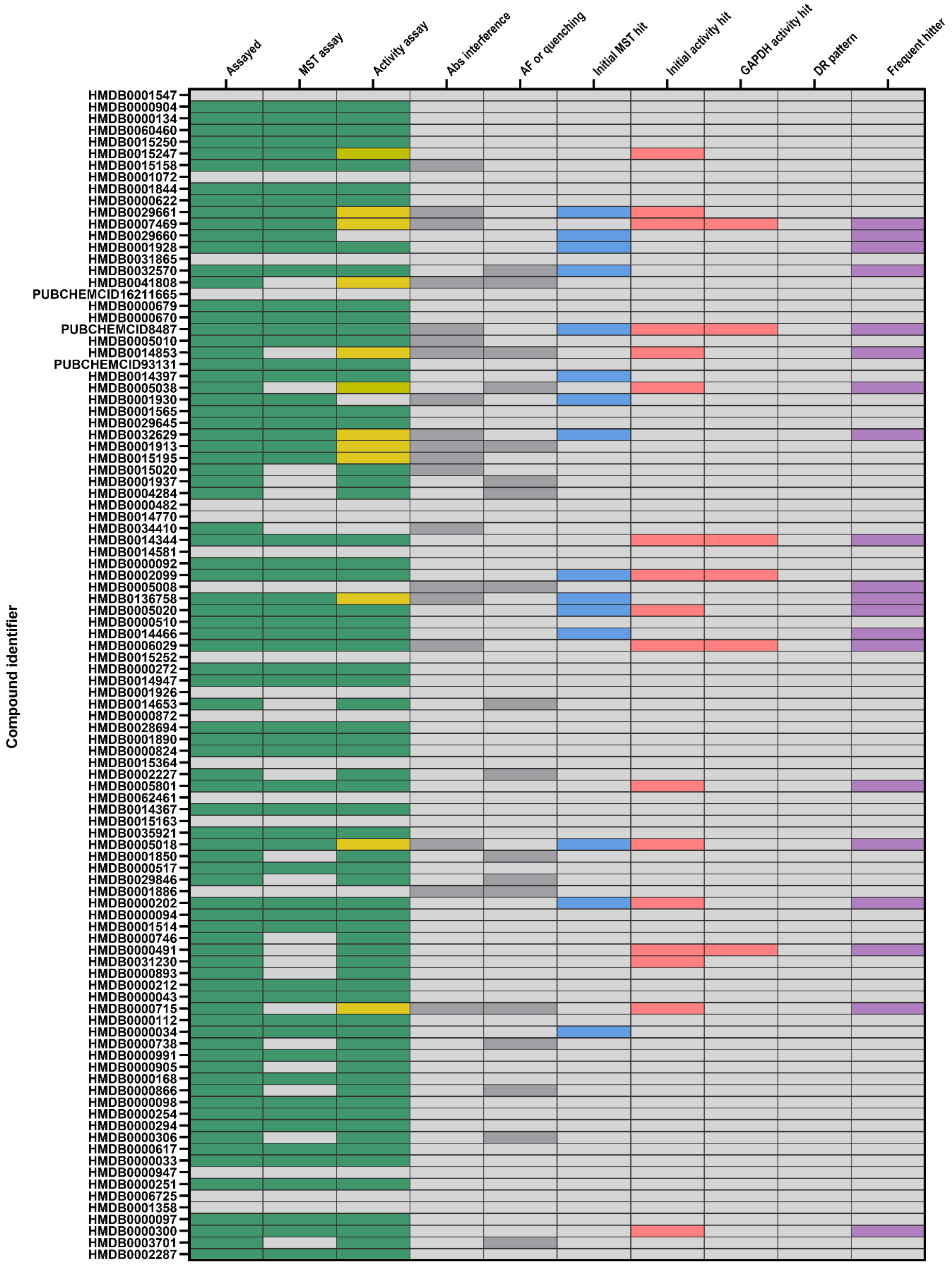

### TPI screening: MetaSci – Box 5

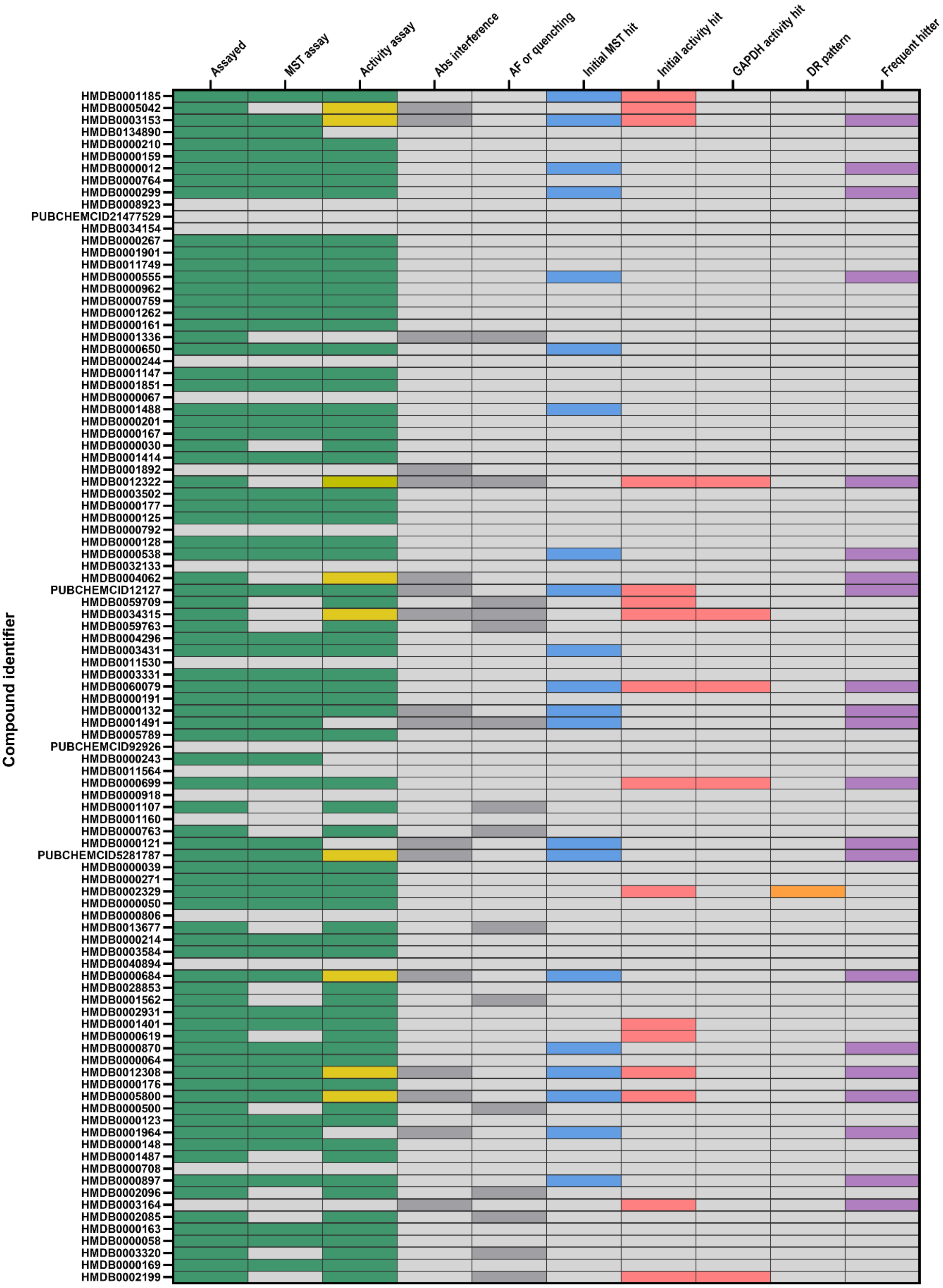

### TPI screening: MetaSci – Box 6

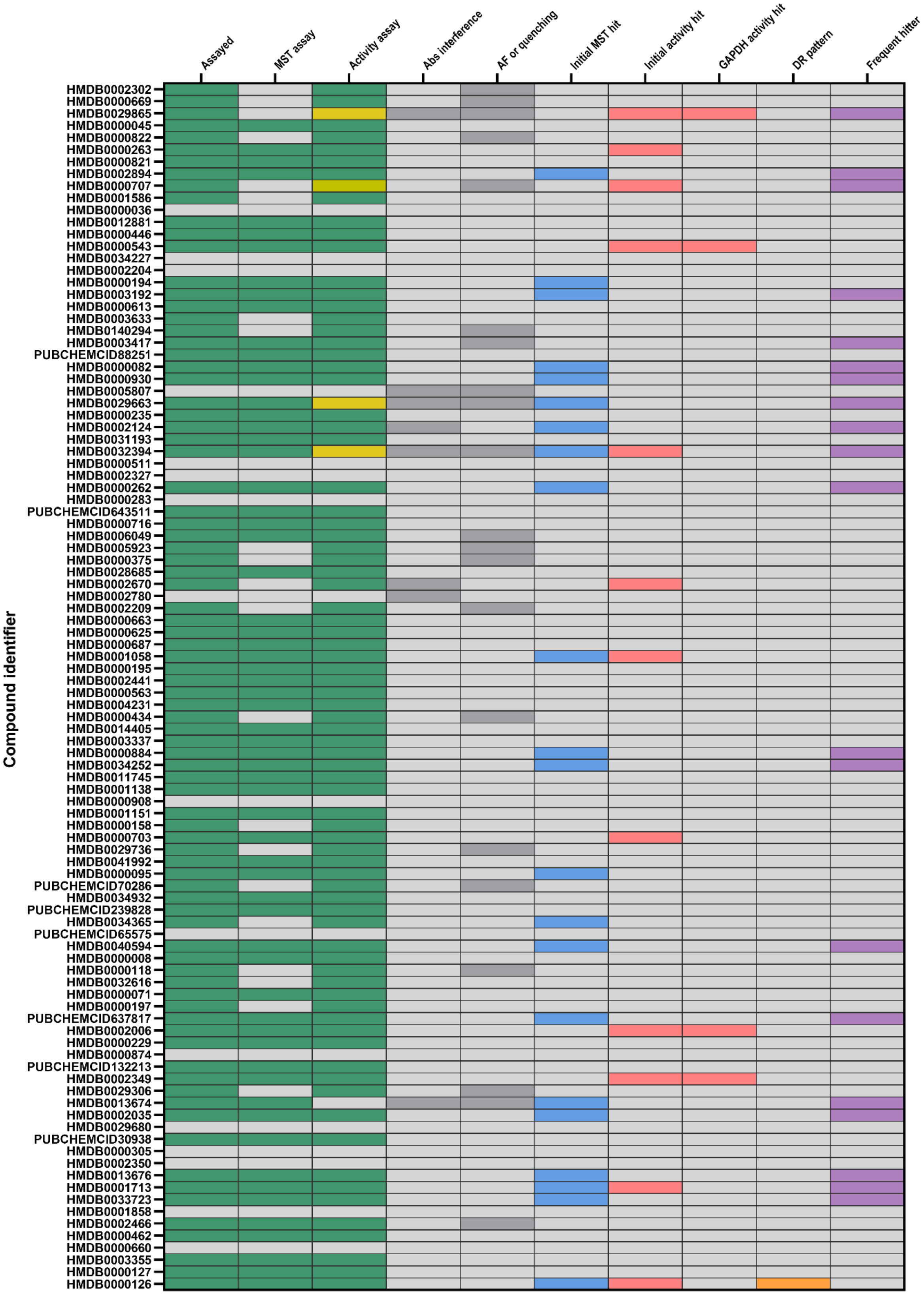

TPI screening: MetaSci – Box 7

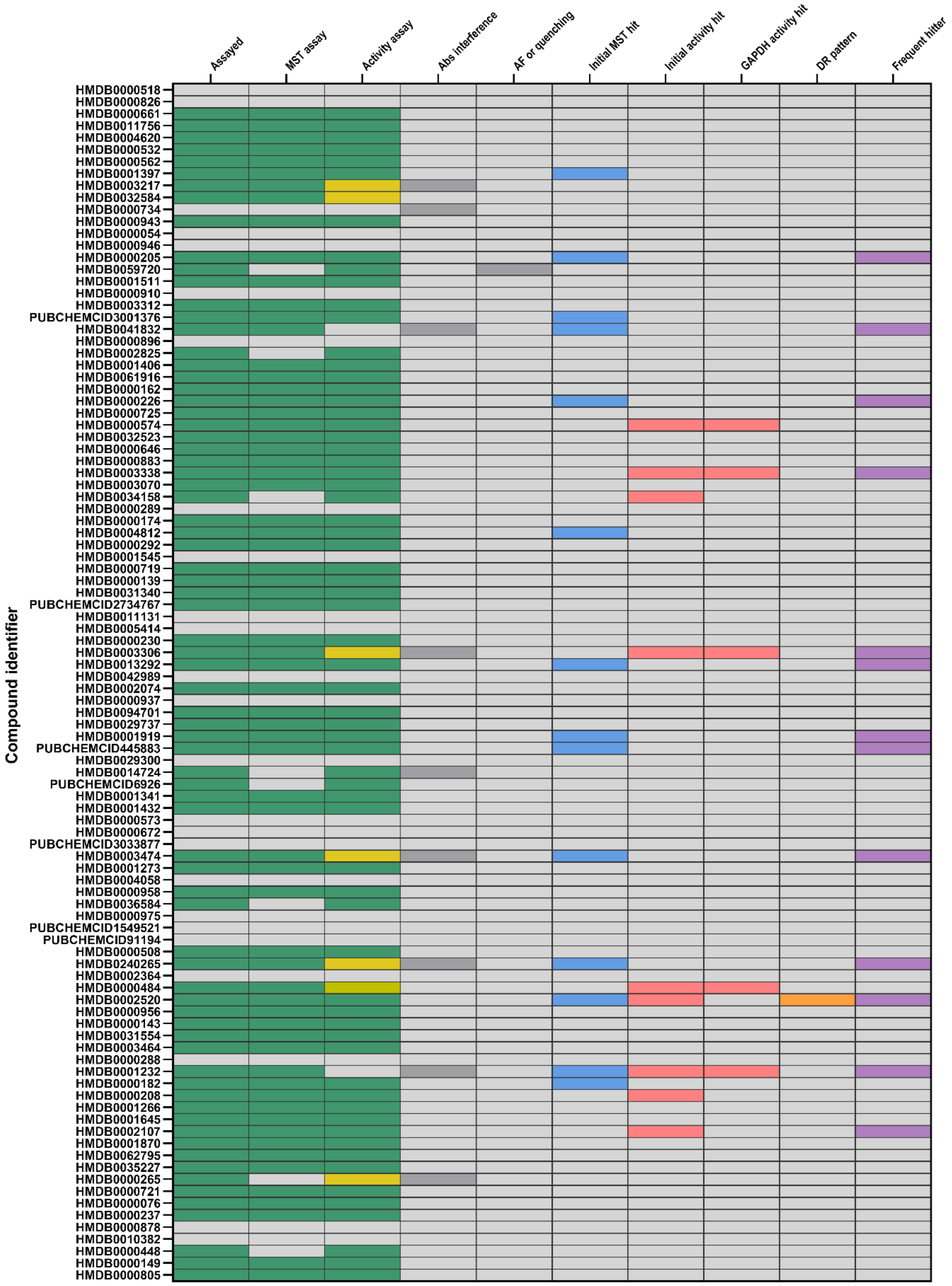

### TPI screening: MetaSci – Box 8

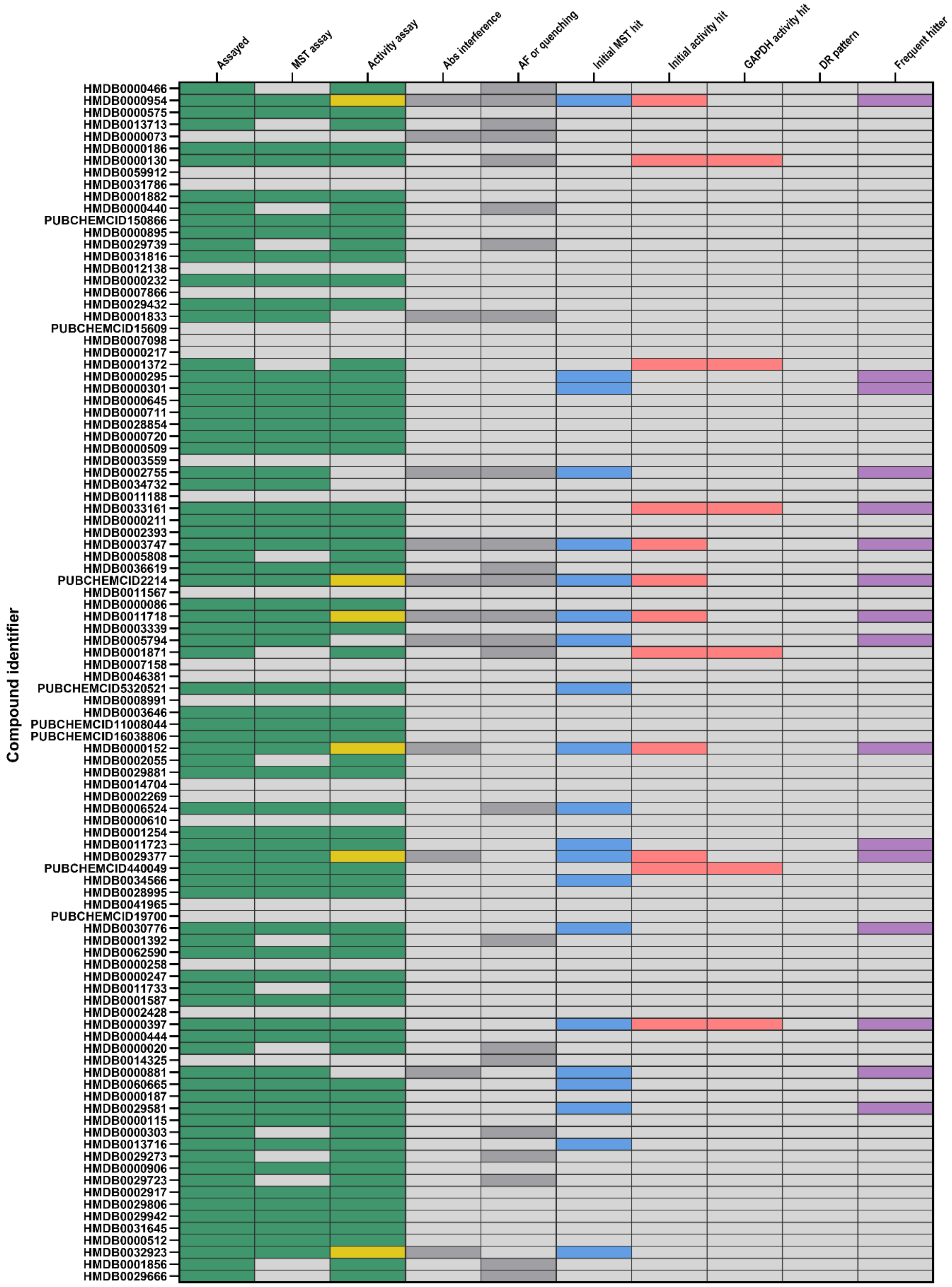

### TPI screening: MetaSci – Box 9

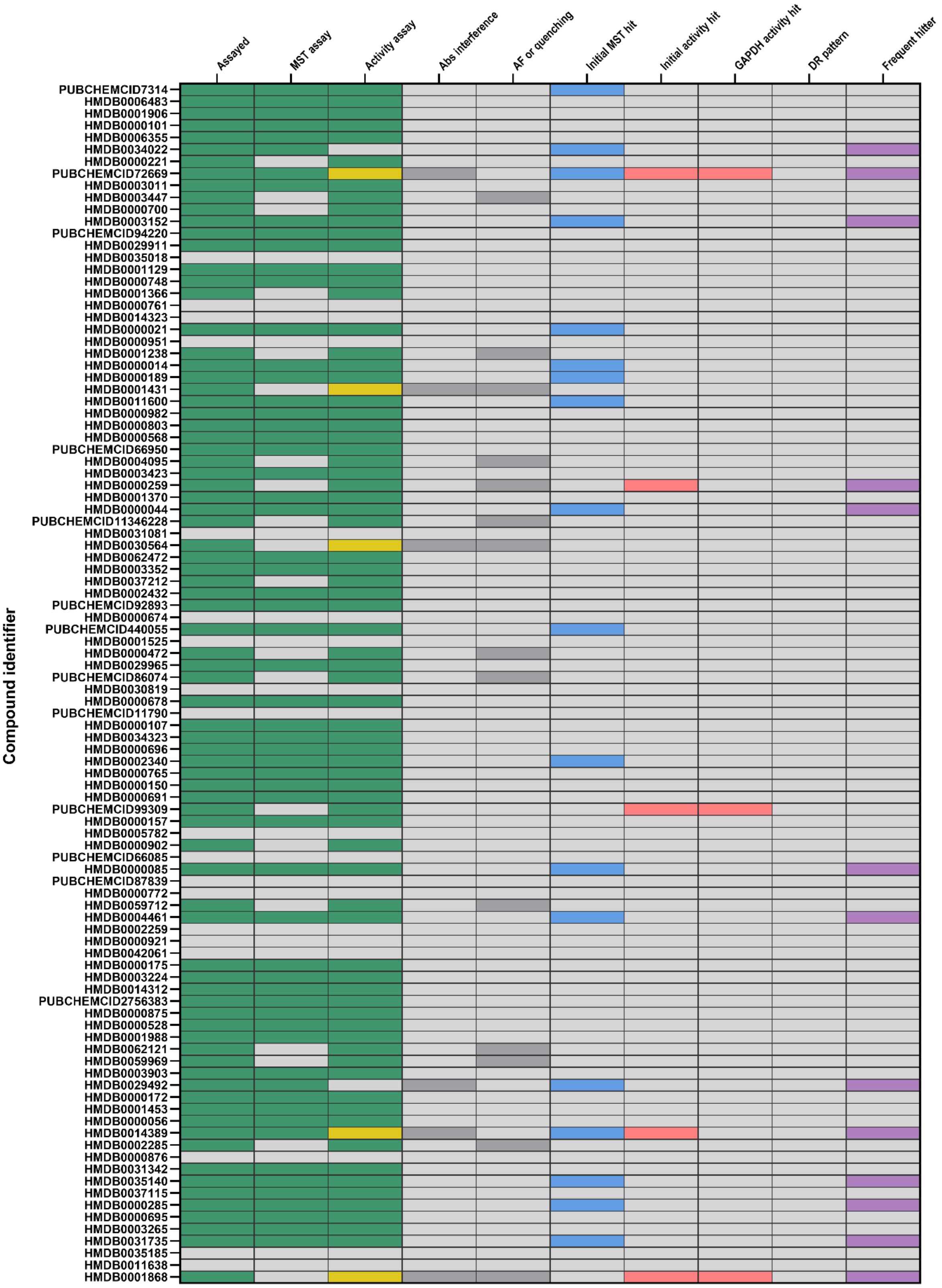

### TPI screening: MetaSci – Box 10

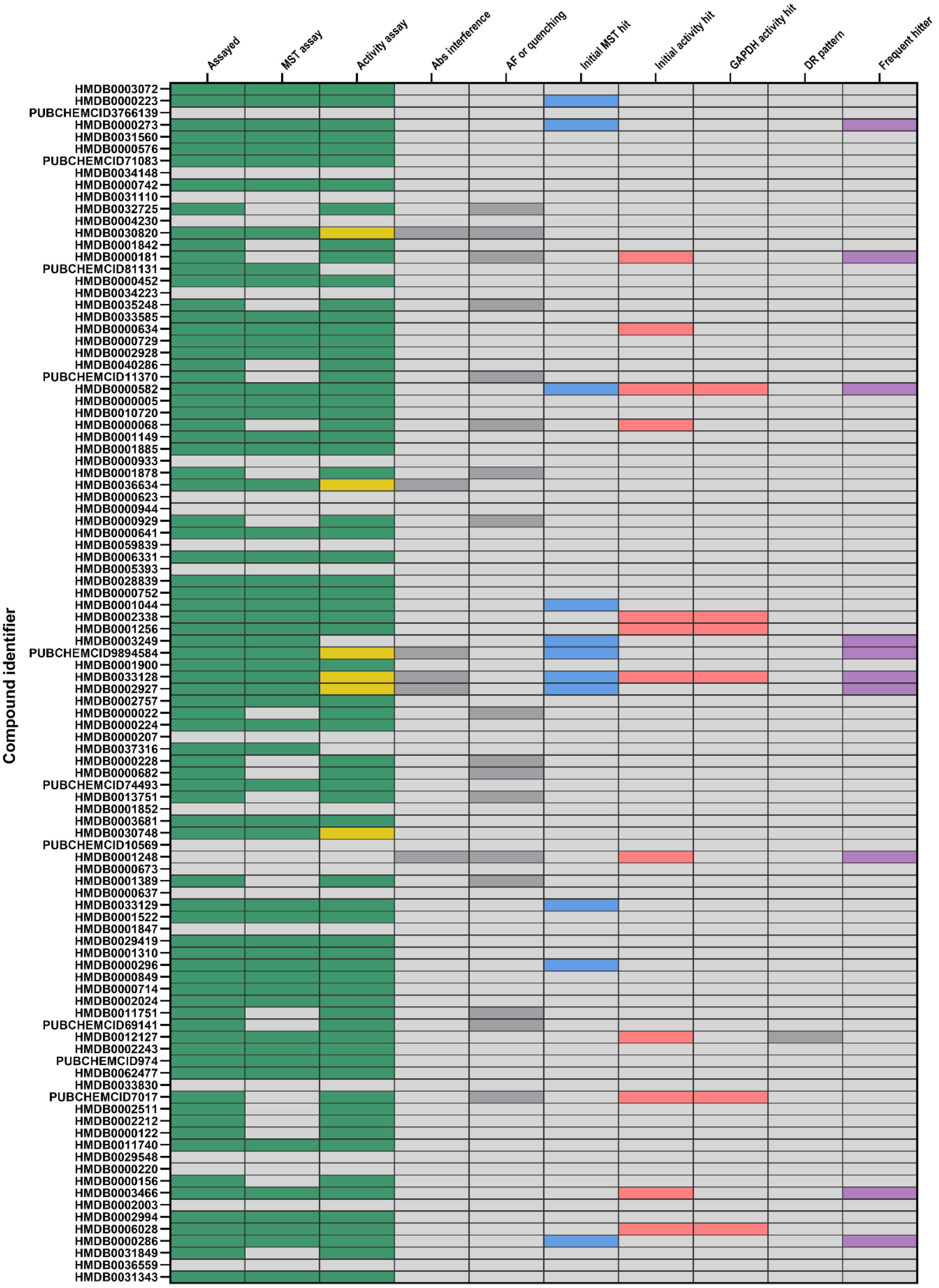

### TPI screening: MetaSci – Box 11

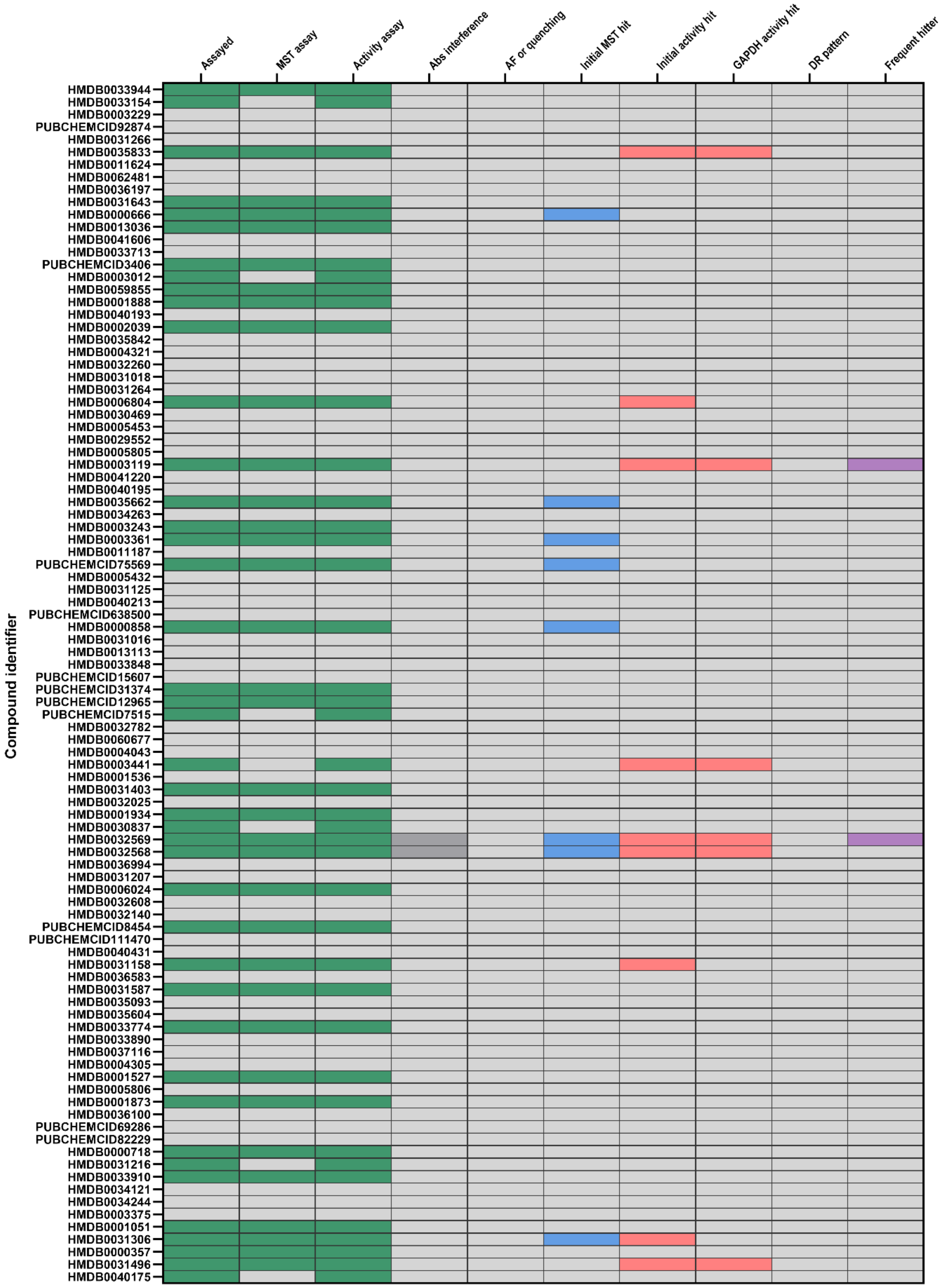

TPI screening: MetaSci – Box 12

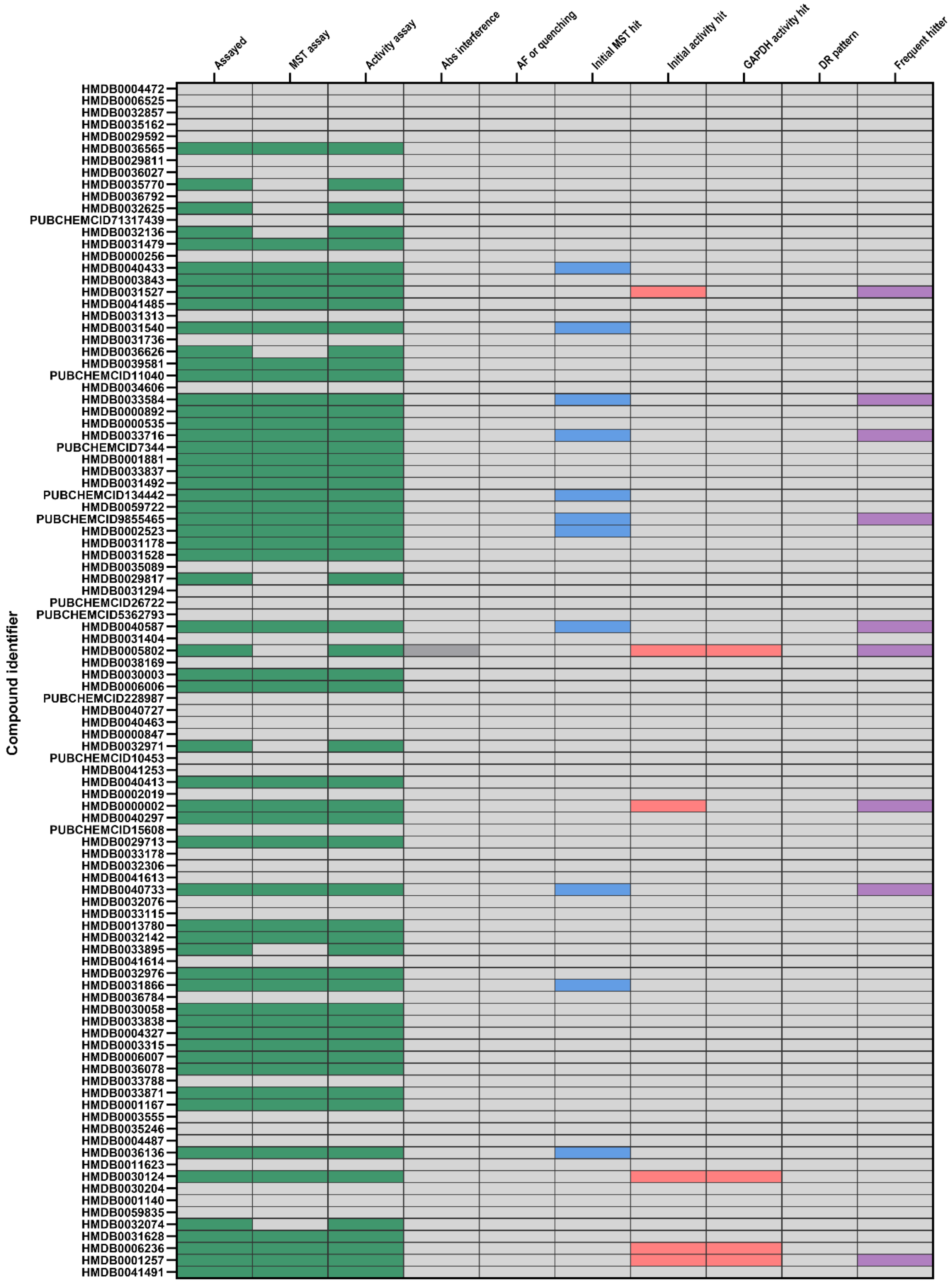

### TPI screening: MetaSci – Box 13 + Custom library

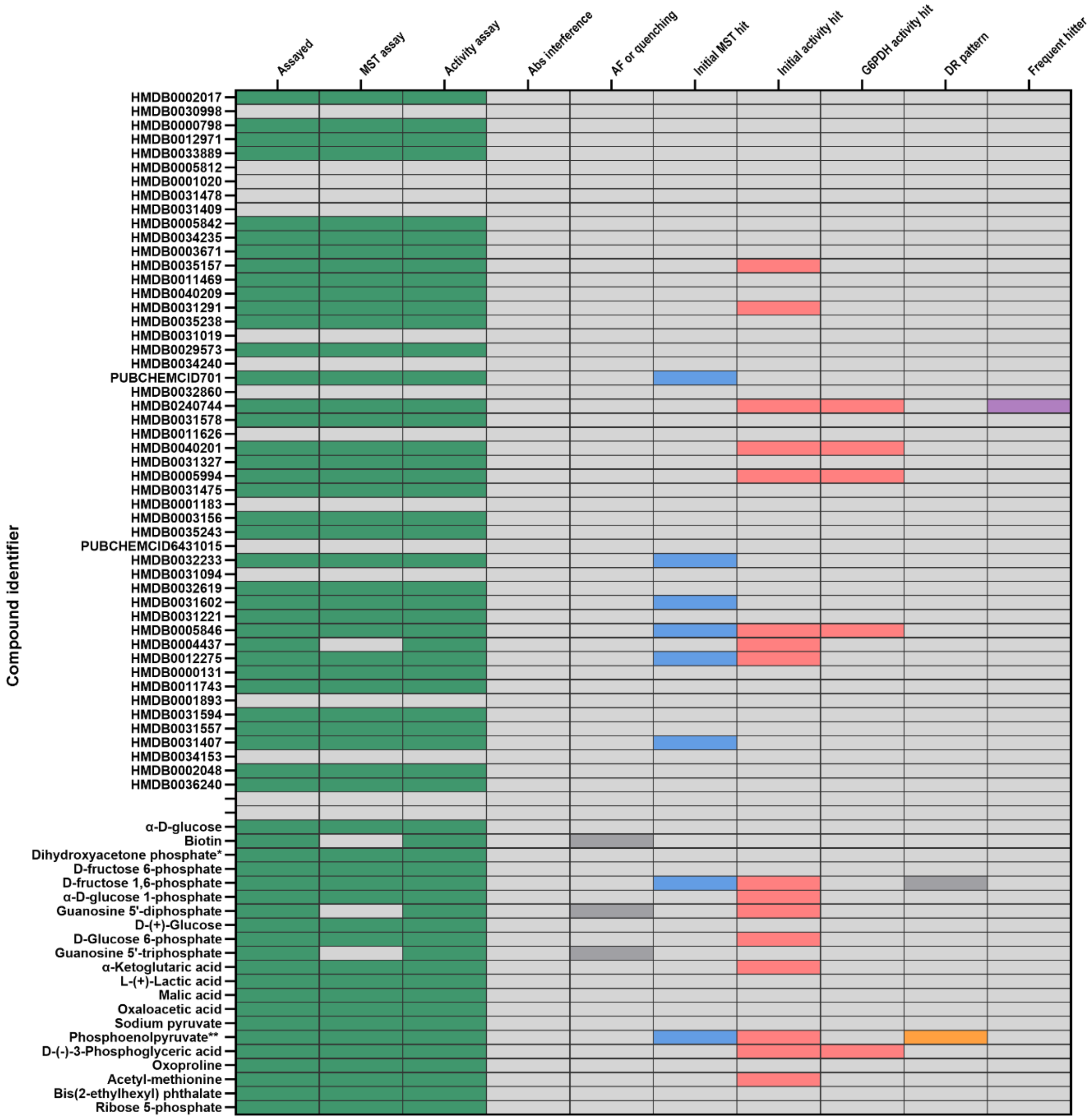


* Enzyme substrate

** Known inhibitor
