## Supplementary data C for "Human Glycolysis Isomerases are Inhibited by Weak Metabolite Modulators"

Known GPI inhibitors from literatures:

| Name | Structure | Species | PDB code | Protein-ligand interaction diagram |
|---|---|---|---|---|
| Erythrose-4-phospahte | 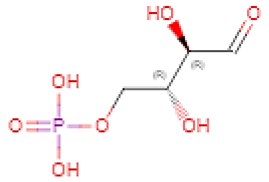 | Human | 1IRI | 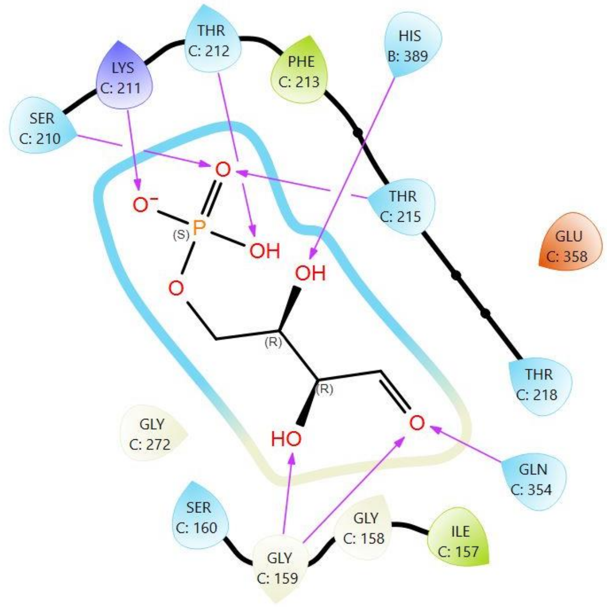 |

<table>
<tr>
<td>O1B<br>(2R,3R,4S)-5-((2-aminoethyl)amino)-2,3,4-trihydroxy-5-oxopentyl dihydrogen phosphate</td>
<td>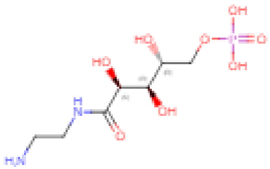</td>
<td>Human</td>
<td>6XUH</td>
<td>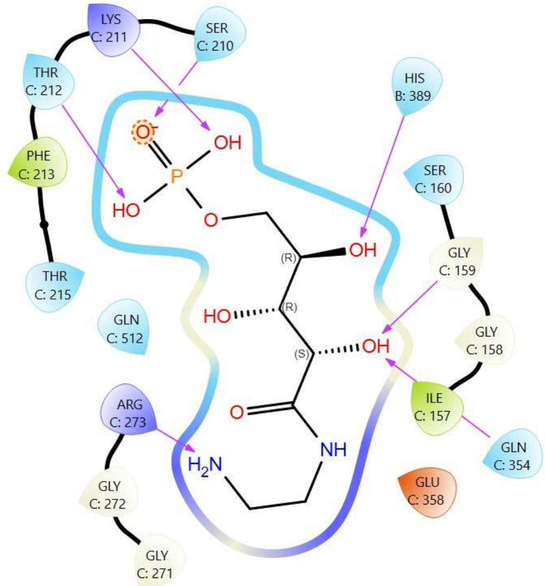
</td>
</tr>
<tr>
<td>PA5<br>5-Phosphoarabinoic acid</td>
<td>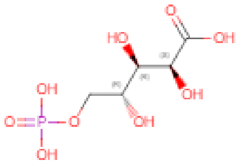</td>
<td>Human</td>
<td>6XUI</td>
<td>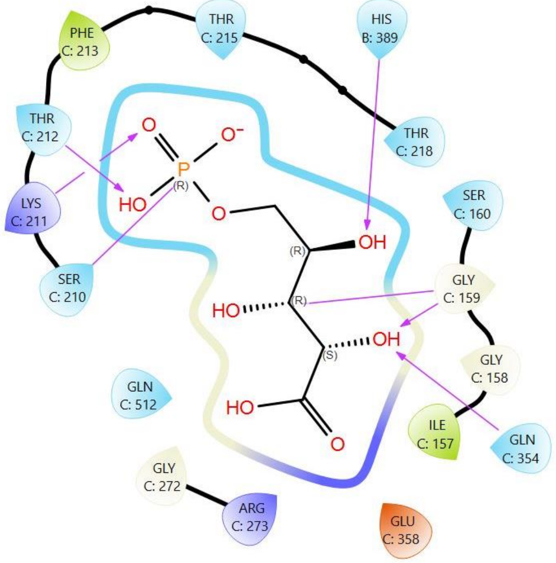
</td>
</tr>
</table>

| | | | | |
|---|---|---|---|---|
| F6P<br>Fructose-6-phosphate | 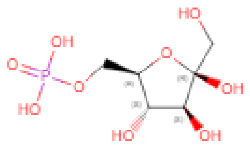 | Rabbit | 1HOX | 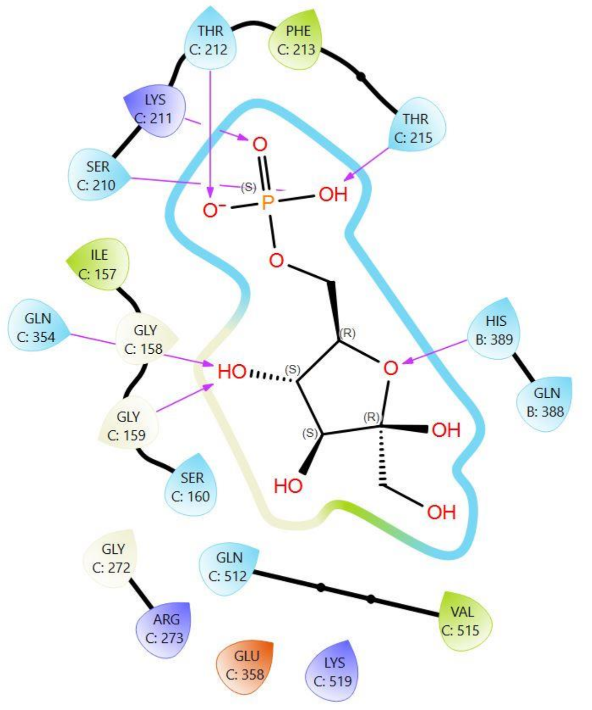 |

| | | | | |
|---|---|---|---|---|
| S6P<br>Sorbitol-6-phosphate | 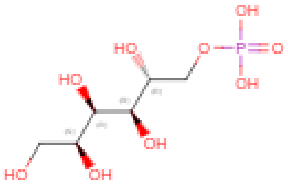 | Rabbit | 1XTB | 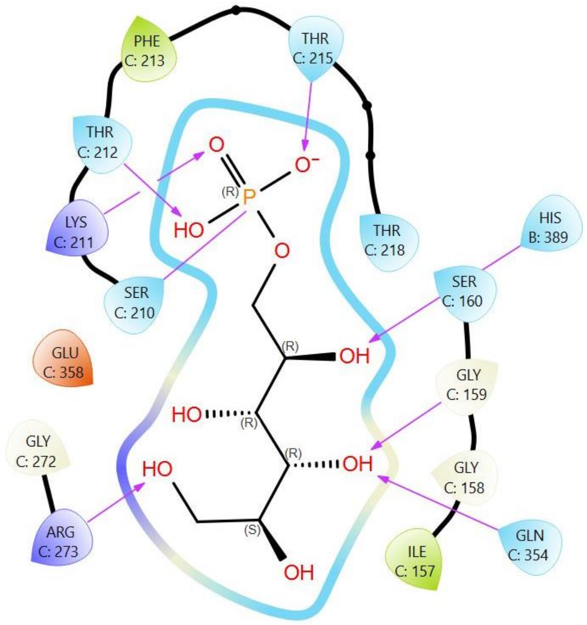 |

| | | | | |
|---|---|---|---|---|
| G6P<br>Glucose-6-phosphate | 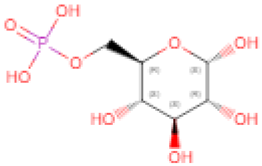 | Mouse | 1U0F | 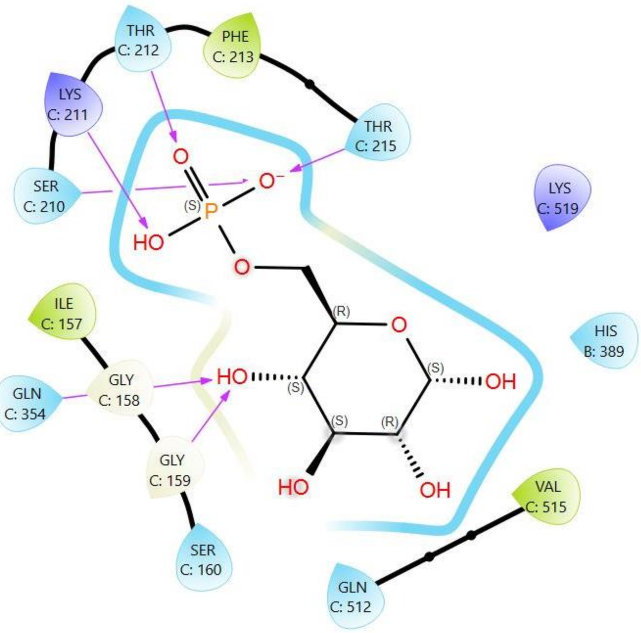 |

| | | | | |
|---|---|---|---|---|
| 6PGc<br>6-phosphogluconic acid | 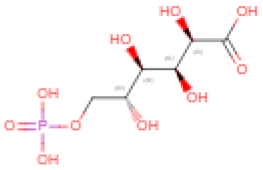 | Mouse | 2CXR | 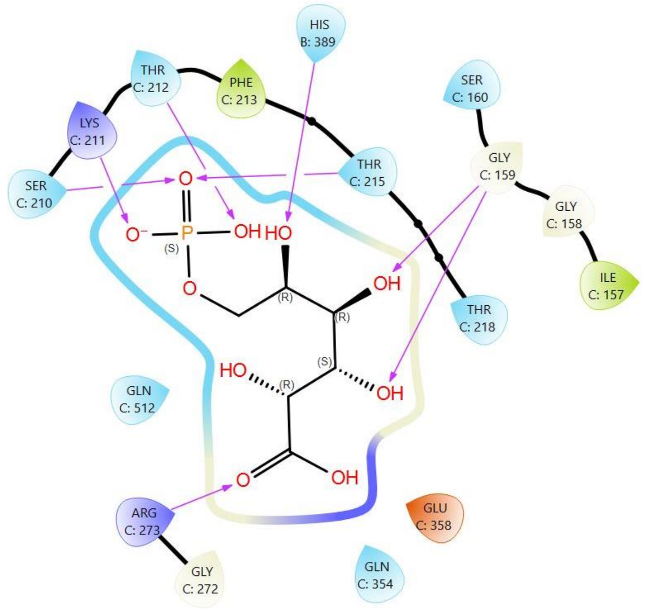  |

Known TPI inhibitors from literatures:

| Name | Structure | Species | PDB code | Protein-ligand interactions |
|---|---|---|---|---|

| | | | | |
|---|---|---|---|---|
| 2PG<br>2-Phosphoglycolic aicd | 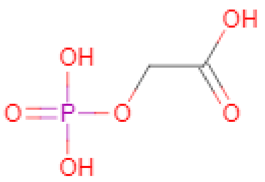 | Human | 1HTI | 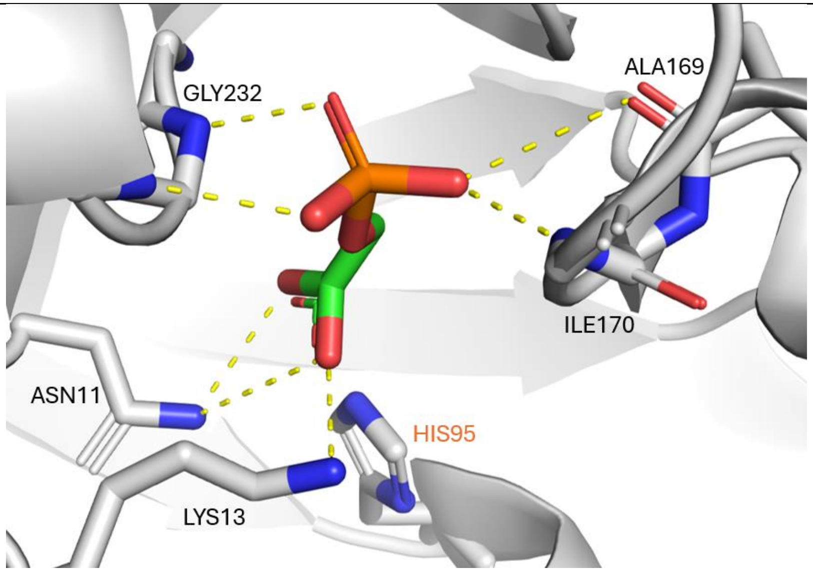 |
| PEP<br>Phosphoenol-pyruvate | 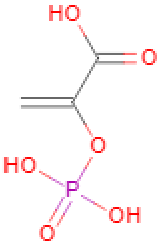 | Rabbit | 4OWG | 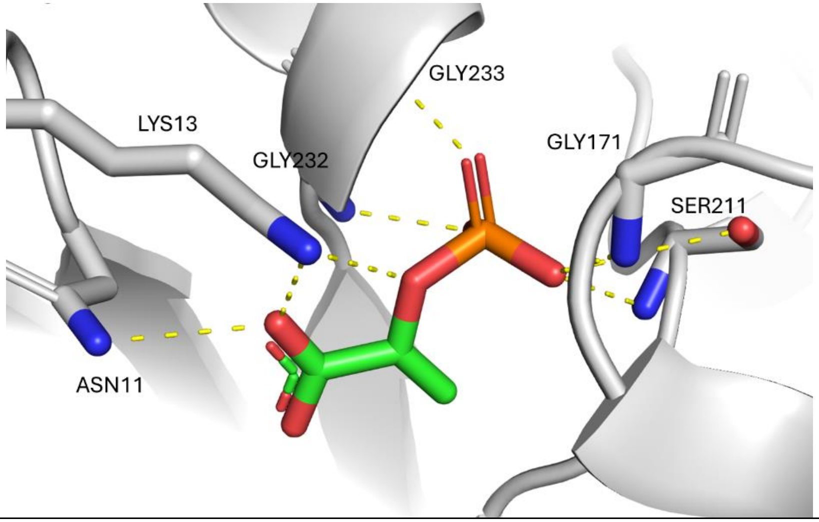 |

| | | | | |
|---|---|---|---|---|
| PGH<br>Phosphoglycolo-hydroxamic acid | 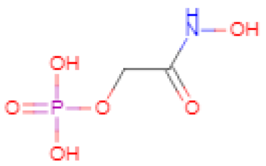 | Chicken | 1TPH | 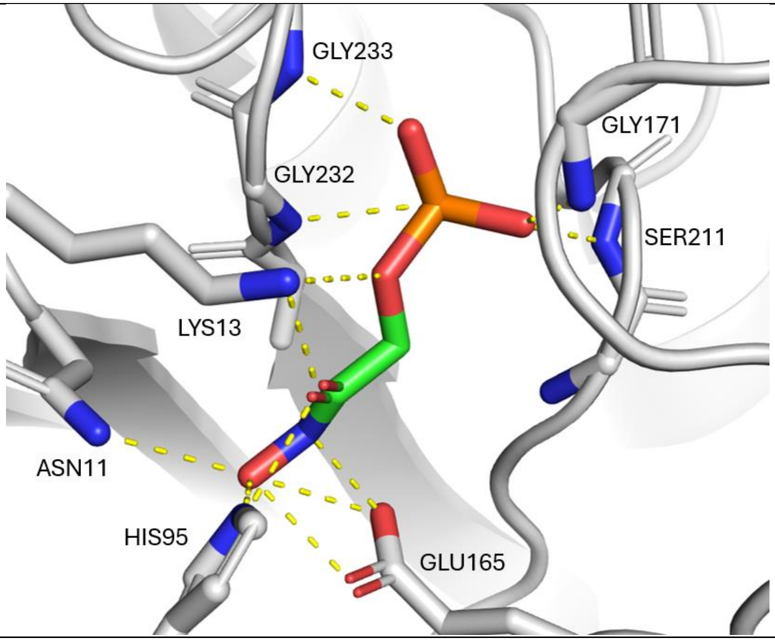<br> |

| | | | | |
|---|---|---|---|---|
| DHAP Dihydroxyacetone -phosphate | 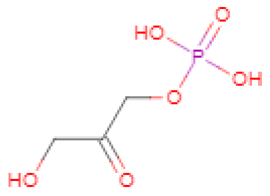 | Saccharomyces cerevisiae | 1NEY | 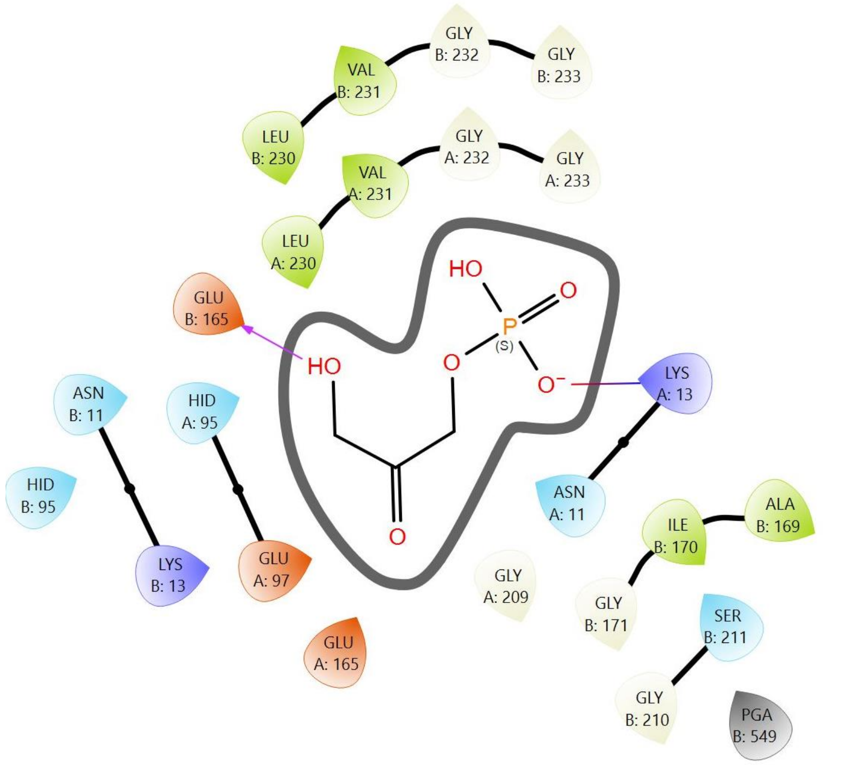 |

| | | | | |
|---|---|---|---|---|
| H4PB N-hydroxy-4-phosphono-butanamide | 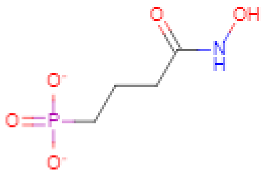 | Trypanosoma brucei | 1TSI | 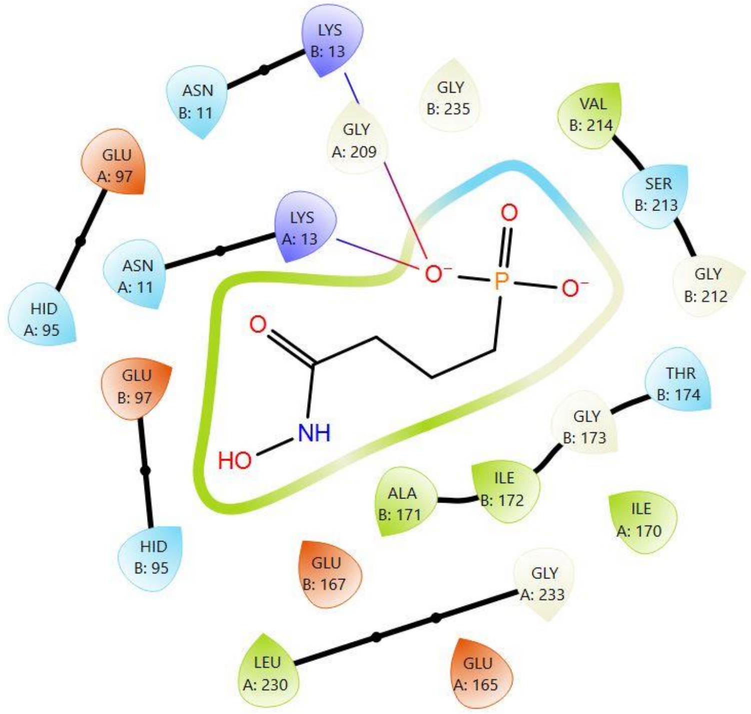 |

| | | | | |
|---|---|---|---|---|
| 2PG 2-phospholyceric acid | 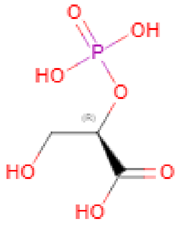  | Trypanosoma brucei | 4TIM | 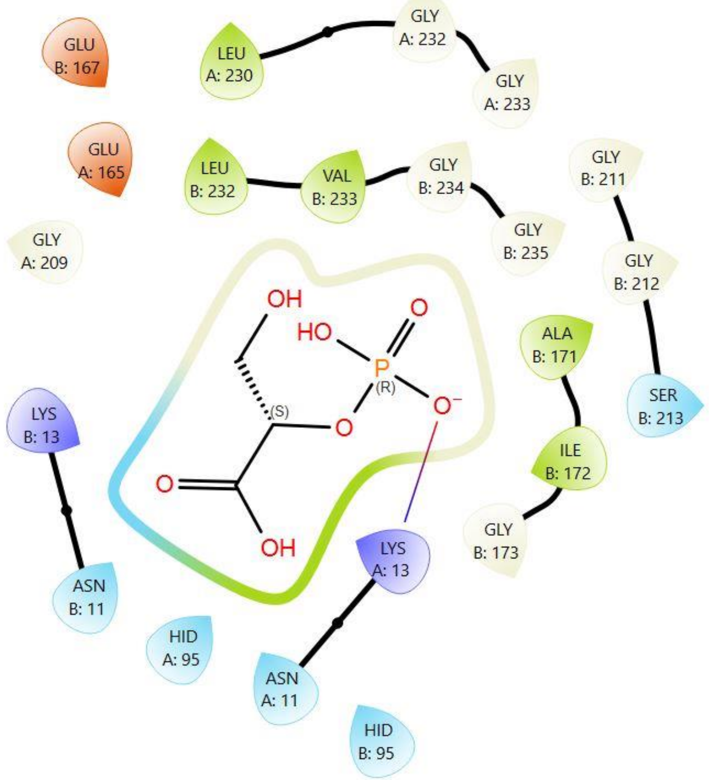  |

| | | | | |
|---|---|---|---|---|
| G3P Glycerol-3-phosphate | 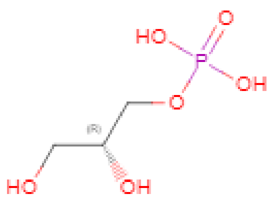  | Trypanosoma brucei | 6TIM | 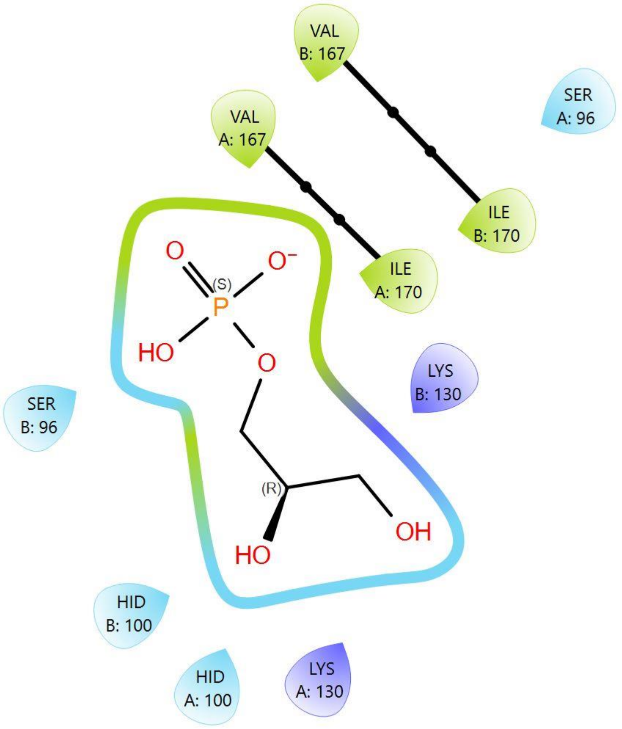  |

| | | | | |
|---|---|---|---|---|
| 3PG<br><br>3-phosphoglycerate | 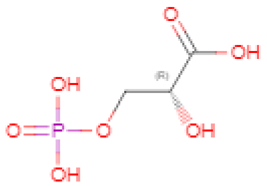 | Trypanosoma brucei | 1IIH | 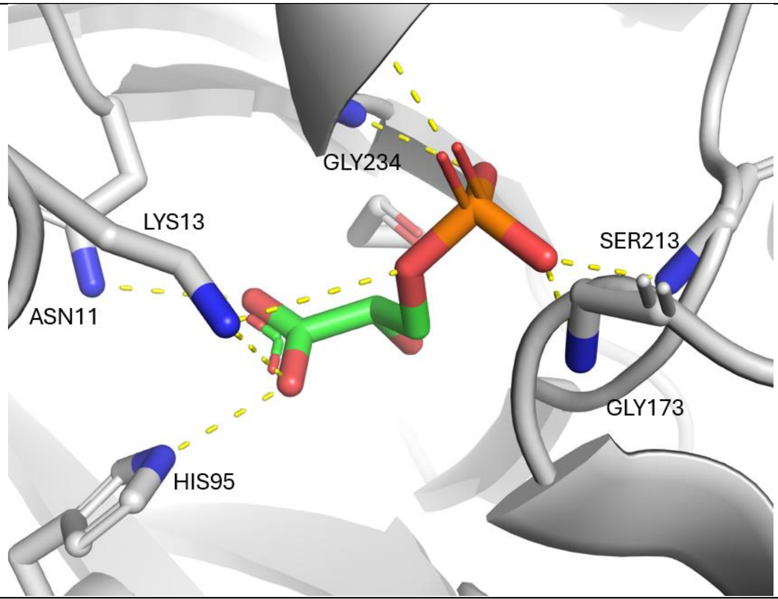 |
