## Appendix A - Ligand solubility guide for "Human Glycolysis Isomerases are Inhibited by Weak Metabolite Modulators"

| Column1 | identifier | name | box | row | column | mg/mL | %DMSO | Acid/Base | Comment | mass (mg) | Mw (g/mol) |
|---|---|---|---|---|---|---|---|---|---|---|---|
| 600 | HMDB0000407 | 2-Hydroxy-3-methylbutyric acic | 1 | A | 1 | **74** | **100** | | | 14.8 | 118.1 |
| 284 | HMDB0004815 | 4-Hydroxy-3-methylbenzoic aci | 1 | A | 2 | **100** | **94** | | | 11.3 | 152.2 |
| 555 | HMDB0029676 | Diosmetin | 1 | A | 3 | **175** | **90** | | | 9 | 300.3 |
| 906 | PUBCHEMCID14077841 | Cholesteryl Heptadecanoate | 1 | A | 4 | | | | | 6.3 | 639.1 |
| 329 | HMDB0002043 | 5-Phenylvaleric acid | 1 | A | 5 | **60** | **70** | | | 20 | 178.2 |
| 931 | HMDB0000766 | N-Acetyl-L-alanine | 1 | A | 6 | **162** | **80** | | | 16.2 | 131.1 |
| 1119 | HMDB0000827 | Stearic acid | 1 | A | 7 | **103** | **0** | | | 14.3 | 284.5 |
| 805 | HMDB0000638 | Dodecanoic acid | 1 | A | 8 | **87** | **90** | | | 17.4 | 200.3 |
| 715 | HMDB0000133 | Guanosine | 1 | A | 9 | **65** | **90** | | | 13.1 | 283.2 |
| 1015 | HMDB0014928 | Perindopril | 1 | A | 10 | | | | | 6.5 | 441.6 |
| 275 | HMDB0014378 | Aminosalicylic Acid | 1 | B | 1 | **100** | **65** | | | 13.7 | 153.1 |
| 235 | PUBCHEMCID117765 | N-(2-Acetamido)-2-Iminodiacet | 1 | B | 2 | **32** | **100** | **NaOH** | **Partially soluble** | 16 | 190.2 |
| 289 | HMDB0137904 | 4-hydroxy-2H-chromen-2-one | 1 | B | 3 | **240** | **100** | | | 12.7 | 162.1 |
| 1055 | HMDB0000239 | Pyridoxine | 1 | B | 4 | **119** | **0** | | | 23.8 | 205.6 |
| 1014 | HMDB0014944 | Pentoxifylline | 1 | B | 5 | | | | | 19.4 | 278.3 |
| 690 | PUBCHEMCID2915 | Cystamine dihydrochloride | 1 | B | 6 | **280** | **0** | | | 28.3 | 225.2 |
| 952 | HMDB0000828 | Ureidosuccinic acid | 1 | B | 7 | **42** | **90** | | | 17 | 176.1 |
| 474 | PUBCHEMCID22324 | Hexadecylpyridinium chloride | 1 | B | 8 | | | | | 12.3 | 340.0 |
| 812 | HMDB0000192 | L-Cystine | 1 | B | 9 | **27** | **0** | **0.5 M HCl** | | 13.5 | 240.3 |
| 413 | HMDB0030094 | Betulinic acid | 1 | B | 10 | **35** | **100** | | **Partially soluble** | 14.9 | 456.7 |
| 614 | PUBCHEMCID12332 | Hexanamide | 1 | C | 1 | **172** | **100** | | | 17.2 | 115.2 |
| 393 | HMDB0014712 | Atropine | 1 | C | 2 | **39.3** | **100** | | | 17.7 | 694.8 |
| 814 | HMDB0001918 | Thyroxine | 1 | C | 3 | **102.8** | **60** | | | 25.7 | 776.9 |
| 802 | HMDB0029416 | L-Targinine | 1 | C | 4 | **100** | **0** | | | 12.7 | 261.2 |
| 647 | HMDB0015279 | Flavoxate | 1 | C | 5 | | | | | 11.6 | 427.9 |
| 138 | PUBCHEMCID67215 | Leucomalachite green | 1 | C | 6 | **33.2** | **100** | | | 12.6 | 330.5 |
| 576 | PUBCHEMCID3935589 | Acid Yellow 36 | 1 | C | 7 | **44.6** | **0** | | | 22.3 | 375.4 |
| 605 | PUBCHEMCID164825 | Acid Yellow 23 | 1 | C | 8 | **96** | **0** | | | 19.2 | 534.4 |
| 366 | HMDB0015636 | Agomelatine | 1 | C | 9 | **116** | **100** | | | 23.2 | 243.3 |
| 346 | HMDB0001859 | Acetaminophen | 1 | C | 10 | **295** | **100** | | | 29.5 | 151.2 |
| 569 | HMDB0000193 | Isocitric acid | 1 | D | 1 | **100** | **0** | | | 10.3 | 258.1 |
| 1239 | HMDB0000561 | beta-Carotene | 1 | D | 2 | **100** | **100** | | | 11.4 | 536.9 |
| 1072 | HMDB0003648 | Retinyl palmitate | 1 | D | 3 | | | | | 23.2 | 524.9 |
| 492 | HMDB0014997 | Penicillamine | 1 | D | 4 | **68.5** | **0** | | | 13.7 | 149.2 |
| 1050 | PUBCHEMCID6578 | Propionamide | 1 | D | 5 | **60.6** | **0** | | | 12.1 | 73.1 |
| 917 | PUBCHEMCID10930 | 3-Methylbutyramide | 1 | D | 6 | **100** | **100** | | | 19.8 | 101.2 |
| 422 | HMDB0033870 | Butyramide | 1 | D | 7 | **59** | **0** | | | 11.8 | 87.1 |
| 352 | HMDB0014925 | Aciclovir | 1 | D | 8 | **46** | **80** | | | 23.3 | 225.2 |
| 1082 | HMDB0042010 | Rutaecarpine | 1 | D | 9 | **18.5** | **90** | | | 14.6 | 287.3 |
| 861 | PUBCHEMCID442088 | (+)-Evodiamine | 1 | D | 10 | **21** | **100** | | | 5.7 | 303.4 |
| 918 | PUBCHEMCID12298 | Valeramide | 1 | E | 1 | **53.5** | **0** | | | 10.7 | 101.2 |
| 527 | HMDB0000124 | Fructose 6-phosphate | 1 | E | 2 | **211** | **0** | | | 21.1 | 305.0 |
| 697 | HMDB0000631 | Deoxycholic acid glycine conjug | 1 | E | 3 | **89** | **100** | | | 8.9 | 467.6 |
| 757 | HMDB0005785 | Indole-3-carbinol | 1 | E | 4 | **100** | **100** | | | 11.4 | 147.2 |
| 535 | PUBCHEMCID91746269 | Ac-L-Orn-OH | 1 | E | 5 | **63** | **0** | | | 6.3 | 174.2 |
| 1062 | PUBCHEMCID101 | 3-Hydroxybenzaldehyde | 1 | E | 6 | **231** | **100** | | | 23.1 | 122.1 |
| 747 | HMDB0000733 | Hyodeoxycholic acid | 1 | E | 7 | | | | | 13 | 392.6 |
| 336 | HMDB0000032 | 7-Dehydrocholesterol | 1 | E | 8 | **165** | **0** | | | 32.9 | 384.6 |
| 341 | PUBCHEMCID1048 | Pyrazole | 1 | E | 9 | **98.5** | **0** | | | 19.7 | 68.1 |
| 1147 | HMDB0001889 | Theophylline | 1 | E | 10 | **17.4** | **70** | **1.7** | | 12.2 | 180.2 |
| 414 | HMDB0011727 | Bicine | 1 | F | 1 | **165** | **0** | | | 33 | 163.2 |
| 262 | HMDB0001904 | 3-Nitrotyrosine | 1 | F | 2 | **53.2** | **90** | **1** | | 13.3 | 226.2 |
| 695 | HMDB0000138 | Glycocholic acid | 1 | F | 3 | **146** | **0** | | **100% MeOH** | 14.6 | 465.6 |
| 131 | HMDB0033826 | 2,6-Di-tert-butyl-4-methylphen | 1 | F | 4 | **95** | **90** | | | 16.2 | 220.3 |
| 1042 | HMDB0001830 | Progesterone | 1 | F | 5 | **130** | **0** | | **100% MeOH** | 13 | 314.5 |
| 409 | HMDB0003409 | Berberine | 1 | F | 6 | | | | | 11.3 | 371.8 |
| 632 | HMDB0034156 | Ethyl stearate | 1 | F | 7 | | | | | 13.9 | 312.5 |
| 959 | HMDB0037834 | Ethyl menthane carboxamide | 1 | F | 8 | **210** | **90** | | | 36.1 | 211.3 |
| 11 | HMDB0059838 | Camphor | 1 | F | 9 | | | | | 11.2 | 152.2 |
| 67 | HMDB0041971 | P-Dichlorobenzene | 1 | F | 10 | **70** | **90** | | | 18.2 | 147.0 |
| 608 | HMDB0000151 | Estradiol | 1 | G | 1 | | | | | 13.3 | 272.4 |
| 552 | HMDB0004983 | Dimethyl sulfone | 1 | G | 2 | **100** | **0** | **0.5** | | 13 | 94.1 |
| 1131 | HMDB0004826 | p-Synephrine | 1 | G | 3 | **17** | **0** | | | 12.7 | 167.2 |
| 1061 | HMDB0000842 | Quinaldic acid | 1 | G | 4 | **70** | **75** | | | 14.2 | 173.2 |
| 1207 | HMDB0001877 | Valproic acid | 1 | G | 5 | **110** | **0** | | | 11 | 166.2 |
| 789 | HMDB0003966 | Selenomethionine | 1 | G | 6 | **57** | **0** | | | 11.4 | 196.1 |
| 1091 | HMDB0036827 | Sclareol | 1 | G | 7 | | | | | 21.7 | 308.5 |
| 386 | HMDB0030353 | Arecoline | 1 | G | 8 | **100** | **70** | | | 11.4 | 236.1 |
| 961 | HMDB0001015 | N-Formyl-L-methionine | 1 | G | 9 | **61** | **50** | | | 6.1 | 177.2 |
| 24 | HMDB0035819 | (-)-Isoborneol | 1 | G | 10 | **118** | **80** | | | 11.8 | 154.3 |
| 657 | HMDB0001933 | Furosemide | 1 | H | 1 | **150** | **60** | **NaOH** | | 20.3 | 330.7 |
| 442 | HMDB0062455 | cholest-5-en-3beta-yl (11Z,14Z- | 1 | H | 2 | | | | | 5.5 | 677.1 |
| 1033 | HMDB0015217 | Pilocarpine | 1 | H | 3 | **146** | **50** | | | 23.4 | 244.7 |
| 1067 | HMDB0014351 | Reserpine | 1 | H | 4 | | | | | 30.2 | 608.7 |
| 1232 | HMDB0001388 | alpha-Linolenic acid | 1 | H | 5 | **53** | **100** | | | 5.3 | 300.4 |
| 636 | HMDB0015109 | Edetic Acid | 1 | H | 6 | **45** | **0** | | | 17.4 | 292.2 |
| 557 | HMDB0001927 | Diphenhydramine | 1 | H | 7 | **100** | **40** | | | 12.1 | 291.8 |
| 878 | HMDB0030017 | Menatetrenone | 1 | H | 8 | | | | | 5.3 | 444.7 |
| 585 | HMDB0001936 | Doxylamine | 1 | H | 9 | **100** | **30** | | | 13.6 | 388.5 |
| 455 | HMDB0030282 | Cinchonidine | 1 | H | 10 | **82** | **85** | **0.25** | | 16.5 | 294.4 |

| | | | | | | | | | | | |
|---|---|---|---|---|---|---|---|---|---|---|---|
| 1035 | HMDB0041481 | 4-(3,4-Methylenedioxyphenyl)- | 1 | I | 1 | **64** | **74** | | | 14.9 | 192.2 |
| 523 | HMDB0000626 | Deoxycholic acid | 1 | I | 2 | **123** | **0** | | **100% MeOH** | 12.3 | 392.6 |
| 884 | PUBCHEMCID2723 | 4-Chloro-3,5-dimethylphenol | 1 | I | 3 | **94** | **56** | | | 15.1 | 156.6 |
| 1186 | HMDB0000925 | Trimethylamine N-oxide | 1 | I | 4 | **100** | **0** | | | 19.9 | 75.1 |
| 228 | HMDB0059966 | 3,5-Dimethoxyphenol | 1 | I | 5 | **134.5** | **30** | | | 26.9 | 154.2 |
| 81 | HMDB0003424 | 1-Hexadecanol | 1 | I | 6 | | | | | 15.2 | 242.4 |
| 1 | HMDB0001046 | Cotinine | 1 | I | 7 | **69** | **50** | | | 6.9 | 176.2 |
| 1135 | PUBCHEMCID15684 | 2,4-dichlorobenzyl alcohol | 1 | I | 8 | **20** | **95** | | **Partially soluble** | 13.2 | 177.0 |
| 496 | HMDB0000055 | Cellobiose | 1 | I | 9 | **45.3** | **16.7** | | | 13.6 | 342.3 |
| 218 | HMDB0059965 | 3,4-Dihydroxybenzaldehyde | 1 | I | 10 | **100** | **38.4** | | | 12.9 | 138.1 |
| 484 | HMDB0000089 | Cytidine | 1 | J | 1 | **56** | **40** | | | 14 | 243.2 |
| 602 | HMDB0002899 | Ellagic acid | 1 | J | 2 | **157** | **50** | | **Partially soluble** | 15.7 | 338.2 |
| 470 | HMDB0001218 | Coumarin | 1 | J | 3 | | | | | 11.9 | 146.1 |
| 711 | HMDB0036648 | 7-Isopropyl-1,4-dimethylazulen | 1 | J | 4 | | | | | 17.6 | 198.3 |
| 65 | HMDB0002144 | 1,3-Dimethyluracil | 1 | J | 5 | **78** | **66** | | | 11.7 | 140.1 |
| 433 | HMDB0000957 | Pyrocatechol | 1 | J | 6 | **100** | **0** | | | 12.2 | 110.1 |
| 610 | HMDB0000145 | Estrone | 1 | J | 7 | | | | | 10.6 | 270.4 |
| 398 | HMDB0011143 | MG(O-18:0/0:0/0:0) | 1 | J | 8 | | | | | 15.3 | 344.6 |
| 190 | HMDB0000329 | 2-Phenylbutyric acid | 1 | J | 9 | **122** | **58** | | | 18.3 | 164.2 |
| 977 | HMDB0032574 | Propylparaben | 1 | J | 10 | **61** | **60** | | | 15.4 | 180.2 |
| 584 | HMDB0014399 | Doxycycline | 2 | A | 1 | **35** | **78** | | | 13.3 | 462.5 |
| 412 | HMDB0036838 | Betulin | 2 | A | 2 | **6.44** | **100** | | | 11.6 | 442.7 |
| 1039 | HMDB0031188 | Ethanethioic acid | 2 | A | 3 | **56** | **33** | | | 17 | 114.2 |
| 1046 | HMDB0033835 | Propyl gallate | 2 | A | 4 | **68.5** | **50** | | | 13.7 | 212.2 |
| 430 | HMDB0029415 | S-Carboxymethyl-L-cysteine | 2 | A | 5 | **27.6** | **60** | | | 11.5 | 179.2 |
| 63 | HMDB0059963 | 1,3,5-Trimethoxybenzene | 2 | A | 6 | **68** | **70** | | | 13.7 | 168.2 |
| 1159 | HMDB0014447 | Tranexamic Acid | 2 | A | 7 | **100** | **0** | | | 17 | 157.2 |
| 598 | PUBCHEMCID71098 | D-Tyrosine | 2 | A | 8 | **68** | **50** | | | 14.7 | 181.2 |
| 525 | HMDB0001944 | Chlorpheniramine | 2 | A | 9 | **81.3** | **33** | | | 12.2 | 390.9 |
| 951 | HMDB0014869 | Nateglinide | 2 | A | 10 | **52** | **77** | | | 11.5 | 317.4 |
| 111 | HMDB0029657 | 2,4'-Dihydroxyacetophenone | 2 | B | 1 | **100** | **38** | | | 13.3 | 152.2 |
| 274 | HMDB0001169 | 4-Aminophenol | 2 | B | 2 | **68** | **78** | | | 12.4 | 109.1 |
| 388 | HMDB0015643 | Artemether | 2 | B | 3 | | | | | 16.3 | 298.4 |
| 354 | PUBCHEMCID3423265 | Sodium lauryl sulfate | 2 | B | 4 | **134** | **0** | | | 13.4 | 288.4 |
| 390 | HMDB0240267 | Artesunate | 2 | B | 5 | | | | | 14.7 | 384.4 |
| 330 | HMDB0000253 | Pregnenolone | 2 | B | 6 | **124** | **50** | | **50% MeOH** | 12.4 | 316.5 |
| 1134 | HMDB0014813 | Tamoxifen | 2 | B | 7 | | | | | 11.1 | 563.6 |
| 237 | PUBCHEMCID0099999 | 1,5,5-Trimethylhydantoin | 2 | B | 8 | **100** | **50** | | | 10.2 | 142.2 |
| 389 | HMDB0061088 | Dihydroartemisinin (DHA) | 2 | B | 9 | | | | **Poor water/DMSO solubil** | 11.7 | 282.3 |
| 513 | HMDB0003416 | D-Arginine | 2 | B | 10 | **26.2** | **10** | **0.5 M HCl** | | 13.1 | 174.2 |
| 514 | HMDB0033780 | D-Asparagine | 2 | C | 1 | **100** | **0** | **0.5 M HCl** | | 12.2 | 132.1 |
| 1096 | HMDB0030583 | Silibinin | 2 | C | 2 | | | | **Poor water solubility** | 15.8 | 482.4 |
| 1215 | HMDB0000607 | Cyanocobalamin | 2 | C | 3 | | | | **Poor water solubility** | 14 | 1355.4 |
| 784 | HMDB0000216 | Norepinephrine | 2 | C | 4 | **70** | **0** | | | 7 | 337.3 |
| 958 | HMDB0002368 | Nervonic acid | 2 | C | 5 | | | | **Poor water solubility** | 5.1 | 366.6 |
| 686 | HMDB0005356 | TG(16:0/16:0/16:0) | 2 | C | 6 | | | | | 11.3 | 807.3 |
| 1176 | HMDB0001940 | Triamterene | 2 | C | 7 | | | | **Poor water solubility** | 10.7 | 253.3 |
| 443 | HMDB0006740 | CE(20:0) | 2 | C | 8 | | | | | 9.9 | 681.2 |
| 773 | PUBCHEMCID102548 | 1,2-o-Dihexadecyl-Sn-glycerol | 2 | C | 9 | | | | **Poor water solubility** | 10.9 | 541.0 |
| 813 | PUBCHEMCID6683 | Purpurin | 2 | C | 10 | **40** | **80** | | | 16.1 | 256.2 |
| 905 | HMDB0014903 | Metyrosine | 2 | D | 1 | **37** | **75** | **0.5 M HCl** | | 7.4 | 195.2 |
| 4 | PUBCHEMCID80281 | 1-Nonadecanol | 2 | D | 2 | | | | **Poor water solubility** | 12.3 | 284.5 |
| 1000 | HMDB0000786 | Oxypurinol | 2 | D | 3 | **100** | **0** | **1 M NaOH** | | 16.2 | 152.1 |
| 257 | HMDB0034181 | 3-Methylcyclopentadecanone | 2 | D | 4 | | | | | 15.2 | 238.4 |
| 990 | PUBCHEMCID15610 | Nonadecanoic acid methyl este | 2 | D | 5 | | | | | 13.2 | 312.5 |
| 578 | HMDB0011730 | Melezitose | 2 | D | 6 | **100** | **0** | | | 16.4 | 522.5 |
| 682 | HMDB0000548 | Glycerol Tridecanoate | 2 | D | 7 | | | | | 7.5 | 554.8 |
| 565 | HMDB0013773 | D-Leucine | 2 | D | 8 | **69** | **50** | **0.75 M HCl** | | 13.8 | 131.2 |
| 1066 | HMDB0034950 | Rebaudioside A | 2 | D | 9 | **89** | **60** | | | 15.1 | 967.0 |
| 589 | HMDB0003411 | D-Proline | 2 | D | 10 | **100** | **0** | | | 13.3 | 115.1 |
| 909 | HMDB0000639 | Galactaric acid | 2 | E | 1 | **26** | **10** | | | 14.4 | 210.1 |
| | HMDB0000206 | N6-Acetyl-L-lysine | 2 | E | 2 | **100** | **0** | | | 11.9 | 188.2 |
| 301 | HMDB0000873 | 4-Methylcatechol | 2 | E | 3 | **100** | **38** | | | 13.3 | 124.1 |
| 502 | HMDB0003213 | Raffinose | 2 | E | 4 | **75** | **0** | | | 14.9 | 594.5 |
| 887 | HMDB0033833 | Methyl cinnamate | 2 | E | 5 | **82.5** | **75** | | | 16.5 | 162.2 |
| 806 | HMDB0038720 | Dodecyl gallate | 2 | E | 6 | | | | | 13.4 | 338.4 |
| 1080 | HMDB0042008 | Ritalinic acid | 2 | E | 7 | **65** | **50** | **0.6 M NaOH** | | 12.5 | 219.3 |
| 1068 | HMDB0032037 | 1,3-Benzenediol | 2 | E | 8 | **55** | **0** | | | 12.3 | 110.1 |
| 521 | HMDB0001264 | Dehydroascorbic acid | 2 | E | 9 | **50** | **0** | **0.4 M NaOH** | | 6.6 | 174.1 |
| 710 | HMDB0001398 | Guaiacol | 2 | E | 10 | **50** | **0** | | | 15.6 | 124.1 |
| 1081 | HMDB0034436 | Rotenone | 2 | F | 1 | **25** | **92** | | | 14.8 | 394.4 |
| 983 | HMDB0004825 | p-Octopamine | 2 | F | 2 | **60** | **0** | | | 14.8 | 189.6 |
| 100 | HMDB0041683 | 4-Hydroxyphenyl-2-propionic a | 2 | F | 3 | **63.5** | **30** | | | 12.7 | 166.2 |
| 873 | HMDB0014922 | Mefenamic acid | 2 | F | 4 | **85** | **90** | | | 17 | 241.3 |
| 969 | HMDB0240291 | Meglumine | 2 | F | 5 | **75** | **0** | | | 14.9 | 195.2 |
| 55 | HMDB0009059 | PE(18:1(9Z)/18:1(9Z)) | 2 | F | 6 | | | | | 5.5 | 744.1 |
| 101 | PUBCHEMCID14389738 | Arachidic acid methyl ester | 2 | F | 7 | | | | | 13.6 | 326.6 |
| 1073 | HMDB0032876 | Rhein | 2 | F | 8 | **14.2** | **95** | | | 5.7 | 284.2 |
| 136 | HMDB0031861 | 2-Acetylpyrazine | 2 | F | 9 | **65** | **68** | | | 14.2 | 122.1 |
| 718 | HMDB0002345 | Heneicosanoic acid | 2 | F | 10 | | | | | 17.1 | 326.6 |
| 460 | HMDB0002080 | Petroselinic acid | 2 | G | 1 | | | | | 7.1 | 282.5 |
| 134 | HMDB0001879 | Aspirin | 2 | G | 2 | **165** | **0** | | **100% MeOH** | 16.5 | 180.2 |

| | | | | | | | | | | | |
|---|---|---|---|---|---|---|---|---|---|---|---|
| 311 | HMDB0000469 | 5-Hydroxymethyluracil | 2 | G | 3 | **50** | **0** | **0.4 M NaOH** | | 12.5 | 142.1 |
| 538 | HMDB0014411 | Dicumarol | 2 | G | 4 | **42** | **10** | **0.1 M NaOH** | | 14.9 | 336.3 |
| 936 | HMDB0061684 | N-Acetylisoleucine | 2 | G | 5 | **58.6** | **0** | **0.2 M NaOH** | | 17.6 | 173.2 |
| 232 | HMDB0013222 | Beta-Guanidinopropionic acid | 2 | G | 6 | **50** | **0** | **0.4 M NaOH** | | 14.1 | 131.1 |
| 253 | PUBCHEMCID14871 | Monolaurin | 2 | G | 7 | | | | | 10.5 | 274.4 |
| 1013 | PUBCHEMCID23342 | Creatinol phosphate | 2 | G | 8 | **50** | **0** | **0.1 M NaOH** | | 11.4 | 197.1 |
| 743 | HMDB0002434 | Hydroquinone | 2 | G | 9 | **50** | **0** | | | 13.2 | 110.1 |
| 638 | HMDB0014887 | Etodolac | 2 | G | 10 | **50** | **65** | | | 13.2 | 287.4 |
| 875 | HMDB0014952 | Meloxicam | 2 | H | 1 | | | | **Poor water solubility** | 15.6 | 351.4 |
| 880 | HMDB0001921 | 1,1-Dimethylbiguanide | 2 | H | 2 | **50** | **0** | | | 11.4 | 165.6 |
| 760 | PUBCHEMCID84815 | D-Methionine | 2 | H | 3 | **60** | **0** | **0.4 M HCl** | | 16.9 | 149.2 |
| 611 | HMDB0014731 | Ethosuximide | 2 | H | 4 | **60** | **0** | | | 13.6 | 141.2 |
| 615 | HMDB0004992 | Benzocaine | 2 | H | 5 | **60** | **0** | **0.5 M HCl** | | 11.7 | 165.2 |
| 21 | HMDB0014535 | Sulpiride | 2 | H | 6 | **59** | **0** | **0.2 M NaOH** | | 11.8 | 341.4 |
| 494 | HMDB0003546 | Salicin | 2 | H | 7 | **40** | **5** | | | 12.2 | 286.3 |
| 594 | HMDB0003406 | D-Serine | 2 | H | 8 | **80** | **0** | | | 11.9 | 105.1 |
| 775 | HMDB0002092 | Itaconic acid | 2 | H | 9 | **63** | **0** | | | 13.8 | 130.1 |
| 54 | HMDB0008821 | PE(14:0/14:0) | 2 | H | 10 | | | | | 5.3 | 635.9 |
| 902 | HMDB0240266 | Methyl vanillate | 2 | I | 1 | **60** | **0** | **0.2 M NaOH** | | 11.3 | 182.2 |
| 548 | HMDB0014487 | Diltiazem | 2 | I | 2 | **71** | **0** | | | 14.2 | 451.0 |
| 1129 | HMDB0014408 | Sulfisoxazole | 2 | I | 3 | **82** | **40** | **0.1 M NaOH** | | 16.4 | 267.3 |
| 1075 | HMDB0014949 | Ribavirin | 2 | I | 4 | **75** | **0** | | | 12.1 | 244.2 |
| 985 | HMDB0033375 | Octyl gallate | 2 | I | 5 | | | | | 15.3 | 282.3 |
| | PUBCHEMCID99474 | Diosgenin | 2 | I | 6 | | | | | 14.4 | 414.6 |
| 1063 | HMDB0014624 | Raloxifene | 2 | I | 7 | | | | **Poor water solubility** | 11.9 | 510.0 |
| 883 | HMDB0029738 | Indole-3-methyl acetate | 2 | I | 8 | **70** | **50** | | | 14.1 | 189.2 |
| 848 | HMDB0014372 | Lovastatin | 2 | I | 9 | **30** | **90** | | **Poorly soluble** | 12.1 | 404.5 |
| 1117 | HMDB0014565 | Spironolactone | 2 | I | 10 | | | | | 14.4 | 416.6 |
| 847 | HMDB0005000 | Loratadine | 2 | J | 1 | **68** | **75** | | | 13.6 | 382.9 |
| 728 | PUBCHEMCID221071 | Santonin | 2 | J | 2 | | | | **Poor water solubility** | 14.2 | 246.3 |
| 1004 | HMDB0061859 | Methyl hexadecanoic acid | 2 | J | 3 | | | | | 14.1 | 270.5 |
| 249 | PUBCHEMCID13067 | Ethyl 3-indoleacetate | 2 | J | 4 | **68** | **60** | | | 11.6 | 203.2 |
| 793 | HMDB0000740 | Lactulose | 2 | J | 5 | **79** | **0** | | | 15.8 | 342.3 |
| 834 | HMDB0001938 | Lisinopril | 2 | J | 6 | **54** | **0** | **0.2 M NaOH** | | 11.9 | 441.5 |
| 882 | PUBCHEMCID71563 | D-Valine | 2 | J | 7 | **65** | **0** | **0.5 M HCl** | | 13.6 | 117.2 |
| 765 | HMDB0032987 | Ipriflavone | 2 | J | 8 | **70** | **100** | | | 13.9 | 280.3 |
| 1170 | HMDB0003066 | Chalcone | 2 | J | 9 | **70** | **75** | | | 14.7 | 208.3 |
| 792 | PUBCHEMCID71567 | D-Phenylalanine | 2 | J | 10 | **59.6** | **0** | **0.5 M HCl** | | 14.9 | 165.2 |
| 1162 | HMDB0000393 | 3-Hexenedioic acid | 3 | A | 1 | **63** | **50** | | | 16.6 | 144.1 |
| 505 | HMDB0013609 | D-Tryptophan | 3 | A | 2 | **49** | **70** | **0.4 M NaOH** | | 10.8 | 204.2 |
| 1036 | HMDB0014694 | Piroxicam | 3 | A | 3 | **20** | **90** | **0.5 M HCl** | | 16.1 | 331.4 |
| 1217 | HMDB0000900 | Ergocalciferol | 3 | A | 4 | **20** | **95** | **0.5 M HCl** | | 15.6 | 396.7 |
| 1021 | HMDB0006344 | Phenylacetylglutamine | 3 | A | 5 | **47** | **0** | **0.17 M NaOH** | | 14.2 | 264.3 |
| 1158 | HMDB0015256 | Tolbutamide | 3 | A | 6 | **46** | **70** | | | 12.4 | 270.4 |
| 563 | HMDB0000291 | Vanillylmandelic acid | 3 | A | 7 | **50** | **0** | | | 11 | 198.2 |
| 495 | HMDB0003418 | D-Tagatose | 3 | A | 8 | **50** | **0** | | | 13.4 | 180.2 |
| 1040 | HMDB0014998 | Prednisolone | 3 | A | 9 | **76** | **100** | | **Poor water solubility** | 11.2 | 360.4 |
| 943 | HMDB0011757 | N-Acetylvaline | 3 | A | 10 | **50** | **0** | | | 15.2 | 159.2 |
| 1028 | HMDB0013675 | 1,3,5-Trihydroxybenzene | 3 | B | 1 | **50** | **22** | | | 11.4 | 126.1 |
| 749 | HMDB0001925 | Ibuprofen | 3 | B | 2 | **48** | **70** | **0.1 M NaOH** | | 13.3 | 206.3 |
| 722 | HMDB0000857 | Pimelic acid | 3 | B | 3 | **50** | **20** | **0.1 M NaOH** | | 11.8 | 160.2 |
| 963 | HMDB0014760 | Nicardipine | 3 | B | 4 | | | | **Poor water solubility** | 13.3 | 516.0 |
| 1111 | HMDB0015470 | Salicylate-sodium | 3 | B | 5 | **50** | **0** | | | 12.9 | 160.1 |
| 10 | HMDB0014394 | Idoxuridine | 3 | B | 6 | **50** | **0** | **0.1 M NaOH** | | 15.5 | 355.0 |
| 1016 | HMDB0014988 | Perphenazine | 3 | B | 7 | **40** | **80** | **0.1 M HCl** | | 11 | 404.0 |
| 1041 | HMDB0015169 | Procainamide | 3 | B | 8 | **56** | **0** | | | 16.7 | 271.8 |
| 698 | PUBCHEMCID164619 | D-Pinitol | 3 | B | 9 | **37** | **0** | | | 5.2 | 194.2 |
| 739 | HMDB0013318 | Tryptophanamide | 3 | B | 10 | **60** | **0** | | | 12.3 | 239.7 |
| 379 | HMDB0015193 | Amoxicillin | 3 | C | 1 | **60** | **0** | **0.25 M NaOH** | | 12 | 419.5 |
| 487 | HMDB0000630 | Cytosine | 3 | C | 2 | **45** | **0** | **1 M NaOH** | | 13 | 111.1 |
| 72 | HMDB0015051 | Amantadine | 3 | C | 3 | **50** | **0** | **0.8 M HCl** | | 10.9 | 151.3 |
| 1233 | PUBCHEMCID5801 | 2-Aminophenol | 3 | C | 4 | **62** | **0** | **0.6 M HCl** | | 18.5 | 109.1 |
| 709 | HMDB0035762 | 3-(Dimethylaminomethyl)indol | 3 | C | 5 | **50** | **0** | **0.2 M HCl** | | 15.6 | 174.3 |
| 333 | HMDB0002658 | 6-Hydroxynicotinic acid | 3 | C | 6 | **50** | **0** | **0.6 M NaOH** | | 15.9 | 139.1 |
| 428 | HMDB0015328 | Captopril | 3 | C | 7 | **50** | **0** | **0.28 M NaOH** | | 10.2 | 217.3 |
| 426 | HMDB0003033 | Canrenone | 3 | C | 8 | | | | **Poor water solubility** | 11.4 | 340.5 |
| 661 | HMDB0015139 | Ganciclovir | 3 | C | 9 | **50** | **25** | **0.24 M HCl** | | 12.4 | 255.2 |
| 649 | HMDB0015227 | Fluvastatin | 3 | C | 10 | **50** | **60** | | | 13.6 | 433.5 |
| 391 | HMDB0001894 | Aspartame | 3 | D | 1 | **50** | **0** | **0.27 M HCl** | | 10.9 | 294.3 |
| 343 | HMDB0015324 | Acebutolol | 3 | D | 2 | **50** | **0** | | | 13.9 | 372.9 |
| 646 | HMDB0003075 | Flavone | 3 | D | 3 | | | | **Poor water solubility** | 12.4 | 222.2 |
| 466 | HMDB0015372 | Clomipramine | 3 | D | 4 | **25** | **25** | | | 13.4 | 351.3 |
| 462 | HMDB0001413 | Citicoline | 3 | D | 5 | **35** | **0** | | | 10.4 | 546.3 |
| 437 | HMDB0014620 | Chlorpromazine | 3 | D | 6 | **50** | **0** | **0.5 M HCl** | | 5.8 | 318.9 |
| 469 | HMDB0002802 | Cortisone | 3 | D | 7 | **58** | **100** | | **Poor water solubility** | 5.8 | 360.4 |
| 347 | HMDB0014957 | Acetazolamide | 3 | D | 8 | **50** | **0** | **0.4 M NaOH** | | 11.3 | 222.3 |
| 56 | HMDB0000593 | PC(18:1(9Z)/18:1(9Z)) | 3 | D | 9 | | | | **Poor water solubility** | 18 | 786.1 |
| 404 | HMDB0014390 | Benzatropine | 3 | D | 10 | **50** | **0** | | | 10.7 | 403.5 |
| 449 | HMDB0000885 | Cholesteryl palmitic acid | 3 | E | 1 | | | | **Poor water solubility** | 10.5 | 625.1 |
| 658 | HMDB0015570 | Fusidic Acid | 3 | E | 2 | **50** | **0** | **0.2 M NaOH** | | 6 | 538.7 |
| 399 | HMDB0014538 | Beclometasone dipropionate | 3 | E | 3 | | | | **Poor water solubility** | 6.7 | 521.4 |
| 679 | PUBCHEMCID57449 | D-Lysine | 3 | E | 4 | **50** | **0** | | | 12.6 | 146.2 |

| | | | | | | | | | | | |
|---|---|---|---|---|---|---|---|---|---|---|---|
| 609 | HMDB0000153 | Estriol | 3 | E | 5 | | | | Poor water solubility | 17.6 | 288.4 |
| 693 | PUBCHEMCID186078 | Glycolaldehyde dimer | 3 | E | 6 | 34 | 0 | | | 11.7 | 120.1 |
| 778 | HMDB0014608 | Ketorolac | 3 | E | 7 | 75 | 0 | | | 15.1 | 376.4 |
| 1130 | HMDB0014743 | Sulindac | 3 | E | 8 | | | | Poor water solubility | 12 | 356.4 |
| 385 | HMDB0029943 | Arbutin | 3 | E | 9 | 56.4 | 0 | | | 14.1 | 272.3 |
| 1153 | HMDB0034155 | Thiourea | 3 | E | 10 | 30 | 0 | | | 13.6 | 76.1 |
| 193 | PUBCHEMCID69435 | D-Threonine | 3 | F | 1 | 50 | 0 | 0.2 M HCl | | 13.8 | 119.1 |
| 900 | PUBCHEMCID150923 | 3-Methyl-2-oxindole | 3 | F | 2 | 50 | 67 | | | 14 | 147.2 |
| 777 | HMDB0015144 | Ketoprofen | 3 | F | 3 | 50 | 70 | | | 10.9 | 254.3 |
| 74 | HMDB0036458 | 1-Aminocyclopropanecarboxyli | 3 | F | 4 | 50 | 0 | | | 12.9 | 101.1 |
| 61 | PUBCHEMCID10243 | Quinoline-4-carboxylic acid | 3 | F | 5 | 50 | 0 | | | 13.1 | 173.2 |
| 1065 | PUBCHEMCID5280451 | Maleamic acid | 3 | F | 6 | 40 | 0 | 0.22 M NaOH | | 13.7 | 115.1 |
| 324 | PUBCHEMCID439559 | Palatinose | 3 | F | 7 | 50 | 0 | | | 15.9 | 342.3 |
| 1047 | HMDB0059933 | p-Toluenesulfonic acid | 3 | F | 8 | 60 | 0 | | | 18.5 | 190.2 |
| 103 | HMDB0059964 | 2,3,4-Trihydroxybenzoic acid | 3 | F | 9 | 50 | 0 | 0.35 M NaOH | | 14.9 | 170.1 |
| 1087 | HMDB0015687 | Salicylamide | 3 | F | 10 | 22 | 50 | | | 10.8 | 137.1 |
| 395 | HMDB0014327 | Baclofen | 3 | G | 1 | 50 | 0 | 0.4 M HCl | | 14.3 | 213.7 |
| 666 | HMDB0015151 | Glyburide | 3 | G | 2 | 42 | 88 | | Poor water solubility | 15.8 | 494.0 |
| 488 | HMDB0035030 | Amygdalin | 3 | G | 3 | 50 | 0 | | | 14.1 | 457.4 |
| 876 | HMDB0015177 | Memantine | 3 | G | 4 | 50 | 36 | | | 13.8 | 215.8 |
| 955 | HMDB0030542 | Neohesperidin dihydrochalcon | 3 | G | 5 | 50 | 50 | | | 10.7 | 612.6 |
| 267 | HMDB0014395 | Dapsone | 3 | G | 6 | 56 | 0 | 0.5 M HCl | | 18 | 248.3 |
| 1078 | HMDB0001520 | Flavin mononucleotide | 3 | G | 7 | 60 | 200uL H2O | | | 12 | 478.3 |
| 454 | HMDB0014644 | Cimetidine | 3 | G | 8 | 50 | 0 | 0.25 M HCl | | 12.2 | 252.3 |
| 742 | HMDB0014879 | Hydrocortisone | 3 | G | 9 | | | | Poor water solubility | 10.3 | 362.5 |
| 1133 | HMDB0014958 | Tadalafil | 3 | G | 10 | | | | Poor water solubility | 13.7 | 389.4 |
| 372 | HMDB0014732 | Amiloride | 3 | H | 1 | 32 | 75 | | | 14.4 | 302.1 |
| 807 | HMDB0000062 | L-Carnitine | 3 | H | 2 | 50 | 0 | | | 17.2 | 161.2 |
| 648 | HMDB0014615 | Fluoxetine | 3 | H | 3 | 42 | 33 | | | 12.7 | 345.8 |
| 914 | PUBCHEMCID3084068 | 2,6-Dihydroxypyridine HCl | 3 | H | 4 | 18 | | | Poor water solubility | 10.8 | 147.6 |
| 20 | HMDB0001849 | Propranolol | 3 | H | 5 | 50 | 40 | | | 10 | 295.8 |
| 1152 | HMDB0014817 | Thioridazine | 3 | H | 6 | 50 | 18 | | | 13.8 | 407.0 |
| 966 | HMDB0014723 | Nizatidine | 3 | H | 7 | 60 | 0 | 0.2 M HCl | | 18.2 | 331.5 |
| 1043 | HMDB0015202 | Promethazine | 3 | H | 8 | 57 | 0 | | | 17 | 320.9 |
| 401 | HMDB0014520 | Trihexyphenidyl | 3 | H | 9 | 20 | 95 | | Poor water solubility | 14.3 | 337.9 |
| 670 | HMDB0015381 | Gliquidone | 3 | H | 10 | 58.4 | 80 | | | 14.6 | 527.6 |
| 491 | HMDB0059876 | Pantolactone | 3 | I | 1 | 50 | 0 | | | 16.4 | 130.1 |
| 421 | HMDB0001510 | Bupropion | 3 | I | 2 | 54 | 0 | | | 13.9 | 276.2 |
| 583 | HMDB0015273 | Doxepin | 3 | I | 3 | 50 | 0 | | | 10.5 | 315.8 |
| 543 | HMDB0014400 | Diethylstilbestrol | 3 | I | 4 | | | | Poor water solubility | 13.8 | 268.4 |
| 432 | HMDB0015267 | Carvedilol | 3 | I | 5 | 18 | 50 | | | 11.1 | 406.5 |
| 986 | HMDB0005012 | Olanzapine | 3 | I | 6 | | | | Poor water solubility | 10.1 | 312.4 |
| 669 | HMDB0015200 | Glipizide | 3 | I | 7 | | | | Poor water solubility | 10.8 | 445.5 |
| 355 | PUBCHEMCID90301 | 2,4-Dihydroxypyrimidine-5-carl | 3 | I | 8 | 36 | 0 | 0.5 M NaOH | | 14.9 | 156.1 |
| 392 | HMDB0001924 | Atenolol | 3 | I | 9 | 50 | 0 | 0.25 M HCl | | 10.3 | 266.3 |
| 342 | HMDB0014429 | Acarbose | 3 | I | 10 | 74 | 0 | | | 14.8 | 645.6 |
| 411 | HMDB0014586 | Betamethasone | 3 | J | 1 | | | | Poor water solubility | 13.9 | 392.5 |
| 434 | HMDB0005032 | Cetirizine | 3 | J | 2 | 50 | 0 | | | 12.9 | 461.8 |
| 438 | HMDB0014810 | Chlorpropamide | 3 | J | 3 | 50 | 68 | | | 11.1 | 276.7 |
| 483 | HMDB0002991 | Cysteamine | 3 | J | 4 | 40 | 0 | | | 13.3 | 113.6 |
| 967 | PUBCHEMCID74300 | 10-Hydroxydecanoic acid | 3 | J | 5 | | | | Low water solubility | 12.4 | 188.3 |
| 1213 | HMDB0015000 | Vardenafil | 3 | J | 6 | | | | Poor water solubility | 14.1 | 525.6 |
| 934 | HMDB0000812 | N-Acetyl-L-aspartic acid | 3 | J | 7 | 50 | 0 | | | 17 | 175.1 |
| 371 | HMDB0014622 | Amikacin | 3 | J | 8 | 50 | 0 | | | 5.4 | 781.8 |
| 644 | HMDB0015173 | Fenofibrate | 3 | J | 9 | | | | Poor water solubility | 11.8 | 360.8 |
| 872 | HMDB0001939 | Medroxyprogesterone | 3 | J | 10 | | | | Poor water solubility | 10.6 | 344.5 |
| 468 | HMDB0001547 | Corticosterone | 4 | A | 1 | | | | Poor water solubility | 14.3 | 346.5 |
| 809 | HMDB0000904 | Citrulline | 4 | A | 2 | 50 | 0 | | | 18.6 | 175.2 |
| 655 | HMDB0000134 | Fumaric acid | 4 | A | 3 | 43 | 42 | | | 15.3 | 116.1 |
| 459 | HMDB0060460 | cis-4-Hydroxy-D-proline | 4 | A | 4 | 46 | 0 | | | 13.9 | 131.1 |
| 376 | HMDB0015250 | Amiodarone | 4 | A | 5 | 38 | 100 | | | 16.6 | 681.8 |
| 965 | HMDB0015247 | Nifedipine | 4 | A | 6 | 30 | 100 | | Poor water solubility | 12.5 | 346.3 |
| 643 | HMDB0015158 | Felodipine | 4 | A | 7 | 48 | 100 | | | 17.6 | 384.3 |
| 467 | HMDB0001072 | Coenzyme Q10 | 4 | A | 8 | 183 | 0 | | Lipid soluble / 60uL Chlorof | 11 | 863.3 |
| 181 | HMDB0001844 | Methylsuccinic acid | 4 | A | 9 | 44 | 0 | | | 13.3 | 132.1 |
| 637 | HMDB0000622 | Ethylmalonic acid | 4 | A | 10 | 40 | 0 | | | 12.3 | 132.1 |
| 217 | HMDB0029661 | 3',4'-Dihydroxyacetophenone | 4 | B | 1 | 37 | 0 | | | 11.2 | 152.2 |
| 285 | HMDB0007469 | 4-acetylphenol | 4 | B | 2 | 50 | 50 | | | 17.2 | 136.2 |
| 123 | HMDB0029660 | 2',6'-Dihydroxyacetophenone | 4 | B | 3 | 50 | 50 | | | 14.5 | 152.2 |
| 740 | HMDB0001928 | Hydrochlorothiazide | 4 | B | 4 | 40 | 0 | 0.2 M NaOH | | 18.5 | 297.7 |
| 910 | HMDB0031865 | 4'-tert-Butyl-2',6'-dimethyl-3',5 | 4 | B | 5 | | | | Poor water solubility | 15.8 | 294.3 |
| 297 | HMDB0032570 | 4'-Methoxyacetophenone | 4 | B | 6 | 38 | 56 | | | 13.8 | 150.2 |
| 268 | HMDB0041808 | 4,4'-Methylenedianiline | 4 | B | 7 | 46 | 50 | | | 18 | 198.3 |
| 353 | PUBCHEMCID16211665 | Brucine sulfate heptahydrate | 4 | B | 8 | | | | Poor water solubility | 13.7 | 1013.1 |
| 825 | HMDB0000679 | Homocitrulline | 4 | B | 9 | 32 | 0 | | | 11.8 | 189.2 |
| 824 | HMDB0000670 | Homo-L-arginine | 4 | B | 10 | 39 | 0 | | | 11.7 | 224.7 |
| 43 | PUBCHEMCID8487 | 3-Hydroxyacetophenone | 4 | C | 1 | 36 | 60 | | | 11.6 | 136.2 |
| 1094 | HMDB0005010 | Sertraline | 4 | C | 2 | 50 | 50 | | | 13.6 | 342.7 |
| 1006 | HMDB0014853 | Paroxetine | 4 | C | 3 | 40 | 50 | | | 14.7 | 365.8 |
| 137 | PUBCHEMCID93131 | Glycylsarcosine | 4 | C | 4 | 46 | 0 | 0.2 M HCl | | 15 | 146.2 |
| 1025 | HMDB0014397 | Phenytoin | 4 | C | 5 | 50 | 0 | 0.25 M NaOH | | 15.4 | 252.3 |
| 461 | HMDB0005038 | Citalopram | 4 | C | 6 | 43.2 | 0 | | | 10.8 | 405.3 |

| | | | | | | | | | | | |
|---|---|---|---|---|---|---|---|---|---|---|---|
| 1064 | HMDB0001930 | Ranitidine | 4 | C | 7 | 50 | 0 | | | 11 | 350.9 |
| 1030 | HMDB0001565 | Phosphorylcholine | 4 | C | 8 | 50 | 0 | | | 14.4 | 329.7 |
| 1224 | HMDB0029645 | Xanthoxylin | 4 | C | 9 | 50 | 85 | | | 17.4 | 196.2 |
| 116 | HMDB0032629 | 2',5'-Dihydroxyacetophenone | 4 | C | 10 | 50 | 80 | | | 13.5 | 152.2 |
| 989 | HMDB0001913 | Omeprazole | 4 | D | 1 | 40 | 85 | | | 12.8 | 345.4 |
| 1001 | HMDB0015195 | Oxybutynin | 4 | D | 2 | 50 | 0 | | | 15.9 | 357.5 |
| 465 | HMDB0015020 | Clomifene | 4 | D | 3 | 50 | 50 | | | 10.5 | 598.1 |
| 1086 | HMDB0001937 | Salbutamol | 4 | D | 4 | 50 | 50 | | | 11.5 | 239.3 |
| 291 | HMDB0004284 | Tyrosol | 4 | D | 5 | 40 | 40 | | | 12 | 138.2 |
| 1102 | HMDB0000482 | Caprylic acid | 4 | D | 6 | 147 | 0 | | | 14.7 | 166.2 |
| 77 | HMDB0014770 | Docosanol | 4 | D | 7 | | | | Poor water solubility | 25.9 | 326.6 |
| 868 | HMDB0034410 | Mangiferin | 4 | D | 8 | 28 | 45 | | | 6 | 422.3 |
| 606 | HMDB0014344 | Erythromycin | 4 | D | 9 | 45 | 85 | | | 13.5 | 733.9 |
| 369 | HMDB0014581 | Allopurinol | 4 | D | 10 | 131 | 100 | | Insoluble | 13.1 | 136.1 |
| 916 | HMDB0000092 | Dimethylglycine | 4 | E | 1 | 35 | 0 | | | 17.6 | 103.1 |
| 922 | HMDB0002099 | 6-Methyladenine | 4 | E | 2 | 29.2 | 80 | | | 7.3 | 149.2 |
| 799 | HMDB0005008 | Lansoprazole | 4 | E | 3 | 40 | 68 | 0.2 M HCl | | 13.7 | 369.4 |
| 315 | HMDB0136758 | 5,7-Dihydroxy-4-methylcoumar | 4 | E | 4 | 50 | 75 | | | 11.2 | 192.2 |
| 1079 | HMDB0005020 | Risperidone | 4 | E | 5 | 50 | 0 | 0.2 M HCl | | 13.5 | 410.5 |
| 373 | HMDB0000510 | Aminoadipic acid | 4 | E | 6 | 50 | 0 | 0.2 M HCl | | 16.1 | 161.2 |
| 377 | HMDB0014466 | Amitriptyline | 4 | E | 7 | 50 | 0 | | | 13.7 | 313.9 |
| 978 | HMDB0006029 | N-Acetylglutamine | 4 | E | 8 | 32 | 3 | | | 12.6 | 188.2 |
| 667 | HMDB0015252 | Gliclazide | 4 | E | 9 | | | | Poor water solubility | 10.2 | 323.4 |
| 1031 | HMDB0000272 | Phosphoserine | 4 | E | 10 | 30 | 0 | | | 13.7 | 185.1 |
| 1188 | HMDB0014947 | Tropicamide | 4 | F | 1 | 50 | 0 | 0.2 M HCl | | 10.6 | 284.4 |
| 71 | HMDB0001926 | 17a-Ethynylestradiol | 4 | F | 2 | | | | Poor water solubility | 11.4 | 296.4 |
| 1209 | HMDB0014653 | Vancomycin | 4 | F | 3 | 75 | 0 | | | 12.7 | 1485.7 |
| 1143 | HMDB0000872 | Tetradecanedioic acid | 4 | F | 4 | | | | Poor water solubility | 17.5 | 258.4 |
| 797 | HMDB0028694 | Alanylphenylalanine | 4 | F | 5 | 50 | 0 | | | 13.2 | 236.3 |
| 935 | HMDB0001890 | Acetylcysteine | 4 | F | 6 | 40 | 0 | | | 12 | 163.2 |
| 1045 | HMDB0000824 | Propionylcarnitine | 4 | F | 7 | 50 | 0 | | | 14.8 | 253.7 |
| 524 | HMDB0015364 | Dexamethasone | 4 | F | 8 | | | | Poor water solubility | 17.8 | 392.5 |
| 427 | HMDB0002227 | Capsaicin | 4 | F | 9 | 50 | 80 | | | 6.1 | 305.4 |
| 776 | HMDB0005801 | Kaempferol | 4 | F | 10 | 50 | 80 | | | 11.4 | 286.2 |
| 450 | HMDB0062461 | cholest-5-en-3beta-yl octadeca | 4 | G | 1 | | | | Poor water solubility | 13 | 653.1 |
| 668 | HMDB0014367 | Glimepiride | 4 | G | 2 | 29 | 100 | | | 17.5 | 490.6 |
| 766 | HMDB0015163 | Irbesartan | 4 | G | 3 | | | | Poor water solubility | 10.3 | 428.5 |
| 829 | HMDB0035921 | Limonin | 4 | G | 4 | 28 | 100 | | | 5.6 | 470.5 |
| 378 | HMDB0005018 | Amlodipine | 4 | G | 5 | 48 | 100 | | | 10.6 | 408.9 |
| 22 | HMDB0001850 | Verapamil | 4 | G | 6 | 44 | 0 | | | 13.2 | 491.1 |
| 801 | HMDB0000517 | L-Arginine | 4 | G | 7 | 41 | 0 | | | 16.3 | 174.2 |
| 975 | HMDB0029846 | Nonivamide | 4 | G | 8 | 50 | 100 | | | 12.6 | 293.4 |
| 261 | HMDB0001886 | 3-Methylxanthine | 4 | G | 9 | 34 | 50 | 0.4 M NaOH | | 10.5 | 166.1 |
| 179 | HMDB0000202 | Methylmalonic acid | 4 | G | 10 | 34 | 0 | | | 15.4 | 118.1 |
| 464 | HMDB0000094 | Citric acid | 4 | H | 1 | 46 | 0 | | | 18.3 | 192.1 |
| 499 | HMDB0001514 | Glucosamine | 4 | H | 2 | 40 | 0 | | | 12.2 | 215.6 |
| 838 | HMDB0000746 | Hydroxyisocaproic acid | 4 | H | 3 | 15 | 80 | 0.4 M HCl | | 12.3 | 132.2 |
| 254 | HMDB0000491 | 3-Methyl-2-oxovaleric acid | 4 | H | 4 | 12 | 0 | 0.4 M HCl | | 5.7 | 298.4 |
| 1101 | HMDB0031230 | 2-Ethylhexanoic acid | 4 | H | 5 | 46 | 16 | | | 14 | 166.2 |
| 1124 | HMDB0000893 | Suberic acid | 4 | H | 6 | 42 | 50 | | | 12.6 | 174.2 |
| 924 | HMDB0000212 | N-Acetylgalactosamine | 4 | H | 7 | 50 | 0 | | | 11.4 | 221.2 |
| 410 | HMDB0000043 | Betaine | 4 | H | 8 | 33.6 | 0 | | | 16.8 | 117.2 |
| 780 | HMDB0000715 | Kynurenic acid | 4 | H | 9 | 50 | 35 | 0.2 M NaOH | | 14.7 | 189.2 |
| 1246 | HMDB0000112 | gamma-Aminobutyric acid | 4 | H | 10 | 29 | 0 | | | 17.3 | 103.1 |
| 358 | HMDB0000034 | Adenine | 4 | I | 1 | 26 | 0 | 0.1 M HCl | | 15.7 | 171.6 |
| 753 | HMDB0000738 | Indole | 4 | I | 2 | 36 | 50 | | | 14.5 | 117.2 |
| 560 | HMDB0000991 | DL-2-Aminooctanoic acid | 4 | I | 3 | 22 | 50 | | | 14.8 | 159.2 |
| 144 | HMDB0000905 | Deoxyadenosine monophospha | 4 | I | 4 | 48 | 0 | 0.2 M NaOH | | 14.6 | 331.2 |
| 803 | HMDB0000168 | L-Asparagine | 4 | I | 5 | 30 | 0 | 0.2 M HCl | | 11.9 | 132.1 |
| 942 | HMDB0000866 | N-Acetyl-L-tyrosine | 4 | I | 6 | 35 | 0 | | | 10.6 | 223.2 |
| 507 | HMDB0000098 | D-Xylose | 4 | I | 7 | 48 | 0 | | | 18 | 150.1 |
| 1125 | HMDB0000254 | Succinic acid | 4 | I | 8 | 35 | 0 | 0.1 M NaOH | | 10.8 | 118.1 |
| 1197 | HMDB0000294 | Urea | 4 | I | 9 | 34 | 0 | | | 17.3 | 60.1 |
| 1191 | HMDB0000306 | Tyramine | 4 | I | 10 | 30 | 0 | 0.2 M NaOH | | 11.9 | 137.2 |
| 157 | HMDB0000617 | 2-Furoic acid | 4 | J | 1 | 31 | 0 | 0.1 M NaOH | | 12.5 | 112.1 |
| 808 | HMDB0000033 | Carnosine | 4 | J | 2 | 50 | 0 | | | 13.5 | 226.2 |
| 1195 | HMDB0000947 | Undecanoic acid | 4 | J | 3 | | | | Poor water solubility | 12.9 | 186.3 |
| 1136 | HMDB0000251 | Taurine | 4 | J | 4 | 36.6 | 0 | | | 18.3 | 125.2 |
| 447 | HMDB0006725 | CE(14:0) | 4 | J | 5 | | | | Poor water solubility | 12.5 | 597.0 |
| 367 | HMDB0001358 | Retinal | 4 | J | 6 | 59 | 100 | | Poor water solubility | 5.9 | 284.4 |
| 452 | HMDB0000097 | Choline | 4 | J | 7 | 36 | 0 | | | 19 | 139.6 |
| 1196 | HMDB0000300 | Uracil | 4 | J | 8 | 27 | 40 | | | 13.5 | 112.1 |
| 314 | HMDB0003701 | Dimethylbenzimidazole | 4 | J | 9 | 40 | 100 | | | 14.2 | 146.2 |
| 567 | HMDB0002287 | Homocysteine thiolactone | 4 | J | 10 | 40 | 0 | | | 12.5 | 153.6 |
| 1084 | HMDB0001185 | S-Adenosylmethionine | 5 | A | 1 | 50 | 0 | | | 13.7 | 570.7 |
| 387 | HMDB0005042 | Aripiprazole | 5 | A | 2 | 43 | 90 | 0.5 M HCl | | 17.5 | 448.4 |
| 3 | HMDB0003153 | Epigallocatechin gallate | 5 | A | 3 | 47 | 30 | | | 11.8 | 476.4 |
| 396 | HMDB0134890 | 5,6,7-trihydroxy-2-phenyl-4H-cl | 5 | A | 4 | 28 | 85 | 0.2 M HCl | | 11.2 | 270.2 |
| 1106 | HMDB0000210 | Pantothenic acid | 5 | A | 5 | 36.7 | 0 | | | 11 | 241.2 |
| 849 | HMDB0000159 | L-Phenylalanine | 5 | A | 6 | 40 | 0 | 0.25 M HCl | | 18.3 | 165.2 |
| 150 | HMDB0000012 | Deoxyuridine | 5 | A | 7 | 48 | 0 | | | 12.4 | 228.2 |
| 741 | HMDB0000764 | Hydrocinnamic acid | 5 | A | 8 | 31.5 | 50 | | | 12.6 | 150.2 |

| | | | | | | | | | | | |
|---|---|---|---|---|---|---|---|---|---|---|---|
| 1223 | HMDB0000299 | Xanthosine | 5 | A | 9 | **35** | **0** | **0.4 M NaOH** | | 5.3 | 320.3 |
| 58 | HMDB0008923 | PE(16:0/16:0) | 5 | A | 10 | | | | **Poor water solubility** | 7.4 | 692.0 |
| 349 | PUBCHEMCID21477529 | Glycine lauryl ester HCl | 5 | B | 1 | | | | **Poor water solubility** | 5.3 | 279.9 |
| 1120 | HMDB0034154 | Methyl stearate | 5 | B | 2 | | | | **Poor water solubility** | 10.3 | 298.5 |
| 851 | HMDB0000267 | Pyroglutamic acid | 5 | B | 3 | **40** | **0** | | | 15.7 | 129.1 |
| 332 | HMDB0001901 | Aminocaproic acid | 5 | B | 4 | **38** | **0** | | | 15.1 | 131.2 |
| 196 | HMDB0011749 | 2-Piperidinone | 5 | B | 5 | **28** | **75** | | | 11.1 | 99.1 |
| 32 | HMDB0000555 | 3-Methyladipic acid | 5 | B | 6 | **40** | **0** | | | 14.8 | 160.2 |
| 575 | HMDB0000962 | Lipoamide | 5 | B | 7 | **43.3** | **100** | | **Poor water solubility** | 13 | 205.3 |
| 703 | HMDB0000759 | Glycylleucine | 5 | B | 8 | **31** | **0** | **0.6 M HCl** | | 16.3 | 188.2 |
| 866 | HMDB0001262 | Maltotriose | 5 | B | 9 | **52.5** | **0** | | | 10.5 | 504.4 |
| 795 | HMDB0000161 | L-Alanine | 5 | B | 10 | **29.2** | **0** | | | 11.7 | 89.1 |
| 219 | HMDB0001336 | 3,4-Dihydroxybenzeneacetic ac | 5 | C | 1 | **43** | **0** | | | 12.9 | 168.2 |
| 599 | HMDB0000650 | D-alpha-Aminobutyric acid | 5 | C | 2 | **32** | **0** | | | 12.3 | 103.1 |
| 1077 | HMDB0000244 | Riboflavin | 5 | C | 3 | **60** | **50** | | | 12 | 376.4 |
| 374 | HMDB0001147 | Aminomalonic acid | 5 | C | 4 | **38** | **0** | | | 15.1 | 119.1 |
| 800 | HMDB0001851 | L-Arabitol | 5 | C | 5 | **32.8** | **0** | | | 13.1 | 152.2 |
| 441 | HMDB0000067 | Cholesterol | 5 | C | 6 | **139** | **0** | | **100% chloroform** | 13.9 | 386.7 |
| 964 | HMDB0001488 | Nicotinic acid | 5 | C | 7 | **36** | **0** | **0.3 M NaOH** | | 14.4 | 123.1 |
| 979 | HMDB0000201 | L-Acetylcarnitine | 5 | C | 8 | **60** | **0** | | | 19 | 239.7 |
| 856 | HMDB0000167 | L-Threonine | 5 | C | 9 | **42.6** | **0** | | | 12.8 | 119.1 |
| 417 | HMDB0000030 | Biotin | 5 | C | 10 | **32** | **90** | | **Low water solubility** | 16.3 | 246.3 |
| 66 | HMDB0001414 | Putrescine | 5 | D | 1 | **50** | **0** | | | 17.1 | 161.1 |
| 877 | HMDB0001892 | Menadione | 5 | D | 2 | **41** | **90** | | **Low water solubility** | 14 | 172.2 |
| 183 | HMDB0012322 | 2-Naphthol | 5 | D | 3 | **46** | **46** | | | 14.8 | 144.2 |
| 1108 | HMDB0003502 | myo-Inositol hexakisphosphate | 5 | D | 4 | **90** | **0** | | | 12.7 | 923.8 |
| 822 | HMDB0000177 | L-Histidine | 5 | D | 5 | **34** | **0** | | | 14.1 | 155.2 |
| 820 | HMDB0000125 | Glutathione | 5 | D | 6 | **50** | **0** | | | 11 | 307.3 |
| 1092 | HMDB0000792 | Sebacic acid | 5 | D | 7 | **86.6** | **33** | | **66% MeOH** | 13 | 202.3 |
| 696 | HMDB0000128 | Guanidoacetic acid | 5 | D | 8 | **25** | **40** | **1 M NaOH** | | 11 | 117.1 |
| 362 | HMDB0000538 | Adenosine triphosphate | 5 | D | 9 | **50** | **0** | | | 12.3 | 551.1 |
| 419 | HMDB0032133 | Bisphenol A | 5 | D | 10 | | | | **Poor water solubility** | 11.1 | 228.3 |
| 117 | HMDB0004062 | Gentisate aldehyde | 5 | E | 1 | **44.5** | **65** | | | 17.8 | 138.1 |
| 730 | PUBCHEMCID12127 | 3-Hydroxy-4-methoxybenzaldel | 5 | E | 2 | **44** | **50** | | | 15.8 | 152.2 |
| 162 | HMDB0059709 | 2-Hydroxybenzyl alcohol | 5 | E | 3 | **34.5** | **50** | | | 13.8 | 124.1 |
| 223 | HMDB0034315 | 3-(3,4-Dimethoxyphenyl)-2-pro | 5 | E | 4 | **38** | **75** | | | 11.3 | 208.2 |
| 222 | HMDB0059763 | 3,4-Dimethoxybenzoic acid | 5 | E | 5 | **30** | **60** | | | 14.9 | 182.2 |
| 356 | HMDB0004296 | Acrylamide | 5 | E | 6 | **29.6** | **0** | | | 14.8 | 71.8 |
| 823 | HMDB0003431 | L-Histidinol | 5 | E | 7 | **36** | **0** | | | 10.7 | 214.1 |
| 675 | HMDB0011530 | MG(0:0/14:0/0:0) | 5 | E | 8 | | | | **Poor water solubility** | 6.3 | 302.5 |
| 85 | HMDB0003331 | 1-Methyladenosine | 5 | E | 9 | **20** | **60** | **1 M HCl** | | 5.5 | 281.3 |
| 1193 | HMDB0060079 | [(2R,3S,4R,5R)-5-(2,4-dioxo-1,2, | 5 | E | 10 | **38** | **0** | | | 6.2 | 610.3 |
| 804 | HMDB0000191 | L-Aspartic acid | 5 | F | 1 | **35** | **0** | **1 M NaOH** | | 12.7 | 133.1 |
| 714 | HMDB0000132 | Guanine | 5 | F | 2 | **25** | **0** | **1 M NaOH** | | 17.6 | 151.1 |
| 1052 | HMDB0001491 | Pyridoxal 5'-phosphate | 5 | F | 3 | **38** | **0** | **0.2 M NaOH** | | 11.7 | 247.1 |
| 1144 | HMDB0005789 | Tetrahydrocurcumin | 5 | F | 4 | **45** | **60** | | | 11.8 | 372.4 |
| 33 | PUBCHEMCID92926 | Glycerol monodecanoate | 5 | F | 5 | | | | **Poor water solubility** | 6.4 | 246.3 |
| 1110 | HMDB0000243 | Pyruvic acid | 5 | F | 6 | **125** | **0** | | | 12.5 | 110.0 |
| 677 | HMDB0011564 | MG(16:0/0:0/0:0) | 5 | F | 7 | | | | **Poor water solubility** | 5.5 | 330.5 |
| 88 | HMDB0000699 | 1-Methylnicotinamide | 5 | F | 8 | **38** | **0** | | | 15.1 | 172.6 |
| 448 | HMDB0000918 | Cholesteryl oleate | 5 | F | 9 | | | | **Poor water solubility** | 5.4 | 651.1 |
| 338 | HMDB0001107 | 7-Methylguanosine | 5 | F | 10 | **28** | **0** | | | 5.6 | 297.3 |
| 1178 | HMDB0001160 | Tricosanoic acid | 5 | G | 1 | | | | **Poor water solubility** | 5.8 | 354.6 |
| 320 | HMDB0000763 | 5-Hydroxyindoleacetic acid | 5 | G | 2 | **37** | **46** | | | 11.1 | 191.2 |
| 650 | HMDB0000121 | Folic acid | 5 | G | 3 | **55** | **0** | **0.6 M NaOH** | | 13.9 | 441.4 |
| 894 | PUBCHEMCID5281787 | Caffeic acid phenethyl ester | 5 | G | 4 | **50** | **60** | | | 14.8 | 284.3 |
| 423 | HMDB0000039 | Butyric acid | 5 | G | 5 | **40** | **0** | | | 12 | 110.1 |
| 1090 | HMDB0000271 | Sarcosine | 5 | G | 6 | **33.2** | **0** | | | 13.3 | 89.1 |
| 999 | HMDB0002329 | Oxalic acid | 5 | G | 7 | **40** | **0** | | | 17.6 | 90.0 |
| 359 | HMDB0000050 | Adenosine | 5 | G | 8 | **63** | **0** | **0.6 M NaOH** | | 19 | 267.2 |
| 912 | HMDB0000806 | Myristic acid | 5 | G | 9 | | | | **Poor water solubility** | 16.2 | 228.4 |
| 225 | HMDB0013677 | 3,5-Dihydroxybenzoic acid | 5 | G | 10 | **50** | **42** | | | 16.9 | 154.1 |
| 993 | HMDB0000214 | Ornithine | 5 | H | 1 | **49** | **0** | | | 14.6 | 168.6 |
| 713 | HMDB0003584 | Taurocyamine | 5 | H | 2 | **45** | **0** | | | 11.7 | 167.2 |
| 26 | HMDB0040894 | Ampelopsin D | 5 | H | 3 | | | | **Poor water solubility** | 5.3 | 304.3 |
| 570 | HMDB0000684 | L-Kynurenine | 5 | H | 4 | **26** | **0** | **0.4 M HCl** | | 5.7 | 208.2 |
| 705 | HMDB0028853 | Glycyltyrosine | 5 | H | 5 | **48** | **0** | **0.2 M HCl** | | 10.5 | 238.2 |
| 651 | HMDB0001562 | Folinic acid | 5 | H | 6 | **50** | **42** | | | 11.6 | 511.5 |
| 925 | HMDB0002931 | N-Acetylserine | 5 | H | 7 | **25** | **0** | | | 5.1 | 147.1 |
| 529 | HMDB0001401 | Glucose 6-phosphate | 5 | H | 8 | **103** | **0** | | | 10.3 | 282.1 |
| 451 | HMDB0000619 | Cholic acid | 5 | H | 9 | **80** | **100** | | **Partially soluble** | 16.1 | 408.6 |
| 736 | HMDB0000870 | Histamine | 5 | H | 10 | **50** | **0** | | | 11.2 | 184.1 |
| 471 | HMDB0000064 | Creatine | 5 | I | 1 | **50** | **0** | | | 17.5 | 131.1 |
| 1211 | HMDB0012308 | Vanillin | 5 | I | 2 | **50** | **0** | | | 12.2 | 152.2 |
| 862 | HMDB0000176 | Maleic acid | 5 | I | 3 | **40** | **0** | | | 16.1 | 116.1 |
| 859 | HMDB0005800 | Luteolin | 5 | I | 4 | **50** | **90** | | | 10.7 | 286.2 |
| 287 | HMDB0000500 | 4-Hydroxybenzoic acid | 5 | I | 5 | **50** | **63** | | | 16.2 | 138.1 |
| 692 | HMDB0000123 | Glycine | 5 | I | 6 | **38** | **0** | | | 12.1 | 75.1 |
| 424 | HMDB0001964 | Caffeic acid | 5 | I | 7 | **50** | **34** | | | 14.8 | 180.2 |
| 817 | HMDB0000148 | L-Glutamic acid | 5 | I | 8 | **50** | **0** | **0.2 M NaOH** | | 15.8 | 147.1 |
| 946 | HMDB0001487 | NADH | 5 | I | 9 | **147** | **0** | | | 14.7 | 709.4 |
| 699 | HMDB0000708 | Glycoursodeoxycholic acid | 5 | I | 10 | | | | **Poor water solubility** | 5.9 | 449.6 |

| | | | | | | | | | | | |
|---|---|---|---|---|---|---|---|---|---|---|---|
| 337 | HMDB0000897 | 7-Methylguanine | 5 | J | 1 | **35** | **0** | **0.2 M NaOH** | | 7 | 165.2 |
| 245 | HMDB0002096 | 3-Indolebutyric acid | 5 | J | 2 | **48** | **50** | | | 11.4 | 203.2 |
| 436 | HMDB0003164 | Chlorogenic acid | 5 | J | 3 | **54** | **40** | | | 13.4 | 354.3 |
| 1132 | HMDB0002085 | Syringic acid | 5 | J | 4 | **48** | **50** | | | 13.6 | 198.2 |
| 865 | HMDB0000163 | D-Maltose | 5 | J | 5 | **60** | **0** | | | 15 | 342.3 |
| 363 | HMDB0000058 | Cyclic AMP | 5 | J | 6 | **33** | **0** | **0.1 M NaOH** | | 6.1 | 329.2 |
| 759 | HMDB0003320 | Indole-3-carboxylic acid | 5 | J | 7 | **50** | **62** | | | 13 | 161.2 |
| 577 | HMDB0000169 | D-Mannose | 5 | J | 8 | **50** | **0** | | | 13.5 | 180.2 |
| 205 | HMDB0002199 | Desaminotyrosine | 5 | J | 9 | **50** | **45** | | | 14.1 | 166.2 |
| 394 | HMDB0000784 | Azelaic acid | 5 | J | 10 | **118** | **100** | | **Poor water solubility** | 11.8 | 188.2 |
| 246 | HMDB0002302 | Indole-3-propionic acid | 6 | A | 1 | **50** | **50** | | | 14.2 | 189.2 |
| 995 | HMDB0000669 | ortho-Hydroxyphenylacetic aci | 6 | A | 2 | **50** | **50** | | | 16.4 | 152.2 |
| 1194 | HMDB0029865 | Umbelliferone | 6 | A | 3 | **50** | **73** | | | 10.9 | 162.1 |
| 361 | HMDB0000045 | Adenosine monophosphate | 6 | A | 4 | **68** | **0** | **0.2 M NaOH** | | 16.9 | 347.2 |
| 290 | HMDB0000822 | p-Hydroxymandelic acid | 6 | A | 5 | **42** | **50** | | | 19.1 | 186.2 |
| 1029 | HMDB0000263 | Phosphoenolpyruvic acid | 6 | A | 6 | **146** | **0** | | | 14.6 | 206.1 |
| 1022 | HMDB0000821 | Phenylacetylglycine | 6 | A | 7 | **50** | **40** | | | 12.3 | 193.2 |
| 326 | HMDB0002894 | 5-Methylcytosine | 6 | A | 8 | **40** | **0** | **0.3 M HCl** | | 12.4 | 125.1 |
| 293 | HMDB0000707 | 4-Hydroxyphenylpyruvic acid | 6 | A | 9 | **50** | **0** | | | 11.8 | 180.2 |
| 1230 | HMDB0001586 | Glucose 1-phosphate | 6 | A | 10 | **141** | **0** | | | 14.1 | 376.2 |
| 1137 | HMDB0000036 | Taurocholic acid | 6 | B | 1 | **115** | **100** | | **Poor water solubility** | 11.5 | 537.7 |
| 923 | HMDB0012881 | Acetylcarnosine | 6 | B | 2 | **54** | **0** | | | 10.9 | 268.3 |
| 938 | HMDB0000446 | N-alpha-Acetyl-L-lysine | 6 | B | 3 | **50** | **0** | | | 12 | 188.2 |
| | HMDB0000543 | Benzenebutanoic acid | 6 | B | 4 | **50** | **62** | | | 15.7 | 164.2 |
| 1219 | HMDB0034227 | alpha-Tocopherol acetate | 6 | B | 5 | **126** | **0** | | | 12.6 | 472.7 |
| 370 | HMDB0002204 | Astaxanthin | 6 | B | 6 | **107** | **0** | | **100% MeOH** | 10.7 | 596.8 |
| 798 | HMDB0000194 | Anserine | 6 | B | 7 | **50** | **0** | | | 12.8 | 240.3 |
| 316 | HMDB0003192 | 5-Aminoimidazole-4-carboxam | 6 | B | 8 | **40** | **0** | **0.2 M HCl** | | 13.4 | 126.1 |
| 1037 | HMDB0000613 | Erythronic acid | 6 | B | 9 | **50** | **0** | | | 18.5 | 174.2 |
| 971 | HMDB0003633 | N-Methyltyramine | 6 | B | 10 | **50** | **0** | | | 11.8 | 187.7 |
| 298 | HMDB0140294 | 2-hydroxy-2-(4-methoxyphenyl | 6 | C | 1 | **50** | **37** | | | 13.5 | 182.2 |
| 517 | HMDB0003417 | D-Cysteine | 6 | C | 2 | **48** | **0** | **0.2 M HCl** | | 10.7 | 121.2 |
| 444 | PUBCHEMCID88251 | (R)-2-Amino-3-methoxypropan | 6 | C | 3 | **50** | **0** | | | 16.9 | 119.1 |
| 485 | HMDB0000082 | Cytidine triphosphate | 6 | C | 4 | **74** | **0** | | | 11.1 | 527.1 |
| 1172 | HMDB0000930 | trans-Cinnamic acid | 6 | C | 5 | **40** | **60** | | | 10.2 | 148.2 |
| 660 | HMDB0005807 | Gallic acid | 6 | C | 6 | **50** | **0** | **0.2 M NaOH** | | 14.4 | 170.1 |
| 1212 | HMDB0029663 | Vanillin acetate | 6 | C | 7 | **35** | **85** | | | 17.4 | 194.2 |
| 1149 | HMDB0000235 | Thiamine | 6 | C | 8 | **50** | **0** | | | 20.6 | 337.3 |
| 382 | HMDB0002124 | Apigenin | 6 | C | 9 | **32** | **75** | | **Poor water solubility** | 12.8 | 270.2 |
| 1177 | HMDB0031193 | 1,2,3-Propanetricarboxylic acid | 6 | C | 10 | **68** | **0** | | | 13.9 | 176.1 |
| 334 | HMDB0032394 | 6-Methylcoumarin | 6 | D | 1 | **42** | **78** | | | 18.8 | 160.2 |
| 519 | HMDB0000511 | Capric acid | 6 | D | 2 | **10** | **100** | | **Poor water solubility** | 14.4 | 172.3 |
| 1179 | HMDB0002327 | 1,11-Undecanedicarboxylic aci | 6 | D | 3 | | | | **Poor water solubility** | 10.1 | 244.3 |
| 1155 | HMDB0000262 | Thymine | 6 | D | 4 | **42** | **0** | **0.2 M NaOH** | | 10.3 | 126.1 |
| 591 | HMDB0000283 | D-Ribose | 6 | D | 5 | **129** | **0** | | | 12.9 | 150.1 |
| 1044 | PUBCHEMCID643511 | (S)-(+)-5-Hydroxymethyl-2-pyrr | 6 | D | 6 | **42** | **36** | | | 11.9 | 115.1 |
| 785 | HMDB0000716 | L-Pipecolic acid | 6 | D | 7 | **45** | **0** | | | 12.6 | 129.2 |
| 991 | HMDB0006049 | O-Phosphotyrosine | 6 | D | 8 | **50** | **0** | **0.15 M NaOH** | | 13.2 | 261.2 |
| 920 | HMDB0005923 | N4-Acetylcytidine | 6 | D | 9 | **35** | **60** | **0.2 M NaOH** | | 11.3 | 285.3 |
| 202 | HMDB0000375 | 3-(3-Hydroxyphenyl)propanoic | 6 | D | 10 | **50** | **40** | | | 11.6 | 166.2 |
| 796 | HMDB0028685 | Alanylglutamine | 6 | E | 1 | **60** | **0** | | | 12.1 | 217.2 |
| 19 | HMDB0002670 | Naringenin | 6 | E | 2 | **66.5** | **50** | | | 13.3 | 272.3 |
| 12 | HMDB0002780 | Catechin | 6 | E | 3 | **62** | **50** | | | 12.4 | 290.3 |
| 603 | HMDB0002209 | Equol | 6 | E | 4 | **50** | **60** | | | 12.7 | 242.3 |
| 593 | HMDB0000663 | Glucaric acid | 6 | E | 5 | **58** | **0** | **0.2 M NaOH** | | 10.3 | 248.2 |
| 1105 | HMDB0000625 | Gluconic acid | 6 | E | 6 | **58** | **0** | | | 16 | 218.1 |
| 839 | HMDB0000687 | L-Leucine | 6 | E | 7 | **50** | **0** | **0.2 M HCl** | | 12.1 | 131.2 |
| 526 | HMDB0001058 | Fructose 1,6-bisphosphate | 6 | E | 8 | **108** | **0** | | | 10.8 | 550.2 |
| 762 | HMDB0000195 | Inosine | 6 | E | 9 | **52** | **0** | | | 15.7 | 268.2 |
| 214 | HMDB0002441 | 3,3-Dimethylglutaric acid | 6 | E | 10 | **50** | **0** | | | 15.9 | 160.2 |
| 508 | HMDB0000563 | D-Phenyllactic acid | 6 | F | 1 | **50** | **0** | **0.4 M NaOH** | | 10.6 | 166.2 |
| 586 | HMDB0004231 | Panthenol | 6 | F | 2 | **54** | **0** | | | 21.8 | 205.3 |
| 738 | HMDB0000434 | Homoveratric acid | 6 | F | 3 | **48** | **26** | | | 15.2 | 196.2 |
| 516 | HMDB0014405 | Cycloserine | 6 | F | 4 | **45** | **0** | | | 11.4 | 102.1 |
| 819 | HMDB0003337 | Oxidized glutathione | 6 | F | 5 | **100** | **0** | | | 10.5 | 612.6 |
| 328 | HMDB0000884 | Ribothymidine | 6 | F | 6 | **50** | **0** | | | 15.2 | 258.2 |
| 574 | HMDB0034252 | 2-Aminoheptanedioic acid | 6 | F | 7 | **32** | **70** | **0.25 M NaOH** | | 14.2 | 175.2 |
| 939 | HMDB0011745 | N-Acetyl-L-methionine | 6 | F | 8 | **50** | **0** | **0.2 M NaOH** | | 14.9 | 191.3 |
| 929 | HMDB0001138 | N-Acetyl-L-glutamic acid | 6 | F | 9 | **46** | **0** | **0.3 M NaOH** | | 16.2 | 189.2 |
| 439 | HMDB0000908 | 5alpha-Cholestanol | 6 | F | 10 | **80** | **50** | | **50% DMSO, 50% chlorofo** | 15.2 | 388.7 |
| 511 | HMDB0001151 | Allose | 6 | G | 1 | **50** | **0** | | | 13.9 | 180.2 |
| 858 | HMDB0000158 | L-Tyrosine | 6 | G | 2 | **50** | **60** | **0.5 M HCl** | | 16.7 | 181.2 |
| 867 | HMDB0000703 | Mandelic acid | 6 | G | 3 | **50** | **40** | **0.1 M NaOH** | | 12 | 152.2 |
| 83 | HMDB0029736 | 1H-Imidazole-1-acetic acid | 6 | G | 4 | **50** | **0** | | | 13.5 | 126.1 |
| 1184 | HMDB0041992 | Pivalic acid | 6 | G | 5 | **40** | **0** | | | 13 | 102.1 |
| 486 | HMDB0000095 | Cytidine monophosphate | 6 | G | 6 | **50** | **0** | **0.2 M NaOH** | | 11.4 | 323.2 |
| 597 | PUBCHEMCID70286 | 3-(Furan-2-yl)propanoic acid | 6 | G | 7 | **48** | **0** | **0.2 M NaOH** | | 15.1 | 140.1 |
| 16 | HMDB0034932 | Fenchol | 6 | G | 8 | **50** | **32** | | | 15.4 | 154.3 |
| 277 | PUBCHEMCID239828 | 3-(Dimethylamino)propanoic a | 6 | G | 9 | **50** | **0** | **0.1 M NaOH** | | 13.1 | 153.6 |
| 854 | HMDB0034365 | L-Theanine | 6 | G | 10 | **50** | **0** | | | 13.9 | 174.2 |
| 339 | PUBCHEMCID65575 | (+)-Cedrol | 6 | H | 1 | | | | **Poor water solubility** | 16.3 | 222.4 |
| 656 | HMDB0040594 | (±)-Furaneol | 6 | H | 2 | **40** | **0** | | | 10.6 | 128.1 |

| | | | | | | | | | | | |
|---|---|---|---|---|---|---|---|---|---|---|---|
| 561 | HMDB0000008 | 2-Hydroxybutyric acid | 6 | H | 3 | **43** | **0** | | | 12.8 | 126.1 |
| 283 | HMDB0000118 | Homovanillic acid | 6 | H | 4 | **50** | **60** | | | 12.6 | 182.2 |
| 1097 | HMDB0032616 | Sinapic acid | 6 | H | 5 | **57** | **50** | | | 11.5 | 224.2 |
| 149 | HMDB0000071 | Deoxyinosine | 6 | H | 6 | **50** | **0** | **0.1 M NaOH** | | 15.9 | 252.2 |
| 756 | HMDB0000197 | Indoleacetic acid | 6 | H | 7 | **60** | **50** | | | 12.1 | 197.2 |
| 7 | PUBCHEMCID637817 | α-Methyl-cinnamic acid | 6 | H | 8 | **50** | **57** | | | 11.2 | 162.2 |
| 559 | HMDB0002006 | 2,3-Diaminopropionic acid | 6 | H | 9 | **40** | **0** | | | 10.3 | 140.6 |
| 1245 | HMDB0000229 | Nicotinamide ribotide | 6 | H | 10 | **55** | **0** | | | 11 | 334.2 |
| 1139 | HMDB0000874 | Tauroursodeoxycholic acid | 6 | I | 1 | **134** | **0** | | **Poor water solubility** | 13.4 | 521.7 |
| 445 | PUBCHEMCID132213 | Acetyl-D-alanine | 6 | I | 2 | **48** | **0** | | | 12.5 | 131.1 |
| 1161 | HMDB0002349 | trans-trans-Muconic acid | 6 | I | 3 | **24** | **75** | | | 11.4 | 142.1 |
| 280 | HMDB0029306 | 4-Ethylphenol | 6 | I | 4 | **37** | **50** | | | 14.8 | 122.2 |
| 1057 | HMDB0013674 | 1,2,3-Trihydroxybenzene | 6 | I | 5 | **46** | **30** | | | 18.8 | 126.1 |
| 1007 | HMDB0002035 | 4-Hydroxycinnamic acid | 6 | I | 6 | **44** | **40** | | | 15.3 | 164.2 |
| 126 | HMDB0029680 | 2,6-Dimethoxy-4-methylphenol | 6 | I | 7 | | | | **Poor water solubility** | 17.3 | 168.2 |
| 306 | PUBCHEMCID30938 | 3-(2-Methylphenyl)propanoic a | 6 | I | 8 | **43** | **67** | | | 13 | 164.2 |
| 1214 | HMDB0000305 | Vitamin A | 6 | I | 9 | | | | **Poor water solubility** | 5.3 | 286.5 |
| 1121 | HMDB0002350 | Octadecanol | 6 | I | 10 | | | | **Poor water solubility** | 14.8 | 270.5 |
| 124 | HMDB0013676 | 2,6-Dihydroxybenzoic acid | 6 | J | 1 | **44** | **26** | | | 16.8 | 154.1 |
| 240 | HMDB0001713 | m-Coumaric acid | 6 | J | 2 | **50** | **34** | | | 16.3 | 164.2 |
| 265 | HMDB0033723 | 4-(4-Hydroxyphenyl)-2-butanor | 6 | J | 3 | **48** | **40** | | | 11.5 | 164.2 |
| 1008 | HMDB0001858 | p-Cresol | 6 | J | 4 | **50** | **40** | | **Toxic** | 12.6 | 108.1 |
| 238 | HMDB0002466 | 3-Hydroxybenzoic acid | 6 | J | 5 | **50** | **33** | | | 15.3 | 138.1 |
| 368 | HMDB0000462 | Allantoin | 6 | J | 6 | **24** | **72** | | | 11.8 | 158.1 |
| 490 | HMDB0000660 | D-Fructose | 6 | J | 7 | **125** | **0** | | | 12.5 | 180.2 |
| 319 | HMDB0003355 | 5-Aminopentanoic acid | 6 | J | 8 | **40** | **0** | | | 12.5 | 117.2 |
| 531 | HMDB0000127 | D-Glucuronic acid | 6 | J | 9 | **53** | **0** | | | 15.7 | 208.1 |
| 1100 | HMDB0000126 | Glycerol 3-phosphate | 6 | J | 10 | **34** | **0** | | | 5.4 | 370.4 |
| 435 | HMDB0000518 | Chenodeoxycholic acid | 7 | A | 1 | **106** | **100** | | **Poor water solubility** | 10.6 | 392.6 |
| 1012 | HMDB0000826 | Pentadecanoic acid | 7 | A | 2 | | | | **Poor water solubility** | 14.6 | 242.4 |
| 671 | HMDB0000661 | Glutaric acid | 7 | A | 3 | **43** | | | | 17.3 | 132.1 |
| 937 | HMDB0011756 | N-Acetylleucine | 7 | A | 4 | **48** | **30** | | | 14 | 173.2 |
| 932 | HMDB0004620 | N-a-Acetyl-L-arginine | 7 | A | 5 | **60** | **0** | | | 12.2 | 252.3 |
| 930 | HMDB0000532 | Acetylglycine | 7 | A | 6 | **40** | **0** | | | 17.5 | 117.1 |
| 473 | HMDB0000562 | Creatinine | 7 | A | 7 | **45** | **0** | | | 18.1 | 113.1 |
| 717 | HMDB0001397 | Guanosine monophosphate | 7 | A | 8 | **87** | **0** | | | 17.4 | 407.2 |
| 662 | HMDB0003217 | Genistein | 7 | A | 9 | **47** | **56** | | **Low water solubility** | 17 | 270.2 |
| 288 | HMDB0032584 | 4-Hydroxychalcone | 7 | A | 10 | **45** | **75** | | **Low water solubility** | 19.1 | 224.3 |
| 1165 | HMDB0000734 | Indoleacrylic acid | 7 | B | 1 | **37** | **66** | | | 11.7 | 187.2 |
| 855 | HMDB0000943 | Threonic acid | 7 | B | 2 | **54** | **0** | **0.5 M HCl** | | 13.6 | 310.3 |
| 415 | HMDB0000054 | Bilirubin | 7 | B | 3 | **100** | **100** | | **Poor water solubility** | 10 | 584.7 |
| 1203 | HMDB0000946 | Ursodeoxycholic acid | 7 | B | 4 | | | | **Poor water solubility** | 11.7 | 392.6 |
| 186 | HMDB0000205 | Phenylpyruvic acid | 7 | B | 5 | **40** | **57** | | | 13.7 | 164.2 |
| 572 | HMDB0059720 | Meta-Tyrosine | 7 | B | 6 | **50** | **0** | | | 10.7 | 181.2 |
| 472 | HMDB0001511 | Phosphocreatine | 7 | B | 7 | **66** | **0** | | | 13.2 | 327.1 |
| 1180 | HMDB0000910 | Tridecanoic acid | 7 | B | 8 | | | | **Poor water solubility** | 12.7 | 214.3 |
| 509 | HMDB0003312 | Daidzein | 7 | B | 9 | **40** | **80** | | **Low water solubility** | 10 | 254.2 |
| 125 | PUBCHEMCID3001376 | Thiamine disulfide | 7 | B | 10 | **100** | **0** | **0.5 M HCl** | | 10 | 562.7 |
| 397 | HMDB0041832 | Baicalin | 7 | C | 1 | **42** | **70** | | | 11.9 | 446.4 |
| 1138 | HMDB0000896 | Taurodeoxycholic acid | 7 | C | 2 | | | | **Poor water solubility** | 13.1 | 539.7 |
| 1146 | HMDB0002825 | Theobromine | 7 | C | 3 | **12** | **75** | | **Partially soluble** | 17.9 | 180.2 |
| 962 | HMDB0001406 | Niacinamide | 7 | C | 4 | **40** | **0** | | | 15.7 | 122.1 |
| 1023 | HMDB0061916 | Phenylglyoxal | 7 | C | 5 | **36** | **70** | | | 12.6 | 152.2 |
| 850 | HMDB0000162 | L-Proline | 7 | C | 6 | **38** | **0** | | | 15 | 115.1 |
| 994 | HMDB0000226 | Orotic acid | 7 | C | 7 | **50** | **0** | **0.5 M NaOH** | | 10.3 | 156.1 |
| 1167 | HMDB0000725 | 4-Hydroxyproline | 7 | C | 8 | **36** | **0** | | | 10.8 | 131.1 |
| 811 | HMDB0000574 | L-Cysteine | 7 | C | 9 | **34** | **0** | **0.5 M HCl** | | 15.7 | 121.2 |
| 1126 | HMDB0032523 | Succinic anhydride | 7 | C | 10 | **34** | **64** | | | 13.7 | 100.1 |
| 787 | HMDB0000646 | L-Arabinose | 7 | D | 1 | **50** | **0** | | | 12.1 | 150.1 |
| 860 | HMDB0000883 | L-Valine | 7 | D | 2 | **40** | **0** | | | 13.6 | 117.2 |
| 745 | HMDB0003338 | Hydroxylamine | 7 | D | 3 | **50** | **0** | | | 13 | 164.1 |
| 1095 | HMDB0003070 | Shikimic acid | 7 | D | 4 | **50** | **0** | | | 12.9 | 174.2 |
| 128 | HMDB0034158 | 2,6-Dimethoxyphenol | 7 | D | 5 | **50** | **36** | | | 14.1 | 154.2 |
| 1198 | HMDB0000289 | Uric acid | 7 | D | 6 | **15** | **20** | **1 M NaOH** | | 14.3 | 168.1 |
| 816 | HMDB0000174 | L-Fucose | 7 | D | 7 | **50** | **0** | | | 11.5 | 164.2 |
| 120 | HMDB0004812 | 2,5-Furandicarboxylic acid | 7 | D | 8 | **40** | **0** | **0.6 M NaOH** | | 14.2 | 156.1 |
| 1222 | HMDB0000292 | Xanthine | 7 | D | 9 | **38** | **20** | **0.3 M NaOH** | | 17.2 | 152.1 |
| 1053 | HMDB0001545 | Pyridoxal | 7 | D | 10 | **50** | **0** | | | 11.6 | 203.6 |
| 826 | HMDB0000719 | L-Homoserine | 7 | E | 1 | **40** | **0** | | | 11.9 | 119.1 |
| 782 | HMDB0000139 | Glyceric acid | 7 | E | 2 | **50** | **0** | | | 12.3 | 268.2 |
| 1103 | HMDB0031340 | Cyclamic acid | 7 | E | 3 | **46** | **0** | | | 11.5 | 201.2 |
| 482 | PUBCHEMCID2734767 | β-Alanine methyl ester hydrocl | 7 | E | 4 | **40** | **0** | | | 15.9 | 139.6 |
| 678 | HMDB0011131 | MG(18:0/0:0/0:0) | 7 | E | 5 | | | | **Poor water solubility** | 5.3 | 358.6 |
| 683 | HMDB0005414 | TG(20:0/20:0/20:0) | 7 | E | 6 | | | | **Poor water solubility** | 6 | 975.7 |
| 944 | HMDB0000230 | N-Acetylneuraminic acid | 7 | E | 7 | **58** | **0** | | | 11.5 | 309.3 |
| 1026 | HMDB0003306 | Phloretin | 7 | E | 8 | **50** | **50** | | | 11.6 | 274.3 |
| 304 | HMDB0013292 | p-Methylhippuric acid | 7 | E | 9 | **50** | **45** | | | 11 | 193.2 |
| 687 | HMDB0042989 | TG(15:0/15:0/15:0) | 7 | E | 10 | | | | **Poor water solubility** | 7.4 | 765.2 |
| 102 | HMDB0002074 | 2,2-Dimethylsuccinic acid | 7 | F | 1 | **50** | **0** | | | 11.7 | 146.1 |
| 1122 | HMDB0000937 | Stigmasterol | 7 | F | 2 | **137** | **0** | | **100% chloroform** | 13.7 | 412.7 |
| 941 | HMDB0094701 | N-Acetylproline | 7 | F | 3 | **50** | **0** | | | 13.9 | 157.2 |
| 758 | HMDB0029737 | Indole-3-carboxaldehyde | 7 | F | 4 | **31** | **75** | | | 12.5 | 145.2 |

| | | | | | | | | | | | |
|---|---|---|---|---|---|---|---|---|---|---|---|
| 641 | HMDB0001919 | Famotidine | 7 | F | 5 | **61** | **0** | **0.4 M HCl** | | 18.3 | 337.5 |
| 960 | PUBCHEMCID445883 | L-Isoglutamine | 7 | F | 6 | **40** | **0** | **0.1 M HCl** | | 11.8 | 146.1 |
| 276 | HMDB0029300 | p-Anisidine | 7 | F | 7 | | | | **Poor water solubility** | 13.9 | 123.2 |
| 537 | HMDB0014724 | Diclofenac | 7 | F | 8 | **25** | **70** | | | 10.8 | 296.2 |
| 596 | PUBCHEMCID6926 | 2-Aminobenzenesulfonic acid | 7 | F | 9 | **45** | **6** | **0.2 M HCl** | | 13.6 | 173.2 |
| 360 | HMDB0001341 | ADP | 7 | F | 10 | **80** | **0** | | | 16 | 465.3 |
| 365 | HMDB0001432 | Agmatine | 7 | G | 1 | **48** | **0** | | | 14.4 | 228.3 |
| 601 | HMDB0000573 | Elaidic acid | 7 | G | 2 | | | | **Poor water solubility** | 5.9 | 282.5 |
| 726 | HMDB0000672 | Hexadecanedioic acid | 7 | G | 3 | **135** | **100** | | **Poor water solubility** | 13.5 | 286.4 |
| 201 | PUBCHEMCID3033877 | Glycerol-monocaprylate | 7 | G | 4 | | | | **Low water solubility** | 8.5 | 218.3 |
| 227 | HMDB0003474 | 3,5-Diiodo-L-tyrosine | 7 | G | 5 | **50** | **0** | **0.6 M HCl** | | 10.2 | 433.0 |
| 716 | HMDB0001273 | Guanosine triphosphate | 7 | G | 6 | **123** | **0** | | | 7.4 | 621.2 |
| 313 | HMDB0004058 | 5,6-Dihydroxyindole | 7 | G | 7 | | | | **Poor water solubility** | 5.4 | 149.2 |
| 1168 | HMDB0000958 | trans-Aconitic acid | 7 | G | 8 | **48** | **0** | | | 14.8 | 174.1 |
| 104 | HMDB0036584 | Tetramethylpyrazine | 7 | G | 9 | **45** | **0** | **0.5 M HCl** | | 14.6 | 136.2 |
| 504 | HMDB0000975 | Trehalose | 7 | G | 10 | **112** | **0** | | | 11.6 | 378.3 |
| 1181 | PUBCHEMCID1549521 | 2-Furanacrolein | 7 | H | 1 | | | | **Poor water solubility** | 11.4 | 122.1 |
| 6 | PUBCHEMCID91194 | Cholesteryl Erucate | 7 | H | 2 | | | | **Poor water solubility** | 6 | 707.2 |
| 1076 | HMDB0000508 | Ribitol | 7 | H | 3 | **40** | **0** | | | 12.1 | 152.2 |
| 1048 | HMDB0240265 | Puerarin | 7 | H | 4 | **63** | **40** | | | 15.8 | 416.4 |
| 987 | HMDB0002364 | Oleanolic acid | 7 | H | 5 | | | | **Poor water solubility** | 12.3 | 456.7 |
| 1210 | HMDB0000484 | Vanillic acid | 7 | H | 6 | **45** | **40** | | | 11.3 | 168.2 |
| 1241 | HMDB0002520 | beta-Glycerophosphoric acid | 7 | H | 7 | **50** | **0** | | | 15.1 | 306.1 |
| 790 | HMDB0000956 | Tartaric acid | 7 | H | 8 | **46** | **0** | | | 18.5 | 150.1 |
| 498 | HMDB0000143 | D-Galactose | 7 | H | 9 | **50** | **0** | | | 16.9 | 180.2 |
| 1127 | HMDB0031554 | Sucralose | 7 | H | 10 | **73** | **0** | | | 21.9 | 397.6 |
| 281 | HMDB0003464 | 4-Guanidinobutanoic acid | 7 | I | 1 | **46** | **0** | | | 14.2 | 145.2 |
| 1202 | HMDB0000288 | Uridine 5'-monophosphate | 7 | I | 2 | **138** | **100uL H2O** | | | 13.8 | 368.1 |
| 307 | HMDB0001232 | 4-Nitrophenol | 7 | I | 3 | **40** | **32** | | | 11.5 | 139.1 |
| 840 | HMDB0000182 | L-Lysine | 7 | I | 4 | **42** | **0** | | | 12.5 | 146.2 |
| 187 | HMDB0000208 | Oxoglutaric acid | 7 | I | 5 | **46** | **0** | | | 19.1 | 146.1 |
| 853 | HMDB0001266 | L-Sorbose | 7 | I | 6 | **50** | **0** | | | 16.4 | 180.2 |
| 845 | HMDB0001645 | L-Norleucine | 7 | I | 7 | **38** | **0** | **0.4 M HCl** | | 11.4 | 131.2 |
| 1032 | HMDB0002107 | Phthalic acid | 7 | I | 8 | **50** | **48** | | | 21.4 | 166.1 |
| 402 | HMDB0001870 | Benzoic acid | 7 | I | 9 | **46** | **50** | | | 18.2 | 122.1 |
| 953 | HMDB0062795 | N-carbamoylglutamic Acid | 7 | I | 10 | **45** | **0** | **0.4 M NaOH** | | 10 | 190.2 |
| 746 | HMDB0035227 | Hydroxypropanedioic acid | 7 | J | 1 | **28** | **0** | | | 5.6 | 120.1 |
| 212 | HMDB0000265 | Liothyronine | 7 | J | 2 | **22** | **70** | **0.2 M HCl** | | 9.5 | 651.0 |
| 704 | HMDB0000721 | Glycylproline | 7 | J | 3 | **44** | **0** | | | 11.5 | 172.2 |
| 545 | HMDB0000076 | Dihydrouracil | 7 | J | 4 | **32** | **38** | **0.6 M NaOH** | | 12.7 | 114.1 |
| 1109 | HMDB0000237 | Propionic acid | 7 | J | 5 | **30** | **0** | | | 11.3 | 96.1 |
| 604 | HMDB0000878 | Ergosterol | 7 | J | 6 | | | | **Poor water solubility** | 13.7 | 396.7 |
| 94 | HMDB0010382 | LysoPC(16:0/0:0) | 7 | J | 7 | **109** | **0** | | **50% MeOH 50% Chlorofor** | 10.9 | 495.6 |
| 364 | HMDB0000448 | Adipic acid | 7 | J | 8 | **38** | **45** | | | 15.3 | 146.1 |
| 139 | HMDB0000149 | Ethanolamine | 7 | J | 9 | **32** | **0** | | | 16.4 | 97.5 |
| 590 | HMDB0000805 | Pyrrolidonecarboxylic acid | 7 | J | 10 | **40** | **0** | | | 11.8 | 129.1 |
| 259 | HMDB0000466 | 3-Methylindole | 8 | A | 1 | **25** | **50** | | | 10.2 | 131.2 |
| 282 | HMDB0000954 | Ferulic acid | 8 | A | 2 | **45** | **45** | | | 16.8 | 194.2 |
| 568 | HMDB0000575 | DL-Homocystine | 8 | A | 3 | **50** | **0** | **0.5 M HCl** | | 11.1 | 268.4 |
| 926 | HMDB0013713 | N-Acetyltryptophan | 8 | A | 4 | **50** | **60** | **0.2 M NaOH** | | 12.8 | 246.3 |
| 582 | HMDB0000073 | Dopamine | 8 | A | 5 | **50** | **0** | | | 11.2 | 189.6 |
| 1231 | HMDB0000186 | Alpha-Lactose | 8 | A | 6 | **52** | **0** | | | 15.6 | 360.3 |
| 737 | HMDB0000130 | Homogentisic acid | 8 | A | 7 | **28** | **50** | | | 5.6 | 168.2 |
| 752 | HMDB0059912 | Indigo Carmine | 8 | A | 8 | **44** | **75** | | | 17.5 | 466.4 |
| 1074 | HMDB0031786 | Rhodamine B | 8 | A | 9 | | | | **Poor water solubility** | 17.3 | 479.0 |
| 546 | HMDB0001882 | Dihydroxyacetone | 8 | A | 10 | **32** | **0** | | | 12.6 | 90.1 |
| 242 | HMDB0000440 | 3-Hydroxyphenylacetic acid | 8 | B | 1 | **24** | **70** | | | 12.1 | 152.2 |
| 156 | PUBCHEMCID150866 | 2-(4-Methylphenyl)propionic ac | 8 | B | 2 | **50** | **50** | | | 14.8 | 164.2 |
| 350 | HMDB0000895 | Acetylcholine | 8 | B | 3 | **50** | **0** | | | 15.4 | 181.7 |
| 755 | HMDB0029739 | Indole-3-acetamide | 8 | B | 4 | **34** | **50** | | | 10.1 | 174.2 |
| 241 | HMDB0031816 | 3-Hydroxyflavone | 8 | B | 5 | **12** | **90** | | **Low water solubility** | 11 | 238.2 |
| 90 | HMDB0012138 | 1-Naphthol | 8 | B | 6 | | | | **Poor water solubility** | 16.5 | 144.2 |
| 109 | HMDB0000232 | Quinolinic acid | 8 | B | 7 | **20** | **50** | | | 15.7 | 167.1 |
| 53 | HMDB0007866 | PC(14:0/14:0) | 8 | B | 8 | | | | **Poor water solubility** | 12.2 | 677.9 |
| 1098 | HMDB0029432 | (S)C(S)S-S-Methylcysteine sulfo | 8 | B | 9 | **37** | **0** | | | 7.4 | 151.2 |
| 375 | HMDB0001833 | Aminopterin | 8 | B | 10 | **26** | **0** | | | 4 | 440.4 |
| 110 | PUBCHEMCID15609 | Heptadecanoic Acid methyl est | 8 | C | 1 | | | | **Poor water solubility** | 12.6 | 284.5 |
| 673 | HMDB0007098 | DG(16:0/16:0/0:0) | 8 | C | 2 | | | | **Poor water solubility** | 5 | 568.9 |
| 947 | HMDB0000217 | NADP | 8 | C | 3 | **52** | **0** | | | 5.2 | 765.4 |
| 1150 | HMDB0001372 | Thiamine pyrophosphate | 8 | C | 4 | **156** | **0** | | | 15.6 | 460.8 |
| 1200 | HMDB0000295 | Uridine 5'-diphosphate | 8 | C | 5 | **68** | **0** | | | 11.5 | 448.1 |
| 1174 | HMDB0000301 | Urocanic acid | 8 | C | 6 | **30** | **50** | **0.2 M HCl** | | 11.5 | 138.1 |
| 1229 | HMDB0000645 | Galactose 1-phosphate | 8 | C | 7 | **50** | **0** | | | 5.1 | 336.3 |
| 165 | HMDB0000711 | Hydroxyoctanoic acid | 8 | C | 8 | **46** | **45** | | | 10.1 | 160.2 |
| 706 | HMDB0028854 | Glycylvaline | 8 | C | 9 | **50** | **0** | | | 13.1 | 174.2 |
| 815 | HMDB0000720 | Levulinic acid | 8 | C | 10 | **45** | **0** | | | 18 | 116.1 |
| 213 | HMDB0000509 | Senecioic acid | 8 | D | 1 | **23** | **30** | | | 14 | 100.1 |
| 665 | HMDB0003559 | Gibberellin A3 | 8 | D | 2 | | | | **Poor water solubility** | 16 | 346.4 |
| 911 | HMDB0002755 | Myricetin | 8 | D | 3 | **42** | **50** | | | 12.7 | 318.2 |
| 1157 | HMDB0034732 | Thymoquinone | 8 | D | 4 | **16** | **80** | | **Low water solubility** | 10.4 | 164.2 |
| 48 | HMDB0011188 | TG(12:0/12:0/12:0) | 8 | D | 5 | | | | **Poor water solubility** | 14.4 | 639.0 |
| 132 | HMDB0033161 | 2,6-Pyridinedicarboxylic acid | 8 | D | 6 | **44** | **50** | | | 17.9 | 167.1 |

| | | | | | | | | | | | |
|---|---|---|---|---|---|---|---|---|---|---|---|
| 479 | HMDB0000211 | myo-Inositol | 8 | D | 7 | 48 | 0 | | | 15.2 | 180.2 |
| 968 | HMDB0002393 | N-Methyl-D-aspartic acid | 8 | D | 8 | 28 | 0 | | | 5.7 | 147.1 |
| 1069 | HMDB0003747 | Resveratrol | 8 | D | 9 | 50 | 60 | | | 15.4 | 228.2 |
| 654 | HMDB0005808 | Formononetin | 8 | D | 10 | 17 | 80 | | Poor water solubility | 10.1 | 268.3 |
| 453 | HMDB0036619 | 5,7-Dihydroxyflavone | 8 | E | 1 | 33 | 80 | | | 15.9 | 254.2 |
| 206 | PUBCHEMCID2214 | Acetovanillone | 8 | E | 2 | 31 | 50 | | | 12.5 | 166.2 |
| 676 | HMDB0011567 | MG(18:1(9Z)/0:0/0:0) | 8 | E | 3 | | | | Poor water solubility | 7.7 | 356.5 |
| 1099 | HMDB0000086 | Glycerophosphocholine | 8 | E | 4 | 50 | 0 | | | 14.5 | 258.2 |
| 286 | HMDB0011718 | 4-Hydroxybenzaldehyde | 8 | E | 5 | 36.5 | 25 | | | 14.6 | 122.1 |
| 532 | HMDB0003339 | D-Glutamic acid | 8 | E | 6 | 38 | 0 | | | 15.3 | 147.1 |
| 1060 | HMDB0005794 | Quercetin | 8 | E | 7 | 50 | 66 | | | 14.9 | 302.2 |
| 2 | HMDB0001871 | Epicatechin | 8 | E | 8 | 41 | 33 | | | 12.2 | 290.3 |
| 60 | HMDB0007158 | DG(18:0/18:0/0:0) | 8 | E | 9 | | | | Poor water solubility | 7.6 | 625.0 |
| 46 | HMDB0046381 | TG(22:0/22:0/22:0) | 8 | E | 10 | | | | Poor water solubility | 10.1 | 1059.8 |
| 573 | PUBCHEMCID5320521 | Raspberry ketone glucoside | 8 | F | 1 | 50 | 45 | | | 10.9 | 326.3 |
| 59 | HMDB0008991 | PE(18:0/18:0) | 8 | F | 2 | | | | Poor water solubility | 5.3 | 748.1 |
| 86 | HMDB0003646 | N-Methylhydantoin | 8 | F | 3 | 38 | 33 | | | 11.6 | 114.1 |
| 881 | PUBCHEMCID11008044 | N,N-dimethyl-L-Valine | 8 | F | 4 | 35 | 33 | | | 10.5 | 145.2 |
| 420 | PUBCHEMCID16038806 | O-Methyl-L-serine hydrochloric | 8 | F | 5 | 50 | 0 | | | 10.1 | 155.6 |
| 118 | HMDB0000152 | Gentisic acid | 8 | F | 6 | 46 | 33 | | | 13.9 | 154.1 |
| 981 | HMDB0002055 | o-Cresol | 8 | F | 7 | 31.5 | 50 | | Toxic | 12.6 | 108.1 |
| 592 | HMDB0029881 | D-Glucaric acid | 8 | F | 8 | 29 | 0 | | | 5.8 | 210.1 |
| 429 | HMDB0014704 | Carbamazepine | 8 | F | 9 | 116 | | | Low water solubility | 17.4 | 236.3 |
| 476 | HMDB0002269 | Curcumin | 8 | F | 10 | 21 | 100 | | Poor water solubility | 11.1 | 368.4 |
| 244 | HMDB0006524 | 3-Indoleacetonitrile | 8 | G | 1 | 38 | 50 | | | 15.5 | 156.2 |
| 446 | HMDB0000610 | CE(18:2(9Z,12Z)) | 8 | G | 2 | | | | | 5 | 649.1 |
| 528 | HMDB0001254 | Glucosamine 6-phosphate | 8 | G | 3 | 31 | 0 | | | 6.2 | 259.2 |
| 178 | HMDB0011723 | 2-Methylhippuric acid | 8 | G | 4 | 38 | 33 | | | 11.5 | 193.2 |
| 1034 | HMDB0029377 | Piperine | 8 | G | 5 | 30 | 85 | | | 10.6 | 285.3 |
| 15 | PUBCHEMCID440049 | D-Mannosamine HCl | 8 | G | 6 | 38 | 0 | | | 11.4 | 215.6 |
| 956 | HMDB0034566 | Neotame | 8 | G | 7 | 30 | 65 | | | 15.4 | 378.5 |
| 708 | HMDB0028995 | Phenylalanylglycine | 8 | G | 8 | 33 | 50 | 0.4 M HCl | | 13.1 | 222.2 |
| 997 | HMDB0041965 | O-Toluidine | 8 | G | 9 | | | | Carcinogenic | 10.3 | 143.6 |
| 1089 | PUBCHEMCID19700 | Erioglaucine disodium salt | 8 | G | 10 | 50 | 0 | | | 10.6 | 792.9 |
| 234 | HMDB0030776 | Maltol | 8 | H | 1 | 14 | 80 | | | 12.5 | 126.1 |
| 273 | HMDB0001392 | p-Aminobenzoic acid | 8 | H | 2 | 34 | 32 | | | 10.4 | 137.1 |
| 1187 | HMDB0062590 | Tropate | 8 | H | 3 | 38 | 32 | | | 11.6 | 166.2 |
| 1128 | HMDB0000258 | Sucrose | 8 | H | 4 | 103 | 0 | | | 10.3 | 342.3 |
| 595 | HMDB0000247 | Sorbitol | 8 | H | 5 | 42 | 0 | | | 12.6 | 182.2 |
| 700 | HMDB0011733 | Glycyl-glycine | 8 | H | 6 | 42 | 0 | | | 12.7 | 132.1 |
| 403 | HMDB0001587 | Phenylglyoxylic acid | 8 | H | 7 | 32 | 50 | | | 12.7 | 150.1 |
| 1140 | HMDB0002428 | Terephthalic acid | 8 | H | 8 | | | | Poor water solubility | 11.1 | 166.1 |
| 107 | HMDB0000397 | 2-Pyrocatechuic acid | 8 | H | 9 | 20 | 20 | 0.3 M NaOH | | 10.9 | 154.1 |
| 231 | HMDB0000444 | 3-Furoic acid | 8 | H | 10 | 36 | 33 | | | 10.7 | 112.1 |
| 292 | HMDB0000020 | p-Hydroxyphenylacetic acid | 8 | I | 1 | 36 | 32 | | | 11 | 152.2 |
| 976 | HMDB0014325 | Masoprocol | 8 | I | 2 | 48 | 45 | | | 11.9 | 302.4 |
| 1225 | HMDB0000881 | Xanthurenic acid | 8 | I | 3 | 20 | 30 | | | 10.7 | 205.2 |
| 772 | HMDB0060665 | Isonicotinic acid | 8 | I | 4 | 40 | 30 | | | 13.8 | 123.1 |
| 852 | HMDB0000187 | L-Serine | 8 | I | 5 | 33 | 0 | | | 10 | 105.1 |
| 1038 | HMDB0029581 | (2E,4E)-2,4-Hexadienoic acid | 8 | I | 6 | 47 | 0 | | | 14.2 | 150.2 |
| 744 | HMDB0000115 | Glycolic acid | 8 | I | 7 | 26 | 0 | | | 11.5 | 76.1 |
| 1189 | HMDB0000303 | Tryptamine | 8 | I | 8 | 36 | 40 | | | 10.7 | 160.2 |
| 846 | HMDB0013716 | Norvaline | 8 | I | 9 | 38 | 0 | 0.2 M HCl | | 12.2 | 117.2 |
| 127 | HMDB0029273 | 2,6-Dimethoxybenzoic acid | 8 | I | 10 | 30 | 30 | | | 10 | 182.2 |
| 1185 | HMDB0000906 | Trimethylamine | 8 | J | 1 | 30 | 0 | | | 12.4 | 95.6 |
| 1085 | HMDB0029723 | Saccharin | 8 | J | 2 | 50 | 0 | | | 13.2 | 241.2 |
| 1226 | HMDB0002917 | D-Xylitol | 8 | J | 3 | 38 | 0 | | | 11.5 | 152.2 |
| 895 | HMDB0029806 | Methyl nicotinate | 8 | J | 4 | 42 | 30 | | | 13.4 | 137.1 |
| 489 | HMDB0029942 | D-Arabinose | 8 | J | 5 | 42 | 0 | | | 12.7 | 150.1 |
| 345 | HMDB0031645 | Acetamide | 8 | J | 6 | 20 | 0 | | | 12.3 | 59.1 |
| 940 | HMDB0000512 | N-Acetyl-L-phenylalanine | 8 | J | 7 | 45 | 0 | 0.5 M NaOH | | 10 | 207.2 |
| 779 | HMDB0032923 | Kojic acid | 8 | J | 8 | 39 | 30 | | | 11.7 | 142.1 |
| 220 | HMDB0001856 | Protocatechuic acid | 8 | J | 9 | 31 | 38 | | | 10 | 154.1 |
| 112 | HMDB0029666 | 2,4-Dihydroxybenzoic acid | 8 | J | 10 | 40 | 30 | 0.2 M NaOH | | 13 | 154.1 |
| 69 | PUBCHEMCID7314 | Lactobionic acid | 9 | A | 1 | 66 | 0 | | | 13.2 | 358.3 |
| 515 | HMDB0006483 | D-Aspartic acid | 9 | A | 2 | 30 | 0 | | | 10.8 | 133.1 |
| 140 | HMDB0001906 | 2-Aminoisobutyric acid | 9 | A | 3 | 34.5 | 0 | | | 13.8 | 103.1 |
| 143 | HMDB0000101 | Deoxyadenosine | 9 | A | 4 | 25 | 60 | 0.3 M HCl | | 13.1 | 251.2 |
| 530 | HMDB0006355 | D-Glucurono-6,3-lactone | 9 | A | 5 | 40 | 0 | | | 11.9 | 176.1 |
| 652 | HMDB0034022 | 6-Hydroxy-5-[(4-sulfophenyl)az | 9 | A | 6 | 37 | 0 | | | 11 | 452.4 |
| 948 | HMDB0000221 | NADPH | 9 | A | 7 | 112 | 0 | | | 5.6 | 833.4 |
| 14 | PUBCHEMCID72669 | Methyl 3-aminopyrazine-2-carb | 9 | A | 8 | 9 | 80 | | | 11.1 | 153.1 |
| 980 | HMDB0003011 | O-Acetylserine | 9 | A | 9 | 50 | 0 | | | 11 | 183.6 |
| 1190 | HMDB0003447 | Tryptophol | 9 | A | 10 | 50 | 50 | | | 10 | 161.2 |
| 243 | HMDB0000700 | Hydroxypropionic acid | 9 | B | 1 | 24 | 0 | | | 7.1 | 90.1 |
| 970 | HMDB0003152 | N-Methylnicotinamide | 9 | B | 2 | 39 | 38 | | | 10.1 | 136.2 |
| 168 | PUBCHEMCID94220 | (R)-(+)-Lactamide | 9 | B | 3 | 26 | 0 | | | 10.3 | 89.1 |
| 1002 | HMDB0029911 | 1-O-alpha-D-Glucopyranosyl-D- | 9 | B | 4 | 100 | 0 | | | 10.3 | 688.6 |
| 1017 | HMDB0035018 | 2-Phenylethyl 3-phenyl-2-prope | 9 | B | 5 | | | | Low water solubility | 12.6 | 252.3 |
| 927 | HMDB0001129 | N-Acetylmannosamine | 9 | B | 6 | 50 | 0 | | | 10 | 221.2 |
| 38 | HMDB0000748 | L-3-Phenyllactic acid | 9 | B | 7 | 36 | 0 | | | 10.9 | 166.2 |
| 1049 | HMDB0001366 | Purine | 9 | B | 8 | 33 | 0 | | | 10 | 120.1 |

| | | | | | | | | | | | |
|---|---|---|---|---|---|---|---|---|---|---|---|
| 837 | HMDB0000761 | Lithocholic acid | 9 | B | 9 | **103** | **100** | | **Poor water solubility** | 10.2 | 376.6 |
| 1208 | HMDB0014323 | Valsartan | 9 | B | 10 | | | | **Poor water solubility** | 10.9 | 435.5 |
| 247 | HMDB0000021 | Iodotyrosine | 9 | C | 1 | **23** | **68** | | | 10.1 | 307.1 |
| 1112 | HMDB0000951 | Taurochenodesoxycholic acid | 9 | C | 2 | | | | **Poor water solubility** | 12.7 | 521.7 |
| 945 | HMDB0001238 | N-Acetylserotonin | 9 | C | 3 | **25.5** | **50** | | | 5.1 | 218.3 |
| 145 | HMDB0000014 | Deoxycytidine | 9 | C | 4 | **50** | **0** | | | 10.4 | 227.2 |
| 763 | HMDB0000189 | Inosine triphosphate | 9 | C | 5 | **53** | **0** | | | 5.3 | 574.1 |
| 1054 | HMDB0001431 | Pyridoxamine | 9 | C | 6 | **50** | **0** | | | 12.7 | 241.1 |
| 255 | HMDB0011600 | 3-Methyladenine | 9 | C | 7 | **26** | **0** | **0.4 M HCl** | | 5.8 | 149.2 |
| 325 | HMDB0000982 | 5-Methylcytidine | 9 | C | 8 | **34** | **0** | | | 5.1 | 257.3 |
| 928 | HMDB0000803 | beta-N-Acetylglucosamine | 9 | C | 9 | **50** | **0** | | | 14.1 | 221.2 |
| 512 | HMDB0000568 | D-Arabitol | 9 | C | 10 | **38** | **0** | | | 11.6 | 152.2 |
| 294 | PUBCHEMCID66950 | Isocytosine | 9 | D | 1 | **32** | **0** | **0.3 M HCl** | | 10.2 | 111.1 |
| 323 | HMDB0004095 | 5-Methoxytryptamine | 9 | D | 2 | **33** | **50** | | | 13.3 | 190.2 |
| 533 | HMDB0003423 | D-Glutamine | 9 | D | 3 | **40** | **0** | | | 13.4 | 146.1 |
| 1093 | HMDB0000259 | Serotonin | 9 | D | 4 | **50** | **0** | | | 10.8 | 212.7 |
| 121 | HMDB0001370 | Diaminopimelic acid | 9 | D | 5 | **50** | **0** | | | 13 | 190.2 |
| 1216 | HMDB0000044 | Ascorbic acid | 9 | D | 6 | **36** | **0** | | | 11.4 | 176.1 |
| 170 | PUBCHEMCID11346228 | N4-Acetyl-2'-deoxycytidin | 9 | D | 7 | **50** | **44** | **0.4 M HCl** | | 11.5 | 269.3 |
| 1011 | HMDB0031081 | 2-Pentadecanone | 9 | D | 8 | | | | **Poor water solubility** | 11.1 | 226.4 |
| 216 | HMDB0030564 | trans-Piceid | 9 | D | 9 | **50** | **44** | | | 11.5 | 390.4 |
| 691 | HMDB0062472 | glycinamide | 9 | D | 10 | **32** | **0** | | | 11.3 | 110.5 |
| 25 | HMDB0003352 | Menthol | 9 | E | 1 | **27** | **25** | | | 10.8 | 156.3 |
| 842 | HMDB0037212 | Menthyl lactate | 9 | E | 2 | **52** | **30** | | | 10.3 | 228.3 |
| 321 | HMDB0002432 | Sumiki's acid | 9 | E | 3 | **36** | **33** | | | 10.8 | 142.1 |
| 707 | PUBCHEMCID92893 | D-histidine methyl ester dihydr | 9 | E | 4 | **50** | **0** | | | 10 | 242.1 |
| 57 | HMDB0000674 | PA(16:0/16:0) | 9 | E | 5 | | | | **Poor water solubility** | 5.2 | 670.9 |
| 835 | PUBCHEMCID440055 | 5-Methyl-2'-deoxycytidine | 9 | E | 6 | **50** | **0** | | | 10.9 | 241.3 |
| 750 | HMDB0001525 | Imidazole | 9 | E | 7 | | | | | 10.8 | 68.1 |
| 791 | HMDB0000472 | 5-Hydroxy-L-tryptophan | 9 | E | 8 | **48** | **48** | | | 10.2 | 220.2 |
| 885 | HMDB0029965 | Methyl beta-D-glucopyranoside | 9 | E | 9 | **50** | **0** | | | 11.2 | 194.2 |
| 36 | PUBCHEMCID86074 | Indoline-2-carboxylic acid | 9 | E | 10 | **29** | **75** | | | 11.5 | 163.2 |
| 331 | HMDB0030819 | Aesculetin | 9 | F | 1 | **47** | **60** | | **Poor water solubility** | 11.9 | 178.1 |
| 99 | HMDB0000678 | Isovalerylglycine | 9 | F | 2 | **37** | **0** | | | 10.7 | 159.2 |
| 1204 | PUBCHEMCID11790 | α-Naphthoflavone | 9 | F | 3 | | | | **Poor water solubility** | 10.7 | 272.3 |
| 659 | HMDB0000107 | Galactitol | 9 | F | 4 | **20** | **0** | | | 10.1 | 182.2 |
| 1088 | HMDB0034323 | S-Allylcysteine | 9 | F | 5 | **34** | **28** | **0.4 M HCl** | | 12.1 | 161.2 |
| 843 | HMDB0000696 | L-Methionine | 9 | F | 6 | **37** | **0** | **0.3 M HCl** | | 11 | 149.2 |
| 996 | HMDB0002340 | 2-Methylbenzoic acid | 9 | F | 7 | **32** | **30** | | | 10.8 | 136.2 |
| 869 | HMDB0000765 | Mannitol | 9 | F | 8 | **50** | **0** | | | 11 | 182.2 |
| 1249 | HMDB0000150 | Gluconolactone | 9 | F | 9 | **40** | **0** | | | 11.4 | 178.1 |
| 863 | HMDB0000691 | Malonic acid | 9 | F | 10 | **36** | **0** | | | 11.4 | 148.0 |
| | PUBCHEMCID99309 | N4-Acetylcytosine | 9 | G | 1 | **12** | **85** | | **Low water solubility** | 11.1 | 153.1 |
| 748 | HMDB0000157 | Hypoxanthine | 9 | G | 2 | **23** | **44** | **0.3 M NaOH** | | 10.3 | 136.1 |
| 724 | HMDB0005782 | Hesperetin | 9 | G | 3 | | | | **Poor water solubility** | 11.9 | 302.3 |
| 1244 | HMDB0000902 | NAD | 9 | G | 4 | **121** | **0** | | | 12.1 | 663.4 |
| 769 | PUBCHEMCID66085 | 8-Aminooctanoic acid | 9 | G | 5 | | | | **Low water solubility** | 11.1 | 159.2 |
| 133 | HMDB0000085 | Deoxyguanosine | 9 | G | 6 | **48** | **42** | **0.3 M HCl** | | 11.5 | 285.3 |
| 113 | PUBCHEMCID87839 | (S)-cis-Verbenol | 9 | G | 7 | | | | **Poor water solubility** | 12 | 152.2 |
| 972 | HMDB0000772 | Nonadecanoic acid | 9 | G | 8 | | | | **Poor water solubility** | 10.8 | 298.5 |
| 239 | HMDB0059712 | 3-Hydroxybenzyl alcohol | 9 | G | 9 | **33** | **33** | | | 10.1 | 124.1 |
| 400 | HMDB0004461 | Benzamide | 9 | G | 10 | **30.5** | **25** | | | 12.2 | 121.1 |
| 719 | HMDB0002259 | Heptadecanoic acid | 9 | H | 1 | | | | | 11.3 | 270.5 |
| 440 | HMDB0000921 | Cholestenone | 9 | H | 2 | | | | | 11.5 | 384.6 |
| 685 | HMDB0042061 | TG(14:0/14:0/14:0) | 9 | H | 3 | | | | | 12.6 | 723.2 |
| 764 | HMDB0000175 | Inosinic acid | 9 | H | 4 | **70** | **0** | | | 11.3 | 536.3 |
| 147 | HMDB0003224 | Deoxyribose | 9 | H | 5 | **34** | **0** | | | 12.2 | 134.1 |
| 28 | HMDB0014312 | (S)-Lipoic acid | 9 | H | 6 | **50** | **50** | **0.3 M NaOH** | | 12.4 | 206.3 |
| 1148 | PUBCHEMCID2756383 | Methyl 4-aminobutyrate HCl | 9 | H | 7 | **40** | **0** | | | 14.6 | 153.6 |
| 1183 | HMDB0000875 | Trigonelline | 9 | H | 8 | **35** | **0** | | | 10.6 | 173.6 |
| 827 | HMDB0000528 | 4,5-Dihydroorotic acid | 9 | H | 9 | **30** | **40** | | | 12 | 158.1 |
| 1166 | HMDB0001988 | 4-Hydroxycyclohexylcarboxylic | 9 | H | 10 | **35** | **33** | | | 10.6 | 144.2 |
| 204 | HMDB0062121 | Dihydroferulic acid | 9 | I | 1 | **44** | **50** | **0.3 M NaOH** | | 13.1 | 196.2 |
| 248 | HMDB0059969 | 3-Methoxyphenylacetic acid | 9 | I | 2 | **40** | **33** | | | 12 | 166.2 |
| 163 | HMDB0003903 | 2-Hydroxyethanesulfonate | 9 | I | 3 | **41** | **0** | | | 12.4 | 148.1 |
| 211 | HMDB0029492 | 3,3',7-Trihydroxy-4'-methoxyfla | 9 | I | 4 | **32** | **60** | | | 10.2 | 286.2 |
| 836 | HMDB0000172 | L-Isoleucine | 9 | I | 5 | **33** | **32** | **0.3 M HCl** | | 10.7 | 131.2 |
| 1113 | HMDB0001453 | Thiocyanate | 9 | I | 6 | **24.6** | **0** | | | 12.3 | 81.1 |
| 1237 | HMDB0000056 | beta-Alanine | 9 | I | 7 | **28** | **0** | | | 11.1 | 89.1 |
| 318 | HMDB0014389 | Mesalazine | 9 | I | 8 | **37** | **32** | | | 11.8 | 153.1 |
| 754 | HMDB0002285 | 2-Indolecarboxylic acid | 9 | I | 9 | **32** | **35** | **0.2 M NaOH** | | 11.3 | 161.2 |
| 1218 | HMDB0000876 | Vitamin D3 | 9 | I | 10 | | | | **Poor water solubility** | 12 | 384.6 |
| 478 | HMDB0031342 | Cyclohexanecarboxylic acid | 9 | J | 1 | **36** | **42** | | | 12.8 | 128.2 |
| 42 | HMDB0035140 | (S)-Abscisic acid | 9 | J | 2 | **28** | **0** | **0.4 M NaOH** | | 5 | 264.3 |
| 160 | HMDB0037115 | (±)-2-Hydroxy-4-(methylthio)bu | 9 | J | 3 | **44** | **40** | **0.4 M HCl** | | 10.6 | 338.5 |
| 1201 | HMDB0000285 | Uridine triphosphate | 9 | J | 4 | **62** | **0** | | | 12.3 | 550.1 |
| 299 | HMDB0000695 | Ketoleucine | 9 | J | 5 | **40** | **50** | **0.4 M HCl** | | 12.2 | 152.1 |
| 725 | HMDB0003265 | Hesperidin | 9 | J | 6 | **23** | **50** | **0.2 M NaOH** | | 11.4 | 610.6 |
| 152 | HMDB0031735 | Ethyl maltol | 9 | J | 7 | **32** | **35** | **0.1 M NaOH** | | 10.7 | 140.1 |
| 1071 | HMDB0035185 | Retinol acetate | 9 | J | 8 | **62** | **50%** | | **50% DMSO, 50% MeOH** | 12.2 | 328.5 |
| 97 | HMDB0011638 | Tetradecanol | 9 | J | 9 | | | | **Poor water solubility** | 10 | 214.4 |
| 322 | HMDB0001868 | 5-Methoxysalicylic acid | 9 | J | 10 | **34** | **31** | **0.2 M NaOH** | | 11 | 168.2 |

| | | | | | | | | | | | |
|---|---|---|---|---|---|---|---|---|---|---|---|
| 493 | HMDB0003072 | Quinic acid | 10 | A | 1 | **37** | **0** | | | 11.1 | 192.2 |
| 998 | HMDB0000223 | Oxalacetic acid | 10 | A | 2 | **28** | **25** | | | 11.1 | 132.1 |
| 1003 | PUBCHEMCID3766139 | Solvent Blue 35 | 10 | A | 3 | | | | **Dye** | 11.2 | 350.5 |
| 1154 | HMDB0000273 | Thymidine | 10 | A | 4 | **35** | **50** | | | 10.4 | 242.2 |
| 1164 | HMDB0031560 | 2-Methyl-2-pentenoic acid | 10 | A | 5 | **25** | **40** | | | 12.4 | 114.1 |
| 627 | HMDB0000576 | Monoethyl malonic acid | 10 | A | 6 | **36** | **0** | | | 10.9 | 170.2 |
| 664 | PUBCHEMCID71083 | D-Histidine | 10 | A | 7 | **28** | **0** | **0.2 M HCl** | | 12.6 | 155.2 |
| 199 | HMDB0034148 | 2-Tridecanone | 10 | A | 8 | | | | **Poor water solubility** | 11.2 | 198.3 |
| 566 | HMDB0000742 | Homocysteine | 10 | A | 9 | **23** | **0** | | | 13.9 | 135.2 |
| 689 | HMDB0031110 | Glycerol tritridecanoate | 10 | A | 10 | | | | **Poor water solubility** | 5.2 | 681.1 |
| 1221 | HMDB0032725 | Acetomenaphthone | 10 | B | 1 | **17** | **70** | | | 11.9 | 130.2 |
| 1058 | HMDB0004230 | Pyrrole-2-carboxylic acid | 10 | B | 2 | **119** | **0** | | **100% MeOH** | 11.9 | 111.1 |
| 607 | HMDB0030820 | Aesculin | 10 | B | 3 | **44** | **40** | | | 10.9 | 340.3 |
| 712 | HMDB0001842 | Guanidine | 10 | B | 4 | **100** | **0** | | **Chaotropic agent** | 10 | 95.5 |
| 221 | HMDB0000181 | L-Dopa | 10 | B | 5 | **32** | **30** | **0.38 M HCl** | | 10.2 | 197.2 |
| 89 | PUBCHEMCID81131 | 2-Acetylphenothiazine | 10 | B | 6 | **17** | **83** | | **Low water solubility** | 10.3 | 241.3 |
| 786 | HMDB0000452 | L-alpha-Aminobutyric acid | 10 | B | 7 | **26** | **0** | | | 10.3 | 103.1 |
| 44 | HMDB0034223 | 2,3,4,5,6-Penta-O-acetyl-D-gluc | 10 | B | 8 | | | | **Poor water solubility** | 11.3 | 390.3 |
| 130 | HMDB0035248 | 2,6-Dimethylpyrazine | 10 | B | 9 | **30** | **25** | | | 11.8 | 108.1 |
| 344 | HMDB0033585 | Acesulfame | 10 | B | 10 | **33** | **0** | | | 10 | 201.2 |
| 463 | HMDB0000634 | Citraconic acid | 10 | C | 1 | **34** | **0** | | | 10.2 | 131.0 |
| 164 | HMDB0000729 | alpha-Hydroxyisobutyric acid | 10 | C | 2 | **28** | **0** | | | 14 | 104.1 |
| 864 | HMDB0002928 | Maltitol | 10 | C | 3 | **60** | **0** | | | 12.1 | 344.3 |
| 406 | HMDB0040286 | Benzyl cinnamate | 10 | C | 4 | **50** | **60** | | **Poor water solubility** | 15.7 | 238.3 |
| 1234 | PUBCHEMCID11370 | 2-Methoxybenzoic acid | 10 | C | 5 | **36** | **50** | | | 14.3 | 152.2 |
| 226 | HMDB0000582 | 3,5-Diiodothyronine | 10 | C | 6 | **30** | **60** | | | 12.6 | 525.1 |
| 167 | HMDB0000005 | 2-Ketobutyric acid | 10 | C | 7 | **30** | **0** | | | 12 | 124.1 |
| 475 | HMDB0010720 | But-2-enoic acid | 10 | C | 8 | **27** | **25** | | | 10.8 | 86.9 |
| 794 | HMDB0000068 | Epinephrine | 10 | C | 9 | **49** | **0** | **0.5 M HCl** | | 10.8 | 183.2 |
| 317 | HMDB0001149 | 5-Aminolevulinic acid | 10 | C | 10 | **41** | **0** | | | 12.4 | 167.6 |
| 230 | HMDB0001885 | 3-Chlorotyrosine | 10 | D | 1 | **30** | **0** | **0.2 M HCl** | | 10 | 215.6 |
| 1175 | HMDB0000933 | Traumatic acid | 10 | D | 2 | | | | **Poor water solubility** | 7.3 | 228.3 |
| 1156 | HMDB0001878 | Thymol | 10 | D | 3 | **24.4** | **60** | | | 12.2 | 150.2 |
| 1027 | HMDB0036634 | Phlorizin | 10 | D | 4 | **47** | **45** | | | 10.4 | 436.4 |
| 581 | HMDB0000623 | Dodecanedioic acid | 10 | D | 5 | | | | **Poor water solubility** | 18.1 | 230.3 |
| 580 | HMDB0000944 | Behenic acid | 10 | D | 6 | | | | **Poor water solubility** | 13.3 | 340.6 |
| 857 | HMDB0000929 | L-Tryptophan | 10 | D | 7 | **38** | **30** | | | 12.4 | 204.2 |
| 818 | HMDB0000641 | L-Glutamine | 10 | D | 8 | **41** | **0** | **0.2 M HCl** | | 16.6 | 146.1 |
| 23 | HMDB0059839 | Camphene | 10 | D | 9 | | | | **Poor water solubility** | 12.9 | 136.2 |
| 456 | HMDB0006331 | cis,cis-Muconic acid | 10 | D | 10 | **16** | **40** | **0.2 M NaOH** | | 12.4 | 142.1 |
| 52 | HMDB0005393 | TG(18:0/18:0/18:0) | 10 | E | 1 | | | | | 10.7 | 891.5 |
| 702 | HMDB0028839 | Glycyl-Glutamine | 10 | E | 2 | **43** | **0** | **0.1 M HCl** | | 12.8 | 203.2 |
| 258 | HMDB0000752 | Methylglutaric acid | 10 | E | 3 | **34** | **0** | | | 10.2 | 146.1 |
| 148 | HMDB0001044 | 2'-Deoxyguanosine 5'-monopho | 10 | E | 4 | **52** | **0** | | | 10.7 | 391.2 |
| 416 | HMDB0002338 | Biochanin A | 10 | E | 5 | **20** | **68** | | | 10.2 | 284.3 |
| 1116 | HMDB0001256 | Spermine | 10 | E | 6 | **50** | **0** | | | 10.5 | 348.2 |
| 1083 | HMDB0003249 | Rutin | 10 | E | 7 | **58** | **46** | **0.2 M NaOH** | | 12.9 | 610.5 |
| 554 | PUBCHEMCID9894584 | Naringin dihydrochalcone | 10 | E | 8 | **55** | **50** | | | 11 | 582.6 |
| 503 | HMDB0001900 | Ribonolactone | 10 | E | 9 | **34** | **0** | | | 10.2 | 148.1 |
| 264 | HMDB0033128 | 4-(2-Furanyl)-3-buten-2-one | 10 | E | 10 | **28** | **50** | | | 11.5 | 136.2 |
| 949 | HMDB0002927 | Naringin | 10 | F | 1 | **60** | **50** | | | 11.9 | 580.5 |
| 810 | HMDB0002757 | Cysteic acid | 10 | F | 2 | **40** | **0** | | | 10.1 | 187.2 |
| 250 | HMDB0000022 | 3-Methoxytyramine | 10 | F | 3 | **40** | **32** | | | 12.5 | 203.7 |
| 992 | HMDB0000224 | O-Phosphoethanolamine | 10 | F | 4 | **40** | **0** | | | 12.2 | 141.1 |
| 988 | HMDB0000207 | Oleic acid | 10 | F | 5 | | | | **Poor water solubility** | 10 | 304.4 |
| 771 | HMDB0037316 | Isoliquiritigenin | 10 | F | 6 | **26** | **50** | | | 10.3 | 256.3 |
| 1018 | HMDB0000228 | Phenol | 10 | F | 7 | **25** | **0** | | | 24.7 | 94.1 |
| 761 | HMDB0000682 | Indoxyl sulfate | 10 | F | 8 | **68** | **0** | | | 20.5 | 251.3 |
| 195 | PUBCHEMCID74493 | D-(+)-Galactosamine HCl | 10 | F | 9 | **49** | **0** | | | 19.7 | 215.6 |
| 166 | HMDB0013751 | 2-Hydroxypyridine | 10 | F | 10 | **28** | **50** | | | 17 | 95.1 |
| 1070 | HMDB0001852 | all-trans-Retinoic acid | 10 | G | 1 | **101** | **100** | | **Poor water solubility** | 10.1 | 300.4 |
| 266 | HMDB0003681 | 4-Acetamidobutanoic acid | 10 | G | 2 | **30** | **30** | | | 10 | 145.2 |
| 954 | HMDB0030748 | Hesperetin 7-neohesperidoside | 10 | G | 3 | **51** | **60** | | | 18.4 | 610.6 |
| 256 | PUBCHEMCID10569 | Abietic Acid | 10 | G | 4 | | | | **Low water solubility** | 11.6 | 302.5 |
| 645 | HMDB0001248 | FAD | 10 | G | 5 | **83** | **0** | | | 5 | 829.5 |
| 833 | HMDB0000673 | Linoleic acid | 10 | G | 6 | **107** | **0** | | **30% chloroform** | 15 | 302.4 |
| 874 | HMDB0001389 | Melatonin | 10 | G | 7 | **35** | **33** | | | 10.4 | 232.3 |
| 694 | HMDB0000637 | Chenodeoxycholic acid glycine | 10 | G | 8 | **50** | **0** | | **80% MeOH** | 5 | 471.6 |
| 1104 | HMDB0033129 | 3-Acetyl-4-hydroxy-6-methyl-2l | 10 | G | 9 | **40** | **0** | | | 11.9 | 190.1 |
| 903 | HMDB0001522 | Methylguanidine | 10 | G | 10 | **41** | **0** | | | 12.4 | 109.6 |
| 425 | HMDB0001847 | Caffeine | 10 | H | 1 | **12.4** | | | **Poor water solubility** | 12.4 | 194.2 |
| 701 | HMDB0029419 | Glycylglycylglycine | 10 | H | 2 | **40** | **16** | **0.2 M HCl** | | 12.2 | 189.2 |
| 510 | HMDB0001310 | D-Alanine | 10 | H | 3 | **24** | **0** | | | 12 | 89.1 |
| 1199 | HMDB0000296 | Uridine | 10 | H | 4 | **50** | **0** | | | 10.1 | 244.2 |
| 788 | HMDB0000849 | Rhamnose | 10 | H | 5 | **36** | **0** | | | 11.6 | 182.2 |
| 735 | HMDB0000714 | Hippuric acid | 10 | H | 6 | **42** | **40** | | | 10.7 | 179.2 |
| 751 | HMDB0002024 | Imidazoleacetic acid | 10 | H | 7 | **40** | **0** | | | 12.4 | 162.6 |
| 203 | HMDB0011751 | 3-Methoxybenzenepropanoic a | 10 | H | 8 | **30** | **30** | **0.1 M NaOH** | | 10.2 | 180.2 |
| 684 | PUBCHEMCID69141 | 4-Hydroxyquinoline | 10 | H | 9 | **40** | **60** | | | 10.1 | 145.2 |
| 564 | HMDB0012127 | (R)-2-Benzylsuccinate | 10 | H | 10 | **40** | **30** | | | 13 | 208.2 |
| 194 | HMDB0002243 | Picolinic acid | 10 | I | 1 | **30** | **30** | | | 10 | 123.1 |
| 950 | PUBCHEMCID974 | 2-Amino-2-oxoacetic acid | 10 | I | 2 | **20** | **10** | **0.2 M NaOH** | | 10 | 89.1 |

| | | | | | | | | | | | |
|---|---|---|---|---|---|---|---|---|---|---|---|
| 146 | HMDB0062477 | 2-Deoxyglucose | 10 | I | 3 | **41** | **0** | | | 12.3 | 164.2 |
| 171 | HMDB0033830 | Cassiastearoptene | 10 | I | 4 | | | | **Poor water solubility** | 18.9 | 162.2 |
| 534 | PUBCHEMCID7017 | 2-Phenylphenol | 10 | I | 5 | **35** | **66** | | | 10.5 | 170.2 |
| 215 | HMDB0002511 | 3,4,5-Trimethoxycinnamic acid | 10 | I | 6 | | | | **Poor water solubility** | 12.1 | 238.2 |
| 383 | HMDB0002212 | Arachidic acid | 10 | I | 7 | | | | **Poor water solubility** | 11.2 | 312.5 |
| 500 | HMDB0000122 | D-Glucose | 10 | I | 8 | **184** | **0** | | | 14.8 | 180.2 |
| 506 | HMDB0011740 | Turanose | 10 | I | 9 | **50** | **0** | | | 10 | 342.3 |
| 556 | HMDB0029548 | Diosmin | 10 | I | 10 | **15** | **80** | | **Poor water solubility** | 11.7 | 608.5 |
| 727 | HMDB0000220 | Palmitic acid | 10 | J | 1 | | | | **Poor water solubility** | 10.9 | 256.4 |
| 783 | HMDB0000156 | Malic acid | 10 | J | 2 | **116** | **0** | | | 11.6 | 134.1 |
| 821 | HMDB0003466 | L-Gulonolactone | 10 | J | 3 | **38** | **0** | | | 11.3 | 178.1 |
| 828 | HMDB0002003 | Tetracosanoic acid | 10 | J | 4 | | | | **Poor water solubility** | 6 | 368.6 |
| 879 | HMDB0002994 | Erythritol | 10 | J | 5 | **38** | **0** | | | 19 | 122.1 |
| 933 | HMDB0006028 | N-Acetylasparagine | 10 | J | 6 | **37** | **0** | **0.1 M HCl** | | 11.1 | 174.2 |
| 1192 | HMDB0000286 | Uridine diphosphate glucose | 10 | J | 7 | **28** | **0** | | | 5.6 | 610.3 |
| 155 | HMDB0031849 | 2-Ethylpyrazine | 10 | J | 8 | **12** | **13** | | | 15.0 µL | 134.2 |
| 8 | HMDB0036559 | (-)-beta-Pinene | 10 | J | 9 | | | | **Poor water solubility** | 15.0 µL | 136.2 |
| 888 | HMDB0031343 | Methyl cyclohexanecarboxylate | 10 | J | 10 | **14** | **28** | | | 15.0 µL | 134.2 |
| 191 | HMDB0033944 | 2-Phenylethanol | 11 | A | 1 | **15** | **10** | | | 15.0 µL | 122.2 |
| 312 | HMDB0033154 | 5,6,7,8-Tetrahydroquinoxaline | 11 | A | 2 | **14** | **7** | | | 15.0 µL | 134.2 |
| 1005 | HMDB0003229 | Palmitoleic acid | 11 | A | 3 | | | | **Poor water solubility** | 5.0 µL | 254.4 |
| 674 | PUBCHEMCID92874 | (-)-Verbenone | 11 | A | 4 | | | | **Poor water solubility** | 15.0 µL | 150.2 |
| 185 | HMDB0031266 | 2-Nonanone | 11 | A | 5 | | | | **Low water solubility** | 15.0 µL | 142.2 |
| 1141 | HMDB0035833 | p-Menth-1-en-4-ol | 11 | A | 6 | **11** | **12** | | | 15.0 µL | 154.3 |
| 76 | HMDB0011624 | Decyl alcohol | 11 | A | 7 | | | | **Poor water solubility** | 15.0 µL | 158.3 |
| 296 | HMDB0062481 | iso-nLc8Cer | 11 | A | 8 | | | | **Low water solubility** | 15.0 µL | 135.2 |
| 27 | HMDB0036197 | (-)-alpha-Bisabolol | 11 | A | 9 | | | | **Poor water solubility** | 15.0 µL | 222.4 |
| 629 | HMDB0031643 | Ethyl pyruvate | 11 | A | 10 | **13** | **13** | | | 15.0 µL | 116.1 |
| 723 | HMDB0000666 | Heptanoic acid | 11 | B | 1 | **15** | **17** | | | 15.0 µL | 130.2 |
| 95 | HMDB0013036 | 1-Pentanol | 11 | B | 2 | **10** | **13** | | | 15.0 µL | 88.2 |
| 549 | HMDB0041606 | Dimethyl adipate | 11 | B | 3 | | | | **Poor water solubility** | 15.0 µL | 174.2 |
| 200 | HMDB0033713 | 2-Undecanone | 11 | B | 4 | | | | **Poor water solubility** | 15.0 µL | 170.3 |
| 1114 | PUBCHEMCID3406 | 4-Methylpyrazole | 11 | B | 5 | **10** | **10** | | | 15.0 µL | 82.1 |
| 380 | HMDB0003012 | Aniline | 11 | B | 6 | **10** | **10** | | | 15.0 µL | 93.1 |
| 871 | HMDB0059855 | m-Dichlorobenzene | 11 | B | 7 | **7** | **60** | | | 15.0 µL | 147.0 |
| 913 | HMDB0001888 | N,N-Dimethylformamide | 11 | B | 8 | **10** | **8** | | | 15.0 µL | 73.1 |
| 625 | HMDB0040193 | Ethyl nonanoate | 11 | B | 9 | | | | **Low water solubility** | 15.0 µL | 186.3 |
| 198 | HMDB0002039 | 2-Pyrrolidinone | 11 | B | 10 | **10** | **9** | | | 15.0 µL | 85.1 |
| 39 | HMDB0035842 | (S)-Citronellal | 11 | C | 1 | | | | **Poor water solubility** | 15.0 µL | 154.3 |
| 40 | HMDB0004321 | (-)-Limonene | 11 | C | 2 | | | | **Poor water solubility** | 15.0 µL | 136.2 |
| 635 | HMDB0032260 | Ethylene glycol distearate | 11 | C | 3 | | | | **Poor water solubility** | 15.0 µL | 62.1 |
| 890 | HMDB0031018 | Methyl dodecanoate | 11 | C | 4 | | | | **Poor water solubility** | 15.0 µL | 214.3 |
| 896 | HMDB0031264 | Methyl nonanoate | 11 | C | 5 | | | | **Poor water solubility** | 15.0 µL | 172.3 |
| 197 | HMDB0006804 | Propynoic acid | 11 | C | 6 | **8.5** | **8** | | | 15.0 µL | 70.1 |
| 899 | HMDB0030469 | Methyl tetradecanoate | 11 | C | 7 | | | | **Poor water solubility** | 15.0 µL | 242.4 |
| 50 | HMDB0005453 | TG(18:1(9Z)/18:1(9Z)/18:1(9Z)) | 11 | C | 8 | | | | **Poor water solubility** | 15.0 µL | 885.4 |
| 634 | HMDB0029552 | Ethyl undecanoate | 11 | C | 9 | | | | **Poor water solubility** | 15.0 µL | 214.3 |
| 1009 | HMDB0005805 | p-Cymene | 11 | C | 10 | | | | **Poor water solubility** | 15.0 µL | 134.2 |
| 405 | HMDB0003119 | Benzyl alcohol | 11 | D | 1 | **11** | **10** | | | 15.0 µL | 108.4 |
| 536 | HMDB0041220 | Dibutyl decanedioate | 11 | D | 2 | | | | **Poor water solubility** | 15.0 µL | 314.5 |
| 624 | HMDB0040195 | Ethyl octanoate | 11 | D | 3 | | | | **Poor water solubility** | 15.0 µL | 172.3 |
| 1173 | HMDB0035662 | Nerolidol | 11 | D | 4 | **17.6** | **40** | | | 15.0 µL | 222.4 |
| 1182 | HMDB0034263 | Triethyl citrate | 11 | D | 5 | | | | **Poor water solubility** | 15.0 µL | 276.3 |
| 233 | HMDB0003243 | Acetoin | 11 | D | 6 | **10** | **10** | | | 15.0 µL | 88.1 |
| 1056 | HMDB0003361 | Pyrimidine | 11 | D | 7 | **10** | **10** | | | 15.0 µL | 80.1 |
| 49 | HMDB0011187 | TG(8:0/8:0/8:0) | 11 | D | 8 | | | | **Poor water solubility** | 5.1 | 470.7 |
| 135 | PUBCHEMCID75569 | 2-Picolinic acid methyl ester | 11 | D | 9 | **15** | **13** | | | 15.0 µL | 137.1 |
| 51 | HMDB0005432 | TG(16:1(9Z)/16:1(9Z)/16:1(9Z)) | 11 | D | 10 | | | | **Poor water solubility** | 5.0 µL | 801.3 |
| 47 | HMDB0031125 | Glycerol trihexanoate | 11 | E | 1 | | | | **Poor water solubility** | 5.0 µL | 386.5 |
| 458 | HMDB0040213 | cis-3-Hexenyl isobutyrate | 11 | E | 2 | | | | **Poor water solubility** | 15.0 µL | 170.3 |
| 41 | PUBCHEMCID638500 | Farnesyl acetate (mixture of iso | 11 | E | 3 | | | | **Poor water solubility** | 15.0 µL | 264.4 |
| 908 | HMDB0000858 | Monomethyl glutaric acid | 11 | E | 4 | **15** | **13** | | | 15.0 µL | 146.1 |
| 70 | HMDB0031016 | 10-Undecen-1-ol | 11 | E | 5 | | | | **Poor water solubility** | 15.0 µL | 170.3 |
| 98 | HMDB0013113 | 1-Undecanol | 11 | E | 6 | | | | **Poor water solubility** | 15.0 µL | 172.3 |
| 889 | HMDB0033848 | Methyl decanoate | 11 | E | 7 | | | | **Poor water solubility** | 15.0 µL | 186.3 |
| 720 | PUBCHEMCID15607 | Methyl undecanoate | 11 | E | 8 | | | | **Poor water solubility** | 15.0 µL | 200.3 |
| 1107 | PUBCHEMCID31374 | N,N-Dimethylacetamide | 11 | E | 9 | **10** | **0** | | | 15.0 µL | 87.1 |
| 351 | PUBCHEMCID12965 | N,N-Dimethylpropionamide | 11 | E | 10 | **12** | **0** | | | 15.0 µL | 101.2 |
| 91 | PUBCHEMCID7515 | N-Methylaniline | 11 | F | 1 | **10** | **33** | | **Low water solubility** | 15.0 µL | 107.2 |
| 122 | HMDB0032782 | 2,6-Diethylaniline | 11 | F | 2 | | | | **Poor water solubility** | 15.0 µL | 149.2 |
| 129 | HMDB0060677 | 2,6-Dimethylaniline | 11 | F | 3 | | | | **Poor water solubility** | 15.0 µL | 121.2 |
| 1236 | HMDB0004043 | alpha-Terpineol | 11 | F | 4 | | | | **Poor water solubility** | 15.0 µL | 154.3 |
| 1171 | HMDB0003441 | Cinnamaldehyde | 11 | F | 5 | **12** | **10** | | **Low water solubility** | 15.0 µL | 132.2 |
| 653 | HMDB0001536 | Formamide | 11 | F | 6 | **15** | **0** | | **Toxic** | 15.0 µL | 45.0 |
| 477 | HMDB0031403 | Cyclohexaneacetic acid | 11 | F | 7 | **15** | **13** | | | 15.0 µL | 142.2 |
| 295 | HMDB0032025 | 4'-Isopropylacetophenone | 11 | F | 8 | | | | **Poor water solubility** | 15.0 µL | 162.2 |
| 844 | HMDB0001934 | Nicotine | 11 | F | 9 | **18** | **14** | | | 15.0 µL | 162.2 |
| 1169 | HMDB0030837 | Anethole | 11 | F | 10 | **5** | **60** | | | 15.0 µL | 148.2 |
| 169 | HMDB0032569 | 2'-Methoxyacetophenone | 11 | G | 1 | **16** | **50** | | | 15.0 µL | 150.2 |
| 161 | HMDB0032568 | 2'-Hydroxyacetophenone | 11 | G | 2 | **15** | **35** | | | 15.0 µL | 136.2 |
| 1142 | HMDB0036994 | Terpinolene | 11 | G | 3 | | | | **Poor water solubility** | 15.0 µL | 136.2 |
| 901 | HMDB0031207 | Methyl pentanoate | 11 | G | 4 | | | | **Low water solubility** | 15.0 µL | 116.2 |

| | | | | | | | | | | |
|---|---|---|---|---|---|---|---|---|---|---|
| 571 | HMDB0006024 | Mevalonolactone | 11 | G | 5 | **10** | **20** | | 5.0 µL | 130.1 |
| 271 | HMDB0032608 | 4'-Methylacetophenone | 11 | G | 6 | | | **Poor water solubility** | 15.0 µL | 134.2 |
| 114 | HMDB0032140 | 2',4'-Dimethylacetophenone | 11 | G | 7 | | | **Poor water solubility** | 15.0 µL | 148.2 |
| 236 | PUBCHEMCID8454 | 1-Methylpyrrolidine | 11 | G | 8 | **10** | **15** | | 15.0 µL | 85.1 |
| 919 | PUBCHEMCID111470 | 8-Methyl nonanoic acid | 11 | G | 9 | | | **Poor water solubility** | 5.0 µL | 172.3 |
| 732 | HMDB0040431 | Hexyl benzoate | 11 | G | 10 | | | **Poor water solubility** | 15.0 µL | 206.3 |
| 180 | HMDB0031158 | 2-Methyl-4-pentenoic acid | 11 | H | 1 | **12.5** | **0** | | 15.0 µL | 114.1 |
| 893 | HMDB0036583 | Methyl jasmonate | 11 | H | 2 | | | **Poor water solubility** | 15.0 µL | 224.3 |
| 176 | HMDB0031587 | 2-Methylheptanoic acid | 11 | H | 3 | **15** | **13** | | 15.0 µL | 144.2 |
| 18 | HMDB0035093 | D-Citronellol | 11 | H | 4 | | | **Poor water solubility** | 5.0 µL | 156.3 |
| 35 | HMDB0035604 | Pulegone | 11 | H | 5 | | | **Poor water solubility** | 15.0 µL | 152.2 |
| 260 | HMDB0033774 | 3-Methylpentanoic acid | 11 | H | 6 | **12** | **13** | | 15.0 µL | 116.2 |
| 886 | HMDB0033890 | Methyl butyrate | 11 | H | 7 | | | **Poor water solubility** | 15.0 µL | 102.1 |
| 522 | HMDB0037116 | delta-Decalactone | 11 | H | 8 | | | **Poor water solubility** | 15.0 µL | 170.3 |
| 1160 | HMDB0004305 | Farnesol | 11 | H | 9 | | | **Poor water solubility** | 15.0 µL | 222.4 |
| 209 | HMDB0001527 | 3-Methylthiopropionic acid | 11 | H | 10 | **13** | **12** | | 15.0 µL | 120.2 |
| 1248 | HMDB0005806 | gamma-Terpinene | 11 | I | 1 | | | **Poor water solubility** | 15.0 µL | 136.2 |
| 768 | HMDB0001873 | Isobutyric acid | 11 | I | 2 | **10** | **0** | | 15.0 µL | 88.1 |
| 830 | HMDB0036100 | 3,7-Dimethyl-1,6-octadien-3-ol | 11 | I | 3 | | | **Poor water solubility** | 15.0 µL | 154.3 |
| 612 | PUBCHEMCID69286 | Glycerol Triheptanoate | 11 | I | 4 | | | **Poor water solubility** | 5.0 µL | 428.6 |
| 579 | PUBCHEMCID82229 | (-)-Fenchone | 11 | I | 5 | | | **Poor water solubility** | 15.0 µL | 152.2 |
| 774 | HMDB0000718 | Isovaleric acid | 11 | I | 6 | **11** | **0** | | 15.0 µL | 102.1 |
| | HMDB0031216 | Ethyl acetoacetate | 11 | I | 7 | | | **Poor water solubility** | 15.0 µL | 130.1 |
| 348 | HMDB0033910 | Acetophenone | 11 | I | 8 | **12** | **41** | | 15.0 µL | 120.2 |
| 272 | HMDB0034121 | 1-Methoxy-4-(2-propenyl)benz | 11 | I | 9 | | | **Poor water solubility** | 15.0 µL | 148.2 |
| | HMDB0034244 | Isoquinoline | 11 | I | 10 | | | **Poor water solubility** | 15.0 µL | 129.2 |
| 34 | HMDB0003375 | (+)-Limonene | 11 | J | 1 | | | **Poor water solubility** | 15.0 µL | 136.2 |
| 501 | HMDB0001051 | Glyceraldehyde | 11 | J | 2 | **10** | **0** | | 6.8 | 90.1 |
| 269 | HMDB0031306 | 3-Hydroxy-4,5-dimethyl-2(5H)-f | 11 | J | 3 | **13** | **12** | | 15.0 µL | 128.1 |
| 30 | HMDB0000357 | beta-Hydroxybutyric acid | 11 | J | 4 | **11** | **0** | | 15.0 µL | 104.1 |
| 1163 | HMDB0031496 | 2-Hexenal | 11 | J | 5 | **10** | **12** | | 15.0 µL | 98.1 |
| 278 | HMDB0040175 | 4-Ethyl-2-methoxyphenol | 11 | J | 6 | **15** | **40** | | 15.0 µL | 152.2 |
| 31 | HMDB0035619 | alpha-Carene | 11 | J | 7 | | | **Poor water solubility** | 15.0 µL | 136.2 |
| 1206 | HMDB0031208 | 1-Phenyl-1-pentanone | 11 | J | 8 | | | **Low water solubility** | 15.0 µL | 162.2 |
| 832 | HMDB0039522 | Linalyl acetate | 11 | J | 9 | | | **Poor water solubility** | 15.0 µL | 196.3 |
| 640 | HMDB0005809 | Eugenol | 11 | J | 10 | | | **Poor water solubility** | 15.0 µL | 164.2 |
| 639 | HMDB0004472 | Eucalyptol | 12 | A | 1 | | | **Poor water solubility** | 15.0 µL | 154.3 |
| 17 | HMDB0006525 | (+)-alpha-Pinene | 12 | A | 2 | | | **Poor water solubility** | 15.0 µL | 136.2 |
| 688 | HMDB0032857 | Glycerol tripropanoate | 12 | A | 3 | | | **Poor water solubility** | 15.0 µL | 260.3 |
| 841 | HMDB0035162 | (-)-Menthone | 12 | A | 4 | | | **Poor water solubility** | 15.0 µL | 154.3 |
| 680 | HMDB0029592 | Triacetin | 12 | A | 5 | | | **Poor water solubility** | 15.0 µL | 218.2 |
| 1242 | HMDB0036565 | beta-Ionone | 12 | A | 6 | **10** | **65** | | 15.0 µL | 192.3 |
| 620 | HMDB0029811 | Ethyl hexadecanoate | 12 | A | 7 | | | **Poor water solubility** | 15.0 µL | 284.5 |
| 1228 | HMDB0036027 | alpha-Damascone | 12 | A | 8 | | | **Poor water solubility** | 15.0 µL | 192.3 |
| 431 | HMDB0035770 | 5-Isopropyl-2-methylphenol | 12 | A | 9 | **16** | **50** | | 15.0 µL | 150.2 |
| 1240 | HMDB0036792 | beta-Caryophyllene | 12 | A | 10 | | | **Poor water solubility** | 15.0 µL | 204.4 |
| 308 | HMDB0032625 | 4-Propylphenol | 12 | B | 1 | **12** | **40** | | 15.0 µL | 136.2 |
| 587 | PUBCHEMCID71317439 | Geranyl linalool (mixture of iso | 12 | B | 2 | | | **Poor water solubility** | 15.0 µL | 290.5 |
| 303 | HMDB0032136 | 2-Methoxy-4-methylphenol | 12 | B | 3 | **15** | **14** | | 15.0 µL | 138.2 |
| 80 | HMDB0031479 | 1-Heptanol | 12 | B | 4 | **11** | **11** | | 15.0 µL | 116.2 |
| 1118 | HMDB0000256 | Squalene | 12 | B | 5 | | | **Poor water solubility** | 15.0 µL | 410.7 |
| 623 | HMDB0040433 | Ethyl levulinate | 12 | B | 6 | **16** | **10** | | 15.0 µL | 144.2 |
| 1247 | HMDB0003843 | Gamma-Caprolactone | 12 | B | 7 | **12** | **12** | | 15.0 µL | 114.1 |
| 173 | HMDB0031527 | (S)-2-Methyl-1-butanol | 12 | B | 8 | **10** | **0** | | 15.0 µL | 88.2 |
| 407 | HMDB0041485 | Benzyl formate | 12 | B | 9 | **12.4** | **40** | | 15.0 µL | 136.2 |
| 1227 | HMDB0031313 | 2-Pentyl-3-phenyl-2-propenal | 12 | B | 10 | | | **Poor water solubility** | 15.0 µL | 202.3 |
| 302 | HMDB0031540 | 4-Methylcyclohexanone | 12 | C | 1 | **12** | **13** | | 15.0 µL | 112.2 |
| 734 | HMDB0031736 | 2-Hexyl-3-phenyl-2-propenal | 12 | C | 2 | | | **Poor water solubility** | 15.0 µL | 216.3 |
| 544 | HMDB0036626 | 3,4-Dihydro-2H-1-benzopyran-2 | 12 | C | 3 | **15** | **13** | | 15.0 µL | 148.2 |
| 617 | HMDB0039581 | Ethyl crotonate | 12 | C | 4 | **12** | **13** | | 15.0 µL | 114.1 |
| 340 | PUBCHEMCID11040 | Methyl (R)-(+)-lactate | 12 | C | 5 | **12** | **0** | | 15.0 µL | 104.1 |
| 733 | HMDB0034606 | Hexyl propionate | 12 | C | 6 | | | **Poor water solubility** | 15.0 µL | 158.2 |
| 540 | HMDB0033584 | Diethyl tartrate | 12 | C | 7 | **20** | **16** | | 15.0 µL | 206.2 |
| 1205 | HMDB0000892 | Valeric acid | 12 | C | 8 | **12** | **0** | | 15.0 µL | 102.1 |
| 731 | HMDB0000535 | Caproic acid | 12 | C | 9 | **12** | **0** | | 15.0 µL | 116.2 |
| 1024 | HMDB0033716 | 3-Phenylpropanal | 12 | C | 10 | **8** | **60** | | 15.0 µL | 134.2 |
| 497 | PUBCHEMCID7344 | (+)-Ethyl D-lactate | 12 | D | 1 | **12** | **0** | | 15.0 µL | 118.1 |
| 62 | HMDB0001881 | Propylene glycol | 12 | D | 2 | **10** | **0** | | 15.0 µL | 76.1 |
| 551 | HMDB0033837 | Dimethyl succinate | 12 | D | 3 | **15** | **0** | | 15.0 µL | 146.1 |
| 224 | HMDB0031492 | 3,4-Hexanedione | 12 | D | 4 | **9** | **50** | | 15.0 µL | 114.1 |
| 384 | PUBCHEMCID134442 | 3-Methoxypropanoic acid | 12 | D | 5 | **12** | **11** | | 15.0 µL | 104.1 |
| 907 | HMDB0059722 | Mono-methyl-adipate | 12 | D | 6 | **17** | **15** | | 15.0 µL | 160.2 |
| 921 | PUBCHEMCID9855465 | 2-Methoxy-2-methylpropanoic | 12 | D | 7 | **12** | **12** | | 15.0 µL | 118.1 |
| 263 | HMDB0002523 | Oxolan-3-one | 12 | D | 8 | **10** | **9** | | 15.0 µL | 86.1 |
| 172 | HMDB0031178 | 2-Methyltetrahydrofuran-3-one | 12 | D | 9 | **11** | **11** | | 15.0 µL | 100.1 |
| 767 | HMDB0031528 | Isopentyl acetate | 12 | D | 10 | **14** | **16** | | 15.0 µL | 130.2 |
| 781 | HMDB0035089 | (R)-Carvone | 12 | E | 1 | | | **Poor water solubility** | 15.0 µL | 150.2 |
| 630 | HMDB0029817 | Ethyl salicylate | 12 | E | 2 | **6.6** | **50** | | 15.0 µL | 166.2 |
| 982 | HMDB0031294 | 2-Octanone | 12 | E | 3 | | | **Poor water solubility** | 15.0 µL | 128.2 |
| 1238 | PUBCHEMCID26722 | Pentadecanoic Acid methyl est | 12 | E | 4 | | | **Poor water solubility** | 15.0 µL | 256.4 |
| 141 | PUBCHEMCID5362793 | Methyl linoelaidate | 12 | E | 5 | | | **Poor water solubility** | 5.0 µL | 294.5 |
| 154 | HMDB0040587 | 2-Ethylfuran | 12 | E | 6 | **8** | **40** | | 15.0 µL | 96.1 |

| | | | | | | | | | | |
|---|---|---|---|---|---|---|---|---|---|---|
| 73 | HMDB0031404 | Cyclohexylamine | 12 | E | 7 | | | **Poor water solubility** | 15.0 µL | 99.2 |
| 770 | HMDB0005802 | Isoeugenol | 12 | E | 8 | **15** | **50** | | 15.0 µL | 164.2 |
| 1243 | HMDB0038169 | beta-Myrcene | 12 | E | 9 | | | **Poor water solubility** | 15.0 µL | 136.2 |
| 457 | HMDB0030003 | 3-Hexen-1-ol | 12 | E | 10 | **10** | **12** | | 15.0 µL | 100.2 |
| 174 | HMDB0006006 | Isobutanol | 12 | F | 1 | **8** | **0** | | 15.0 µL | 74.1 |
| 207 | PUBCHEMCID228987 | 3-(4-tert-Butylphenyl)isobutyra | 12 | F | 2 | | | **Poor water solubility** | 15.0 µL | 204.3 |
| 831 | HMDB0040727 | (±)-trans-Linalyl oxide | 12 | F | 3 | | | **Poor water solubility** | 15.0 µL | 170.3 |
| 631 | HMDB0040463 | Ethyl sorbate | 12 | F | 4 | | | **Poor water solubility** | 15.0 µL | 140.2 |
| 974 | HMDB0000847 | Pelargonic acid | 12 | F | 5 | | | **Low water solubility** | 15.0 µL | 158.2 |
| 108 | HMDB0032971 | 2,3-Dimethylpyrazine | 12 | F | 6 | **11** | **11** | | 15.0 µL | 108.1 |
| 1051 | PUBCHEMCID10453 | Isophytol | 12 | F | 7 | | | **Poor water solubility** | 15.0 µL | 296.5 |
| 106 | HMDB0041253 | 2,3-Diethylpyrazine | 12 | F | 8 | | | **Poor water solubility** | 15.0 µL | 136.2 |
| 613 | HMDB0040413 | Ethyl 3-(methylthio)propanoate | 12 | F | 9 | **15** | **15** | | 15.0 µL | 148.2 |
| 229 | HMDB0002019 | Phytol | 12 | F | 10 | | | **Poor water solubility** | 15.0 µL | 296.5 |
| 64 | HMDB0000002 | 1,3-Diaminopropane | 12 | G | 1 | **9** | **0** | | 15.0 µL | 74.1 |
| 626 | HMDB0040297 | Ethyl pentanoate | 12 | G | 2 | **13** | **15** | | 15.0 µL | 130.2 |
| 973 | PUBCHEMCID15608 | Tridecanoic Acid methyl ester | 12 | G | 3 | | | **Poor water solubility** | 15.0 µL | 228.4 |
| 1151 | HMDB0029713 | Thiazole | 12 | G | 4 | **10** | **0** | | 15.0 µL | 85.1 |
| 327 | HMDB0033178 | 5-Methylquinoxaline | 12 | G | 5 | | | **Poor water solubility** | 15.0 µL | 144.2 |
| 904 | HMDB0032306 | Diethyl oxalpropionate | 12 | G | 6 | | | **Low water solubility** | 15.0 µL | 202.2 |
| 547 | HMDB0041613 | Bis(1-methylethyl) hexanedioat | 12 | G | 7 | | | **Poor water solubility** | 15.0 µL | 230.3 |
| 1019 | HMDB0040733 | Phenyl acetate | 12 | G | 8 | **14** | **20** | | 15.0 µL | 136.2 |
| 300 | HMDB0032076 | 1-Methoxy-4-methylbenzene | 12 | G | 9 | | | **Poor water solubility** | 15.0 µL | 122.2 |
| 335 | HMDB0033115 | 6-Methylquinoline | 12 | G | 10 | | | **Poor water solubility** | 15.0 µL | 143.2 |
| 553 | HMDB0013780 | Dimethyl trisulfide | 12 | H | 1 | **13** | **11** | | 15.0 µL | 126.3 |
| 115 | HMDB0032142 | 2,4-Dimethylbenzaldehyde | 12 | H | 2 | **9** | **47** | | 15.0 µL | 134.2 |
| 381 | HMDB0033895 | Anisole | 12 | H | 3 | **12** | **40** | | 15.0 µL | 108.1 |
| 558 | HMDB0041614 | Dipropyl hexanedioate | 12 | H | 4 | | | **Poor water solubility** | 15.0 µL | 230.3 |
| 270 | HMDB0032976 | 4,5-Dimethylthiazole | 12 | H | 5 | **12** | **11** | **Poor water solubility** | 15.0 µL | 113.2 |
| 310 | HMDB0031866 | 2-Methyl-4-phenyl-2-butanol | 12 | H | 6 | **13** | **40** | | 15.0 µL | 150.2 |
| 1250 | HMDB0036784 | 6-Hexyltetrahydro-2H-pyran-2- | 12 | H | 7 | | | **Poor water solubility** | 15.0 µL | 184.3 |
| 628 | HMDB0030058 | Ethyl propionate | 12 | H | 8 | **11** | **12** | | 15.0 µL | 102.1 |
| 542 | HMDB0033838 | Diethyl succinate | 12 | H | 9 | **18** | **18** | | 15.0 µL | 174.2 |
| 75 | HMDB0004327 | 1-Butanol | 12 | H | 10 | **9** | **0** | | 15.0 µL | 74.1 |
| 480 | HMDB0003315 | Cyclohexanone | 12 | I | 1 | **10** | **10** | | 15.0 µL | 98.2 |
| 251 | HMDB0006007 | Isopentanol | 12 | I | 2 | **10** | **0** | | 15.0 µL | 88.2 |
| 5 | HMDB0036078 | (-)-Isopulegol | 12 | I | 3 | **15** | **16** | **Low water solubility** | 15.0 µL | 154.3 |
| 622 | HMDB0033788 | Ethyl dodecanoate | 12 | I | 4 | | | **Poor water solubility** | 15.0 µL | 228.4 |
| 408 | HMDB0033871 | Benzylamine | 12 | I | 5 | **11** | **12** | | 15.0 µL | 107.2 |
| 1059 | HMDB0001167 | Pyruvaldehyde | 12 | I | 6 | **10** | **0** | | 15.0 µL | 72.1 |
| 1220 | HMDB0003555 | Vitamin K1 | 12 | I | 7 | | | **Poor water solubility** | 12.9 | 450.7 |
| 1145 | HMDB0035246 | 3,7-Dimethyl-3-octanol | 12 | I | 8 | | | **Poor water solubility** | 15.0 µL | 158.3 |
| 9 | HMDB0004487 | (S)-Carvone | 12 | I | 9 | | | **Poor water solubility** | 15.0 µL | 150.2 |
| 279 | HMDB0036136 | 4-Ethyloctanoic acid | 12 | I | 10 | **12** | **15** | | 15.0 µL | 172.3 |
| 520 | HMDB0011623 | Decanal | 12 | J | 1 | | | **Poor water solubility** | 15.0 µL | 156.3 |
| 252 | HMDB0030124 | Prenol | 12 | J | 2 | **10** | **12** | | 15.0 µL | 86.1 |
| 79 | HMDB0030204 | 1-(2-Furanylmethyl)-1H-pyrrole | 12 | J | 3 | | | **Poor water solubility** | 15.0 µL | 147.2 |
| 984 | HMDB0001140 | Octanal | 12 | J | 4 | | | **Poor water solubility** | 15.0 µL | 128.2 |
| 92 | HMDB0059835 | Nonanal | 12 | J | 5 | | | **Poor water solubility** | 15.0 µL | 142.2 |
| 84 | HMDB0032074 | 1-Methoxy-2-methylbenzene | 12 | J | 6 | | | **Poor water solubility** | 15.0 µL | 122.2 |
| 189 | HMDB0031628 | 2-Phenyl-1-propanol | 12 | J | 7 | **12** | **18** | | 15.0 µL | 136.2 |
| 1020 | HMDB0006236 | Phenylacetaldehyde | 12 | J | 8 | **6.5** | **70** | | 15.0 µL | 120.2 |
| 1115 | HMDB0001257 | Spermidine | 12 | J | 9 | **15** | **0** | | 15.0 µL | 145.3 |
| 210 | HMDB0041491 | 3-(Methylthio)butanal | 12 | J | 10 | **6.6** | **80** | | 15.0 µL | 118.2 |
| 37 | HMDB0002017 | 1-Phenylethylamine | 13 | A | 1 | **13** | **14** | | 15.0 µL | 121.2 |
| 618 | HMDB0030998 | Ethyl decanoate | 13 | A | 2 | | | **Poor water solubility** | 15.0 µL | 200.3 |
| 619 | HMDB0000798 | Oenanthic ether | 13 | A | 3 | **16** | **15** | | 15.0 µL | 158.2 |
| 82 | HMDB0012971 | 1-Hexanol | 13 | A | 4 | **11** | **0** | | 15.0 µL | 102.2 |
| 616 | HMDB0033889 | Ethyl butyrate | 13 | A | 5 | **12** | **13** | | 15.0 µL | 116.2 |
| 957 | HMDB0005812 | Geraniol | 13 | A | 6 | | | **Poor water solubility** | 15.0 µL | 154.3 |
| 915 | HMDB0001020 | N,N-Dimethylaniline | 13 | A | 7 | | | **Poor water solubility** | 15.0 µL | 121.2 |
| 891 | HMDB0031478 | Methyl heptanoate | 13 | A | 8 | | | **Poor water solubility** | 15.0 µL | 144.2 |
| 518 | HMDB0031409 | 2-Decanone | 13 | A | 9 | | | **Poor water solubility** | 15.0 µL | 156.3 |
| 159 | HMDB0005842 | 2-Oxohexane | 13 | A | 10 | **11** | **13** | | 15.0 µL | 100.2 |
| 188 | HMDB0034235 | 2-Pentanone | 13 | B | 1 | **10** | **12** | | 15.0 µL | 86.1 |
| 158 | HMDB0003671 | 2-Heptanone | 13 | B | 2 | **12** | **11** | | 15.0 µL | 114.2 |
| 663 | HMDB0035157 | Geranyl acetate | 13 | B | 3 | **14** | **50** | | 15.0 µL | 196.3 |
| 142 | HMDB0011469 | 2-Butanol | 13 | B | 4 | **8** | **0** | | 15.0 µL | 74.1 |
| 621 | HMDB0040209 | Ethyl hexanoate | 13 | B | 5 | **15** | **15** | | 15.0 µL | 144.2 |
| 897 | HMDB0031291 | Methyl caprylate | 13 | B | 6 | **14** | **16** | **Low water solubility** | 15.0 µL | 158.2 |
| 892 | HMDB0035238 | Methyl hexanoate | 13 | B | 7 | **14** | **16** | | 15.0 µL | 130.2 |
| 151 | HMDB0031019 | 2-Dodecanone | 13 | B | 8 | | | **Poor water solubility** | 15.0 µL | 184.3 |
| 541 | HMDB0029573 | Diethyl malonate | 13 | B | 9 | **17** | **16** | **Soluble in acetone** | 15.0 µL | 160.2 |
| 1123 | HMDB0034240 | Styrene | 13 | B | 10 | | | **Poor water solubility** | 15.0 µL | 104.2 |
| 208 | PUBCHEMCID701 | Ethyl 2-methylacetoacetate | 13 | C | 1 | **15** | **15** | **Low water solubility** | 15.0 µL | 144.2 |
| 87 | HMDB0032860 | 1-Methylnaphthalene | 13 | C | 2 | | | **Poor water solubility** | 15.0 µL | 142.2 |
| 550 | HMDB0240744 | Dimethyl maleate | 13 | C | 3 | **14** | **13** | | 15.0 µL | 144.1 |
| 182 | HMDB0031578 | 2-Methylpentanal | 13 | C | 4 | **10** | **12** | **Low water solubility** | 15.0 µL | 100.2 |
| 78 | HMDB0011626 | Dodecanol | 13 | C | 5 | | | **Poor water solubility** | 15.0 µL | 186.3 |
| 418 | HMDB0040201 | 3,3'-Dithiobis[2-methylfuran] | 13 | C | 6 | **11** | **64** | **Low water solubility** | 15.0 µL | 226.3 |
| 184 | HMDB0031327 | 2-Butoxyethanol | 13 | C | 7 | **12** | **0** | | 15.0 µL | 118.2 |
| 729 | HMDB0005994 | Hexanal | 13 | C | 8 | **10** | **9** | | 15.0 µL | 100.2 |

| | | | | | | | | | | |
|---|---|---|---|---|---|---|---|---|---|---|
| 721 | HMDB0031475 | Heptanal | 13 | C | 9 | **12** | **11** | | 15.0 µL | 114.2 |
| 93 | HMDB0001183 | Octanol | 13 | C | 10 | | | **Poor water solubility** | 15.0 µL | 130.2 |
| 105 | HMDB0003156 | 2,3-Butanediol | 13 | D | 1 | **10** | **0** | | 15.0 µL | 90.1 |
| 96 | HMDB0035243 | 1-Phenyl-1,2-propanedione | 13 | D | 2 | **14** | **35** | | 15.0 µL | 148.2 |
| 13 | PUBCHEMCID6431015 | (−)-α-Cedrene | 13 | D | 3 | | | **Poor water solubility** | 15.0 µL | 204.3 |
| 119 | HMDB0032233 | 2,5-Dimethyl-3(2H)-furanone | 13 | D | 4 | **12** | **10** | | 15.0 µL | 112.1 |
| 681 | HMDB0031094 | Glycerol tributanoate | 13 | D | 5 | | | **Poor water solubility** | 15.0 µL | 302.4 |
| 1235 | HMDB0032619 | 1-Phenylethanol | 13 | D | 6 | **13** | **13** | | 15.0 µL | 122.2 |
| 309 | HMDB0031602 | 4-Pentenoic acid | 13 | D | 7 | **11** | **10** | | 15.0 µL | 100.1 |
| 153 | HMDB0031221 | 2-Ethylbutanoic acid | 13 | D | 8 | **12** | **10** | | 15.0 µL | 116.2 |
| 175 | HMDB0005846 | Ethyl isopropyl ketone | 13 | D | 9 | **10** | **10** | **Low water solubility** | 15.0 µL | 100.2 |
| 539 | HMDB0004437 | Diethanolamine | 13 | D | 10 | **15** | **10** | | 11.7 | 141.6 |
| 192 | HMDB0012275 | Phenylethylamine | 13 | E | 1 | **13** | **10** | | 15.0 µL | 121.2 |
| 672 | HMDB0000131 | Glycerol | 13 | E | 2 | **10** | **0** | | 15.0 µL | 92.1 |
| 562 | HMDB0011743 | 2-Phenylpropionate | 13 | E | 3 | **11** | **34** | | 15.0 µL | 150.2 |
| 29 | HMDB0001893 | alpha-Tocopherol | 13 | E | 4 | | | **Poor water solubility** | 15.6 | 430.7 |
| 177 | HMDB0031594 | 2-Methylhexanoic acid | 13 | E | 5 | **14** | **11** | | 15.0 µL | 130.2 |
| 305 | HMDB0031557 | 4-Methyloctanoic acid | 13 | E | 6 | **14** | **30** | **Low water solubility** | 15.0 µL | 158.2 |
| 481 | HMDB0031407 | Cyclopentanone | 13 | E | 7 | **10** | **8** | **Soluble in acetone** | 15.0 µL | 84.1 |
| 633 | HMDB0034153 | Ethyl tetradecanoate | 13 | E | 8 | | | **Poor water solubility** | 15 | 256.4 |
| 870 | HMDB0002048 | m-Cresol | 13 | E | 9 | **11** | **8** | **Low water solubility** | 15.0 µL | 108.1 |
| 898 | HMDB0036240 | Methyl 4-tert-butylphenylaceta | 13 | E | 10 | **12** | **40** | | 15.0 µL | 206.3 |
| 357 | HMDB0011753 | | | | | | | | | |
| 1010 | HMDB0014444 | | | | | | | | | |
| 45 | HMDB0051629 | 1,2,3-Tri-13(Z)-Docosenoyl-rac-glycerol | | | | | | | | |
| 588 | PUBCHEMCID16129778 | Tannic acid | | | | | | | | |
| 642 | PUBCHEMCID5819661 | Glycerol tri-13(E)-docosenoate | | | | | | | | |
